# Insulin secretion deficits in a Prader-Willi syndrome β-cell model are associated with a concerted downregulation of multiple endoplasmic reticulum chaperones

**DOI:** 10.1101/2021.12.16.473032

**Authors:** Erik A. Koppes, Marie A. Johnson, James J. Moresco, Patrizia Luppi, Dale W. Lewis, Donna B. Stolz, Jolene K. Diedrich, John R. Yates, Ronald C. Wek, Simon C. Watkins, Susanne M. Gollin, Hyun J. Park, Peter Drain, Robert D. Nicholls

**Author notes:** Address correspondence to R.D.N.: Dr. Robert D. Nicholls, Department of Pediatrics, UPMC Children’s Hospital of Pittsburgh, Rangos Research Building, 4401 Penn Avenue, Pittsburgh, PA 15224. Center for Genetics of Host Defense, UT Southwestern Medical Center, Dallas, TX.

## Abstract

Prader-Willi syndrome (PWS) is a multisystem disorder with neurobehavioral, metabolic, and hormonal phenotypes, caused by loss of expression of a paternally-expressed imprinted gene cluster. Prior evidence from a PWS mouse model identified abnormal pancreatic islet development with retention of aged insulin and deficient insulin secretion. To determine the collective roles of PWS genes in β-cell biology, we used genome-editing to generate isogenic, clonal INS-1 insulinoma lines having 3.16 Mb deletions of the silent, maternal (control) or active, paternal PWS-alleles. PWS β-cells demonstrated a significant cell-autonomous reduction in basal and glucose-stimulated insulin secretion. Further, proteomic analyses revealed reduced levels of cellular and secreted hormones, including all insulin peptides and amylin, concomitant with reduction of at least ten endoplasmic reticulum (ER) chaperones, including GRP78 and GRP94. Critically, transcriptomic studies demonstrated that the broad reduction of ER chaperones originated from transcriptional downregulation without corresponding changes for *Ins1* and *Ins2*. In contrast to the dosage compensation previously seen for ER chaperones in *Grp78* or *Grp94* gene knockouts or knockdown, compensation is precluded by the stress-independent deficiency of ER chaperones in PWS β-cells. Consistent with reduced ER chaperones levels, PWS INS-1 β-cells are more sensitive to ER stress, leading to earlier activation of all three arms of the unfolded protein response. These findings suggest that a chronic shortage of ER chaperones in PWS β-cells leads to a deficiency of protein folding and/or delay in ER transit of insulin and other cargo. In summary, our results illuminate the pathophysiological basis of pancreatic β-cell hormone deficits in PWS, with evolutionary implications for the multigenic PWS-domain, and indicate that PWS-imprinted genes coordinate concerted regulation of ER chaperone biosynthesis and β-cell secretory pathway function.

## INTRODUCTION

Prader-Willi syndrome (PWS) is a multisystem disorder caused by loss of expression of a large contiguous cluster of paternally-expressed, imprinted genes from human chromosome 15q11.2 [1–3]. Clinically, PWS is characterized by failure to thrive with hypotonia, developmental and cognitive delay, behavioral problems, short stature, hypogonadism, hyperphagia, and early-onset obesity [3, 4]. Prominent endocrine features of PWS include deficiencies of multiple hormones, including growth hormone, oxytocin, gonadotropins, insulin-like growth factor, thyroid hormones, amylin/IAPP, and pancreatic polypeptide [1, 4–9]. In addition, plasma insulin is lower than expected in PWS relative to the degree of obesity [10–12]. Episodes of hypoglycemia have been reported in PWS patients, suggesting an imbalance in glucose homeostasis [13, 14]. In contrast, plasma ghrelin is grossly elevated in PWS [3, 10, 15], perhaps as a physiological response to glucose imbalance [16, 17]. Although the PWS literature suggests a hypothalamic etiology [3], the mechanisms for endocrine and metabolic dysfunction in PWS have not been elucidated.

The PWS-imprinted domain is comprised of ten paternally-expressed imprinted genes conserved in human and rodents (with an additional two genes each unique to human and rodents), encoding lncRNAs, snoRNAs, miRNAs, or distinct proteins [1, 2, 18–23]. These imprinted genes are predominantly expressed in neuronal [19, 21, 24] and neuroendocrine lineages [23], as well as pancreatic endocrine cells (α-, β-, δ-, and γ-cells, with only low expression in acinar, ductal, or undefined cells) based on recent gene-specific [25] and single cell RNA sequencing studies [26–28]. Further, *SNRPN* has decreased islet expression in β-cell failure due to saturated fatty acids [29] or cytokines [30], while *SNORD116* and *SNORD107* occur in islet exosomes and decrease by IL-1β and IFN-γ treatment [31]. Finally, *SNORD116* RNA levels are greatly reduced in MODY3 β-cells [32]. These observations suggest that PWS-gene products function in the endocrine pancreas including in β-cells.

Our earlier studies of a transgenic-PWS (TgPWS) mouse model harboring a deletion of the orthologous PWS-imprinted domain demonstrated severe failure to thrive, abnormalities in fetal pancreatic islet development and architecture, reduced α- and β-cell mass, increased apoptosis, and postnatal onset of progressive hypoglycemia that led to lethality within the first postnatal week [16, 25]. Plasma insulin and glucagon levels were low during fetal and neonatal life of TgPWS mice [16, 25], and, at postnatal day 1 prior to onset of hypoglycemia, there was significantly reduced basal and glucose-stimulated insulin secretion (GSIS) from cultured TgPWS islets [25]. Furthermore, using an insulin-Timer fluorescent protein biomarker to image postnatal day 1 β-cells *in vivo*, TgPWS mice showed a striking accumulation of aged insulin whereas wildtype control littermates only displayed newly synthesized insulin [25]. These results suggested that PWS-imprinted genes are required for the development and secretory function of pancreatic endocrine cells.

Based on the salient insulin secretion dysfunction in the PWS-mouse model [25], we sought to assess the role of PWS genes in pancreatic β-cell secretory pathway function in a cellular model system. Herein, we used CRISPR/Cas9 genome editing within the INS-1(832/13) insulinoma (β)-cell line [33] to generate deletions of the complete PWS-domain. Following validation of a cell-autonomous insulin secretion deficit in the PWS INS-1 model, we performed molecular profiling by proteomic and transcriptomic approaches. The data revealed deficits in multiple secreted hormones as well as endoplasmic reticulum (ER) chaperones that are components of secretory protein folding and trafficking pathways [34–39], providing mechanistic insight into PWS-gene function in pancreatic β-cells.

## RESULTS

### Generation of INS-1 cell lines with silent, maternal allele PWS-gene deletions

Various insulinoma cell lines, predominantly of rodent origin, are widely utilized to investigate β-cell mechanisms as a tractable *in vitro* model system [40]. To investigate the functions of PWS-genes in β-cells, we used CRISPR/Cas9 genome-editing to target 3.16 Mb deletions encompassing the PWS-genes (**Fig. 1A**; **Fig. S1A**) in rat INS-1(832/13)::mCherry insulinoma cells (hereafter termed INS-1) that secrete rat and human insulin [33] and express a mouse *Ins2* C-peptide-mCherry biosensor in insulin secretory granules [41]. The large deletions were visualized in about 9% of unselected, transfected cells as determined by fluorescence *in situ* hybridization (FISH) (**Fig. S1B**). INS-1 lines with PWS-domain deletions were derived through sequential targeting and clonal isolation of cell lines initially harboring deletions of the silent maternal allele, followed by targeting of the remaining paternal loci, culminating in the creation of homozygous PWS-deletion lines (**Fig. 1B**).

**Figure 1.**
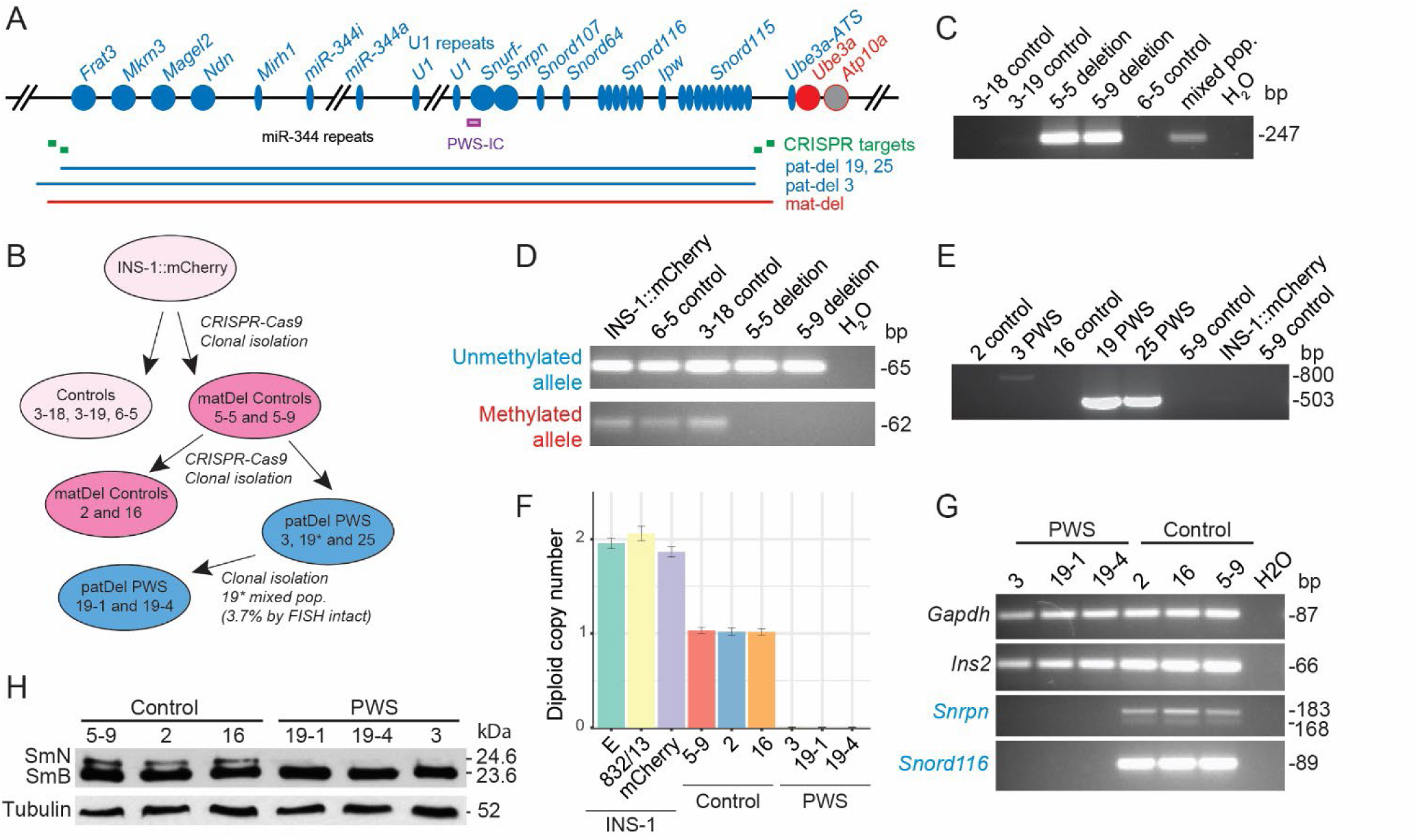
CRISPR/Cas9 genome editing to generate INS-1 lines with 3.16 Mb deletions of the silent (maternal) and active (paternal) PWS-imprinted domain. **(A)** Gene map of the rat PWS-imprinted domain and CRISPR/Cas9-targeted deletions. Symbols: circles, protein-coding genes; thin ovals, RNA genes; blue, (paternal; pat) and red (maternal; mat), imprinted genes; IC, imprinting control region (purple bar); green boxes, CRISPR gRNA target sites; blue and red horizontal bars, extent of deletions. Not all copies of tandemly repeated loci (*miR-344*; U1, *Snord116*, *Snord115*) are shown. **(B)** Schematic showing generation of genome edited, clonal INS-1 lines with 3.16 Mb deletions on the silent maternal allele only (5-9, 2, 16; dark pink) or homozygous deletion of both maternal and active paternal alleles (3, 19-1, 19-4, 25; blue). **(C)** First round deletion-PCR on genomic DNA identifies two INS-1 lines with deletions (5-9, 5-5). **(D)** DNA methylation analysis of bisulfite-modified genomic DNA at the PWS-IC establishes that lines 5-5 and 5-9 have deletions on the maternal allele. **(E)** Second round deletion-PCR identifies three INS-1 lines (3, 19, 25) with deletions on the paternal allele, with line 3 having an alternate proximal breakpoint. **(F)** Gene dosage of *Snord107* normalized to *Ube3a* as determined by ddPCR using genomic DNA from the INS-1 panel of cell lines (also see **Fig. S9**). **(G)** PWS-deletion lines (3, 19-1, 19-4) lack mRNA expression of the PWS-imprinted genes, *Snrpn* and *Snord116*, as detected by RT-PCR, whereas control lines (5-9, 2, 16) express PWS-imprinted genes. All cell lines express the control genes (*Gapdh*, *Ins2*). The full panel of PWS-imprinted genes is shown in **Fig. S10A,B**. **(H)** Use of whole cell extracts for a western blot shows that PWS-deletion lines (3, 19-1, 19-4) lack expression of the spliceosomal SmN polypeptide (24.6 kDa) encoded by *Snrpn*, but retain expression of the paralogous SmB polypeptide (23.6 kDa) encoded by the unlinked *Snrpb* gene. The control is α-Tubulin.

This process first led to isolation of two lines with a PWS-region deletion (5-5, 5-9) identified by deletion-breakpoint PCR (**Fig. 1C**) and DNA sequencing (**Fig. S2A,B**), and confirmed by FISH using BAC probes from within and outside the PWS-domain (**Fig. S1C-F**). One PWS allele was deleted in all interphase and metaphase cells of line 5-5 (**Fig. S1D**), indicating clonal isolation of a cell line with a PWS-domain deletion. Initially, line 5-9 was more complex, as FISH showed two cell types with either no deletion (most cells) or a PWS-domain deletion (**Fig. S1E**); additionally, similar proportions of cells lacked mCherry fluorescence, likely due to epigenetic silencing (as DNA analysis indicated the transgene was present) or were mCherry-positive, respectively. This allowed separation by fluorescence-activated cell sorting (FACS), with establishment of a clonal mCherry-positive cell line (5-9) harboring a PWS-domain deletion (**Fig. S1F**). Clonal engineered cell lines 5-5 and 5-9 were inferred to have maternal-deletions of the rat PWS-domain as evidenced by **1)** detection of only an unmethylated allele at the PWS-imprinting center (PWS-IC) by bisulfite PCR whereas parental INS-1 and control lines had biparental DNA methylation (**Fig. 1D**); and **2)** expression of PWS-imprinted loci by reverse-transcription-PCR (RT-PCR) (**Fig. S1G**), indicating the presence of an active, paternal-allele (**Fig. S1H**). Finally, the presence of mutations at sgRNA target sites on the intact, paternal chromosome, termed scarred mutation alleles, identified a single nucleotide insertion at the sgRNA1 site (**Fig. S1I**; **Fig. S2C-E**) but different nucleotide insertions at the sgRNA3 site (**Fig. S1I**; **Fig. S2F-H**), the latter indicating an independent origin for the cell lines, 5-5 and 5-9. In contrast, no sequence changes occurred at top-ranked off-target sites for sgRNA1 or sgRNA3 (**Fig. S3**).

### Generation of PWS INS-1 cell lines with deletions on the active paternal allele

To specifically target the paternal allele of lines 5-5 and 5-9, we used rat-specific sgRNA sets internal to the first targeting sites to generate slightly smaller 3.143 Mb PWS-deletions (**Fig. S4A-C**). The use of FISH demonstrated that about 5% of unselected, transfected cells had a PWS-deletion (**Fig. S4D**). We chose maternal deletion (control) line 5-9 for a second round of transfections and clonal isolation, generating control lines 2 and 16 (**Fig. 1B,E**; **Fig. S4E,F**) as well as PWS lines 19 and 25, with an expected paternal deletion-PCR product and PWS line 3 with a larger than expected deletion-PCR product (**Fig. 1B,E**; **Fig. S4G**). DNA sequencing confirmed PWS-deletion breakpoints for lines 19 and 25 (**Fig. S5A-C**) as well as for line 3, although the latter arose from a larger deletion at an alternate (alt) proximal alt-sgRNA70-3 targeting site with a DNA repair event that inserted a fragment of *E. coli* DNA in the breakpoint junction (**Fig. S5D**). FISH studies confirmed that lines 3 and 25 had deletions of the PWS-domain on each allele (**Fig. S4H,I**); however, FISH analysis also revealed that line 25 was mosaic for cells with diploid or tetraploid PWS-signals (**Fig. S4I**). Consequently, line 25 was not used further. In contrast, line 19 had a residual fraction of cells (1-3.7%) with an intact paternal allele as shown by FISH (**Fig. S4J**) and droplet digital PCR (ddPCR) (**Fig. S6**) which was removed by cell dilution and isolation of five clonal lines 19-1 through 19-5 (**Fig. 1B**; **Fig. S4K,L**; **Fig. S10A**). Finally, sgRNA target sites not deleted or involved in a deletion breakpoint were assessed for scarred mutation alleles identifying a 28-nt deletion at alt-sgRNA70-3 in PWS lines 19-1 and 19-4 (**Fig. S4N**; **Fig. S5E-L**), while no sequence changes occurred at top-ranked off-target sites for sgRNA70-3 (**Fig. S7**) or sgRNA79-1 (**Fig. S8**).

Genomic copy number was confirmed by ddPCR (**Fig. 1F**; **Fig. S6**; **Fig. S9**), establishing a set of three control lines (5-9, 2, 16) with hemizygosity for maternal-allele deletions (**Fig. S1H**) and three PWS INS-1 lines (3, 19-1, 19-4) with homozygosity for PWS-domain deletions (**Fig. S4M**). An absence of PWS-gene expression in all PWS lines was shown by RT-PCR for each PWS-imprinted gene (**Fig. 1G**; **Fig. S10A,B**) and by western blot analysis for the SmN polypeptide encoded by *Snrpn* (**Fig. 1H**). Intriguingly, PWS INS-1 lines showed a very low level of apparent expression of the PWS-imprinted gene *Snurf* by RT-PCR (**Fig. S10A**; **Fig. S11A,B**). Sequencing of the RT-PCR products identified an expressed *ψSnurf* sequence within a recently evolved *ψSnurf*-*ψSnrpn* locus located in an intron of the *Mon2* gene (**Fig. S11B,C**). Using a *Pml*I variant between *Snurf* and *ψSnurf* (**Fig. S11D**), we confirmed expression specifically of *ψSnurf* with complete silencing of *Snurf* (**Fig. S11B**), as expected for PWS INS-1 lines. Scattered mutations in the *ψSnurf* (**Fig. S11D**) and *ψSnrpn* (**Fig. S11E,F**) segments likely inactivate any coding potential (**Fig. S11G, H**). In summation, these results establish a panel of clonal INS-1 lines homozygous for ∼ 3.16 Mb deletions, also termed PWS β-cell lines, and similarly, of clonal control lines with a deletion only involving the silent, maternal allele.

### PWS INS-1 cell lines show deficits in insulin secretion and ER chaperones

To determine whether PWS INS-1 cell lines have secretory deficits, we carried out insulin secretion assays under low (2.2 mM) and high (22 mM) glucose conditions. The three PWS cell lines displayed a striking deficit in both basal and glucose-induced insulin secretion as compared to the isogenic control INS-1 lines (**Fig. 2**). However, both the PWS and control cell lines had a similar increase from each basal level for GSIS (**Fig. 2**), indicating that PWS β-cells were not deficient in glucose responsiveness.

**Figure 2.**
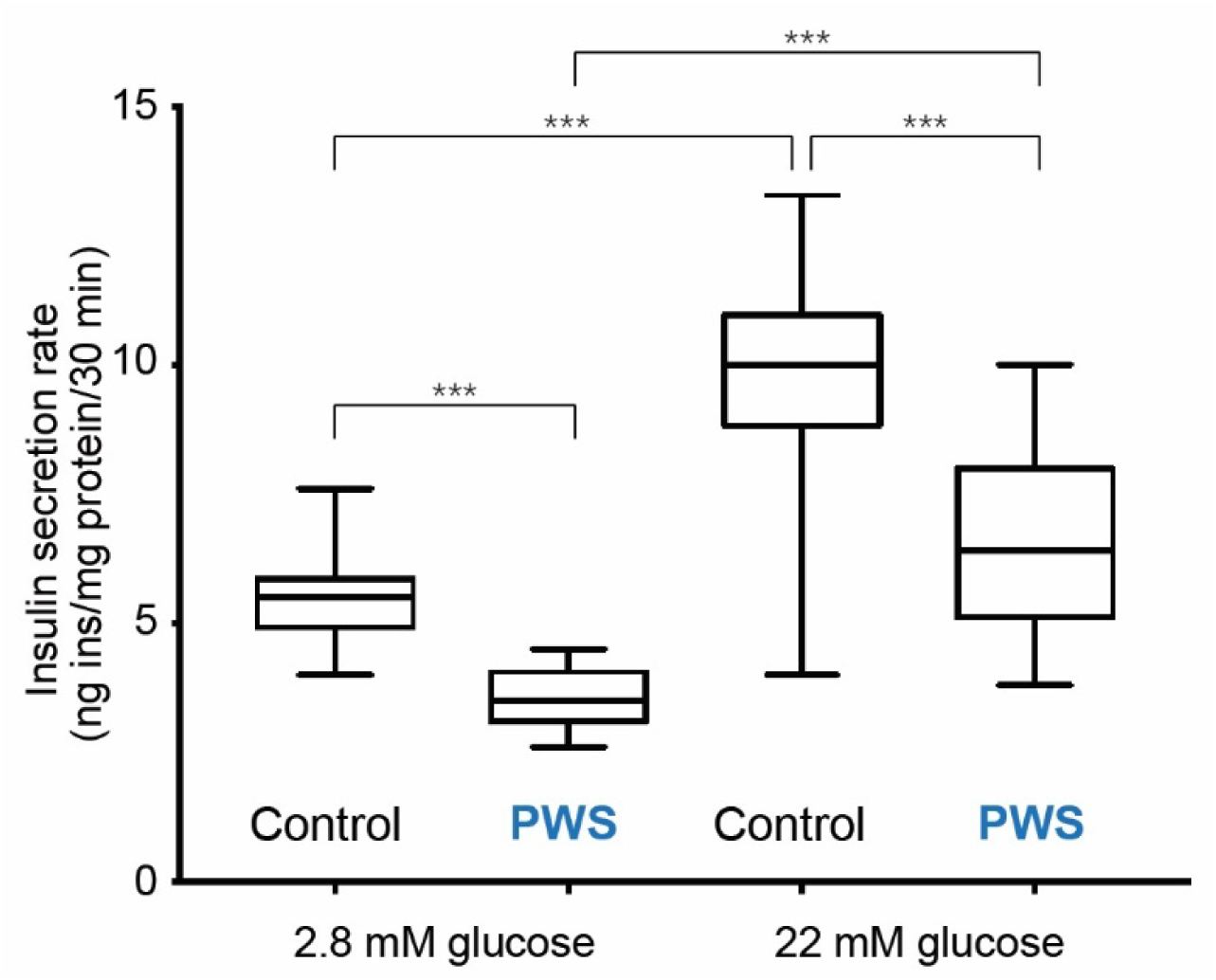
Deficient basal and GSIS in PWS-deletion INS-1 lines. PWS INS-1 cell lines have insulin secretion deficits at both low (2.8 mM) and high (22 mM) glucose, based on the pooled insulin secretory rates for each of PWS (3, 19-1, 19-4) and control (5-9, 2, 16) groups (n=36 biological replicates per group, with n=12 per cell line). Although GSIS increased 1.75-fold for control and 1.85-fold for PWS, loss of the PWS genes decreased the secretory rate by 36% in basal 2.8 mM glucose conditions and by 32% in stimulatory 22 mM glucose conditions. Statistical comparison by ANOVA with Tukey post hoc test (***, *P* < 0.0001).

Next, mass spectrometry was used to assess changes in the cellular proteome and all secreted peptides in PWS *vs.* control INS-1 cell lines, under insulin secretion conditions of 22 mM glucose, separating cellular proteins by methanol-acetic acid extraction into soluble (mostly small) and insoluble (mostly large) protein fractions with the latter analyzed by quantitative proteomics (**Fig. 3A**). Unexpectedly, in PWS INS-1 cells there was a striking deficiency of multiple ER chaperone proteins, including GRP78/BiP (HSPA5), GRP94/endoplasmin (HSP90B1), PDIA4, HYOU1, CRELD2, and DNAJB11, with lesser reductions in SDF2L1, DNAJC3, PDIA6, and PDIA3, and a modest decrease in PPIB (**Fig. 3B**). Similar deficits of residual amounts of many of these ER chaperones as well as MANF were detected in the soluble protein fraction (**Fig. 3C**). Furthermore, in PWS β-cell lines there was reduced abundance of numerous hormones co-secreted from INS-1 cells, including rat insulin-1 and insulin-2, mouse insulin-2, and human insulin (pre-pro-, pro-, C-peptide, and mature versions of each), as well as reductions in processed and precursor forms of IAPP and NPY, detected in both the cellular small protein fraction (**Fig. 3C**) and residual amounts in the large protein fraction by quantitative proteomics (**Fig. 3B**). In contrast, the full-length secretory granule protein, chromogranin B (CHGB), and the C-terminal CHGB (CCB)-peptide were increased 2-fold (**Fig. 3B,C**). Extending both the insulin secretion data (**Fig. 2**) and the cellular protein deficiencies (**Fig. 3B,C**), mass spectrometry analysis of peptides secreted into the culture media demonstrated reduced levels of all forms of insulins and IAPP in PWS β-cell media compared to control (**Fig. 3D**). There was also a reduction in secreted levels of the 14 amino acid WE14 peptide processed from chromogranin A (CHGA), but no changes in the CHGA precursor and processed forms in either secreted or cellular fractions. Secreted levels of chromogranin B were increased in PWS (**Fig. 3D**), further illustrating the concordance between cellular and secreted peptide levels in PWS *vs.* control INS-1 lines. These results indicate that deletion of PWS-genes sharply lowers secretion of insulin and other peptide hormones (IAPP, NPY, CHGA-WE14) and this reduction is associated with deficiencies in many ER chaperones.

**Figure 3.**
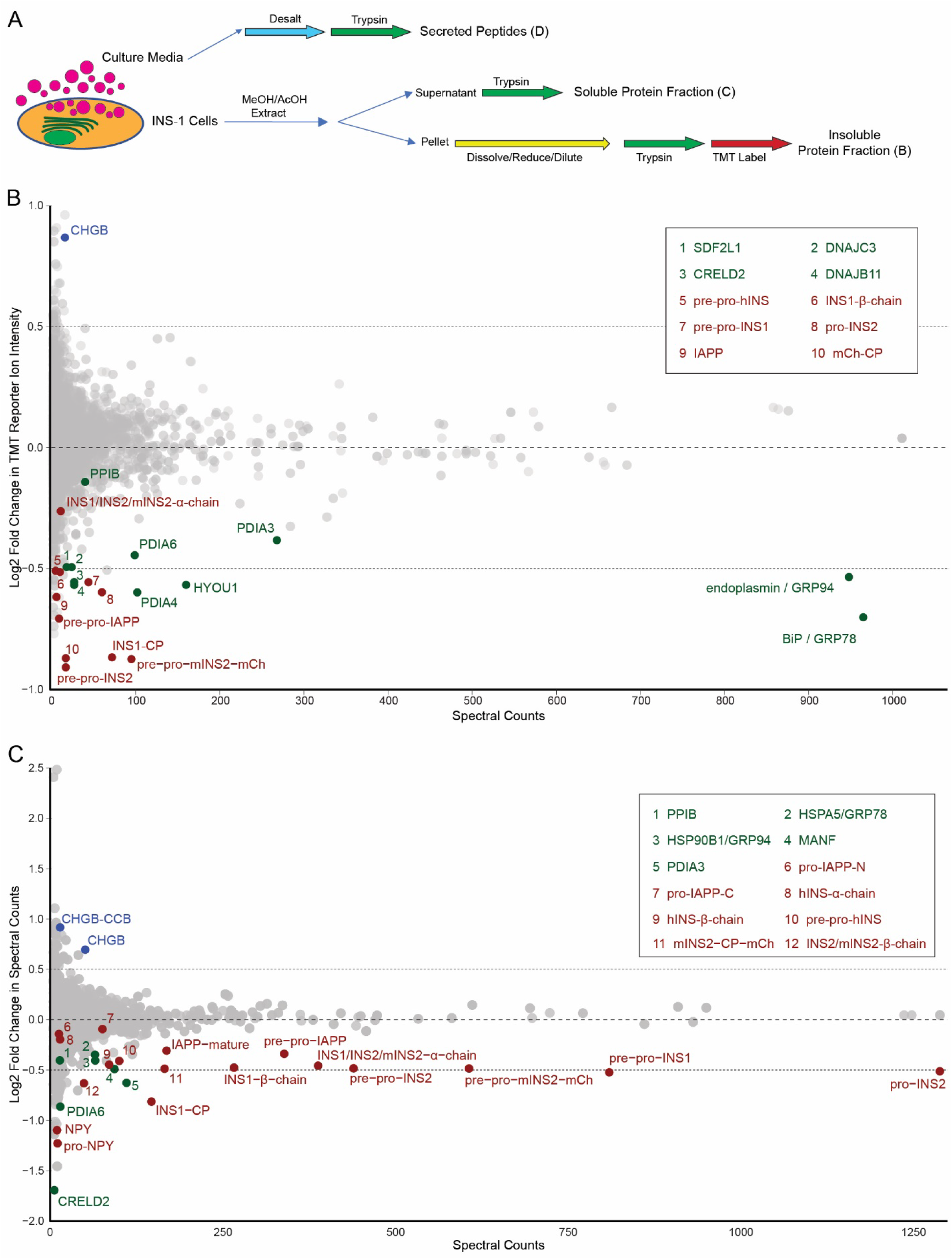
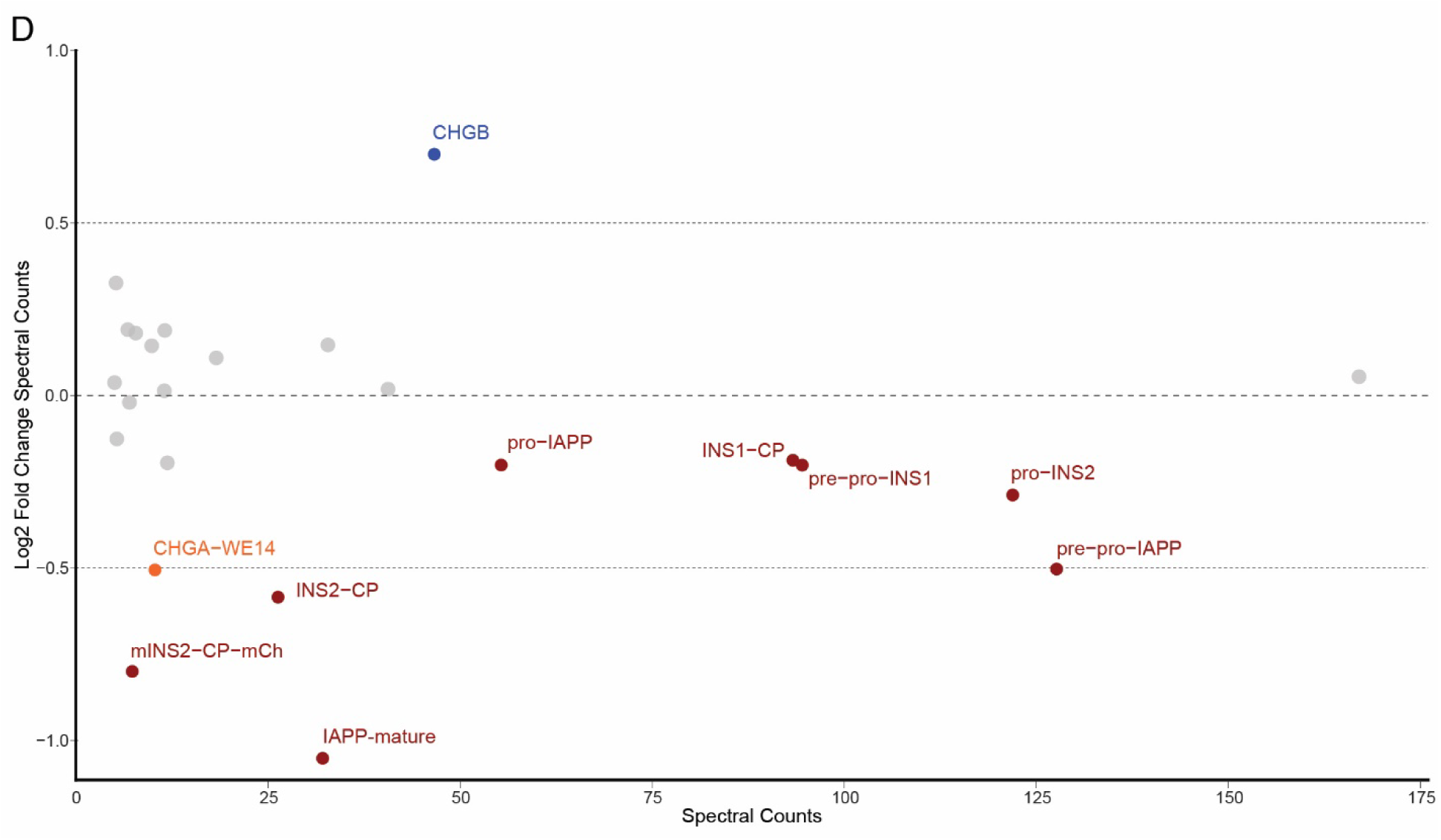
Proteome-wide alterations in PWS-deletion *vs.* control INS-1 lines identifies reductions in levels of ER chaperones and hormones. **(A)** Multiple fractions of PWS and control INS-1 cell cultures grown under GSIS conditions were assessed by mass spectrometry (MS). These included peptide hormones released from secretory granules (pink circles) into the media [see Fig. 3D] and cellular proteins methanol-acetic acid extracted into soluble proteins [see Fig. 3C], or an insoluble protein fraction assessed by quantitative Tandem Mass Tag (TMT) MS [see Fig. 3B]. **(B-D)** Relative protein detection of proteins in PWS of Log2 Fold Change (PWS/Control) of protein detection plotted against spectral counts indicating overall protein abundance. **(B)** Relative comparison of insoluble cellular proteins in PWS and control INS-1 β-cell lines detected by quantitative TMT MS. Protein levels of eleven ER chaperones (green) as well as secreted hormones (red) insulin [various processed forms of INS1, INS2, mouse (m) mINS2-mCherry (mCh), human (h) INS, and C-peptide (CP); mature INS is the 21 amino acid peptide identical between rat INS1, rat INS2 and mouse INS2] and amylin (full length and processed IAPP) were markedly reduced in PWS INS-1 cell lines. In contrast, chromogranin B (CHGB) levels are noticeably increased in PWS INS-1 cell lines (blue). The boxed Key indicates the identity of the 10 numbered peptide spots. **(C)** Comparison of soluble cellular peptides detected in PWS *vs.* control INS-1 lines, with reductions observed in PWS lines for highly expressed peptide hormones (red) including insulins [various processed forms of INS1, INS2, mINS2-mCh and CP], amylin (processed and unprocessed IAPP), and neuropeptide Y (processed and unprocessed NPY), as well as lower levels in the soluble fraction of the ER chaperones (green). Additionally, two isoforms of chromogranin B (full-length CHGB and the CCB C-terminal fragment of chromogranin B) are increased in PWS INS-1 cell lines (blue). The boxed Key indicates the identity of the 12 numbered peptide spots. **(D)** Comparison of secreted peptides highlighting reduced levels for PWS relative to control INS-1 lines for numerous secreted hormones (red) including insulins [various processed forms of INS1, INS2, mINS2-mCh and CP] and amylin (processed and unprocessed IAPP), as well as reduced secreted levels from PWS INS-1 lines of the CHGA-derived WE14 peptide (orange), while secreted levels of CHGB (blue) were increased in PWS INS-1 cell lines.

### Reduced mRNA levels for hormones and ER chaperones in PWS INS-1 cell lines

Many of the PWS-imprinted genes (**Fig. 1A**) are suggested to function in gene expression [18, 19, 21, 22, 42, 43]. To gain insight into the molecular basis for insulin secretion and ER chaperone deficits in PWS INS-1 lines, high-throughput total RNA sequencing (RNA-seq) was used to identify differentially expressed genes (DEGs) (**Fig. 4A-C**; **Tables S1,S2**). Visualization by a heatmap clustergram (**Fig. 4A**) indicates that PWS clonal lines and control clonal lines each grouped together with similar expression profiles. A volcano plot depicting the magnitude, direction, and statistical significance of DEGs in the 3 PWS *vs.* 3 control lines illustrates no appreciable expression (Log2 Fold Change < -5) of PWS-imprinted genes with remaining DEGs symmetrically orientated around the ordinate axis with a near equivalent number of significantly upregulated (105) and downregulated (123) genes (**Fig. 4B**). The PWS transcripts with complete loss of expression specifically in PWS INS-1 lines include *Snurf*, *Snrpn*, *Ipw*, *Mkrn3*, all four snoRNAs (*Snord116, Snord115, Snord107* and Snord*109*), miRNA (*Mir344* isoforms), and duplicated U1-*Snurf* sequences (**Fig. 4B**), as also seen by RT-PCR (**Fig. 1G**; **Fig. S10A,B**; **Fig. S11A,B**). Three PWS-imprinted genes, *Ndn*, *Magel2*, and *Frat3* (*Peg12*), were not detected as DEGs as these are not expressed by RT-PCR or RNA-sequencing in any of parental, control, or PWS INS-1 lines. These loci are present by genomic PCR, suggesting an epigenetic inactivation in the INS-1 founder cell line, although silencing is not widespread, as *Mkrn3* and *Mir-344* are interspersed with the silenced loci (see **Fig. 1A**) and are well-expressed in control INS-1 lines (**Fig. S10B**; **Fig. S12**).

**Figure 4.**
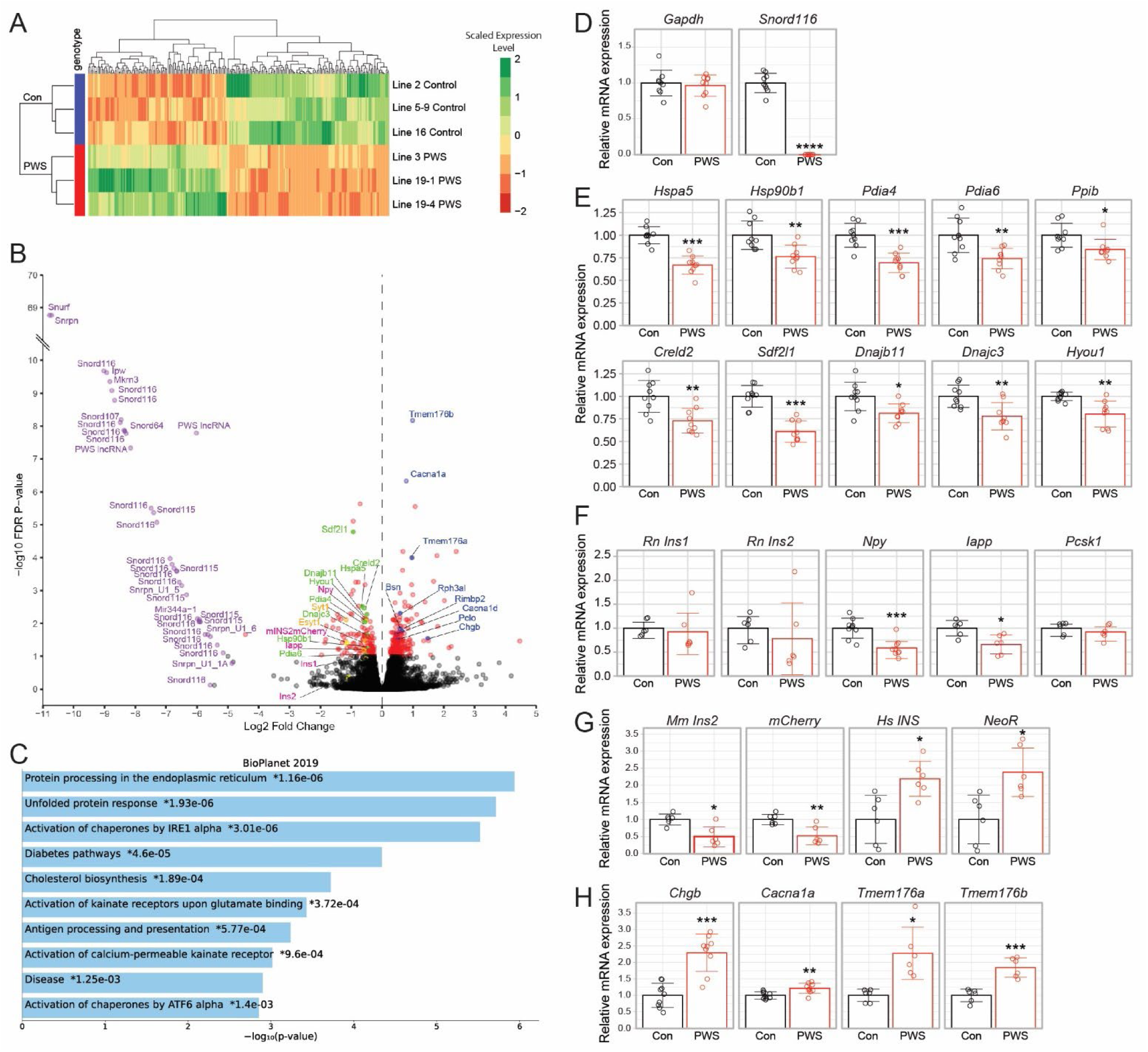
Genome-wide transcriptome alterations in PWS-deletion *vs.* control INS-1 lines identifies significant differentially expressed genes (DEGs) including those encoding ER chaperones and hormones. **(A)** Heatmap clustergram of 228 DEGs demonstrates tight clustering of PWS *vs.* control groups (*Padj* < 0.05). Scale: green (enriched) to red (depleted). RNA-seq was performed for 3 PWS (3, 19-1, 19-4) *vs.* 3 control (5-9, 2, 16) INS-1 cell lines. **(B)** Volcano plot showing statistical significance (-log10 FDR *P*-value) *vs.* magnitude of change (Log2 Fold Change) for all expressed genes. Colored data points indicate PWS-imprinted genes (purple), ER chaperones (green), secreted proteins (pink), synaptotagmins involved in insulin secretion (yellow), proteins involved in neuronal active zone/exocytosis and vesicle acidification (blue), and all other significant genes (red, without labels). **(C)** Gene set enrichment analysis identifies key biological pathways involved in protein processing and the unfolded stress response in the ER. **(D-H)** Relative gene expression of control (Con, black; 5-9, 2, 16) *vs.* PWS (red; 3, 19-1, 19-4) determined by RT-ddPCR normalized to *Gpi* levels and to the average expression in control INS-1 lines, including for **(D)** *Gapdh* and *Snord116* (PWS) control genes, **(E)** ten downregulated DEGs encoding ER chaperones, **(F)** endogenous *Rattus norvegicus* (Rn) genes encoding β-cell hormones and secretory proteins, **(G)** exogenous insulin transgenes in the INS-1 lines (*Mm*, *Mus musculus*; *Hs*, *Homo sapiens*) as well as mCherry and neomycin resistance (*Neo*R) marker proteins, and **(H)** upregulated DEGs involved in the secretory pathway. Statistical comparison by Welch’s t-test: *, *P* < 0.05; **, *P* < 0.005; ***, *P* < 0.0005; ****, *P* < 0.00005.

To ensure complete coverage of the INS-1 transcriptome, RNA-seq of small stable RNAs in the PWS *vs.* control cell lines was carried out, culminating in the identification of 58 significant differentially expressed miRNAs and snoRNAs (**Fig. S12A,B**; **Table S3**). Congruent with the RNA-seq data, all but three correspond to PWS-imprinted small RNAs, including duplicated *MiR-344*, *Snord116* and *Snord115* loci, as well as single copy *Snord107* and *Snord64* loci (**Fig. S12A,B**). In addition, *Mir135b*, *Mir3065*, and *Mir212* were downregulated in PWS INS-1 lines (**Fig. S12B**), although predicted targets were not DEGs by RNA-seq. Eleven of the top 25 highly expressed miRNAs in the rat INS-1 β-cell model (**Table S4**), including *Mir375*, *Mir148a*, *Mir183*, *Mir30d*, *Mir27b*, *Mir25*, *Mir26a2*, *Mir26a*, *Mir192*, *Mir125a*, and *Mir141* are also in the top 25 expressed miRNAs for a human β-cell model [44]. Within INS-1 cells, expression of PWS-region *Snord116* copies make up 13.8% of the top 188 highly expressed snoRNAs and cumulatively would be the thirteenth highest expressed, with *Snord64* also in and *Snord107* just outside the top 100 (**Table S5**), although snoRNAs are poorly studied as a small RNA class.

In addition to the loss of PWS-imprinted gene expression in the PWS INS-1 lines, four additional classes of DEGs were identified by manual annotation analysis of RNA-seq data (**Fig. 4B**). These DEG classes encoded: **1)** hormones (e.g., *Iapp*, *Npy*, *mIns2::mCherry*) that were significantly reduced; **2)** nine ER chaperones that were lowered including *Sdf2l1*, *Hspa5*, *Creld2*, *Dnajb11*, *Hyou1*, *Pdia4*, *Dnajc2*, *Hsp90b1*, and *Pdia6*; **3)** “neuronal active zone” proteins that had increased expression, including *Cacna1a*, *Rph3al*, *Bsn*, *Rimbp2*, *Cacna1d*, *Pclo*, and *Chgb*, many of which also play a role in insulin exocytosis [45–50]; and **4)** two related transmembrane proteins involved in vesicle acidification (*Tmem176a,b*) [51] whose expression were also enhanced. Gene ontology and pathway analysis of downregulated genes (excluding PWS-imprinted genes) through Enrichr or DAVID highlighted enrichment of multiple ER functional groups including those in ER protein processing, and the unfolded protein response (UPR) that are linked with both IRE1 and ATF6 (**Fig. 4C**; **Fig. S13A-C**). Finally, based on the input of downregulated DEGs, Enrichr predicted potential upstream transcriptional regulatory factors, including NFYA/NFYB (a cofactor of ATF6α, hereafter ATF6), CPEB1, RFX5, IRF3, CREB1, SREBF1, XBP1, and PPARB/D (**Fig. S14**).

Transcriptional changes observed by RNA-seq were corroborated by RT-droplet digital PCR (RT-ddPCR) analyses using RNA from independent biological replicates of each PWS and control INS-1 line (**Fig. 4D-H**; **Fig. S15**; **Fig. S16**). We validated equal expression of the housekeeping gene, *Gapdh*, and loss of expression of *Snord116* (**Fig. 4D**), as well as significant down-regulation in PWS INS-1 lines of 10 genes encoding ER chaperones (**Fig. 4E**: *Hspa5*, *Hsp90b1*, *Pdia4*, *Pdia6*, *Ppib*, *Creld2*, *Sdf2l1*, *Dnajb11*, *Dnajc3*, and *Hyou1*), of endogenous hormones (**Fig. 4F**: *Npy*, *Iapp*), and of the mouse *Ins2*-*mCherry* transgene (**Fig. 4G**). Further, another eight downregulated genes in PWS β-cells were verified (**Fig. S16**) including *Syt1*, encoding a Ca^2+^-sensor [52] and *Jph3*, encoding a junctophilin involved in ER-plasma membrane contact bridges [53]; each with roles in insulin secretion [52, 53]; *Derl3*, involved in ER-associated degradation (ERAD) [54], *Mylip*, encoding an E3 ubiquitin ligase regulating the LDL receptor [55], *Atp10a*, encoding a lipid flippase [56], and *Tap1* and *Tap2*, encoding ER antigenic peptide transporters that play a role in type 1 diabetes [57]. Additionally, up-regulation of four genes was confirmed by RT-ddPCR (**Fig. 4H**), including *Cacna1a*, *Chgb*, *Tmem176a*, and *Tmem176b*.

Several other observations merit note. In contrast to the proteomics data wherein all endogenous forms of insulin were decreased in PWS INS-1 lines, neither rat *Ins1* nor *Ins2* mRNA levels were statistically different between PWS and control lines in the RNA-seq or RT-ddPCR (**Fig. 4F**) expression profiling. Only a minority of DEGs identified by RNA-seq did not validate by RT-ddPCR, including several encoding neuronal active zone proteins (**Fig. S16**), possibly arising from cell culture media differences between biological replicates. Additionally, an apparent increase in expression of the human *INS*-*neoR* transgene in PWS lines (**Fig. 4G**) was an artifactual consequence of epigenetic silencing of the transgene, specifically in control line 16 (**Fig. S15A,I-K,M**). Interestingly, although both mRNA (*Pcsk1*) and protein levels of prohormone convertase PC1 were reduced in iPSC-derived neurons from PWS patients and in whole islets of inbred *Snord116*-deficient mice [23, 58], neither PC1 protein nor *Pcsk1* mRNA levels were changed in PWS INS-1 cell lines (**Fig. 4F**). Finally, no markers of activation of apoptotic or other cell death pathways were observed by RNA-seq or proteomics, consistent with the absence of any increase in cell death observed in cultured PWS or control INS-1 cells. Indeed, electron microscopy showed normal mitochondria, rough ER, and other subcellular organelles in PWS and control INS-1 lines (**Fig. S17**). The difference in observations *in vivo* where increased apoptosis was observed in a subset of α- and β-cells in TgPWS fetal islets [25] and *in vitro* (this study) may reflect the use of enriched culture medium with reduced cellular stress under cell culture conditions. In summary, the transcriptome studies show that loss of PWS-gene expression in PWS β-cell lines is accompanied by widespread alterations in mRNA levels most notably encoding secreted peptides and ER chaperones.

### Confirmation of insulin and ER chaperone protein deficits in PWS INS-1 lines

To further address the predicted disruptions of the ER and secretory pathway, we used western blot analyses to measure cellular levels of insulin and ER chaperones in control and PWS INS-1 lines. Levels of numerous forms of insulin polypeptides detected by an anti-insulin antibody were each significantly decreased in PWS INS-1 lines compared to controls, including diminished amounts of pro-insulin, pre-pro-insulin, and pro-mInsulin-2::mCherry bands (**Fig. 5A,D**). Additionally, using an anti-mCherry antibody, we observed that PWS cells have significantly lower levels of the proinsulin form of the mouse insulin2-mCherry transgene but no decrease in the processed C-peptide (CP) form (**Fig. 5B,E**). Indeed, confocal microscopy of the processed CP-mCherry shows punctate cytoplasmic compartmentalization without a discernable difference between PWS and control INS-1 lines (**Fig. S18**). Importantly, use of a KDEL antibody to identify proteins with the ER retention motif confirmed significant deficiencies in levels of the two major ER chaperone proteins, GRP78/BiP and GRP94, in PWS INS-1 cell lines (**Fig. 5C,F**). These results indicate that there are major disruptions of the protein folding machinery and attendant reductions in insulin processing and secretion in the PWS INS-1 cells.

**Figure 5.**
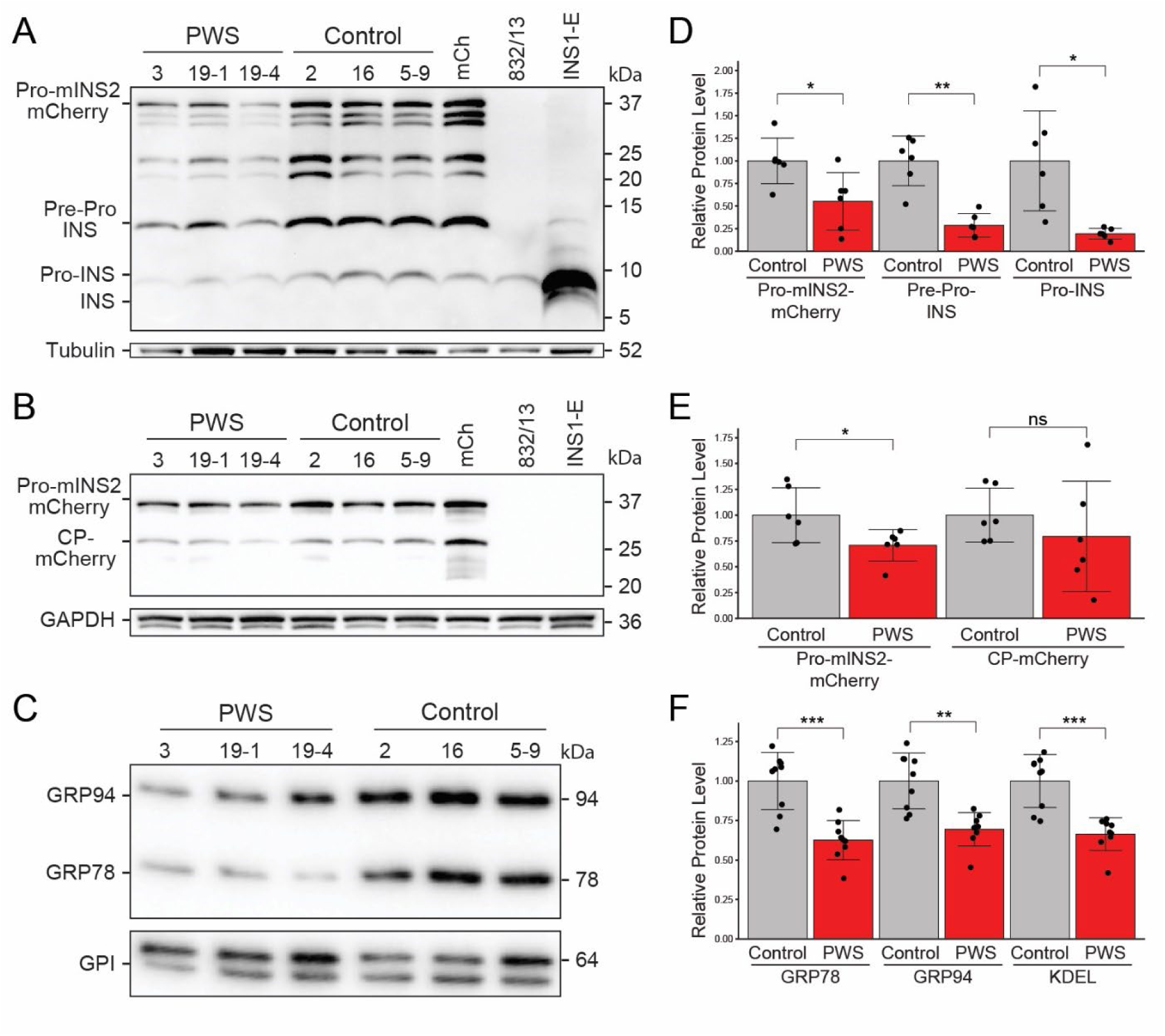
Reductions in insulins and ER chaperone levels in PWS-deletion *vs.* control INS-1 lines. **(A-C)** Western blots of whole cell lysates from PWS (3, 19-1, 19-4) and control (5-9, 2, 16) INS-1 β-cell lines grown under control (DMSO) conditions for 5 h were assessed using a panel of antibodies. **(A)** Anti-insulin, detecting all cellular forms of insulin (Pre-Pro-, Pro-, and fully processed rat INS) as well as mouse proinsulin2 (Pro-mINS2)-mCherry. **(B)** Anti-mCherry, detecting mouse proinsulin2 (Pro-mINS2)-mCherry and C-peptide (CP)-mCherry. **(C)** Anti-KDEL, detecting the two major ER chaperones GRP94 (endoplasmin; HSP90B1) and GRP78 (BiP; HSPA5). Anti-α-Tubulin, anti-GAPDH, and anti-GPI were used as controls for protein loading levels in **(A)**, **(B)**, and **(C)**, respectively. For **(A)** and **(B)**, control cell lines included INS-1::mCherry (mCh), INS-1(832/13) parental, and INS1-E. **(D)** Quantitation of Pro-mINS2-mCherry, Pre-Pro-INS and Pro-INS detected with anti-insulin in the PWS and control INS-1 lines (n=6 each genotype). **(E)** Quantitation of Pro-mINS2-mCherry and CP-mCherry detected with anti-mCherry in the PWS and control INS-1 lines (n=6 each). **(F)** Quantitation of GRP78, GRP94, and total KDEL detected with anti-KDEL in the PWS and control INS-1 lines (n=9 each). For **(D-F)**, statistical comparison by Welch’s t-test: *, *P* < 0.05; **, *P* < 0.005; ***, *P* < 0.0005; ns, not significant.

### PWS INS-1 cells are more sensitive to activation by ER stressors

Chaperones, such as GRP78 and GRP94, not only have essential roles in protein folding and trafficking in the ER but disruptions of chaperone functions sensitize cells to agents that stress this organelle [59–64]. Therefore, we assessed whether the PWS INS-1 lines and their reduced levels of ER chaperones (**Fig. 3**-**5**) would accentuate activation of the three sensory proteins of the UPR. First, ER stress activates IRE1α to catalyze mRNA processing of the *Xbp1* mRNA to generate the functional XBP1 transcription factor [64]; this non-canonical “splicing” of *Xbp1* transcripts occurs at low and equal levels in PWS and control INS-1 cells under DMSO control growth conditions but is enhanced by treatment with thapsigargin to initiate ER stress (**Fig. 6A,D**; **Fig. S19A,B**). Importantly, production of “spliced” *Xbp1* mRNA in thapsigargin-treated cells occurs significantly more robustly at earlier 2 h and 3 h timepoints for PWS than for control INS-1 lines (**Fig. 6A,D**; **Fig. S19A,B**) but normalizes by 4-5 h (**Fig. S19A,B**). These results reveal that while the initial magnitude of IRE1 activation is greater in the PWS INS-1 cells, the duration and cumulative response is comparable. Second, phosphorylation of eIF2α by PERK [63] was assessed, with PWS INS-1 lines showing significantly higher levels of eIF2α pSer51 phosphorylation than control lines when ER stress was induced by 5 h of thapsigargin treatment (**Fig. 6B,E**). Phosphorylated eIF2α inhibits general translation but preferentially translates certain stress adaptive genes, including ATF4 and CHOP; transcripts of these genes were unaffected in unstressed PWS INS-1 cells as measured by RNA-seq.

**Figure 6.**
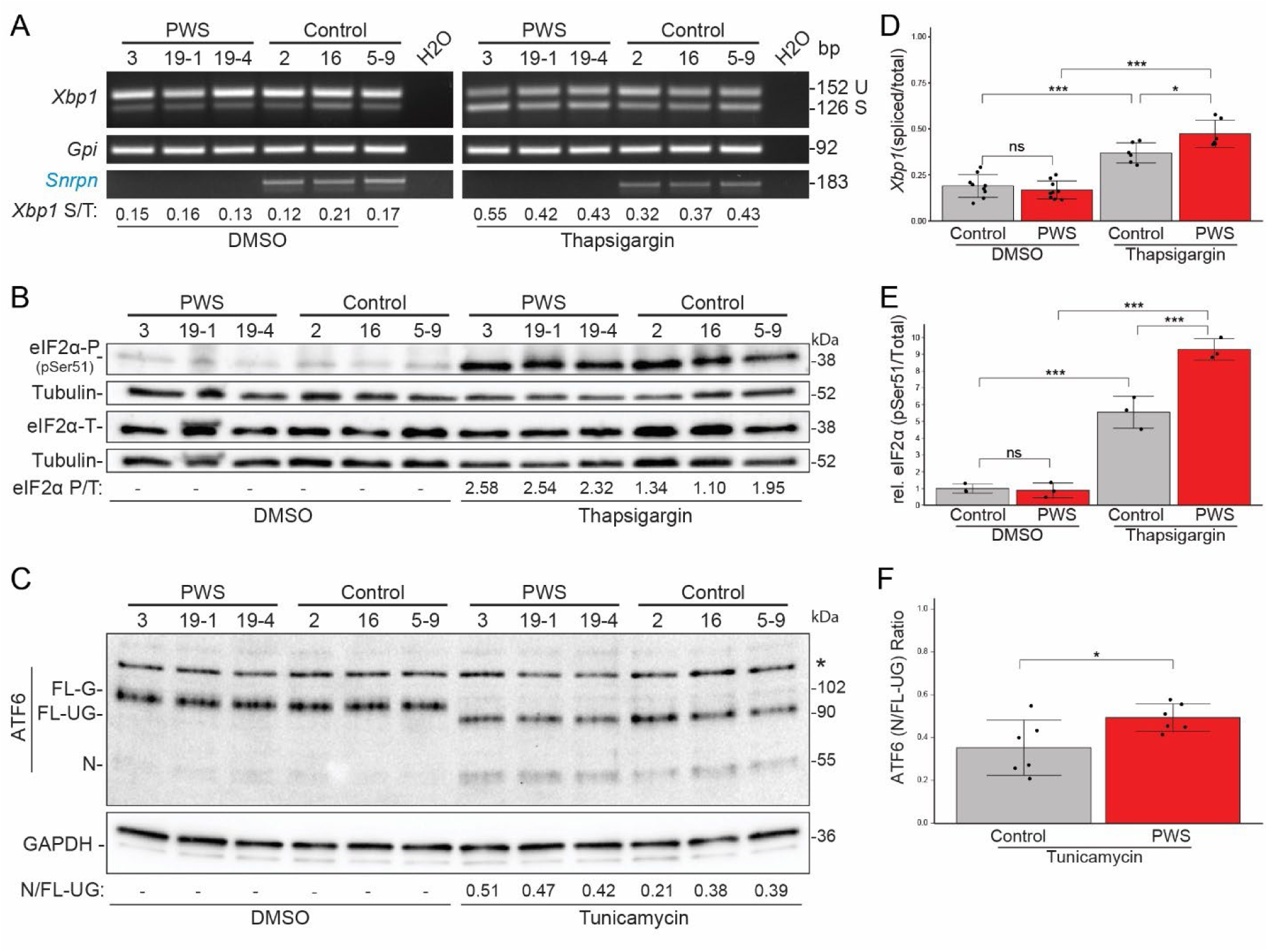
More sensitive activation of all three ER stress master regulatory pathways in PWS-deletion *vs.* control INS-1 β-cell lines. **(A)** For the IRE1α pathway, we assessed its mRNA processing of *Xbp1* from unspliced (U) to spliced (S), with control gene *Gpi* and PWS-*Snord116* also assayed by RT-PCR, using RNA from PWS (3, 19-1, 19-4) and control (5-9, 2, 16) INS-1 cell lines treated with DMSO or 0.1 µM thapsigargin for 3 h. At bottom, the *Xbp1* S/T (spliced/total) ratios for each cell line are shown, under both control (DMSO) and ER stress (thapsigargin) conditions. **(B)** The PERK pathway was assessed by comparison of total eIF2α to PERK phosphorylated eIF2α-P (pSer51), using whole cell lysates from PWS and control INS-1 cell lines treated with DMSO or 0.1 µM thapsigargin for 5 h. Anti-αTUB was used as a control for protein-loading levels. At bottom, the eIF2α P/T (phosphorylated/total) ratios for each cell line are shown under the ER stress (thapsigargin) condition. **(C)** For the ATF6 pathway, we used whole cell lysates from PWS and control INS-1 cell lines treated with DMSO or 10 µg/ml tunicamycin for 5h. Under control DMSO conditions, full-length (FL) and glycosylated (G) ATF6 of ∼ 100-kD is ER-localized. tunicamycin inhibits biosynthesis of N-linked glycans, resulting in unglycosylated (UG) ATF6-FL of 90-kD in the ER, with proteolytic processing in the Golgi to produce nuclear (N) ATF6-N of ∼ 55-kD. *, non-specific band detected by the anti-ATF6 antibody. Anti-GAPDH was used as a control for protein-loading levels. At bottom, the ATF6 N/FL-UG (nuclear/FL-unglycosylated) ratios for each cell line are shown under the ER stress (tunicamycin) condition. **(D)** Quantitation of the ratio of spliced/total (S/T) *Xbp1* mRNA in the PWS and control INS-1 lines under control (DMSO; n=9 each genotype) and thapsigargin (3 h; n=6 each) conditions. PWS β-cells are more sensitive to thapsigargin-induced ER stress with earlier and more robust activation of *Xbp1* mRNA “splicing” than control cell lines. **(E)** Quantitation of the relative (rel.) level of phosphorylated (pSer51) to total eIF2α in the PWS and control INS-1 lines (n=3 each) under control (DMSO) and thapsigargin conditions. PWS cells are more sensitive to thapsigargin-induced ER stress with more robust activation of eIF2α phosphorylation. **(F)** Quantitation of the ratio of nuclear/FL-unglycosylated (N/UG) for ATF6 in the PWS and control INS-1 lines under ER stress (tunicamycin) conditions (n=6 each, from two biological replicates). PWS cells are more sensitive to tunicamycin-induced ER stress with more robust activation of ATF6-N. For **(D-E)**, statistical comparison by ANOVA and for **(F)** by Welch’s t-test with Tukey’s HSD post-hoc: *, *P* < 0.05; **, *P* < 0.005; ***, *P* < 0.0005; ns, not significant.

Third, dissociation of ER-retained ATF6 from GRP78 by ER stressors such as de-glycosylation agents [60, 61] enable it to move to the Golgi, where it is processed to an activated nuclear (N) form, ATF6-N, which regulates expression of genes encoding many ER chaperones, including GRP78 and others. In PWS and control INS-1 cells, ER stress induced by tunicamycin, as expected, de-glycosylates full-length glycosylated (FL-G) ATF6 of ∼ 102-kD to an unglycosylated 90-kD (FL-UG) form and the processed ATF6-N form of 55-kD (**Fig. 6C**; **Fig. S20A**). The loss of the ATF6 FL-G form by deglycosylation occurs at the same rate for PWS and control cell lines (**Fig. S20B**). However, the ratio of the processed ATF6-N (due to higher production in PWS than control INS-1 cells) to full-length unglycosylated 90-kD (with a higher level in control than PWS INS-1 cells) is significantly greater for PWS compared to control INS-1 cells at 2-5 h timepoints (**Fig. 6C,F**; also see **Fig. S20A,B**). Thus, ATF6 is being activated to ATF6-N earlier and more robustly in PWS than in control INS-1 β-cell lines. Combined, these results on ER stress activation of the IRE1α/XBP1, PERK/eIF2α, and ATF6-N pathways indicate that PWS INS-1 β-cells are more sensitive than control INS-1 β-cells to ER stressors.

## DISCUSSION

Although PWS has long been assumed to be a neuroendocrine disease of the hypothalamus [3, 23], studies using mouse models have implicated a role for the pancreatic endocrine system [25, 58]. Here, we generated a novel INS-1 β-cell model through deletion of the ∼ 3.16 Mb PWS-orthologous imprinted domain in which to investigate PWS β-cell biology. The PWS deletion results in a β-cell autonomous defect in both basal and regulated insulin secretion, although the latter is at least in part consequent to the effect on basal insulin secretion. The PWS deletion also results in cell autonomous deficits in the secretion of other peptide hormones and in deficiencies of β-cell ER chaperones required for folding, exit and trafficking of secretory peptides. Presciently, studies reported 25 years ago demonstrated that PWS adults compared to matched obese controls had significantly reduced first- and second-phase insulin secretion during intravenous glucose tolerance tests [11], attesting to the translational impact of the PWS INS-1 model. These findings strongly support a hypothesis that PWS-genes are required for fundamental β-cell mechanisms affecting production of peptide hormones.

### PWS β-cells have concerted deficits in ER chaperone expression and insulin production/secretion

Basal insulin secretion and GSIS are dramatically reduced both *in vivo* in the TgPWS-deletion mouse model [25] and the new *in vitro* PWS INS-1 β-cell model. As PWS β-cells in both models are glucose-responsive this indicates the defect is in a fundamental component of the secretory apparatus. In TgPWS mice, perinatal pancreatic insulin secretion dysfunction was associated with intracellular retention of aged insulin and reduced islet insulin and C-peptide content as well as reduced islet glucagon content and plasma glucagon levels suggestive of broader endocrine cell dysfunction [25]. Extending these observations, in our current study we have found that all insulin isoforms that are expressed in and secreted from the PWS INS-1 β-cell are decreased, as measured by proteomics and western blot analyses. Remarkably, these equivalent insulin secretion deficits between PWS mice [25] and the cell autonomous INS-1 model of PWS (this work) were associated with distinct transcriptional responses. *In vivo*, an apparent physiological compensatory mechanism increased levels of mRNAs encoding all major pancreatic hormones, including insulin, amylin, glucagon, somatostatin, and pancreatic polypeptide, and several other secretory polypeptides [25]. In contrast, in PWS INS-1 cell lines no changes in gene expression were found for the endogenous rat *Ins1* or *Ins2* genes while there was diminished mRNA expression of a *mIns2::mCherry* transgene and those encoding two other secreted peptides (*Iapp* and *Npy*). Taken together, either translation or post-translational protein folding and/or trafficking of insulins must be dysregulated in PWS β-cells.

A further critical finding in the PWS INS-1 model is a concurrent down-regulation at the transcriptional and polypeptide levels of numerous ER chaperones, including for GRP78 and GRP94 that are major facilitators of insulin folding and trafficking in the ER. GRP94 directly binds pro-insulin, and chemical inhibition or shRNA knockdown or genetic ablation of the *Hsp90b1* gene results in diminished insulin processing and secretion, larger but immature insulin granules, and activation of the PERK ER stress pathway [65]. Similarly, knockdown of *Hspa5* in INS-1(832/13) cells reduced insulin biosynthesis and secretion [66], a finding validated in human EndoC-βH1 cells [67]. The acute reduction of single ER chaperones resulting in reduced insulin production and secretion indicates a direct impact at the level of ER insulin protein folding and trafficking that is recapitulated in the PWS INS-1 model where we describe a chronic shortage of multiple ER-chaperones. Therefore, we interpret the broad ER-chaperone deficiency in the PWS INS-1 model as a primary molecular abnormality that directly results in diminished insulin secretion.

### Chronic stress-independent ER chaperone deficits in PWS β-cells prevents ER chaperone dosage compensation

Prior evidence from mouse knockout models of key ER chaperone genes has shown that they are subject to dosage compensation to maintain homeostasis. Although knockouts for either *Hspa5* (GRP78) or *Hsp90b1* (GRP94) are lethal early in mouse embryogenesis, heterozygous mice are viable [34, 68–70]. Intriguingly, upregulation of GRP94 and other ER chaperones occurs in *Hspa5* +/- mice [68] and similarly for GRP78 in *Hsp90b1* -/- ES cells [69], indicative of dosage compensation among ER chaperones. A similar mechanism operates in *Hspa5* +/- mice to attenuate diet-induced obesity and insulin resistance [70]. Compensatory changes with upregulation of GRP94 or GRP78 and other ER chaperones including PDIA6 also occurred with shRNA knockdown of *Hspa5* or *Hsp90b1*, respectively, in a mouse cell line [71]. In contrast, the widespread deficiency of ER chaperones, including GRP78 and GRP94, that we describe for PWS INS-1 β-cells would preclude compensatory increases. Because of an inability to compensate, we hypothesize that PWS β-cells have a chronic deficit in ER chaperone production which would interfere with folding of proinsulin and/or delay ER transit of cargo (e.g., hormones) along the secretory pathway. Consistent with this hypothesis, our previous *in vivo* results using a Timer fluorescent protein demonstrated an accumulation of aged insulin in islets of PWS mice [25].

ER chaperones also function as sensors of ER stress, a physiological quality control mechanism responding to accumulation or failure in degradation of misfolded secretory proteins [3, 59, 62, 72, 73]. ER stress responses, governed by the IRE1α/XBP1, PERK/eIF2α/Ddit3(CHOP), and ATF6 regulatory pathways, aim to recover ER homeostasis by upregulating ER chaperone gene expression as part of the UPR [38, 59, 62]. In unstressed cells, GRP78 forms stable complexes with each of IRE1α, PERK, and ATF6 in the ER lumen [59, 60], whereas ER stress releases GRP78 and induces each pathway. Due to the demand to fold high levels of insulin, a protein notoriously difficult to fold [73], β-cells have a basal level of ATF6 activation to maintain higher levels of GRP78 than for most cell types [74]. In contrast, unresolved ER stress within β-cells leads to diabetes in mouse models and human [38, 75]. For example, insulin gene mutations lead to insulin misfolding with induction of ER-stress and broad upregulation of the UPR [76], a converse mechanism to the reduced ER chaperone levels we demonstrated within PWS INS-1 cells under both basal conditions and during GSIS. Indeed, lower GRP78 levels in PWS β-cells likely accounts for the earlier and more robust ER stress induction by thapsigargin or tunicamycin by lowering the threshold required to dissociate ER stress activators resulting in cells that are poised to a greater degree for response by all three master regulatory pathways. Although normal β-cells *in vivo* cycle through states of elevated and low UPR coordinate with insulin gene expression and ER protein folding load [77–79], our omics data on steady-state mRNA and protein levels of bulk PWS INS-1 cells under basal unstressed or GSIS conditions suggests a stress independent defect in ER chaperone gene expression.

### A putative PWS-ER chaperone gene regulatory network (GRN)

The coordinate downregulation of at least ten ER chaperone genes, including *Hspa5* and *Hsp90b1*, in PWS β-cells, but not the whole suite of UPR genes, supports the hypothesis that a PWS-imprinted gene or genes governs a pathway that coordinately regulates ER chaperones in a unique gene regulatory network (GRN). Although many of the DEGs that encode ER chaperones in PWS INS-1 β-cells are known ATF6 and/or XBP1 targets, numerous ATF6, IRE1α/XBP1, and ATF4/CHOP (PERK pathway) target genes [80–89] are not dysregulated in PWS INS-1 lines, suggesting a novel GRN in PWS β-cells. Candidate transcription factors (TFs) within the PWS-ER chaperone GRN are those with binding site motifs enriched in the promoters of DEGs in PWS β-cells, including, but not limited to the known ATF6 co-factor NFYA, which has roles in insulin secretion and glucose homeostasis [90], as well as PPARB/D also with known roles in β-cell mass and insulin secretion [91]. Intriguingly, a recently published study found that TFs of the nuclear receptor 4A (NR4A) family that participate in long-term memory in mice act through coordinate regulation of numerous ER chaperones [92], with a high degree of concordance with the dysregulated ER chaperones that we identified in PWS INS-1 cell lines; nevertheless, NR4A pathway factors are not DEGs in PWS β-cells.

It also remains to be determined which PWS gene or genes regulates the putative PWS→TF→ER chaperone network. Neither *Magel2* nor *Ndn* have a role, as they are not expressed in INS-1 cells. The top candidate PWS genes that are highly expressed in INS-1 cells are *Snrpn*, encoding the SmN spliceosomal protein [42], *Snurf*, encoding a small arginine-rich nuclear protein [18], and *Snord116*, a tandemly duplicated C/D-box snoRNA that may function as a sno-lncRNA that sequestrates FOX2 and other RNA binding proteins in alternative splicing pathways [22, 93], or through direct binding and regulation of as yet undefined RNA targets. Further studies are necessary to distinguish amongst the candidate PWS genes and to identify downstream GRN steps leading to coordinated ER chaperone expression.

### Evolution of PWS gene functions in secretory endocrine cells

Recent work has shown hypothalamic deficiencies in *Magel2*-null mice as well as for iPSC- and dental-pulp stem cell neuronal models, specifically in secretory granule components including PCSK1, PCSK2, CPE, granins (e.g., CHGB and others), and numerous neuropeptides [23]. These studies established that MAGEL2 plays a role in hypothalamic neuroendocrine cells by regulating neuropeptide production and secretion via endosome recycling to prevent lysosomal degradation of secretory granule proteins [23], a process downstream of the ER in protein trafficking and secretory pathways. As discussed above, our study shows that a different PWS-gene (e.g., *Snrpn*, *Snurf*, and/or *Snord116*) is putatively responsible for regulating ER chaperone and hormone (e.g., insulin) biosynthesis and secretion from pancreatic β-cells. In contrast to the reported reduction of prohormone convertase PC1 in PWS iPSC-derived neurons and in *Snord116*-null mice [58], no deficiency in *Pcsk1* mRNA or PC1/3 levels or in *Pcsk2* or PC2 was observed in PWS β-cell lines, which may reflect differences in cell type, genetics, or experimental conditions. Combined, these observations in hypothalamic neuroendocrine cells and pancreatic β-cells indicate that at least two PWS-imprinted genes regulate related neuropeptide and peptide hormone secretory pathways, suggesting that the PWS domain may function as a “mammalian operon” or synexpression group [94, 95]. As the PWS-domain arose evolutionarily in a eutherian mammalian ancestor [96], this suggests emergence of functions acting as molecular rheostats to regulate secretory pathways in endocrine cells that control growth, metabolism, and neural pathways.

### PWS genes regulate glucose and hormone homeostasis: implications for metabolic disease

Maintaining glucose homeostasis relies on exquisite coordination between secretion of the opposing hormones insulin and glucagon from pancreatic β- and α-cells, respectively, in response to changes in blood glucose, and disruption of these processes contributes to the pathogenesis of metabolic disorders, including type 2 diabetes [38, 97–100]. Although insulin and glucagon play opposite regulatory roles both in normal metabolic homeostasis and dysfunction in type 2 diabetes, PWS mice are remarkable in having low blood levels of both insulin and glucagon [16, 25]. As recent studies indicate that GRP78 interacts with glucagon in α-cells [101], it will be important to further assess dysregulation of ER chaperones and glucagon secretion in PWS α-cells both in cell culture and within PWS mouse models. Intriguingly, quantitative trait locus (QTL) studies in mice link genetic variation in blood insulin and glucose levels in or near to the PWS-orthologous domain [102]. Further studies are warranted to examine the role of PWS-imprinted genes in these metabolic traits in humans, and whether β-cell or other endocrine cell deficits in ER chaperones and hormone secretion contribute to clinical phenotypes, including episodes of hypoglycemia in PWS subjects [13, 14]. As PWS genes regulate islet development, β- and α-cell secretion [16, 25], and a GRN affecting ER chaperone and insulin secretion (this study), identifying mechanisms by which PWS-genes carry out these critical β-cell functions will illuminate the pathogenesis and may reveal effective treatments for not only PWS, but for common disorders with deficits in glycemic homeostasis and islet hormone secretory pathways.

## MATERIALS AND METHODS

### Cell culture

INS-1 lines were cultured using “RPMI 1640 media without glucose” (Life Technologies) supplemented with 10% FBS, 10 mM HEPES, 1 mM sodium pyruvate, 0.05 mM 2-mercaptoethanol, 7.5 mM glucose, and antibiotics (1% Pen-Strep, 10 µg/ml each piperacillin and ciprofloxacin). Chemical treatments included 0.1 µM thapsigargin (Sigma) for either 5 hr or a 0-5 hr time-course, and 10 µg/ml tunicamycin (Sigma) for 5 hr or a 0-8 hr time-course. The INS-1 lines used for genome editing in this study were generated from parental INS-1(832/13) cells that have integrated a human *INS*-neoR transgene [33] as well as a mouse *Ins2*-C-mCherry transgene (Costa RA, et al., manuscript in preparation).

### Genome editing

Pairs of sgRNAs (**Table S6**) were designed (http://crispr.mit.edu/) to target sites flanking the rodent PWS-orthologous domain to delete all paternally-expressed imprinted genes (**Fig. 1A**), and cloned into the pX330 CRISPR/Cas9 vector [103] (Addgene). INS-1 parental cells were transfected using lipofectamine 3000 with pX330-vectors encoding sgRNAs targeting proximal of *Frat3* and distal (between *Snord115* and *Ube3a*) of the PWS-domain and/or the control pEGFP-N3 vector (using 3 µg or 500 ng of DNA per vector per T75 flask or 6-well plate, respectively). After 4 days culture at 30°C or 37°C, cells were harvested for DNA isolation or flow cytometry (FACS ARIAII in the Rangos Flow Cytometry Core Laboratory) with ∼ 300 GFP-positive cells plated in five 96-well plates and clonally expanded to 12-well plates for DNA isolation. Clonal lines were screened by deletion-PCR and positive lines further expanded for DNA, RNA, protein, and molecular cytogenetic analyses. PCR primers for deletion-PCR, inversion-PCR, and scarred- or intact-allele PCRs are in **Table S7**. For DNA sequencing (Genewiz) of deletion breakpoints and scarred or intact sgRNA sites, we Sanger sequenced PCR products directly or from pJET (Thermo Fisher) cloned PCR products. Potential CRISPR/Cas9 off-target sites were predicted *in silico* using CRISPOR (http://crispor.tefor.net/); top predicted off-targets were PCR amplified (**Table S7**) and directly Sanger sequenced.

### Molecular cytogenetics

Cytogenetic studies were carried out in the University of Pittsburgh Cell Culture and Cytogenetics Facility. Briefly, cultured INS-1 cell lines were treated with 0.1 µg/ml Colcemid 1 hr, harvested, fixed, and slides processed for metaphase FISH by standard methods [104]. Fluorescent probes were prepared by labeling BAC (BACPAC Genomics) DNA using nick translation (Enzo Life Sciences, Inc.), including CH230-114P11 from the central PWS-domain (encodes U1A-*Snurf*-*Snrpn*-*Snord107*-*Snord64*) with Orange-dUTP and control CH230-2B12 mapping several Mb distal of the PWS domain (encodes *Cyfip1*-*Nipa2*-*Nipa1*-*Herc2*) with Green-dUTP. Probe and slide preparation, hybridization, and DAPI staining were by standard cytogenetics methods, with FISH analyses on an Olympus BX61 epifluorescence microscope (Olympus Microscopes) and image capture and analysis using the Genus software platform on the Cytovision System (Leica Microsystems) [104].

### Droplet digital PCR (ddPCR) genomic copy number assays

For genomic copy number 5 ng of *EcoR*I digested genomic DNA was used as input with amplification of *Snord107*, PWS-IC, and *Mirh1* as targets and *Ube3a* as a reference; during clonal derivatization EvaGreen chemistry was used whereas Fam/Hex TaqMan probes were used in a final analysis. Absolute copy number of *Snord107* and Poisson confidence intervals were calculated by numerical approximation [105]. Primers and probes for copy number ddPCR are listed in **Table S7**.

### DNA methylation

Genomic DNA was bisulfite converted using the EZ DNA methylation Gold kit (Zymo Research). Outer first round genomic PCR was performed using primers to amplify the *Snurf*-*Snrpn* promoter region (annealing temperature or Ta of 64°C; **Table S7**), with subsequent second round genomic PCR performed using PCR primers specific for the maternal, methylated or paternal, unmethylated alleles (Ta of 60°C; **Table S7**). HotStart Taq polymerase (Qiagen) was used for all methylation PCR.

### Insulin secretion assays

Cells were plated at 1.0 x 10^6^ cells/well, then 24 hr later at ∼ 80% confluency were washed with PBS containing Mg^++^ and Ca^++^ (Gibco), pre-incubated 1.0 hr in KRBH (129 mM NaCl, 5 mM NaHCO_3_, 4.8 mM KCl, 1.2 mM KH2PO_4_, 1.2 mM MgSO_4_, 2.5 mM CaCl_2_, 10 mM HEPES, 0.1% BSA) with low glucose (LG; 2.8 mM) and washed in KRBH (without glucose). Following a 30 min incubation in KRBH-LG, 1.0 ml low glucose secreted fractions were removed and centrifuged 13,000 rpm for 5 min., aliquoted and stored at - 80°C. The cells were then incubated 30 min in KRBH-high glucose (HG; 22 mM), and high glucose secreted fractions similarly centrifuged and stored at -80°C. Finally, cells were washed in PBS containing Mg^++^ and Ca^++^, harvested in RIPA lysis buffer and protease inhibitor, centrifuged at 13,000 rpm for 15 min at 4°C and the supernatant (protein) stored at -80°C. Protein was determined using Pierce BCA protein assay kit (Thermo Fisher). Insulin was measured by the RIA and Biomarkers Core of the Penn Diabetes Research Center, University of Pennsylvania, using an ultrasensitive rat insulin ELISA kit (Alpco Diagnostics). ANOVA followed by the Tukey HSD post-hoc test was used to assess differences between different conditions and genotypes using Prism 8 (GraphPad Software, Inc).

### Reverse transcription-PCR

Total RNA was isolated from PWS and control INS-1 lines grown as above (7.5 mM glucose) by Trizol harvest and miRNeasy (Qiagen) column purification. RNA quality was assessed by RNA TapeStation (Agilent) and quantified by broad range Qubit (Thermo Fisher) fluorometric analysis. First strand cDNA synthesis from 1 µg RNA was carried out using random hexamer primed RT by using Super Script IV (Thermo Fisher). Primers for gene specific RT-PCR amplification are in **Table S8**.

### RT-Droplet digital-PCR (RT-ddPCR) assays

Template cDNA pools were generated from 1 µg RNA using cells grown under standard glucose conditions (as above) by using iScript (Bio-Rad) RT with first-strand synthesis priming from a mixture of random hexamers and oligo-dT. Droplet generation, PCR and reading of EvaGreen based ddPCR reactions was carried out using an automated QX200 ddPCR system (Bio-Rad). A total template input volume of 2 µl of cDNA diluted in a range from undiluted to 1:200 (30 ng to 0.15 ng RNA equivalents per reaction) determined empirically based on the absolute expression level of target genes was used per 20 µl reaction. Primer sequences are listed in **Table S8**. Expression of target genes was normalized to the stable and modestly expressed *Gpi* (1:5 diluted, 6.0 ng RNA equivalents) as the ratio of positive droplet concentration and normalized to the average expression levels of target genes in the control INS-1 lines. Relative expression of target genes in PWS and control lines was compared using the Welch’s t-test for unequal variance with a significant threshold of *P* < 0.05.

### Proteomics

Each of the 3 control cell lines (5-9, 2, 16) and the 3 PWS cell lines (3, 19-1, 19-4) were grown in 6-well plates, harvested and then washed with PBS containing Mg^++^ and Ca^++^, incubated 30 min in KRH-high glucose (22 mM, glucose-induced insulin secretion condition, with BSA excluded from the KRBH buffer) after which the media was collected, centrifuged 10 min 1300 rpm, and supernatant (secretory fractions 1-6; **Fig. 3A**) collected and stored at -80°C. Cellular proteins were harvested by direct addition of 1.0 ml of methanol-acetic acid (90% methanol, 9% water, 1% acetic acid) to each well, transferred to a microcentrifuge tube and centrifuged at 1300 rpm 15 min. The supernatant (soluble protein fractions 7-12; **Fig. 3A**) and pellets (insoluble protein fractions 13-18; **Fig. 3A**) were collected and stored at -80°C. The secretory and cellular fractions were dried *in vacuo* and stored at -80°C. In preparation for mass spectrometry, proteins from each fraction were dissolved in 8 M urea, 100 µM Triethylammonium bicarbonate (TEAB) pH 8.5, reduced with tris(2-carboxyethyl)phosphine (TCEP) and alkylated with chloroacetamide, then diluted to 2 M urea with 100 mM TEAB, addition of 0.5 µg trypsin (Promega) and placed in a 37°C shaker for 16 hrs. Secretory protein fractions were analyzed on an Orbitrap Elite mass spectrometer (Thermo Fisher) while cellular protein fractions were analyzed on an Orbitrap Fusion mass spectrometer (Thermo Fisher). The 6-plex Tandem Mass Tag (TMT) system for quantitative proteomics (Thermo Fisher) was used to compare methanol-acetic acid insoluble cellular proteins (fractions 13-18) for control and PWS cell lines. The TMT labeled samples were analyzed on an Orbitrap Fusion Lumos mass spectrometer (Thermo Fisher).

Peptide/protein identification and quantification were determined using IP2 (Integrated Proteomics Applications). The MS raw data files were converted using RawConverter [106] (version 1.1.0.23) with monoisotopic option. For peptide identification, tandem mass spectra were searched against a database including the UniProt Rat database one entry per gene (21589 entries released 5/30/2021), these entries scrambled between K and R to make a decoy database, common contaminants, peptides and custom proteins using ProLuCID [107], and data filtered using DTASelect [108]. Quantitation of TMT samples was calculated with Census version 2.51 [109] and filtered with an intensity value of 5000 and isobaric purity value of 0.6 [110]. The Quantitative@COMPARE feature of IP2 was used to determine statistical significance.

### Immunoblot analyses

Cells were grown under standard glucose conditions with or without the addition of DMSO, thapsigargin or tunicamycin (as above). Whole cell lysates were harvested by direct lysis in cold radioimmunoprecipitation buffer (RIPA) with the addition of EDTA and combined protease and phosphatase inhibitors (Thermo Fisher) followed by clearing of insoluble material by centrifugation at 13,000 RPM for 10 minutes. Proteins were separated by SDS-PAGE with 2-mercaptoethanol as a reducing agent using criterion sized 4-15% gradient tris-glycine for broad range of molecular weights greater than 30 kDa or 16.5% tris-tricine for high resolution of smaller proteins (Bio-Rad). Proteins were transferred to nitrocellulose membranes and probed with antibodies [commercially obtained except for ATF6 [63], with dilutions as in **Table S9**]. Chemiluminescence detection of HRP-conjugated secondary antibodies was performed with either Clarity or Clarity Max Western ECL detection kits on a Bio-Rad Chemi-Doc XRS+ imager. Densitometry measurements were made using Image Lab (Bio-Rad) software and exported for analysis in R. Statistical comparison of normalized protein levels between PWS and control samples was made by either Welch’s unequal variance t-test or by ANOVA with Tukey’s HSD post-hoc test for multiple comparison where appropriate, with a significance threshold of *P* < 0.05.

### Transmission electron microscopy

Monolayers of INS-1 cells were fixed in 2.5% glutaraldehyde in 100 mM PBS, post-fixed in aqueous 1% osmium tetroxide, 1% Fe_6_CN_3_ for 1 hr, and dehydrated prior to embedding in Polybed 812 resin (Polysciences). Ultrathin cross sections (60 nm) of the cells were obtained on a Riechart Ultracut E microtome, post-stained in 4% uranyl acetate for 10 min and 1% lead citrate for 7 min. Sections were viewed on a JEOL JEM 1400 FLASH transmission electron microscope (JEOL) at 80 KV. Images were taken using a bottom mount AMT digital camera (Advanced Microscopy Techniques).

### Confocal microscopy

For each of the 3 control and 3 PWS INS-1 lines 750,000 live cells were seeded on a 35 mm dish with an uncoated 14 mm glass bottom insert (Mattek) and allowed to grow for 36 hr in basal glucose conditions. Live cell imaging was performed on a Zeiss LSM 710 confocal microscope at 20x magnification to capture mCherry fluorescence and brightfield images.

### RNA-seq and bioinformatics

Total RNA (from cells grown under 7.5 mM glucose) was used to prepare rRNA depleted Illumina TruSeq 75-bp paired-end stranded (fr-firststrand) sequencing libraries and sequenced to a depth of ∼ 40 million reads on a NextSeq500 instrument at the UPMC Children’s Hospital of Pittsburgh Genomics Core. RNA-seq data analysis was performed using computational resources available from the University of Pittsburgh Center for Research Computing. Quality of reads was assessed by FastQC and sequencing adapters trimmed by cutadapt using the combined TrimGalore tool [111]. The sequencing reads were aligned to the Ensembl v98 rat genome to which the sequence and annotation of the human *INS*-neo^R^ and mouse *Ins2*-C-mCherry transgenes had been added along with annotation of previously undefined genomic features (e.g., *Ipw*, *ψSnurf*, and upstream *U1 Snurf-Snrpn* exons) using the STAR splice-aware aligner [112]. Gene feature counts were tabulated with HTSeq using either the default (--nonunique none) or customized (--nonunique all) options to enable the inclusion of multi-copy (i.e., *Snord116*) and overlapping (bicistronic *Snurf-Snrpn*) genes which otherwise were excluded as ambiguous reads under the default options [113]. Differential expression analysis was performed with DESeq2 using a cutoff of *P*adj < 0.1 as calculated by the Benjamini-Hochberg multiple comparison procedure [114]. Similar overall transcriptomic results were obtained with RSEM based analysis [115] but had lower counts for both multi-mapped reads for *Snord116* and for ambiguous bicistronic *Snurf*-*Snrpn* reads. Gene ontology enrichment analysis of up- and down-regulated gene sets was done using DAVID [116]. Additional gene ontology and upstream analysis utilized Enrichr [117].

Sequencing libraries of small RNAs including snoRNAs were made from the same total RNA isolation but using a miRNA-Seq (Qiagen) library prep kit with a modified size selection for up to 200-bp fragments. Single end reads were generated and sequenced. UMI tools [118] and Cutadapt [119] were used to deduplicate and remove adapters in preprocessing of the fastq reads for alignment with Bowtie2 [120]. Similar to the total RNA-seq, HTSeq required the --nonunique all option to accurately quantitate the multicopy snoRNA and miRNA genes, and DESeq2 was used for differential expression analysis to compare genotypes. A full set of bash scripts for analysis of both total and small RNA-seq data sets including custom annotations are provided at the github repository (https://github.com/KoppesEA/INS-1_PWS_RNA-Seq).

## DATA STATEMENT

The RNA-seq FASTQ files and processed data associated from this study have been deposited in the NCBI Gene Expression Omnibus (GEO) under SuperSeries GSE190337 including GSE190334 (total RNA-seq) and GSE190336 (small RNA-seq), assigned to BioProject PRJNA786769.

The TMT quantitative proteomics data files have been deposited at ProteomeXchange under accession number PXD034471.

## ACKNOWLEDGMENTS

We thank Dr. Michael R. Rickels and Heather W. Collins at the Penn Diabetes Center RIA/Biomarkers Core, Mary F. Sanchirico for the protein database, Dr. Régis A. Costa for off-target PCR primer design, and Drs. Rebecca A. Simmons and Dwi Kemaladewi for review of the manuscript. This project used the University of Pittsburgh Health Sciences Sequencing Core at UPMC Children’s Hospital of Pittsburgh, for RNA sequencing. This work was supported by a research grant to R.D.N. from the Foundation for Prader-Willi Research (FPWR) and funding to R.D.N. from the Storr Family Foundation through the Prader-Willi Syndrome Association (PWSA). J.J.M., J.K.D., and J.R.Y. were supported by the National Institute of General Medical Sciences (8P41 GM103533). This manuscript is dedicated to PWS families and especially the Storr family for their support of our PWS research program.

## AUTHOR CONTRIBUTIONS

EAK, MAJ, JJM, PL, DWL, DBS, JKD performed experiments and analyzed data; EAK performed bioinformatics analyses; EAK, JJM, DBS, JRY, SCW, SMG, HJP, PD, RDN designed experiments and interpreted data; RCW provided a critical reagent; EAK, RDN wrote the manuscript; all co-authors reviewed and approved manuscript drafts and final version.

## Abbreviations

ddPCR: droplet digital PCR
DEGs: differentially expressed genes
ER: endoplasmic reticulum
FISH: fluorescence *in situ* hybridization
GRN: gene regulatory network
GSIS: glucose-stimulated insulin secretion
INS-1: INS-1 (832/13)::mCherry insulinoma cells
PWS: Prader-Willi syndrome
PWS-IC: Prader-Willi syndrome imprinting center
RNA-seq: RNA sequencing
RT-PCR: reverse-transcription-PCR
RT-ddPCR: reverse-transcription droplet digital PCR
TgPWS: transgenic-PWS mouse
TFs: transcription factors
UPR: unfolded protein response.

## REFERENCES

1. Costa RA, Ferreira IR, Cintra HA, Gomes LHF, Guida LDC. Genotype-phenotype relationships and endocrine findings in Prader-Willi syndrome. Front Endocrinol. 2019;10:864. doi: 10.3389/fendo.2019.00864.

2. Chung MS, Langouet M, Chamberlain SJ, Carmichael GG. Prader-Willi syndrome: reflections on seminal studies and future therapies. Open Biol. 2020;10(9):200195. doi: 10.1098/rsob.200195.

3. Tauber M, Hoybye C. Endocrine disorders in Prader-Willi syndrome: a model to understand and treat hypothalamic dysfunction. Lancet Diabetes Endocrinol. 2021;9(4):235–46. doi: 10.1016/S2213-8587(21)00002-4.

4. Muscogiuri G, Formoso G, Pugliese G, Ruggeri RM, Scarano E, Colao A, RESTARE. Prader-Willi syndrome: An uptodate on endocrine and metabolic complications. Rev Endocr Metab Disord. 2019;20(2):239–50. doi: 10.1007/s11154-019-09502-2.

5. Tomita T, Greeley G, Jr., Watt L, Doull V, Chance R. Protein meal-stimulated pancreatic polypeptide secretion in Prader-Willi syndrome of adults. Pancreas. 1989;4(4):395–400. doi: 10.1097/00006676-198908000-00001.

6. Lee HJ, Choe YH, Lee JH, Sohn YB, Kim SJ, Park SW, et al. Delayed response of amylin levels after an oral glucose challenge in children with Prader-Willi syndrome. Yonsei Med J. 2011;52(2):257–62. doi: 10.3349/ymj.2011.52.2.257.

7. Deal CL, Tony M, Hoybye C, Allen DB, Tauber M, Christiansen JS, Growth Hormone in Prader-Willi Syndrome Clinical Care Guidelines Workshop Participants. Growth Hormone Research Society workshop summary: consensus guidelines for recombinant human growth hormone therapy in Prader-Willi syndrome. J Clin Endocrinol Metab. 2013;98(6):E1072–87. doi: 10.1210/jc.2012-3888.

8. Fountain MD, Jr., Schaaf CP. MAGEL2 and oxytocin-implications in Prader-Willi syndrome and beyond. Biol Psychiatry. 2015;78(2):78–80. doi: 10.1016/j.biopsych.2015.05.006.

9. Hirsch HJ, Eldar-Geva T, Bennaroch F, Pollak Y, Gross-Tsur V. Sexual dichotomy of gonadal function in Prader-Willi syndrome from early infancy through the fourth decade. Hum Reprod. 2015;30(11):2587–96. doi: 10.1093/humrep/dev213.

10. Goldstone AP, Patterson M, Kalingag N, Ghatei MA, Brynes AE, Bloom SR, et al. Fasting and postprandial hyperghrelinemia in Prader-Willi syndrome is partially explained by hypoinsulinemia, and is not due to peptide YY3-36 deficiency or seen in hypothalamic obesity due to craniopharyngioma. J Clin Endocrinol Metab. 2005;90(5):2681–90. doi: 10.1210/jc.2003-032209.

11. Schuster DP, Osei K, Zipf WB. Characterization of alterations in glucose and insulin metabolism in Prader-Willi subjects. Metabolism. 1996;45(12):1514–20. doi: 10.1016/s0026-0495(96)90181-x.

12. Paik KH, Jin DK, Lee KH, Armstrong L, Lee JE, Oh YJ, et al. Peptide YY, cholecystokinin, insulin and ghrelin response to meal did not change, but mean serum levels of insulin is reduced in children with Prader-Willi syndrome. J Korean Med Sci. 2007;22(3):436–41. doi: 10.3346/jkms.2007.22.3.436.

13. Ma Y, Wu T, Liu Y, Wang Q, Song J, Song F, et al. Nutritional and metabolic findings in patients with Prader-Willi syndrome diagnosed in early infancy. J Pediatr Endocrinol Metab. 2012;25(11-12):1103–9. doi: 10.1515/jpem-2012-0167.

14. Harrington RA, Weinstein DA, Miller JL. Hypoglycemia in Prader-Willi syndrome. Am J Med Genet A. 2014;164A(5):1127–9. doi: 10.1002/ajmg.a.36405.

15. Cummings DE, Clement K, Purnell JQ, Vaisse C, Foster KE, Frayo RS, et al. Elevated plasma ghrelin levels in Prader Willi syndrome. Nat Med. 2002;8(7):643–4. doi: 10.1038/nm0702-643.

16. Stefan M, Ji H, Simmons RA, Cummings DE, Ahima RS, Friedman MI, et al. Hormonal and metabolic defects in a Prader-Willi syndrome mouse model with neonatal failure to thrive. Endocrinology. 2005;146(10):4377–85. doi: 10.1210/en.2005-0371.

17. Mani BK, Shankar K, Zigman JM. Ghrelin’s Relationship to Blood Glucose. Endocrinology. 2019;160(5):1247–61. doi: 10.1210/en.2019-00074.

18. Gray TA, Saitoh S, Nicholls RD. An imprinted, mammalian bicistronic transcript encodes two independent proteins. Proc Natl Acad Sci U S A. 1999;96(10):5616–21. doi: 10.1073/pnas.96.10.5616.

19. Cavaillé J, Buiting K, Kiefmann M, Lalande M, Brannan CI, Horsthemke B, et al. Identification of brain-specific and imprinted small nucleolar RNA genes exhibiting an unusual genomic organization. Proc Natl Acad Sci U S A. 2000;97(26):14311–6. doi: 10.1073/pnas.250426397.

20. Nicholls RD, Knepper JL. Genome organization, function, and imprinting in Prader-Willi and Angelman syndromes. Annu Rev Genomics Hum Genet. 2001;2:153–75. doi: 10.1146/annurev.genom.2.1.153.

21. Kishore S, Stamm S. The snoRNA HBII-52 regulates alternative splicing of the serotonin receptor 2C. Science. 2006;311(5758):230–2. doi: 10.1126/science.1118265.

22. Wu H, Yin QF, Luo Z, Yao RW, Zheng CC, Zhang J, et al. Unusual processing generates SPA lncRNAs that sequester multiple RNA binding proteins. Mol Cell. 2016;64(3):534–48. doi: 10.1016/j.molcel.2016.10.007.

23. Chen H, Victor AK, Klein J, Tacer KF, Tai DJ, de Esch C, et al. Loss of MAGEL2 in Prader-Willi syndrome leads to decreased secretory granule and neuropeptide production. JCI Insight. 2020;5(17):e138576. doi: 10.1172/jci.insight.138576.

24. Castle JC, Armour CD, Lower M, Haynor D, Biery M, Bouzek H, et al. Digital genome-wide ncRNA expression, including snoRNAs, across 11 human tissues using polyA-neutral amplification. PLoS One. 2010;5(7):e11779. doi: 10.1371/journal.pone.0011779.

25. Stefan M, Simmons RA, Bertera S, Trucco M, Esni F, Drain P, et al. Global deficits in development, function, and gene expression in the endocrine pancreas in a deletion mouse model of Prader-Willi syndrome. Am J Physiol Endocrinol Metab. 2011;300(5):E909–22. doi: 10.1152/ajpendo.00185.2010.

26. Benner C, van der Meulen T, Caceres E, Tigyi K, Donaldson CJ, Huising MO. The transcriptional landscape of mouse beta cells compared to human beta cells reveals notable species differences in long non-coding RNA and protein-coding gene expression. BMC Genomics. 2014;15:620. doi: 10.1186/1471-2164-15-620.

27. Li J, Klughammer J, Farlik M, Penz T, Spittler A, Barbieux C, et al. Single-cell transcriptomes reveal characteristic features of human pancreatic islet cell types. EMBO Rep. 2016;17(2):178–87. doi: 10.15252/embr.201540946.

28. Segerstolpe A, Palasantza A, Eliasson P, Andersson EM, Andreasson AC, Sun X, et al. Single-cell transcriptome profiling of human pancreatic islets in health and type 2 diabetes. Cell Metab. 2016;24(4):593–607. doi: 10.1016/j.cmet.2016.08.020.

29. Cnop M, Abdulkarim B, Bottu G, Cunha DA, Igoillo-Esteve M, Masini M, et al. RNA sequencing identifies dysregulation of the human pancreatic islet transcriptome by the saturated fatty acid palmitate. Diabetes. 2014;63(6):1978–93. doi: 10.2337/db13-1383.

30. Eizirik DL, Sammeth M, Bouckenooghe T, Bottu G, Sisino G, Igoillo-Esteve M, et al. The human pancreatic islet transcriptome: expression of candidate genes for type 1 diabetes and the impact of pro-inflammatory cytokines. PLoS Genet. 2012;8(3):e1002552. doi: 10.1371/journal.pgen.1002552.

31. Krishnan P, Syed F, Jiyun Kang N, Mirmira RG, Evans-Molina C. Profiling of RNAs from human islet-derived exosomes in a model of type 1 diabetes. Int J Mol Sci. 2019;20(23):5903. doi: 10.3390/ijms20235903.

32. Low BSJ, Lim CS, Ding SSL, Tan YS, Ng NHJ, Krishnan VG, et al. Decreased GLUT2 and glucose uptake contribute to insulin secretion defects in MODY3/HNF1A hiPSC-derived mutant β cells. Nat Commun. 2021;12(1):3133. doi: 10.1038/s41467-021-22843-4.

33. Hohmeier HE, Mulder H, Chen G, Henkel-Rieger R, Prentki M, Newgard CB. Isolation of INS-1-derived cell lines with robust ATP-sensitive K+ channel-dependent and -independent glucose-stimulated insulin secretion. Diabetes. 2000;49(3):424–30. doi: 10.2337/diabetes.49.3.424.

34. Zhu G, Lee AS. Role of the unfolded protein response, GRP78 and GRP94 in organ homeostasis. J Cell Physiol. 2015;230(7):1413–20. doi: 10.1002/jcp.24923.

35. Rajpal G, Schuiki I, Liu M, Volchuk A, Arvan P. Action of protein disulfide isomerase on proinsulin exit from endoplasmic reticulum of pancreatic β-cells. J Biol Chem. 2012;287(1):43–7. doi: 10.1074/jbc.C111.279927.

36. Sun M, Kotler JLM, Liu S, Street TO. The endoplasmic reticulum (ER) chaperones BiP and Grp94 selectively associate when BiP is in the ADP conformation. J Biol Chem. 2019;294(16):6387–96. doi: 10.1074/jbc.RA118.007050.

37. Kalwat MA, Scheuner D, Rodrigues-Dos-Santos K, Eizirik DL, Cobb MH. The pancreatic ß-cell response to secretory demands and adaption to stress. Endocrinology. 2021;162(11):bqab173. doi: 10.1210/endocr/bqab173.

38. Sharma RB, Landa-Galvan HV, Alonso LC. Living dangerously: protective and harmful ER stress responses in pancreatic β-cells. Diabetes. 2021;70(11):2431–43. doi: 10.2337/dbi20-0033.

39. Yong J, Johnson JD, Arvan P, Han J, Kaufman RJ. Therapeutic opportunities for pancreatic β-cell ER stress in diabetes mellitus. Nat Rev Endocrinol. 2021;17(8):455–67. doi: 10.1038/s41574-021-00510-4.

40. Skelin M, Rupnik M, Cencic A. Pancreatic beta cell lines and their applications in diabetes mellitus research. ALTEX. 2010;27(2):105–13. doi: 10.14573/altex.2010.2.105.

41. Watkins S, Geng X, Li L, Papworth G, Robbins PD, Drain P. Imaging secretory vesicles by fluorescent protein insertion in propeptide rather than mature secreted peptide. Traffic. 2002;3(7):461–71. doi: 10.1034/j.1600-0854.2002.30703.x.

42. Gray TA, Smithwick MJ, Schaldach MA, Martone DL, Graves JA, McCarrey JR, et al. Concerted regulation and molecular evolution of the duplicated SNRPB’/B and SNRPN loci. Nucleic Acids Res. 1999;27(23):4577–84. doi: 10.1093/nar/27.23.4577.

43. Liu Q, He H, Zeng T, Huang Z, Fan T, Wu Q. Neural-specific expression of miR-344-3p during mouse embryonic development. J Mol Histol. 2014;45(4):363–72. doi: 10.1007/s10735-013-9555-y.

44. Lawlor N, Marquez EJ, Orchard P, Narisu N, Shamim MS, Thibodeau A, et al. Multiomic profiling identifies cis-regulatory networks underlying human pancreatic β cell identity and function. Cell Rep. 2019;26(3):788–801. doi: 10.1016/j.celrep.2018.12.083.

45. Fujimoto K, Shibasaki T, Yokoi N, Kashima Y, Matsumoto M, Sasaki T, et al. Piccolo, a Ca2+ sensor in pancreatic beta-cells. Involvement of cAMP-GEFII.Rim2. Piccolo complex in cAMP-dependent exocytosis. J Biol Chem. 2002;277(52):50497–502. doi: 10.1074/jbc.M210146200.

46. Obermuller S, Calegari F, King A, Lindqvist A, Lundquist I, Salehi A, et al. Defective secretion of islet hormones in chromogranin-B deficient mice. PLoS One. 2010;5(1):e8936. doi: 10.1371/journal.pone.0008936.

47. Proverbio MC, Mangano E, Gessi A, Bordoni R, Spinelli R, Asselta R, et al. Whole genome SNP genotyping and exome sequencing reveal novel genetic variants and putative causative genes in congenital hyperinsulinism. PLoS One. 2013;8(7):e68740. doi: 10.1371/journal.pone.0068740.

48. Reinbothe TM, Alkayyali S, Ahlqvist E, Tuomi T, Isomaa B, Lyssenko V, et al. The human L-type calcium channel Cav1.3 regulates insulin release and polymorphisms in CACNA1D associate with type 2 diabetes. Diabetologia. 2013;56(2):340–9. doi: 10.1007/s00125-012-2758-z.

49. Fan F, Matsunaga K, Wang H, Ishizaki R, Kobayashi E, Kiyonari H, et al. Exophilin-8 assembles secretory granules for exocytosis in the actin cortex via interaction with RIM-BP2 and myosin-VIIa. Elife. 2017;6:e26174. doi: 10.7554/eLife.26174.

50. Matsunaga K, Taoka M, Isobe T, Izumi T. Rab2a and Rab27a cooperatively regulate the transition from granule maturation to exocytosis through the dual effector Noc2. J Cell Sci. 2017;130(3):541–50. doi: 10.1242/jcs.195479.

51. Drujont L, Lemoine A, Moreau A, Bienvenu G, Lancien M, Cens T, et al. RORγt+ cells selectively express redundant cation channels linked to the Golgi apparatus. Sci Rep. 2016;6:23682. doi: 10.1038/srep23682.

52. Nakajima-Nagata N, Sugai M, Sakurai T, Miyazaki J, Tabata Y, Shimizu A. Pdx-1 enables insulin secretion by regulating synaptotagmin 1 gene expression. Biochem Biophys Res Commun. 2004;318(3):631–5. doi: 10.1016/j.bbrc.2004.04.071.

53. Li L, Pan ZF, Huang X, Wu BW, Li T, Kang MX, et al. Junctophilin 3 expresses in pancreatic beta cells and is required for glucose-stimulated insulin secretion. Cell Death Dis. 2016;7(6):e2275. doi: 10.1038/cddis.2016.179.

54. Oda Y, Okada T, Yoshida H, Kaufman RJ, Nagata K, Mori K. Derlin-2 and Derlin-3 are regulated by the mammalian unfolded protein response and are required for ER-associated degradation. J Cell Biol. 2006;172(3):383–93. doi: 10.1083/jcb.200507057.

55. van Loon NM, Lindholm D, Zelcer N. The E3 ubiquitin ligase inducible degrader of the LDL receptor/myosin light chain interacting protein in health and disease. Curr Opin Lipidol. 2019;30(3):192–7. doi: 10.1097/MOL.0000000000000593.

56. Naito T, Takatsu H, Miyano R, Takada N, Nakayama K, Shin HW. Phospholipid flippase ATP10A translocates phosphatidylcholine and is involved in plasma membrane dynamics. J Biol Chem. 2015;290(24):15004–17. doi: 10.1074/jbc.M115.655191.

57. Kronenberg-Versteeg D, Eichmann M, Russell MA, de Ru A, Hehn B, Yusuf N, et al. Molecular pathways for immune recognition of preproinsulin signal peptide in type 1 diabetes. Diabetes. 2018;67(4):687–96. doi: 10.2337/db17-0021.

58. Burnett LC, LeDuc CA, Sulsona CR, Paull D, Rausch R, Eddiry S, et al. Deficiency in prohormone convertase PC1 impairs prohormone processing in Prader-Willi syndrome. J Clin Invest. 2017;127(1):293–305. doi: 10.1172/JCI88648.

59. Bertolotti A, Zhang Y, Hendershot LM, Harding HP, Ron D. Dynamic interaction of BiP and ER stress transducers in the unfolded-protein response. Nat Cell Biol. 2000;2(6):326–32. doi: 10.1038/35014014.

60. Shen J, Chen X, Hendershot L, Prywes R. ER stress regulation of ATF6 localization by dissociation of BiP/GRP78 binding and unmasking of Golgi localization signals. Dev Cell. 2002;3(1):99–111. doi: 10.1016/s1534-5807(02)00203-4.

61. Hong M, Luo S, Baumeister P, Huang JM, Gogia RK, Li M, et al. Underglycosylation of ATF6 as a novel sensing mechanism for activation of the unfolded protein response. J Biol Chem. 2004;279(12):11354–63. doi: 10.1074/jbc.M309804200.

62. Braakman I, Bulleid NJ. Protein folding and modification in the mammalian endoplasmic reticulum. Annu Rev Biochem. 2011;80:71–99. doi: 10.1146/annurev-biochem-062209-093836.

63. Teske BF, Wek SA, Bunpo P, Cundiff JK, McClintick JN, Anthony TG, et al. The eIF2 kinase PERK and the integrated stress response facilitate activation of ATF6 during endoplasmic reticulum stress. Mol Biol Cell. 2011;22(22):4390–405. doi: 10.1091/mbc.E11-06-0510.

64. Tsuchiya Y, Saito M, Kadokura H, Miyazaki JI, Tashiro F, Imagawa Y, et al. IRE1-XBP1 pathway regulates oxidative proinsulin folding in pancreatic β cells. J Cell Biol. 2018;217(4):1287–301. doi: 10.1083/jcb.201707143.

65. Ghiasi SM, Dahlby T, Hede Andersen C, Haataja L, Petersen S, Omar-Hmeadi M, et al. Endoplasmic reticulum chaperone glucose-regulated protein 94 is essential for proinsulin handling. Diabetes. 2019;68(4):747–60. doi: 10.2337/db18-0671.

66. Zhang L, Lai E, Teodoro T, Volchuk A. GRP78, but not protein-disulfide isomerase, partially reverses hyperglycemia-induced inhibition of insulin synthesis and secretion in pancreatic β-Cells. J Biol Chem. 2009;284(8):5289–98. doi: 10.1074/jbc.M805477200.

67. Szczerbinska I, Tessitore A, Hansson LK, Agrawal A, Ragel Lopez A, Helenius M, et al. Large-scale functional genomics screen to identify modulators of human β-cell insulin secretion. Biomedicines. 2022;10(1). doi: 10.3390/biomedicines10010103.

68. Luo S, Mao C, Lee B, Lee AS. GRP78/BiP is required for cell proliferation and protecting the inner cell mass from apoptosis during early mouse embryonic development. Mol Cell Biol. 2006;26(15):5688–97. doi: 10.1128/MCB.00779-06.

69. Mao C, Wang M, Luo B, Wey S, Dong D, Wesselschmidt R, et al. Targeted mutation of the mouse Grp94 gene disrupts development and perturbs endoplasmic reticulum stress signaling. PLoS One. 2010;5(5):e10852. doi: 10.1371/journal.pone.0010852.

70. Ye R, Jung DY, Jun JY, Li J, Luo S, Ko HJ, et al. Grp78 heterozygosity promotes adaptive unfolded protein response and attenuates diet-induced obesity and insulin resistance. Diabetes. 2010;59(1):6–16. doi: 10.2337/db09-0755.

71. Eletto D, Maganty A, Eletto D, Dersh D, Makarewich C, Biswas C, et al. Limitation of individual folding resources in the ER leads to outcomes distinct from the unfolded protein response. J Cell Sci. 2012;125(Pt 20):4865–75. doi: 10.1242/jcs.108928.

72. Tiwari A, Schuiki I, Zhang L, Allister EM, Wheeler MB, Volchuk A. SDF2L1 interacts with the ER-associated degradation machinery and retards the degradation of mutant proinsulin in pancreatic β-cells. J Cell Sci. 2013;126(Pt 9):1962–8. doi: 10.1242/jcs.117374.

73. Sun J, Cui J, He Q, Chen Z, Arvan P, Liu M. Proinsulin misfolding and endoplasmic reticulum stress during the development and progression of diabetes. Mol Aspects Med. 2015;42:105–18. doi: 10.1016/j.mam.2015.01.001.

74. Teodoro T, Odisho T, Sidorova E, Volchuk A. Pancreatic β-cells depend on basal expression of active ATF6α-p50 for cell survival even under nonstress conditions. Am J Physiol Cell Physiol. 2012;302(7):C992–1003. doi: 10.1152/ajpcell.00160.2011.

75. Salpea P, Cosentino C, Igoillo-Esteve M. A Review of Mouse Models of Monogenic Diabetes and ER Stress Signaling. Methods Mol Biol. 2020;2128:55–67. doi: 10.1007/978-1-0716-0385-7_4.

76. Hartley T, Siva M, Lai E, Teodoro T, Zhang L, Volchuk A. Endoplasmic reticulum stress response in an INS-1 pancreatic beta-cell line with inducible expression of a folding-deficient proinsulin. BMC Cell Biol. 2010;11:59. doi: 10.1186/1471-2121-11-59.

77. Preissler S, Ron D. Early events in the endoplasmic reticulum unfolded protein response. Cold Spring Harb Perspect Biol. 2019;11(4):a033894. doi: 10.1101/cshperspect.a033894.

78. Sharma RB, O’Donnell AC, Stamateris RE, Ha B, McCloskey KM, Reynolds PR, et al. Insulin demand regulates β cell number via the unfolded protein response. J Clin Invest. 2015;125(10):3831–46. doi: 10.1172/JCI79264.

79. Xin Y, Dominguez Gutierrez G, Okamoto H, Kim J, Lee AH, Adler C, et al. Pseudotime Ordering of single human β-cells reveals states of insulin production and unfolded protein response. Diabetes. 2018;67(9):1783–94. doi: 10.2337/db18-0365.

80. Adachi Y, Yamamoto K, Okada T, Yoshida H, Harada A, Mori K. ATF6 is a transcription factor specializing in the regulation of quality control proteins in the endoplasmic reticulum. Cell Struct Funct. 2008;33(1):75–89. doi: 10.1247/csf.07044.

81. Hassler JR, Scheuner DL, Wang S, Han J, Kodali VK, Li P, et al. The IRE1alpha/XBP1s pathway is essential for the glucose response and protection of β cells. PLoS Biol. 2015;13(10):e1002277. doi: 10.1371/journal.pbio.1002277.

82. Belmont PJ, Tadimalla A, Chen WJ, Martindale JJ, Thuerauf DJ, Marcinko M, et al. Coordination of growth and endoplasmic reticulum stress signaling by regulator of calcineurin 1 (RCAN1), a novel ATF6-inducible gene. J Biol Chem. 2008;283(20):14012–21. doi: 10.1074/jbc.M709776200.

83. Sharma RB, Darko C, Alonso LC. Intersection of the ATF6 and XBP1 ER stress pathways in mouse islet cells. J Biol Chem. 2020;295(41):14164–77. doi: 10.1074/jbc.RA120.014173.

84. Acosta-Alvear D, Zhou Y, Blais A, Tsikitis M, Lents NH, Arias C, et al. XBP1 controls diverse cell type- and condition-specific transcriptional regulatory networks. Mol Cell. 2007;27(1):53–66. doi: 10.1016/j.molcel.2007.06.011.

85. Adamson B, Norman TM, Jost M, Cho MY, Nunez JK, Chen Y, et al. A multiplexed single-cell CRISPR screening platform enables systematic dissection of the unfolded protein response. Cell. 2016;167(7):1867–82 e21. doi: 10.1016/j.cell.2016.11.048.

86. Plate L, Cooley CB, Chen JJ, Paxman RJ, Gallagher CM, Madoux F, et al. Small molecule proteostasis regulators that reprogram the ER to reduce extracellular protein aggregation. Elife. 2016;5::e15550. doi: 10.7554/eLife.15550.

87. Bergmann TJ, Fregno I, Fumagalli F, Rinaldi A, Bertoni F, Boersema PJ, et al. Chemical stresses fail to mimic the unfolded protein response resulting from luminal load with unfolded polypeptides. J Biol Chem. 2018;293(15):5600–12. doi: 10.1074/jbc.RA117.001484.

88. Park SM, Kang TI, So JS. Roles of XBP1s in transcriptional regulation of target genes. Biomedicines. 2021;9(7):791. doi: 10.3390/biomedicines9070791.

89. Wiseman RL, Mesgarzadeh JS, Hendershot LM. Reshaping endoplasmic reticulum quality control through the unfolded protein response. Mol Cell. 2022;82(8):1477–91. doi: 10.1016/j.molcel.2022.03.025.

90. Liu Y, He S, Zhou R, Zhang X, Yang S, Deng D, et al. Nuclear factor-Y in mouse pancreatic β-cells plays a crucial role in glucose homeostasis by regulating β-cell mass and insulin secretion. Diabetes. 2021;70(8):1703–16. doi: 10.2337/db20-1238.

91. Iglesias J, Barg S, Vallois D, Lahiri S, Roger C, Yessoufou A, et al. PPARβ/δ affects pancreatic β cell mass and insulin secretion in mice. J Clin Invest. 2012;122(11):4105–17. doi: 10.1172/JCI42127.

92. Chatterjee S, Bahl E, Mukherjee U, Walsh EN, Shetty MS, Yan AL, et al. Endoplasmic reticulum chaperone genes encode effectors of long-term memory. Sci Adv. 2022;8(12):eabm6063. doi: 10.1126/sciadv.abm6063.

93. Yin QF, Yang L, Zhang Y, Xiang JF, Wu YW, Carmichael GG, et al. Long noncoding RNAs with snoRNA ends. Mol Cell. 2012;48(2):219–30. doi: 10.1016/j.molcel.2012.07.033.

94. Niehrs C, Pollet N. Synexpression groups in eukaryotes. Nature. 1999;402(6761):483–7. doi: 10.1038/990025.

95. Nicholls RD. Incriminating gene suspects, Prader-Willi style. Nat Genet. 1999;23(2):132–4. doi: 10.1038/13758.

96. Rapkins RW, Hore T, Smithwick M, Ager E, Pask AJ, Renfree MB, et al. Recent assembly of an imprinted domain from non-imprinted components. PLoS Genet. 2006;2(10):e182. doi: 10.1371/journal.pgen.0020182.

97. Rorsman P, Braun M. Regulation of insulin secretion in human pancreatic islets. Annu Rev Physiol. 2013;75:155–79. doi: 10.1146/annurev-physiol-030212-183754.

98. Sandoval DA, D’Alessio DA. Physiology of proglucagon peptides: role of glucagon and GLP-1 in health and disease. Physiol Rev. 2015;95(2):513–48. doi: 10.1152/physrev.00013.2014.

99. Roder PV, Wu B, Liu Y, Han W. Pancreatic regulation of glucose homeostasis. Exp Mol Med. 2016;48:e219. doi: 10.1038/emm.2016.6.

100. Chen YC, Taylor AJ, Verchere CB. Islet prohormone processing in health and disease. Diabetes Obes Metab. 2018;20 Suppl 2:64–76. doi: 10.1111/dom.13401.

101. Asadi F, Dhanvantari S. Plasticity in the glucagon interactome reveals novel proteins that regulate glucagon secretion in α-TC1-6 cells. Front Endocrinol. 2018;9:792. doi: 10.3389/fendo.2018.00792.

102. Lawson HA, Lee A, Fawcett GL, Wang B, Pletscher LS, Maxwell TJ, et al. The importance of context to the genetic architecture of diabetes-related traits is revealed in a genome-wide scan of a LG/J x SM/J murine model. Mamm Genome. 2011;22(3-4):197–208. doi: 10.1007/s00335-010-9313-3.

103. Ran FA, Hsu PD, Wright J, Agarwala V, Scott DA, Zhang F. Genome engineering using the CRISPR-Cas9 system. Nat Protoc. 2013;8(11):2281–308. doi: 10.1038/nprot.2013.143.

104. Parikh RA, Appleman LJ, Bauman JE, Sankunny M, Lewis DW, Vlad A, et al. Upregulation of the ATR-CHEK1 pathway in oral squamous cell carcinomas. Genes Chromosomes Cancer. 2014;53(1):25–37. doi: 10.1002/gcc.22115.

105. Dube S, Qin J, Ramakrishnan R. Mathematical analysis of copy number variation in a DNA sample using digital PCR on a nanofluidic device. PLoS One. 2008;3(8):e2876. doi: 10.1371/journal.pone.0002876.

106. He L, Diedrich J, Chu YY, Yates JR, 3rd. Extracting accurate precursor information for tandem mass spectra by RawConverter. Anal Chem. 2015;87(22):11361–7. doi: 10.1021/acs.analchem.5b02721.

107. Xu T, Park SK, Venable JD, Wohlschlegel JA, Diedrich JK, Cociorva D, et al. ProLuCID: An improved SEQUEST-like algorithm with enhanced sensitivity and specificity. J Proteomics. 2015;129:16–24. doi: 10.1016/j.jprot.2015.07.001.

108. Tabb DL, McDonald WH, Yates JR, 3rd. DTASelect and Contrast: tools for assembling and comparing protein identifications from shotgun proteomics. J Proteome Res. 2002;1(1):21–6. doi: 10.1021/pr015504q.

109. Park SK, Aslanian A, McClatchy DB, Han X, Shah H, Singh M, et al. Census 2: isobaric labeling data analysis. Bioinformatics. 2014;30(15):2208–9. doi: 10.1093/bioinformatics/btu151.

110. Sandberg A, Branca RM, Lehtio J, Forshed J. Quantitative accuracy in mass spectrometry based proteomics of complex samples: the impact of labeling and precursor interference. J Proteomics. 2014;96:133–44. doi: 10.1016/j.jprot.2013.10.035.

111. Krueger F. Trim galore: A wrapper tool around Cutadapt and FastQC to consistently apply quality and adapter trimming to fastq files 2015 [cited 2020 March 20]. Available from: http://www.bioinformatics.babraham.ac.uk/projects/trim_galore/.

112. Dobin A, Davis CA, Schlesinger F, Drenkow J, Zaleski C, Jha S, et al. STAR: ultrafast universal RNA-seq aligner. Bioinformatics. 2013;29(1):15–21. doi: 10.1093/bioinformatics/bts635.

113. Anders S, Pyl PT, Huber W. HTSeq--a Python framework to work with high-throughput sequencing data. Bioinformatics. 2015;31(2):166–9. doi: 10.1093/bioinformatics/btu638.

114. Love MI, Huber W, Anders S. Moderated estimation of fold change and dispersion for RNA-seq data with DESeq2. Genome Biol. 2014;15(12):550. doi: 10.1186/s13059-014-0550-8.

115. Li B, Dewey CN. RSEM: accurate transcript quantification from RNA-Seq data with or without a reference genome. BMC Bioinformatics. 2011;12:323. doi: 10.1186/1471-2105-12-323.

116. Dennis G, Jr., Sherman BT, Hosack DA, Yang J, Gao W, Lane HC, et al. DAVID: Database for Annotation, Visualization, and Integrated Discovery. Genome Biol. 2003;4(5):P3.

117. Chen EY, Tan CM, Kou Y, Duan Q, Wang Z, Meirelles GV, et al. Enrichr: interactive and collaborative HTML5 gene list enrichment analysis tool. BMC Bioinformatics. 2013;14:128. doi: 10.1186/1471-2105-14-128.

118. Smith T, Heger A, Sudbery I. UMI-tools: modeling sequencing errors in Unique Molecular Identifiers to improve quantification accuracy. Genome Res. 2017;27(3):491–9. doi: 10.1101/gr.209601.116.

119. Martin M. Cutadapt removes adapter sequences from high-throughput sequencing reads. EMBnetjournal. 2011;17(1):3. doi: 10.14806/ej.17.1.200.

120. Langmead B, Salzberg SL. Fast gapped-read alignment with Bowtie 2. Nat Methods. 2012;9(4):357–9. doi: 10.1038/nmeth.1923.

