## Supporting Information for "Insulin secretion deficits in a Prader-Willi syndrome β-cell model are associated with a concerted downregulation of multiple endoplasmic reticulum chaperones"

[1] Division of Genetic and Genomic Medicine, Department of Pediatrics, UPMC Children's Hospital of Pittsburgh, Pittsburgh, PA 15224; [2] Department of Molecular Medicine and Neurobiology, The Scripps Research Institute, La Jolla, CA 92037; [3] Department of Cell Biology, University of Pittsburgh School of Medicine, Pittsburgh, PA 15261; [4] Department of Human Genetics, University of Pittsburgh School of Public Health, Pittsburgh, PA 15261; [5] Department of Biochemistry and Molecular Biology, Indiana University School of Medicine, Indianapolis, IN 46202

[#] Current address: Center for Genetics of Host Defense, UT Southwestern Medical Center, Dallas, TX 75390

### CONTENTS OF SUPPLEMENTARY DATA:

| Page | Content |
|---|---|
| 2 | **Summary of Tables** (including page numbers for each respective Table, **Tables S1-S9**, each appearing consecutively following the References). |
| 3-33 | **Figures S1-S20**, including legends. |
| 34 | **References**, for Supplementary Data. |

**TABLES**

**Table S1. Differentially expressed genes (DEGs) in PWS *vs.* control INS-1 lines from RNA-seq with HTSeq feature counts.** See pages 35-39.
Data presented from RNA-seq expression bioinformatics pipeline using a custom rat annotation build with STAR aligner, gene level counts with HTSeq (with options --mode union and --nonunique all) and differential expression with DESeq2. Top DEGs with a cutoff of *P*adj < 0.1 are shown. Full gene expression table available at GEO repository GSE190334.

**Table S2. Differentially expressed genes (DEGs) in PWS vs. control INS-1 lines from RNA-seq with RSEM feature counts.** See pages 40-43.
Data presented from RNA-seq expression bioinformatics pipeline using a custom rat annotation build with STAR aligner, gene level counts with RSEM and differential expression with DESeq2. Top DEGs with a cutoff of *P*adj < 0.1 are shown. Full gene expression table available at GEO repository GSE190334.

**Table S3. Significant differentially expressed small RNAs (DE sRNAs) in PWS *vs.* control INS-1 lines from small RNA-seq.** See pages 44-45.
Data presented from small RNA-seq expression bioinformatics pipeline, deduplicated with UMI Tools, mapped using a custom rat annotation build with Bowtie2 aligner, with gene level counts quantified by HTSeq (with option --nonunique all) and differential expression with DESeq2. Top DE sRNAs with a cutoff of *P*adj < 0.1 are shown. PWS sRNAs gene names are color coded *Snord116* (orange), *Snord115* (blue), *Mir344* (red), *Snord64* (green) and *Snord107* (purple). Full gene expression table available at GEO repository GSE190336.

**Table S4. Highly expressed miRNAs in INS-1 lines from small RNA-seq.** See pages 46-51.

**Table S5. Highly expressed snoRNAs in INS-1 lines from small RNA-seq.** See pages 52-55.

**Table S6. sgRNA oligonucleotides.** See page 56.
*Bbs*I site oligonucleotide cloning adapters for sgRNAs. Abbreviations: b/w, between; F, forward.

**Table S7. Genomic PCR and DNA methylation PCR primers.** See pages 57-58.
Abbreviations: F, forward; R, reverse; un, unmethylated; me, methylated; IC, imprinting center; chr, chromosome.

**Table S8. RT-PCR and RT-ddPCR primers.** See pages 59-60.
Abbreviations: F, forward; R, reverse.

**Table S9. Antibodies used in this study.** See page 61.

**Figure S1. CRISPR/Cas9 genome editing of the PWS-region generates maternal deletion INS-1 lines 5-5 and 5-9.**

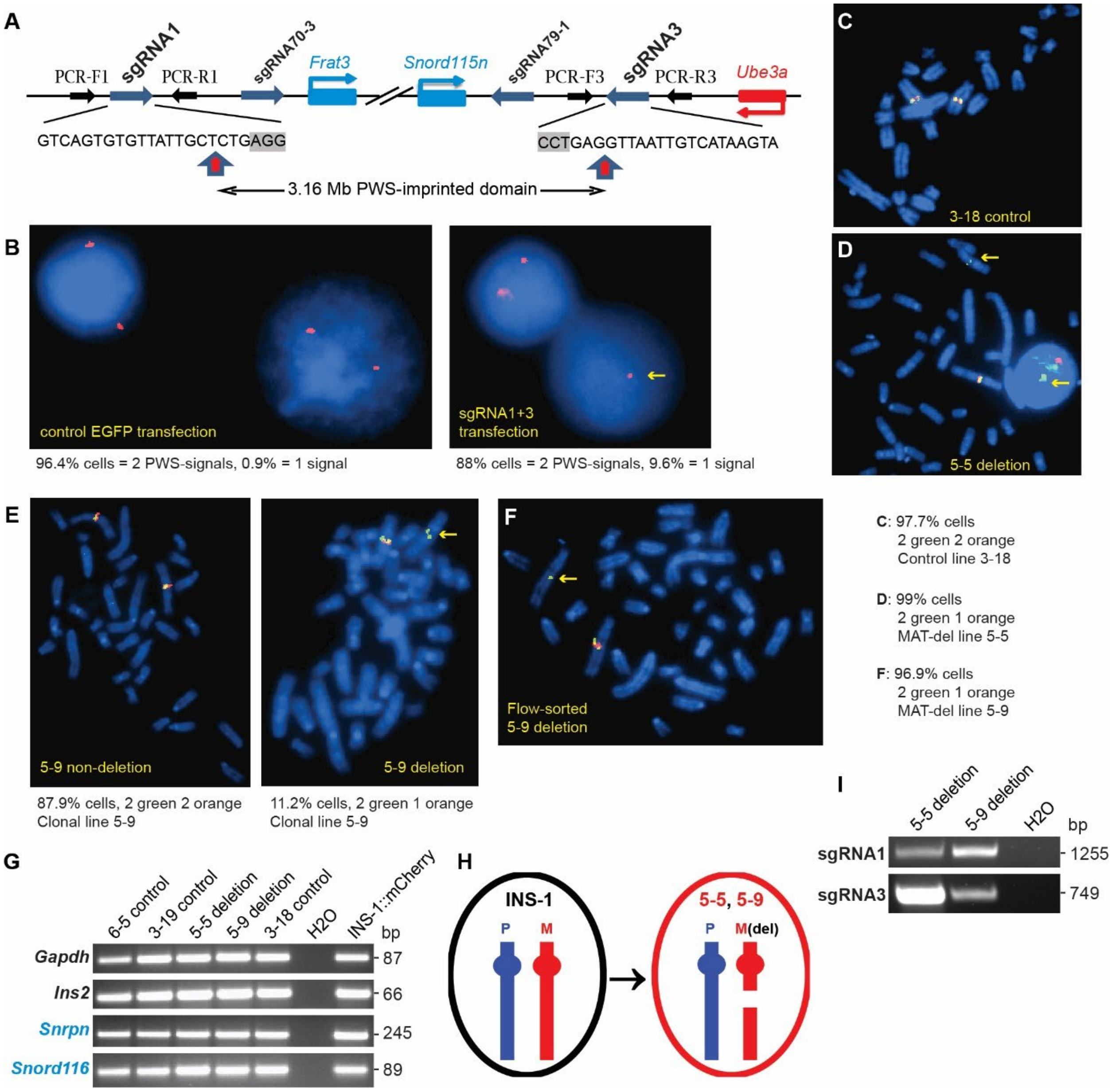

**Figure S1. CRISPR/Cas9 genome editing of the PWS-region generates maternal deletion INS-1 lines 5-5 and 5-9. (A)** Map of proximal sgRNA1 and distal sgRNA3 targeting sites. DNA sequences of the sgRNA targeting sites (blue arrows) are shown, with PAM motifs in grey highlight. Other symbols: blue boxes, proximal and distal PWS paternally-expressed genes (also see **Fig. 1A**); red box, flanking maternally-expressed gene; black arrows, PCR primers (F, forward; R, reverse); vertical arrows, canonical double-strand break (DSB) position catalyzed by CRISPR/Cas9. **(B-F)** Fluorescence *in situ* hybridization (FISH) with rat BAC probes CH230-114P11 spanning the PWS-IC (see **Methods**; **Fig. 1A**) and control CH230-2B12 (outside the PWS-domain) labeled with Orange-dUTP or Green-dUTP, respectively. Representative interphase nuclei or partial metaphases are shown. Yellow arrows, deletion of the PWS-domain. **(B)** 1-color FISH for parental INS-1::mCherry (INS-1) cells transfected with plasmid vectors expressing EGFP (left) or CRISPR (sgRNA1 + sgRNA3)/Cas9 (right). Most INS-1 control cells show diploid PWS BAC (orange) signals, with < 1% of cells having a single PWS-signal due to either a technical artifact (hybridization to a single allele) or to loss of a chromosome (while a small percent of cells shows increased signals due to artifact or to aneusomy). In contrast, genome editing greatly increases the percentage of cells with hemizygosity for the PWS-signal and hence a deletion for the PWS-locus. **(C)** Two-color FISH for control line 3-18, with most cells showing diploid signals for both BAC probes. **(D)** FISH for maternal (MAT)-deletion (del) line 5-5, with virtually all cells showing deletion of the PWS-locus. **(E)** FISH for MAT-del line 5-9 showing mosaicism, one cell population intact (and negative for the mCherry transgene) and one (mCherry-positive) deleted for the PWS-locus. **(F)** FISH on flow sorted mCherry-positive MAT-deletion line 5-9, almost all cells having the PWS-deletion. **(G)** INS-1 deletion lines 5-5 and 5-9 express PWS-imprinted genes. **(H)** Schematic of origin for MAT-deletion INS-1 lines 5-5 and 5-9, based on loss of maternal DNA methylation (**Fig. 1D**) and presence of paternal-gene expression data in **Fig. S1G**. Paternally- (P) and maternally-derived (M) chromosomes shown in blue or red, respectively (del, deletion). **(I)** Gels showing PCR fragments spanning sgRNA1 and sgRNA3 for MAT-del lines 5-5 and 5-9. See **Fig. S2C-H** for Sanger sequencing data on these PCR fragments.

**Figure S2. Sanger sequencing of genome editing events at sgRNA sites in derivation of maternal deletion INS-1 clonal lines.**

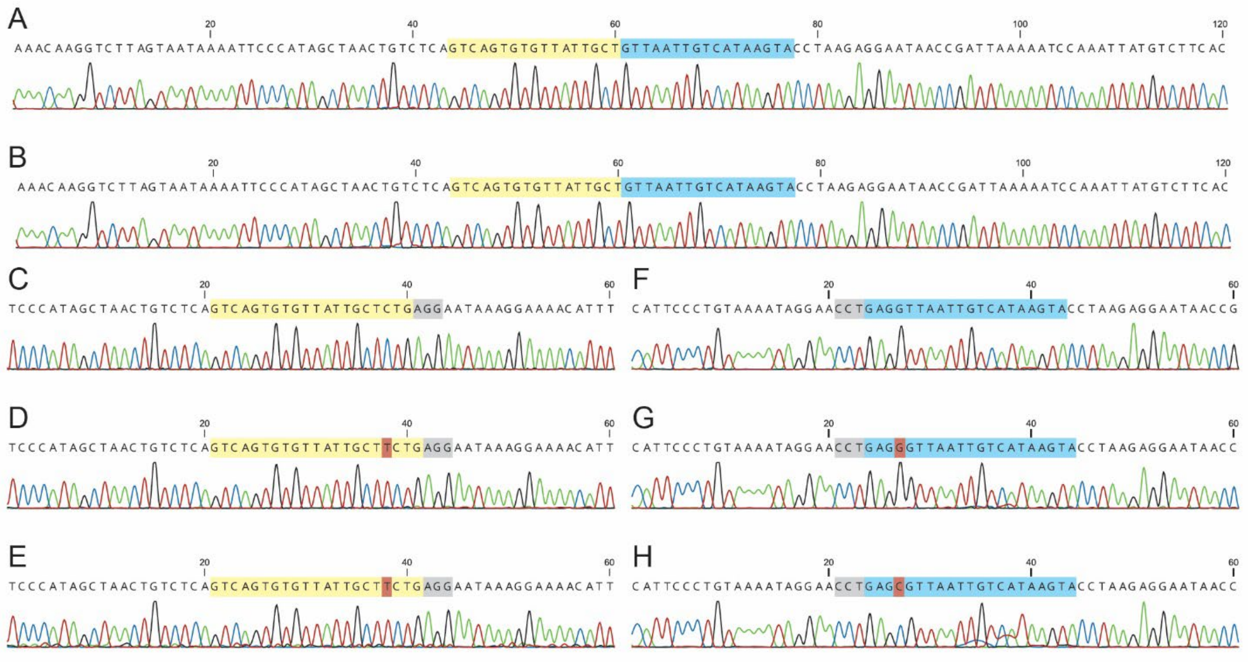

**Figure S2. Sanger sequencing of genome editing events at sgRNA sites in derivation of maternal deletion INS-1 clonal lines. (A-H)** Sanger sequence traces are shown highlighting sgRNA1 (yellow), sgRNA3 (blue), SpCas9 NGG PAMs (grey), and insertion mutations (pink). **(A-B)** Maternal-deletion allele of clonal lines 5-9 **(A)** and 5-5 **(B)** are identical canonical deletions with 3.16 Mb deleted between sgRNA1 and sgRNA3 due to a breakpoint from DNA repair of DSBs occurring at each position 3-nt upstream of the PAM (PAM-3) nuclease sites (see **Fig. S1A**). Deletion-PCR breakpoint fragments are from the **Fig. 1C** gel. **(C-E)** Sequence of intact sgRNA1 site in parental INS-1 **(C)**, and scarred alleles with a single T/A insertion at the canonical PAM-3 DSB position (pink highlight) in clonal lines 5-9 **(D)** and 5-5 **(E)**. Sequenced PCR fragments spanning sgRNA1 are from the **Fig. S1I** gel. **(F-H)** Sequence of intact sgRNA3 site in parental INS-1 **(F)**, and scarred alleles with a G/C insertion or a C/G insertion at the PAM-3 site (pink highlight) in line 5-9 **(G)** or line 5-5 **(H)**, respectively. Sequenced PCR fragments spanning sgRNA3 are from the **Fig. S1I** gel. It may be noted that as the deletions for 5-9 and 5-5 are on the maternal allele, the sgRNA1 and sgRNA3 scarred alleles can be inferred to occur on the paternal allele for each cell line. Further, as each of lines 5-5 and 5-9 have different sgRNA3 scarred alleles (despite sharing deletion breakpoints and sgRNA1 scarred allele mutations) then these two cell lines clearly arose as independent genome editing events.

**Figure S3. Sanger sequencing of sgRNA1 and sgRNA3 off-target sites in maternal deletion INS-1 clonal lines.**

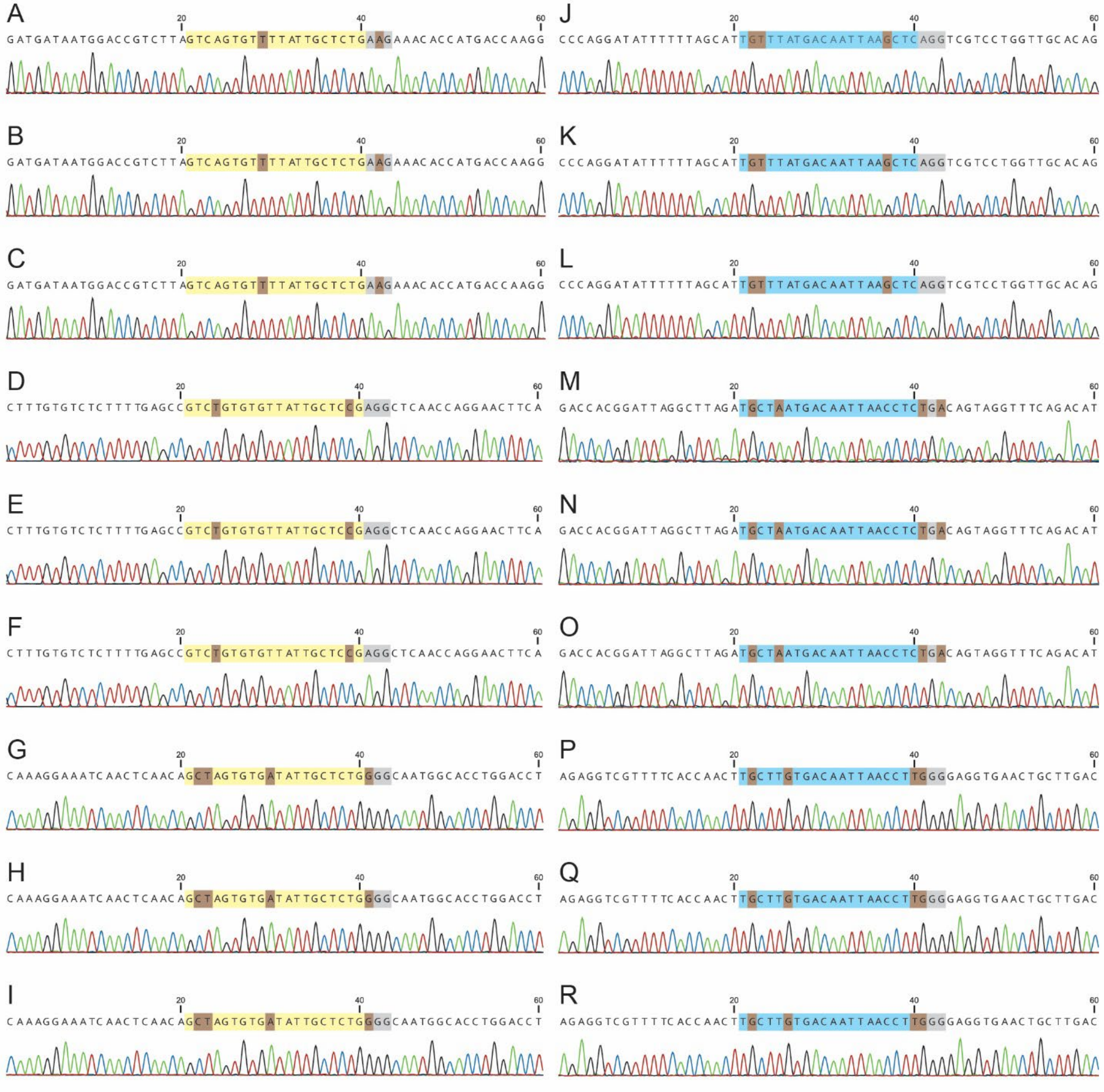

**Figure S3. Sanger sequencing of sgRNA1 and sgRNA3 off-target sites in maternal deletion INS-1 clonal lines.** Sanger sequencing chromatographs of direct sequenced off-target genomic PCRs are shown for the top three ranked predicted off-target sites for sgRNA1 highlighted by the sgRNA seed (yellow) and for sgRNA3 (blue), with SpCas9 NGG PAMs (grey), and deviation in off-target from sgRNA sequence (brown). **(A-C)** Sequence of intact sgRNA1 off-target site at chromosome 2 position 115,451,401-115,451,423 (+; intergenic *Gnb4-Actl6a*) with 1 mismatch in parental INS-1 **(A)** and maternal (MAT)-deletion lines 5-9 **(B)**, and 5-5 **(C)**. **(D-F)** Sequence of intact sgRNA1 off-target site at chromosome 6 position 102,394,187-102,394,209 (-; intron *Rgs6*) with 2 mismatches in parental INS-1 **(D)** and MAT-deletion lines 5-9 **(E)**, and 5-5 **(F)**. **(G-I)**

Sequence of intact sgRNA1 off-target site at chromosome 1 position 185,582,894-185,582,916 (+; intergenic *Htra1-Dmbt1*) with 3 mismatches in parental INS-1 **(G)** and MAT-deletion lines 5-9 **(H)**, and 5-5 **(I)**. **(J-L)** Sequence of intact sgRNA3 off-target site at chromosome 19 position 19,693,031-19,693,053 (+; intergenic *Cbln1-N4bp1*) with 3 mismatches in parental INS-1 **(J)** and MAT-deletion lines 5-9 **(K)**, and 5-5 **(L)**. **(M-O)** Sequence of intact sgRNA3 off-target site at chromosome 6 position 88,102,670-88,102,692 (+; intron *Sos2*) with 2 mismatches in parental INS-1 **(M)** and MAT-deletion lines 5-9 **(N)**, and 5-5 **(O)**. **(P-R)** Sequence of intact sgRNA3 off-target site at chromosome 9 position 105,005,251-105,005,273 (+; intron *Tmem232*) with 3 mismatches in parental INS-1 **(P)** and MAT-deletion lines 5-9 **(Q)**, and 5-5 **(R)**. In all off-target sites analyzed for both sgRNA1 and sgRNA3 there was no evidence of CRISPR-Cas9 induced dsDNA break repair resulting in small insertion-deletion events.

**Figure S4. Further CRISPR/Cas9 genome editing of the PWS-region generates paternal deletion INS-1 lines 3, 19-1, 19-4, and 25.**

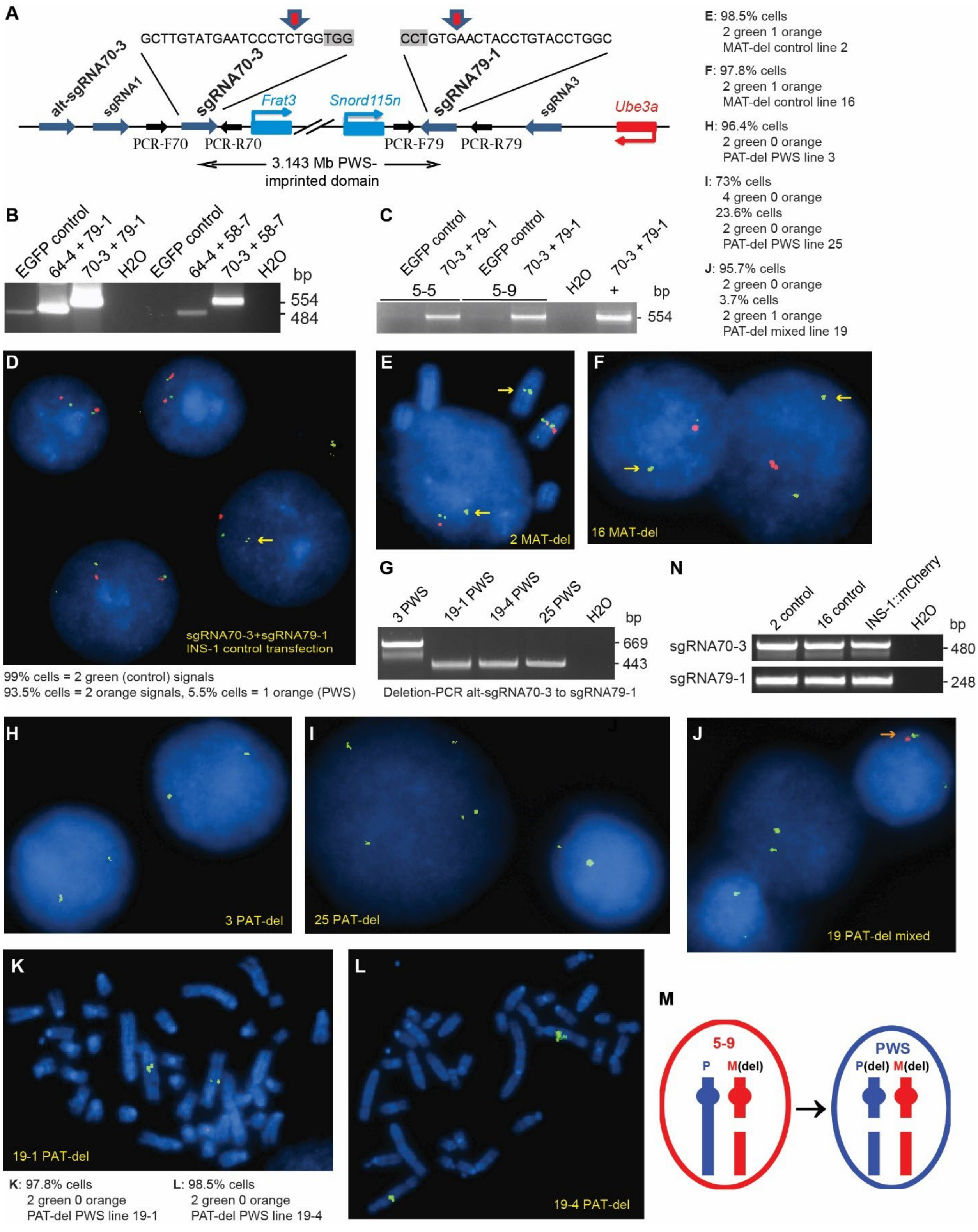

**Figure S4. Further CRISPR/Cas9 genome editing of the PWS-region generates paternal deletion INS-1 lines 3, 19-1, 19-4, and 25. (A)** Map of proximal sgRNA70-3 and distal sgRNA70-3 targeting sites on the paternal (PAT)-allele. Both sgRNA sites are absent on the maternal allele of lines 5-5 and 5-9 (**Fig. S1**), providing the PAT-allele specificity. Further proximal, the alt-sgRNA70-3 site differs at two 5′ nucleotides of the sgRNA and is part of a segmental-duplication (to be described elsewhere; manuscript in preparation). All symbols are as for **Fig. S1A**. **(B)** Deletion-PCR assays for a CRISPR/Cas9 screen of four pairs of sgRNAs flanking the 3.143 Mb PWS-domain, using control INS-1 cells. EGFP represents control transfections. Of the four pairs of sgRNAs, the highest efficiency is obtained for sgRNA70-3 + sgRNA79-1. **(C)** Deletion-PCR assay following transfection of MAT-del lines 5-5 and 5-9 using the CRISPR (sgRNA70-3 + sgRNA79-1)/Cas9-expressing vector shows specific targeting of the paternal allele of the PWS-domain. The positive (+) control is from **Fig. S4B**. **(D-F**, **H-L)** FISH studies using rat FISH probes, as for **Fig. S1B-F**. **(D)** FISH for parental INS-1 cells transfected with plasmid vectors expressing EGFP (not shown) or CRISPR (sgRNA70-3 + sgRNA79-1)/Cas9. Control cells for both probes (not shown) and the control probe (green) show diploid signals in virtually all cells, whereas for the PWS-probe (orange) in genome edited cells 5% of cells display an interphase with a PWS-region deletion (yellow arrow). **(E-N)** Studies on clonal lines derived from transfection of the MAT-del 5-9 cell line with an EGFP control vector (**E-F, N**) or with the CRISPR (sgRNA70-3 + sgRNA79-1)/Cas9-expressing vector (**G-M**). **(E)** Interphase and partial metaphase FISH for control line 2 with a MAT-deletion in virtually all cells. **(F)** FISH for control line 16 with a MAT-deletion in most cells. **(G)** Deletion-PCR for PAT-del line 3 using alt-sgRNA70-3 specific F and sgRNA79-1 R primers. As compared to the faint deletion breakpoint band using a sgRNA70-3 F PCR primer (**Fig. 1E**), this alternate (alt) assay provides a high degree of specificity, with fainter deletion-PCR bands for lines 19-1, 19-4, and 25 due to mismatches in the F primer flanking the sgRNA70-3 site. See **Fig. S5J** for Sanger sequencing data on the line 3 PCR fragment. **(H)** FISH for PAT-del (PWS) line 3 showing homozygous loss of the PWS-region in most cells. **(I)** FISH for PAT-del (PWS) line 25 shows homozygous loss of the PWS-region but mosaicism for cells tetrasomic or disomic for chromosome 1 (PWS-locus). **(J)** FISH for mixed line 19 shows mosaicism with most cells having homozygous loss of the PWS-region but a small % of cells with a MAT-deletion (from the parental 5-9 line) having an intact paternal PWS-region (orange arrow). **(K-L)** Single cells from dilution of mixed line 19 and 96-well plating underwent a further clonal isolation, followed by PCR screening of genomic DNA for loss of PWS-loci (not shown) and FISH, with isolation of 5 sub-lines. **(K)** Metaphase from FISH on PAT-del (PWS) line 19-1 showing homozygous loss of the PWS-region in most cells. **(L)** FISH for PAT-del (PWS) line 19-4 with homozygous loss of the PWS-region in virtually all cells. **(M)** Schematic of origin for PAT-deletion INS-1 lines 3, 19, 19-1 through 19-5, and 25. **(N)** Gels showing PCR fragments spanning sgRNA70-3 and sgRNA79-1 for parental INS-1 and control lines 2 and 16. See **Fig. S5A-F** for Sanger sequencing data on these PCR fragments. Note that there are no scarred allele sites for sgRNA70-3 and sgRNA79-1 in the paternal-deletion lines 19-1, 19-4 and 25 as the maternal allele is deleted for these loci (see **Fig. 1A**; **Fig. S1A**, **S5A**).

**Figure S5. Sanger sequencing of genome editing events at sgRNA sites in derivation of paternal deletion INS-1 clonal lines.**

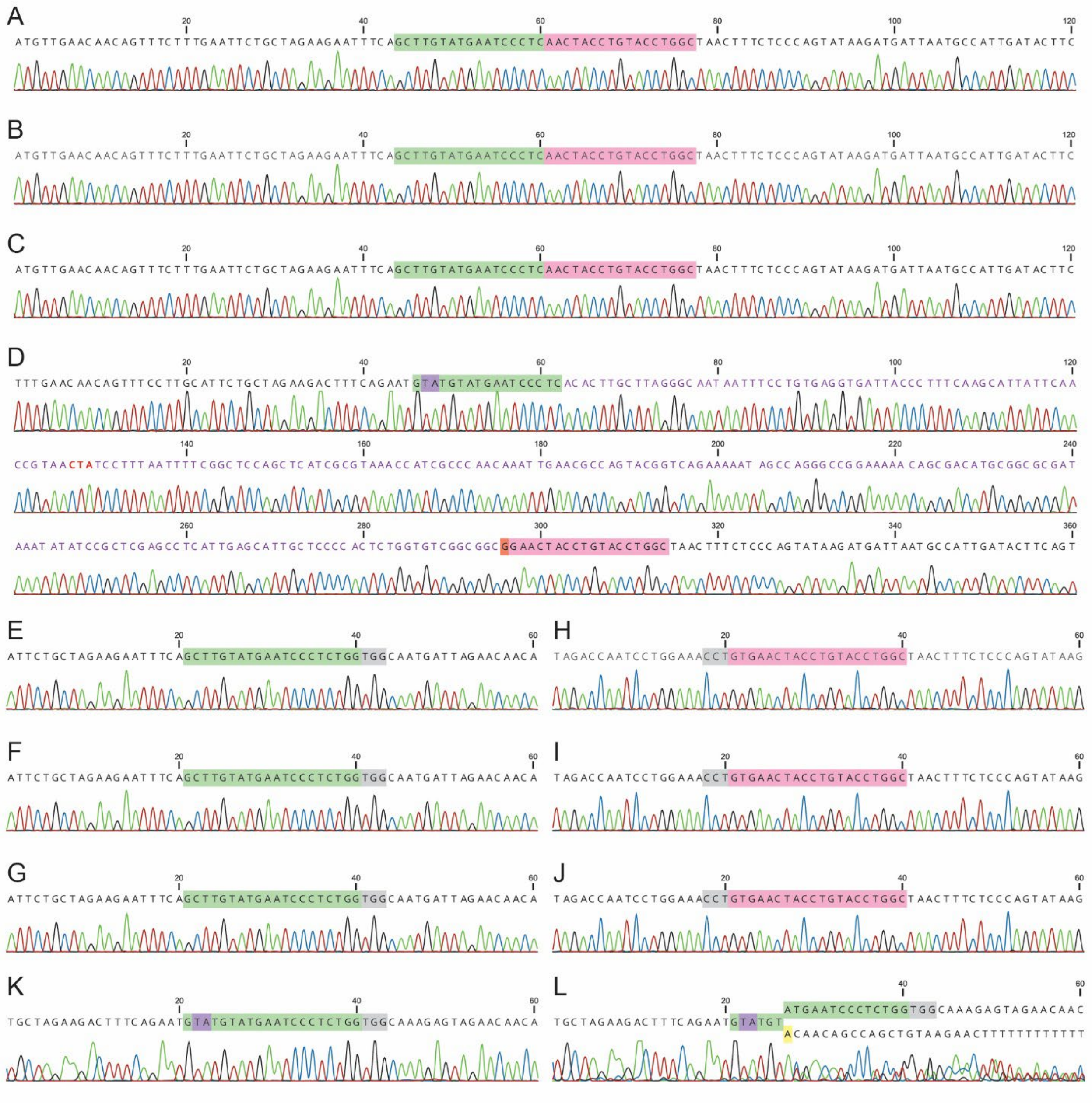

**Figure S5. Sanger sequencing of genome editing events at sgRNA sites in derivation of paternal deletion INS-1 clonal lines. (A-J)** Sanger sequence traces are shown highlighting sgRNA70-3 (green), sgRNA79-1 (pink), and SpCas9 NGG PAMs (grey). **(A-C)** Paternal (PAT)-del allele of clonal line 19-1 **(A)**, 19-4 **(B)**, and 25 **(C)** are identical canonical deletions with 3.143 Mb deleted between the sgRNA70-3 and sgRNA79-1 PAM-3 nuclease sites. Deletion-PCR breakpoint fragments are from gels such as shown in **Fig. 1E**. **(D)** PAT-del line 3 generated an unexpected proximal deletion breakpoint at the canonical PAM-3 position of an alternate (alt) sgRNA70-3 site upstream of *Frat3*, that maps ~ 125 kb upstream of the sgRNA70-3 position (see **Fig. 1A**; **Fig. S4A**).

The alt-sgRNA70-3 site differs only at 2 of the most 5′ nucleotides of the sgRNA (purple highlight) and is part of a segmental-duplication (to be described elsewhere; manuscript in preparation). The line 3 genome-editing event with a 3.268 Mb deletion has at the breakpoint a 233- or 234-bp insertion of sequence with 97% identity to *E. coli* (purple sequence, with *gsiD* amber stop codon indicated in red), followed by a distal deletion breakpoint with a single G/C nucleotide insertion or polymorphism in *E. coli* sequence (orange highlight) that occurred at a DSB at the PAM-2 position of the distal sgRNA79-1 site (pink highlight). The deletion-PCR breakpoint fragment for the PAT-del line 3 is from the **Fig. S4G** gel. **(E-G)** Sequence of intact sgRNA70-3 site in parental INS-1 **(E)** and maternal (MAT)-deletion (del) control lines 2 **(F)** and 16 **(G)**. Sequenced PCR fragments spanning sgRNA70-3 are from the **Fig. S4N** gel. **(H-J)** Sequence of intact sgRNA79-1 site in parental INS-1 **(H)** and MAT-del control lines 2 **(I)** and 16 **(J)**. Sequenced PCR fragments spanning sgRNA79-1 are from the **Fig. S4N** gel. **(K-L)** Sanger sequence traces of the alt-sgRNA70-3 site by direct Sanger sequencing of a genomic PCR product showing an intact site in parental INS-1 **(K)**, and a hemizygous scarred allele with a 28-nt deletion (leaving 6 nt intact at the 5'-end of the alt-sgRNA70-3 sgRNA site) in clonal PAT-del line 19-1 **(L)**. Sequence from the biphasic chromatograph portion in **(L)** was determined from PCR cloning and Sanger sequencing of the intact (upper) and scarred (lower) alleles. Control lines 5-9, 2, 16, and PAT-del lines 3 and 25 have the same intact alt-sgRNA70-3 sequence as INS-1 indicating homozygosity for the control lines and PAT-del line 25 while for PAT-del line 3 this indicates an intact MAT-allele at alt-sgRNA70-3 since that site is involved in the PWS-deletion breakpoint (see **Fig. S5D**). As expected, PAT-del line 19-4 has equivalent results to line 19-1, with heterozygosity for an identical scarred allele.

**Figure S6. Copy number of PWS-loci based on droplet-digital PCR (ddPCR) using EvaGreen for PWS *vs.* control INS-1 lines.**

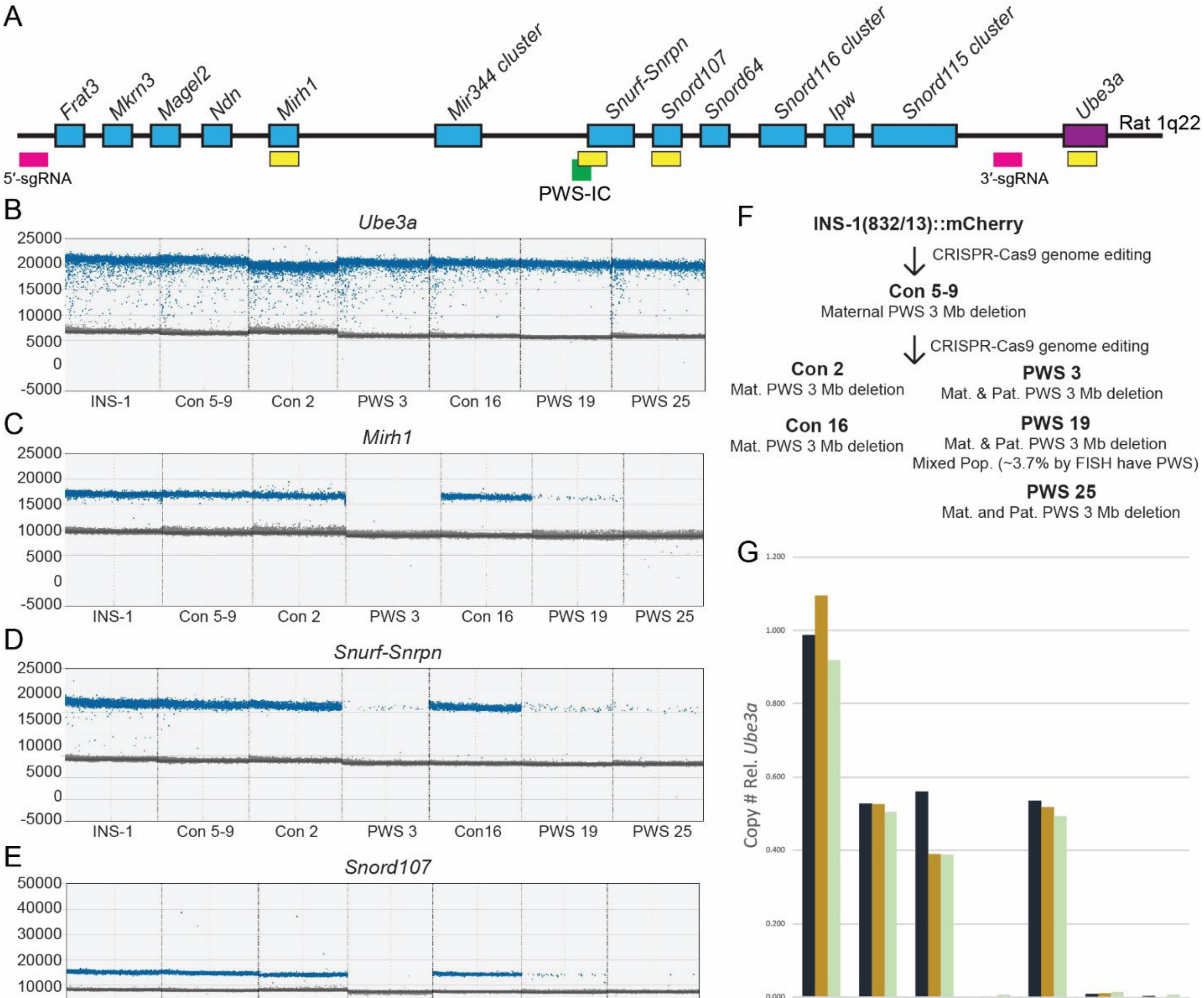

| | INS-1 | Con 5-9 | Con 2 | PWS 3 | Con 16 | PWS 19 | PWS 25 |
|---|---|---|---|---|---|---|---|
| *Snord107* | 0.988 | 0.528 | 0.561 | 0.000 | 0.535 | 0.008 | 0.000 |
| *Mirh1* | 1.095 | 0.525 | 0.390 | 0.000 | 0.519 | 0.010 | 0.000 |
| *Snrpn* | 0.918 | 0.505 | 0.387 | 0.006 | 0.494 | 0.014 | 0.006 |

**Figure S6. Copy number of PWS-loci based on droplet-digital PCR (ddPCR) using EvaGreen for PWS *vs.* control INS-1 lines. (A)** Schematic of the PWS-imprinted domain with paternally expressed genes in blue and the maternally expressed *Ube3a* in purple. The positions of the sgRNAs that mark the PWS-deletion breakpoints are indicated by pink boxes, the PWS-IC by a green box, and the four loci examined by ddPCR by yellow boxes. **(B)** Genomic ddPCR 1d amplitude plot for *Ube3a*, localized outside of the distal PWS-deletion breakpoint and hence intact in all INS-1 cell lines. **(C-E)** Genomic ddPCR 1d amplitude plots for **(C)** *Mirh1*, **(D)** *Snurf-Snrpn*, and **(E)** *Snord107*, each localized within the PWS-deletion region. **(F)** Schematic showing generation in an initial CRISPR/Cas9 genome editing screen of clonal control (Con) cell line 5-9 with an ~ 3 Mb deletion of the PWS-domain on the maternal allele, and additional clonal cell lines generated from a second CRISPR/Cas9 genome editing screen. The latter include two further control lines,

2 and 16, each with a PWS-domain deletion on the maternal allele, and three independent homozygous deletion sublines 3, 19 and 25. **(G)** Graph of genomic ddPCR for the PWS-region demonstrating that sublines 3 and 25 are pure populations of cells with a homozygous PWS-region deletion. However, whereas most cells in subline 19 have a homozygous PWS-deletion, there is a small percentage of cells having an intact PWS-region (i.e., derived from the 5-9 parental line) as a mixed population. FISH also confirmed 3.7% of cells with an intact PWS-region in subline 19 (**Fig. S4J**). Note that subline 19 then had a further screen by plating single cells in a 96-well plate and selecting clonal lines with a homozygous PWS-deletion (i.e., 19-1, 19-2, 19-3, 19-4, 19-5).

**Figure S7. Sanger sequencing of sgRNA70-3 off-target sites in maternal deletion and paternal deletion INS-1 clonal lines.**

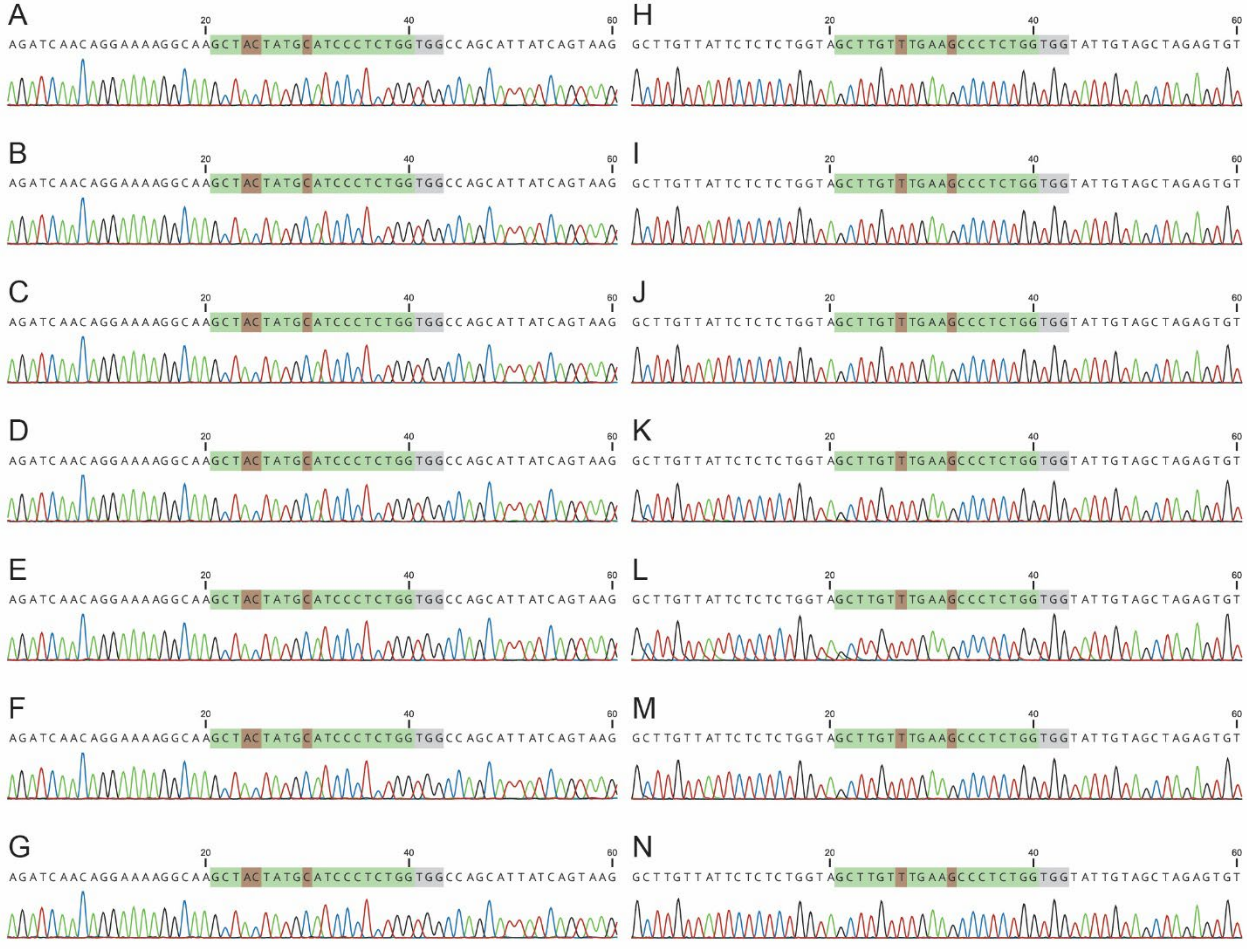

**Figure S7. Sanger sequencing of sgRNA70-3 off-target sites in maternal deletion and paternal deletion INS-1 clonal lines. (A-N)** Sanger sequence traces are shown for the top 2 ranked predicted off-target sites for sgRNA70-3 highlighted by the sgRNA seed (green), SpCas9 NGG PAMs (grey), and deviation in off-target from sequence from sgRNA70-3 (brown). **(A-G)** Sequence of intact sgRNA70-3 off-target site at chromosome 1 position 178,678,292-178,678,314 (-; intron *Hs3st4*) with 3 mismatches in parental INS-1 **(A)** and maternal (MAT)-deletion lines 5-9 **(B)**, line 2 **(C)**, line 16 **(D)**, and paternal (PAT)-deletion lines 3 **(E)**, 19-1 **(F)** and 19-4 **(G)**. **(H-N)** Sequence of intact sgRNA70-3 off-target site at chromosome 8 position 72,506,613-72,506,635 (+; intron *Tcf12*) with 2 mismatches in parental INS-1 **(H)** and MAT-deletion lines 5-9 **(I)**, line 2 **(J)**, line 16 **(K)**, and PAT-deletion lines 3 **(L)**, 19-1 **(M)** and 19-4 **(N)**.

**Figure S8. Sanger sequencing of sgRNA79-1 off-target sites in maternal deletion and paternal deletion INS-1 clonal lines.**

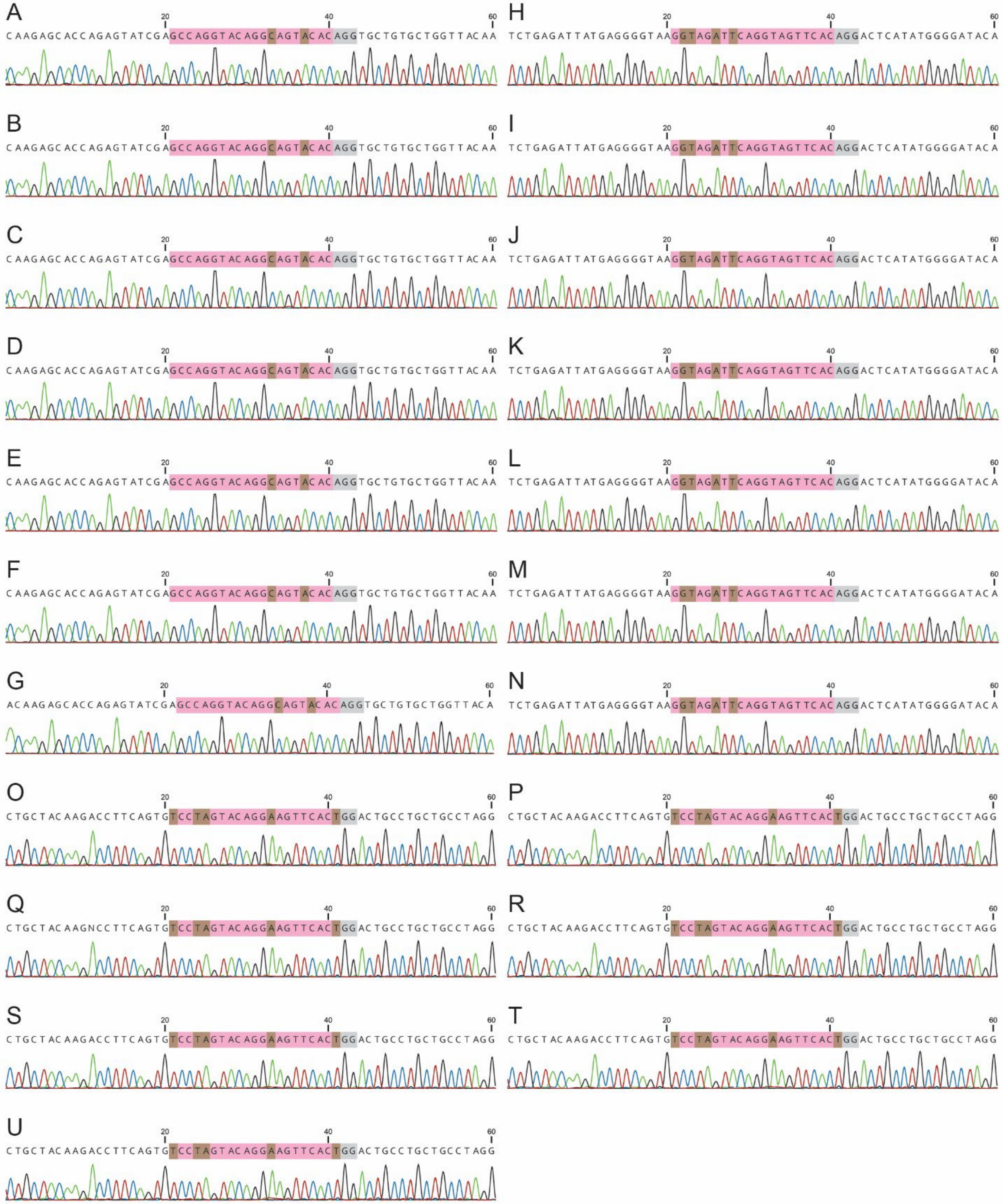

**Figure S8. Sanger sequencing of sgRNA79-1 off-target sites in maternal deletion and paternal deletion INS-1 clonal lines. (A-U)** Sanger sequence traces are shown for the top 3 ranked predicted off-target sites for sgRNA79-1 highlighted by the sgRNA seed (pink), SpCas9 NGG PAMs (grey), and deviation in off-target from sequence from sgRNA79-1 (brown). **(A-G)** Sequence of intact sgRNA79-1 off-target site at chromosome 9 position 58,847,915-58,847,937 (+; intergenic *Satb2*-RGD1306941) with 2 mismatches in parental INS-1 **(A)** and maternal (MAT)-deletion lines 5-9 **(B)**, line 2 **(C)**, line 16 **(D)**, and paternal (PAT)-deletion lines 3 **(E)**, 19-1 **(F)** and 19-4 **(G)**. **(H-N)** Sequence of intact sgRNA79-1 off-target site at chromosome 3 position 128,425,398-128,425,420 (+; intron *Macrod2*) with 4 mismatches in parental INS-1 **(H)** and MAT-deletion lines 5-9 **(I)**, line 2 **(J)**, line 16 **(K)**, and PAT-deletion lines 3 **(L)**, 19-1 **(M)** and 19-4 **(N)**. **(O-U)** Sequence of intact sgRNA79-1 off-target site at chromosome 2 position 47,166,018-47,166,040 (+; intergenic *Itga1*-*Isl1*) with 4 mismatches in parental INS-1 **(O)** and MAT-deletion lines 5-9 **(P)**, line 2 **(Q)**, line 16 **(R)**, and PAT-deletion lines 3 **(S)**, 19-1 **(T)** and 19-4 **(U)**.

**Figure S9. Copy number of PWS-loci based on droplet-digital PCR (ddPCR) using TaqMan probes for PWS *vs.* control INS-1 lines.**

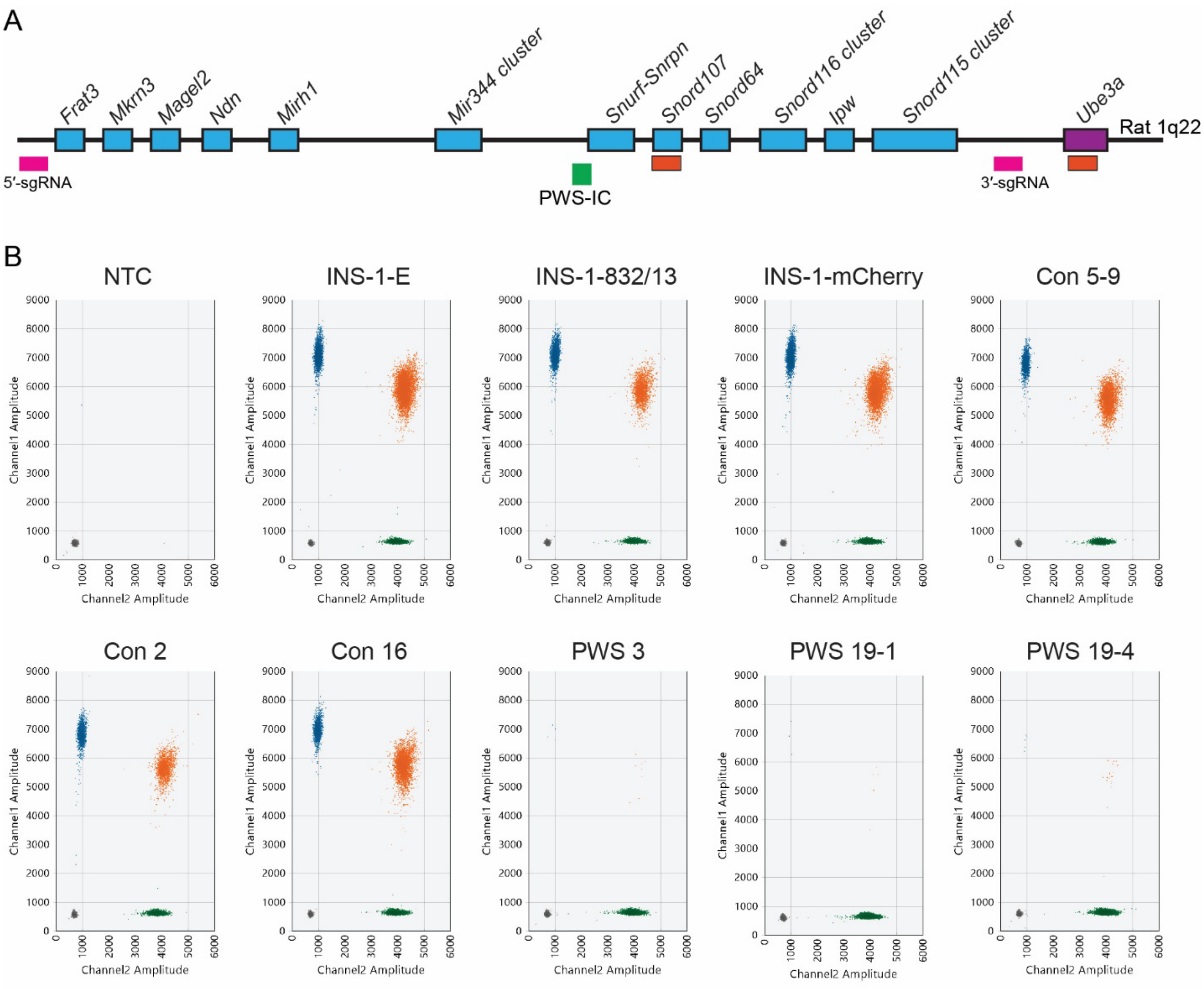

**Figure S9. Copy number of PWS-loci based on droplet-digital PCR (ddPCR) using TaqMan probes for PWS *vs.* control INS-1 lines. (A)** Schematic of the PWS-imprinted domain with paternally expressed genes in blue and the maternally expressed *Ube3a* in purple. The positions of the sgRNAs that mark the PWS-deletion breakpoints are indicated by pink boxes, the PWS-IC by a green box, and the probes used for two loci that were examined by TaqMan ddPCR by orange boxes. **(B)** ddPCR 2d plots for TaqMan probe copy number assay with absorbance amplitude for channel 1 (*Snord107*, FAM) on the y-axis and channel 2 (*Ube3a*, HEX) on the x-axis. Blue dots denote *Snord107* positive droplets, green dots represent *Ube3a* positive droplets and orange indicate double positive droplets. Note the absence of channel 1 *Snord107* positive (blue or orange droplets) in the PWS lines, and 50% reduction of channel 1 positive droplets in maternal-deletion control (Con) lines (also see **Fig. 1F** for graphical data).

**Figure S10. Gene expression for PWS-imprinted genes, *Ube3a* and *Ube3a-ATS* loci.**

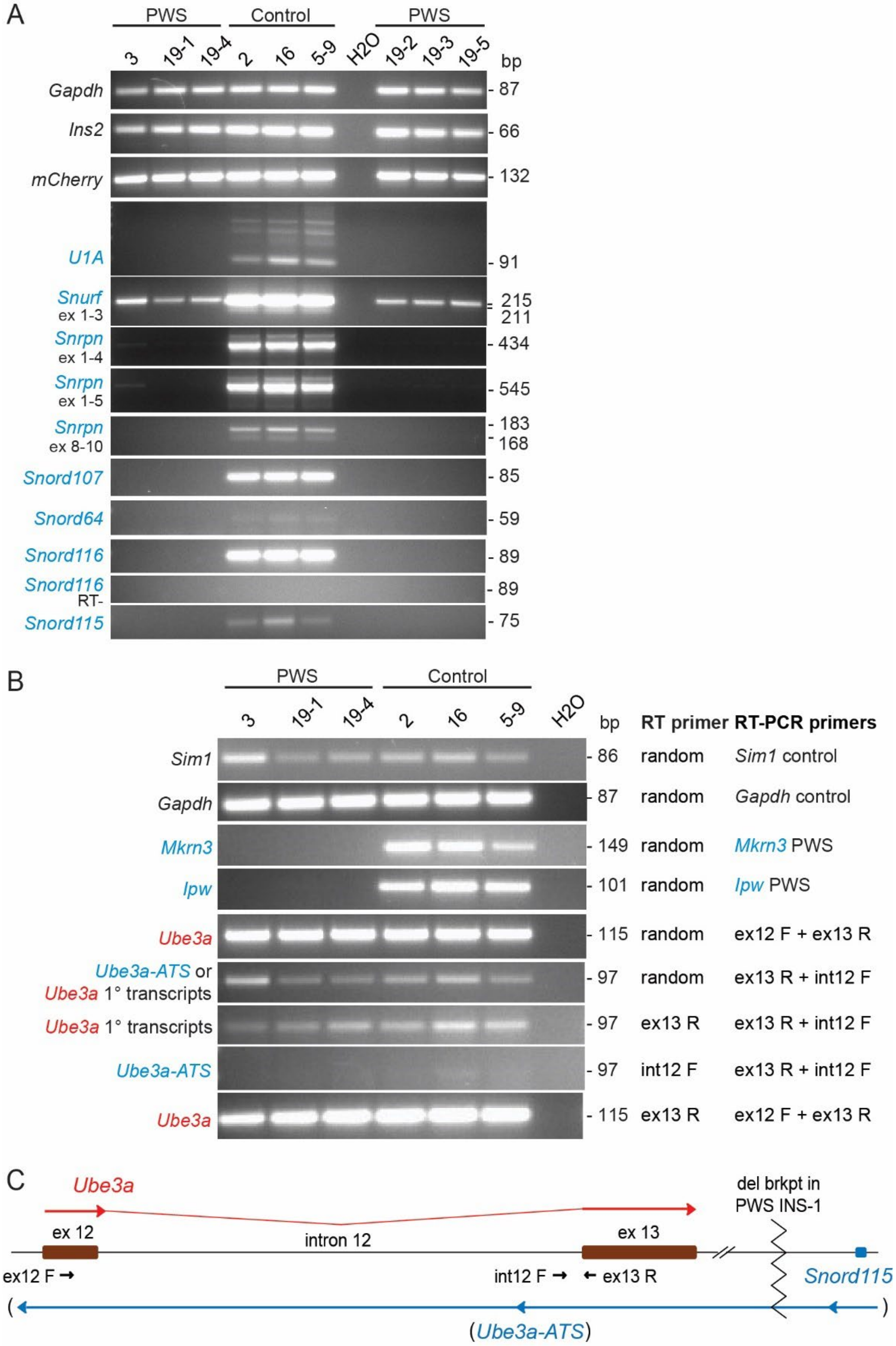

**Figure S10. Gene expression for PWS-imprinted genes, *Ube3a* and *Ube3a-ATS* loci.** **(A)** RT-PCR analyses of 7 PWS-imprinted genes (*U1A*, *Snurf*, *Snrpn*, *Snord107*,

*Snord64*, *Snord116*, *Snord115*; see map in **Fig. 1A**), rat *Ins2*, *mCherry* transgene, and *Gapdh* control gene in the expanded INS-1 panel of 9 cell lines. U1 represents an alternate upstream (U) promoter-first exon that splices into *Snurf*-*Snrpn* exon 2 and has multiple duplicate U1 copies in rodents. Abbreviations: ex, exon; RT-, PCR control using RNA not treated with reverse-transcriptase. The absence of bands in the RT- assay for the multicopy, tandemly repeated *Snord116* locus rules out genomic DNA contamination in the RNA. Note that the RT-PCR gel results for 4 genes (*Gapdh*, *Ins2*, *Snrpn* ex 8-10, *Snord116*) in the 6-cell line panel used in this entire study are reproduced from **Fig. 1G** for direct comparison to the remaining genes in the PWS domain that are presented here, plus this dataset has 3 additional clonal PWS INS-1 cell lines (19-2, 19-3, 19-5). Similarly, while the *Snurf* ex 1-3 results are shown here for the expanded panel of 9 INS-1 cell lines, these same results for the 6-cell line panel used in this entire study are reproduced in **Fig. S11B** for comparison to other loci examined in that dataset. **(B)** RT-PCR analyses of 2 PWS-imprinted genes (*Mkrn3*, *Ipw*) as well as *Ube3a* and *Ube3a-ATS* loci (see maps in **Fig. 1A**; **Fig. S10C**), and control genes (*Gapdh*, *Sim1*) in the 6-cell line panel used in this entire study. The upper 6 rows of gels are from standard RT-PCR assays that use random hexamer primers for the RT primer and typical gene-specific RT-PCR primers, while the lower 3 rows of gels are from RT-PCR assays that use a strand-specific primer for RT followed by RT-PCR with either a *Ube3a* (ex12 F + ex13 R) or *Ube3a-ATS* (ex13 R + int12 F) primer set. Combined, the data show that transcripts from the *Ube3a-ATS*-region that are detected when using random primers for RT derive from *Ube3a* primary (1°) transcripts since there is no expression from the *Ube3a-ATS* strand. **(C)** Map of the *Ube3a* and *Ube3a-ATS* locus. Long arrows represent transcriptional orientation and extent, with *Ube3a* spliced (exon 12-13), and *Ube3a-ATS* transcribed as the 3'-end of long lncRNAs from the *U1A*-*Snurf*-*Snrpn*-snoRNA locus. While *Ube3a* is highly expressed in all INS-1 cell lines, as seen in **Fig. S10B** there is no detectable expression of *Ube3a-ATS* in control or PWS cell lines. Abbreviations: del brkpt, CRISPR/Cas9 deletion breakpoint in PWS INS-1 cell lines; ex, exon; int, intron; short arrows, PCR primers; parentheses [(,)] indicate the lack of *Ube3a-ATS* transcripts.

**Figure S11. Expression of a ψ*Snurf*-ψ*Snrpn* locus within the *Mon2* gene.**

A

| gene_ID | gene_name | Control_5-9 | Control_2 | Control_16 | PWS_3 | PWS_19-1 | PWS_19-4 | log2FoldChange | pvalue | padj |
|---|---|---|---|---|---|---|---|---|---|---|
| ENSRNOG00000054391 | *Snurf* | 3992.4473 | 3159.5016 | 3911.1605 | 2.041794 | 0 | 3.6367 | -10.80977381 | 6.74E-72 | 6.61E-68 |
| ENSRNOG00000059764 | *Snrpn* | 3956.3681 | 3149.4044 | 3874.1853 | 2.041794 | 0 | 3.6367 | -10.79883928 | 8.04E-72 | 6.61E-68 |
| Rnor_G_Custom_004_psSnurfSnrpn | *psSnurfSnrpn* | 502.26031 | 362.5808 | 470.40744 | 27.56422 | 43.04738 | 45.4587 | -3.522981509 | 1.35E-51 | 7.37E-48 |
| ENSRNOG00000004185 | *Mon2* | 3130.3445 | 2880.452 | 3272.3102 | 3161.719 | 3215.4 | 3141.2 | 0.036196296 | 0.671107 | 0.99282 |

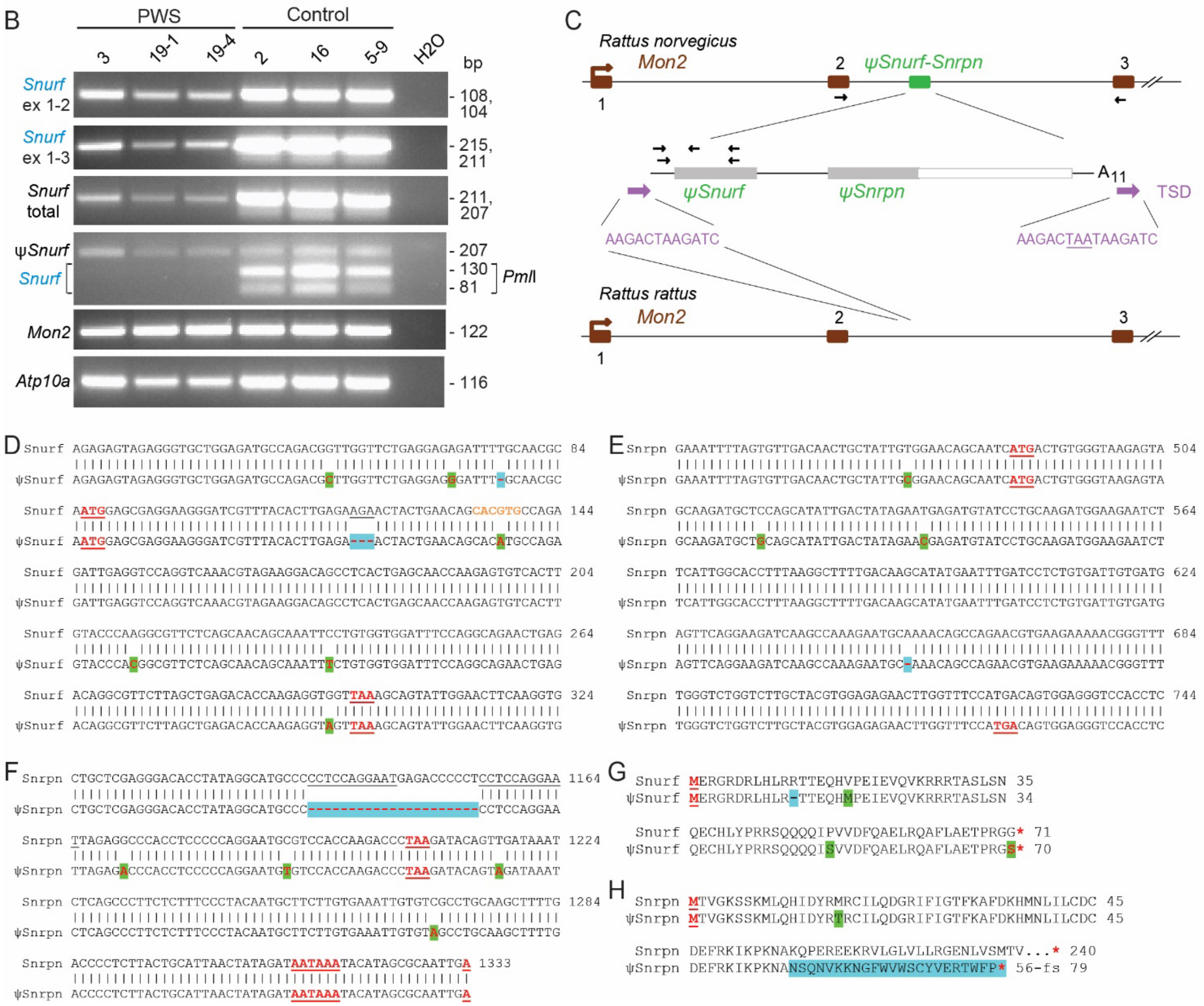

**Figure S11. Expression of a ψ*Snurf*-ψ*Snrpn* locus within the rat *Mon2* gene. (A)** RNA-seq analysis using a custom rat genome build identifies a ψ*Snurf*-*Snrpn* locus expressed in PWS and control INS-1 lines. In contrast, the imprinted *Snurf* and *Snrpn* loci are only expressed in the control INS-1 lines. Expression for the pseudogene host gene, *Mon2*, is also shown. **(B)** RT-PCR with gel analysis for 3 amplicons from *Snurf* as well as ψ*Snurf*-ψ*Snrpn*, *Mon2* and *Atp10a* in the INS-1 panel. The first 2 rows show primer sets designed to amplify *Snurf* exons (ex) 1-2 and 1-3, while row 3 shows RT-PCR with a primer set designed to amplify both *Snurf* and ψ*Snurf* sequences; the latter RT-PCR products were then digested with the *Pml*I restriction endonuclease which distinguishes between *Snurf* and ψ*Snurf* products. Note that *Atp10a* is significantly reduced in PWS *vs.* control lines (also see **Table S1**,**S2**). **(C)** Map location of the expressed ψ*Snurf*-ψ*Snrpn* locus (green box) within the brown rat (*Rattus norvegicus*) *Mon2* gene (brown boxes are 5' exons 1-3). In contrast, the black rat (*Rattus rattus*) has no pseudogene insertion and a single copy of the Target Site Duplication (TSD, purple arrows) present in the brown rat genome at the 5'-end of the inserted pseudogene. Black arrows represent PCR primers, while the underlined TAA nucleotides represents a tandem duplication in the 3'-TSD copy. **(D)** DNA sequence of ψ*Snurf* transcripts in PWS INS-1 lines compared to the endogenous *Snurf* gene. Green shading of red nucleotides indicates missense mutations, and blue shading with a hyphen represents a nucleotide deletion. Orange font represents the *Pml*I cleavage site specifically in the *Snurf* cDNA sequence. Snurf start and stop codons are indicated. **(E)** DNA sequence identifies a frameshift from a single nucleotide deletion and premature stop codon in the 5' ψ*Snrpn* portion of the pseudogene. Symbols as for **(D)**. **(F)** DNA sequence identifies a 21-nucleotide deletion in the 3' ψ*Snrpn* portion of the pseudogene. Symbols as for **(D)**. **(G)** Potential ψSnurf amino acid sequence encoded by the ψ*Snurf* gene. Green or blue highlight represents potential amino acid changes or in-frame deletion, respectively, and red asterisks the stop codons. **(H)** Potential truncated ψSmN amino acid sequence encoded by the ψ*Snrpn* gene. Symbols as for **(G)**, except blue highlight, amino acids resulting from a frame-shift mutation.

Assuming no evolutionary selection for function, we can date the age of rat ψ*Snurf*-ψ*Snrpn* based on a total of 26 mutations/substitutions in 1,288-nt of homologous sequence to rat *Snurf*-*Snrpn*. The 2.02% divergence (97.98% similarity) corresponds using a mutation rate formula for silent site substitutions and intronic sequences of $5 \times 10^{-9}$ to $7 \times 10^{-9}$ mutations/site/year [1] to a pseudogene origin ~ 2.88-4.0 million years ago (mya), which correlates well with prior molecular dates for divergence of black and brown rats about 2.0-2.9 mya [2; 3].

**Figure S12. Genome-wide snoRNA and miRNA transcriptome changes in PWS INS-1 lines from small RNA-seq.**

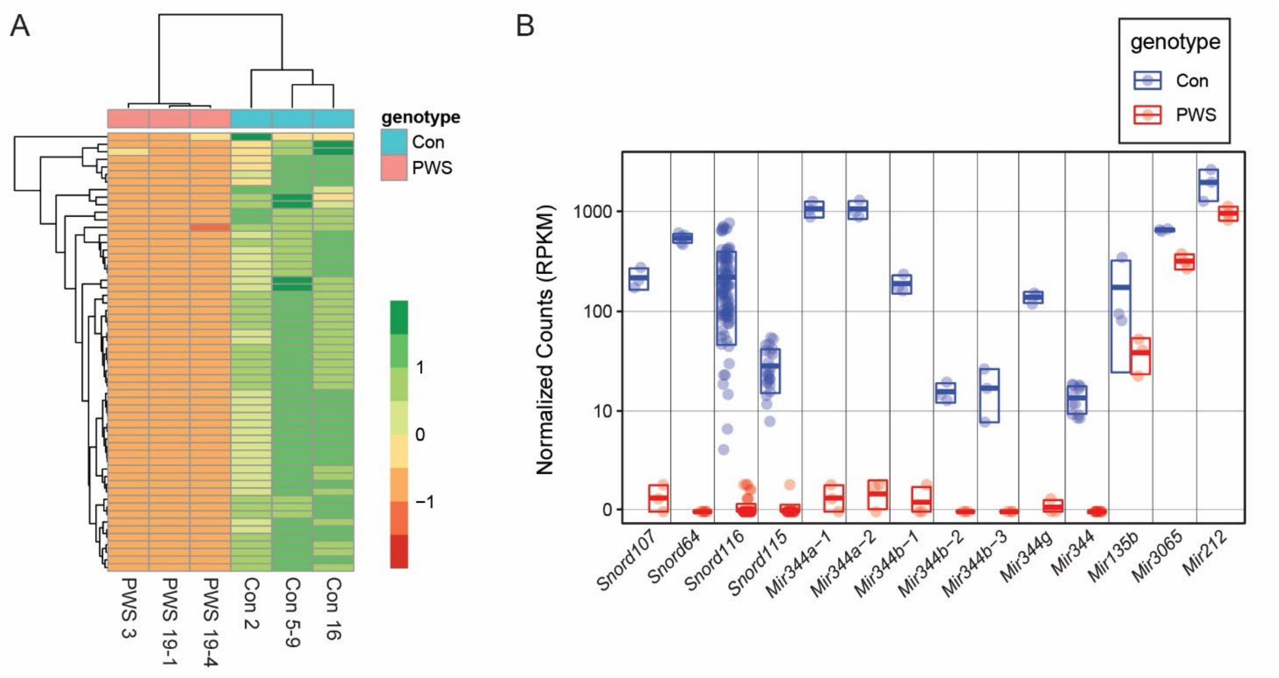

**Figure S12. Genome-wide snoRNA and miRNA transcriptome changes in PWS INS-1 lines from small RNA-seq. (A)** Heatmap clustergram of 58 differentially expressed miRNAs and snoRNAs demonstrates tight clustering of PWS *vs.* control groups (*P*adj < 0.1). Scale: green (enriched) to red (depleted). Small RNA-seq was performed for 3 PWS (3, 19-1, 19-4) *vs.* 3 control (5-9, 2, 16) INS-1 cell lines. **(B)** Normalized expression counts for the top 14 significant differentially expressed miRNAs and snoRNAs, of which only 3 miRNAs identified are not encoded within the PWS-imprinted domain. Box charts of control (blue) and PWS (red) genotypes are shown with underlying data points for each sample. For the multicopy *Snord115*, *Snord116*, and *Mir344* genes the data for the paralogs were binned.

**Figure S13. Gene ontology enrichment of DEGs.**

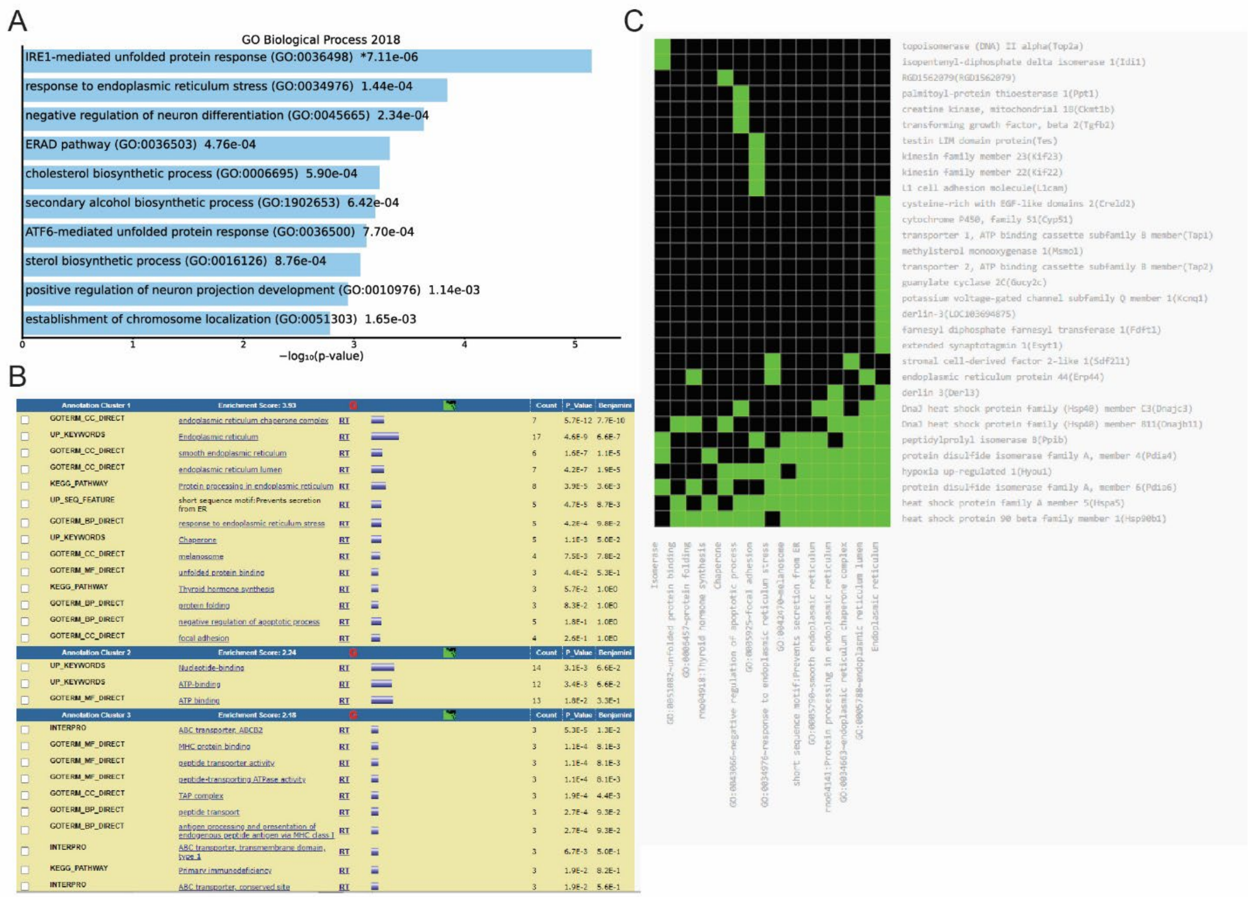

**Figure S13. Gene ontology enrichment of DEGs. (A)** EnrichR gene ontology biological processes enriched in down-regulated DEGs highlighting ER stress mediators including IRE1, ERAD and ATF6 pathways. **(B)** DAVID analysis reveals an annotation grouping ascribed with ER functions and **(C)** many of the components of the UPR clustered together. Analysis performed with down regulated DEGs from adjusted *P* value < 0.1, and fold change decrease of less than -1.25 with all PWS genes removed. Abbreviations: ERAD, Endoplasmic reticulum-associated protein degradation; UPR, unfolded protein response.

**Figure S14. Transcription factor binding site enrichment in the promoters of DEGs.**

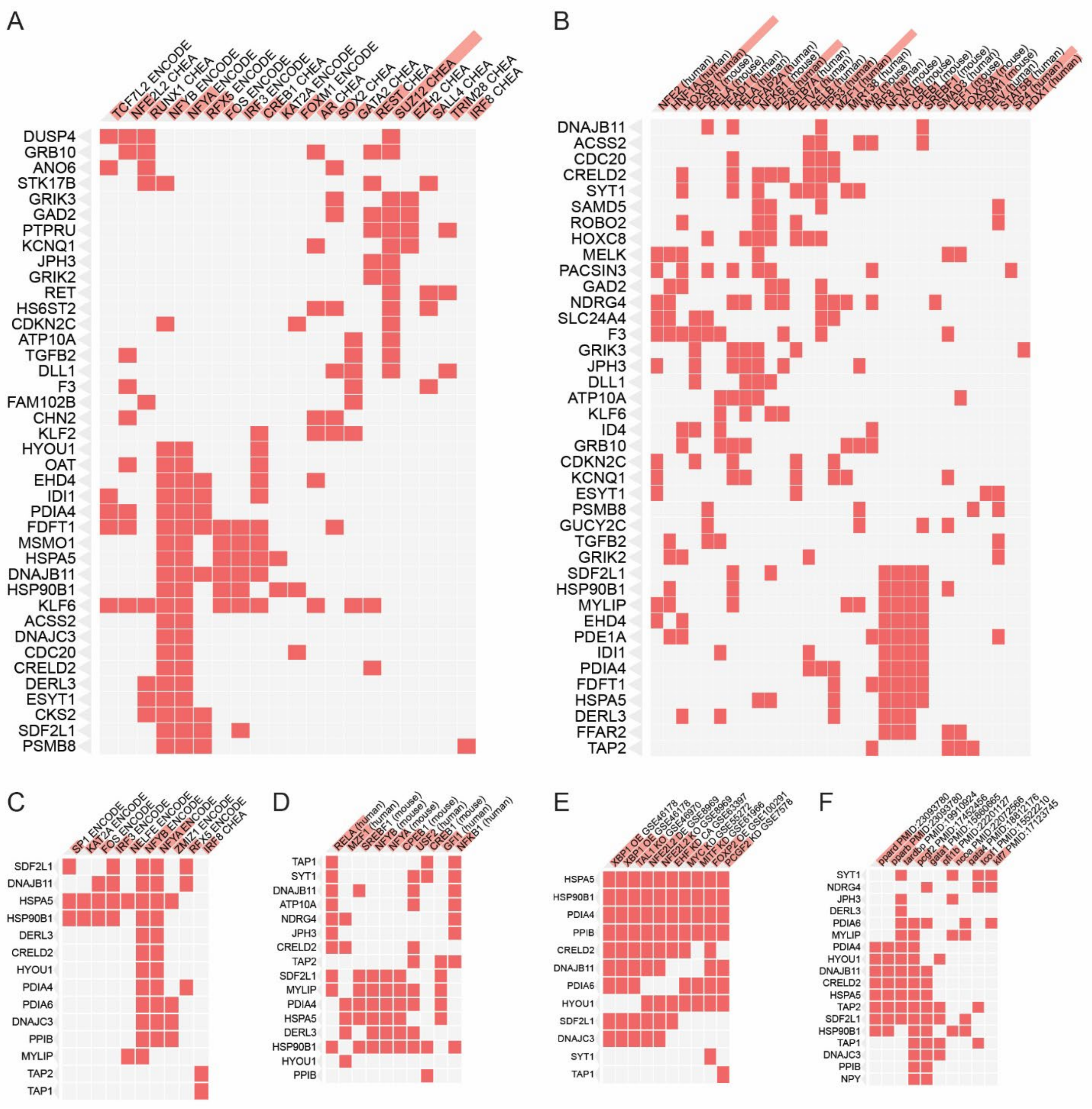

**Figure S14. Transcription factor binding site enrichment in the promoters of DEGs. (A-B)** Enrichr transcription factor (TF) analysis of 94 down-regulated non-PWS DEGs (*P*adj < 0.10 and an absolute fold-change (FC) >1.25) shown as clustergrams of **(A)** top 20 ENCODE and ChEA Consensus TFs from ChIP-X or **(B)** top 30 TRANSFAC and JASPAR PWMs enriched in the gene set. **(C-F)** Enrichr TF analysis for 20 RT-ddPCR validated down-regulated genes (from **Fig. 4E,F** and **Fig. S16**: *Hspa5*, *Hsp90b1*, *Pdia4*, *Pdia6*, *Ppib*, *Creld2*, *Sdf2l1*, *Dnajb11*, *Dnajc3*, *Hyou1*, *Npy*, *Iapp*, *Derl3*, *Mylip*, *Atp10a*, *Jph3*, *Ndrg4*, *Tap1*, *Tap2*, *Syt1*. Further analysis of validated DEGs enriched for TF binding sites from **(C)** ENCODE and ChEA Consensus TFs, **(D)** TRANSFAC and JASPAR motifs, **(E)** TF Perturbations Followed by Expression (DE, differential expression; KD, knockdown; KO, knockout; OE, over-expression) with GEO accession listed, and **(F)** TF-LOF Expression from GEO with PMID reference appended (LOF, loss of function). Clustergrams generated based on enrichment *P*-value significance which shows as the red highlight of TF names. Notably in the downregulated genes in PWS INS-1 cells a prominent cluster of genes including many ER chaperones were enriched for ATF6-cofactor NFYA and NFYB binding sites and other potential regulators including CPEB1, RFX5, IRF3, CREB1 and SREBF1 (**Fig. S14A-D**). Similarities to other model systems revealed analogous gene-expression changes including for *Xbp1* perturbations in an adipose cell line [4] and the *Pparb/d* KO mouse pancreas [5] (**Fig. S14E-F**).

**Figure S15. Specificity of RT-PCR and RT-ddPCR assays for rat, mouse and human insulin genes in the INS-1 cell line panel.**

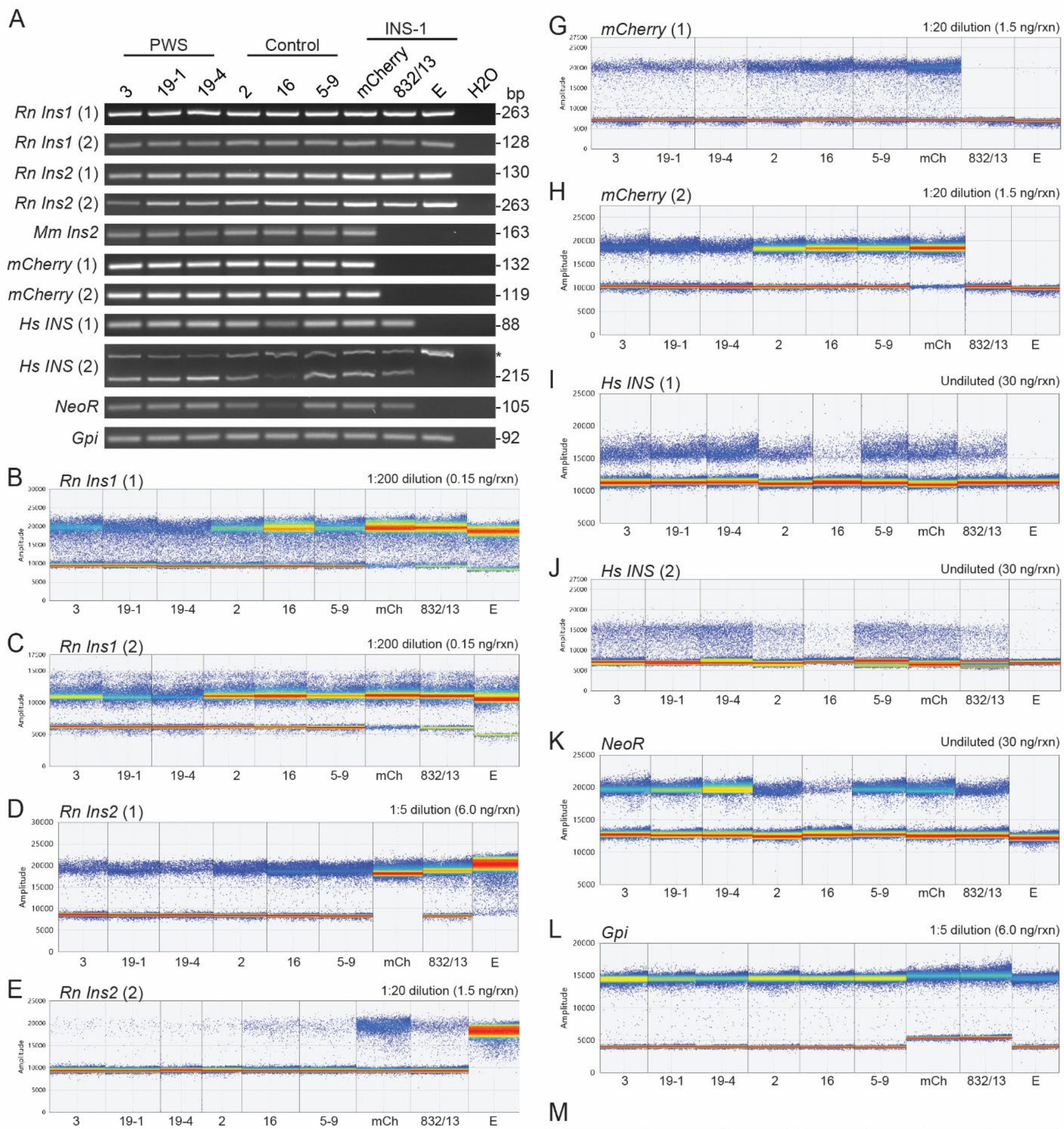

M

| gene_name | Control_5-9 | Control_2 | Control_16 | PWS_3 | PWS_19-1 | PWS_19-4 |
|---|---|---|---|---|---|---|
| *rIns1* | 256090 | 262628 | 282807 | 247273 | 175597 | 133488 |
| *mIns2mcherry* | 34169 | 38160 | 47715 | 25333 | 16273 | 13027 |
| *rIns2* | 5197 | 2310 | 7689 | 2494 | 3135 | 1516 |
| *hINS* | 79.8 | 49.6 | 3.1 | 79.6 | 57.4 | 107.3 |
| *neoR* | 178.5 | 96.4 | 4.1 | 173.6 | 162.6 | 233.7 |

**Figure S15. Specificity of RT-PCR and RT-ddPCR assays for rat, mouse and human insulin genes in the INS-1 cell line panel. (A)** RT-PCR with gel analysis for insulin genes. Abbreviations are: (1), (2), represent amplicons 1 and 2 for a given gene; mCherry, INS-1(832/13)::mCherry cell line; 832/13, INS-1(832/13) cell line; E, INS1-E cell line; *Hs*, *Homo sapiens*; *Mm*, *Mus musculus*; *Rn*, *Rattus norvegicus*; *, non-specific band in the indicated RT-PCR assay (note that this amplified product is not present using the same PCR primer pair in the RT-ddPCR assay shown in **Fig. S15J**, likely due to different chemistry or annealing temperature in the two assays). Note that control line 16 has greatly reduced expression of the *Hs INS* (amplicons 1 and 2) and *NeoR* segments of the *INS*-*NeoR* transgene, as also seen in the RT-ddPCR data [see **Fig. S15I-K**] and RNA-seq data [see **Fig. S15M**], likely reflecting epigenetic inactivation of the *INS*-*NeoR* transgene in a majority of cells for line 16 (as DNA analysis indicated the transgene remained present). **(B-L)** RT-ddPCR assays for the listed amplicons, with ddPCR performed using EvaGreen. All abbreviations are as for **Fig. S15A**. **(B)** RT-ddPCR assay for *Rn Ins1* amplicon 1. **(C)** RT-ddPCR assay for *Rn Ins1* amplicon 2. **(D)** RT-ddPCR assay for *Rn Ins2* amplicon 1. **(E)** RT-ddPCR assay for *Rn Ins2* amplicon 2. **(F)** RT-ddPCR assay for *Mm Ins2*. **(G)** RT-ddPCR assay for *mCherry* amplicon 1. **(H)** RT-ddPCR assay for *mCherry* amplicon 2. **(I)** RT-ddPCR assay for *Hs INS* amplicon 1. **(J)** RT-ddPCR assay for *Hs INS* amplicon 2. **(K)** RT-ddPCR assay for *NeoR*. **(L)** RT-ddPCR assay for control gene *Gpi*. **(M)** RNA-seq analysis of insulin genes expressed in the INS-1 lines. In addition to the endogenous rat (r) *Ins1* and *Ins2* genes, a custom rat genome build identified mouse (m) *Ins2-mCherry* and *Hs INS*-*NeoR* transgene mRNA levels. Red numbers indicate that control line 16 is an outlier with drastically reduced expression of the *Hs INS*-*NeoR* transgene (see **Fig. S15A** legend).

**Figure S16. Additional RT-ddPCR expression data for candidate differentially-expressed genes (DEGs).**

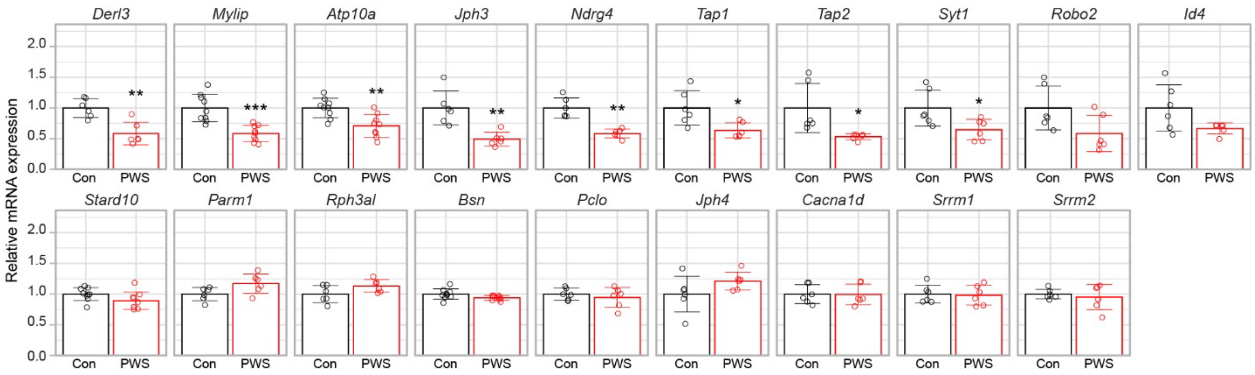

**Figure S16. Additional RT-ddPCR expression data for candidate differentially-expressed genes (DEGs).** Quantitative expression analysis of a panel of 19 additional candidate DEG validated an additional 8 down-regulated genes (*Derl3, Mylip, Atp10a, Jph3, Ndrg4, Tap1, Tap2 and Syt1*), but failed to validate any additional genes that were candidates as up-regulated in the RNA-seq data. Statistical comparison by Welch's t-test: *, $P < 0.05$; **, $P < 0.005$; ***, $P < 0.0005$.

**Figure S17. Electron microscopy establishes normal subcellular structures including ER and mitochondria in PWS and control INS-1 lines.**

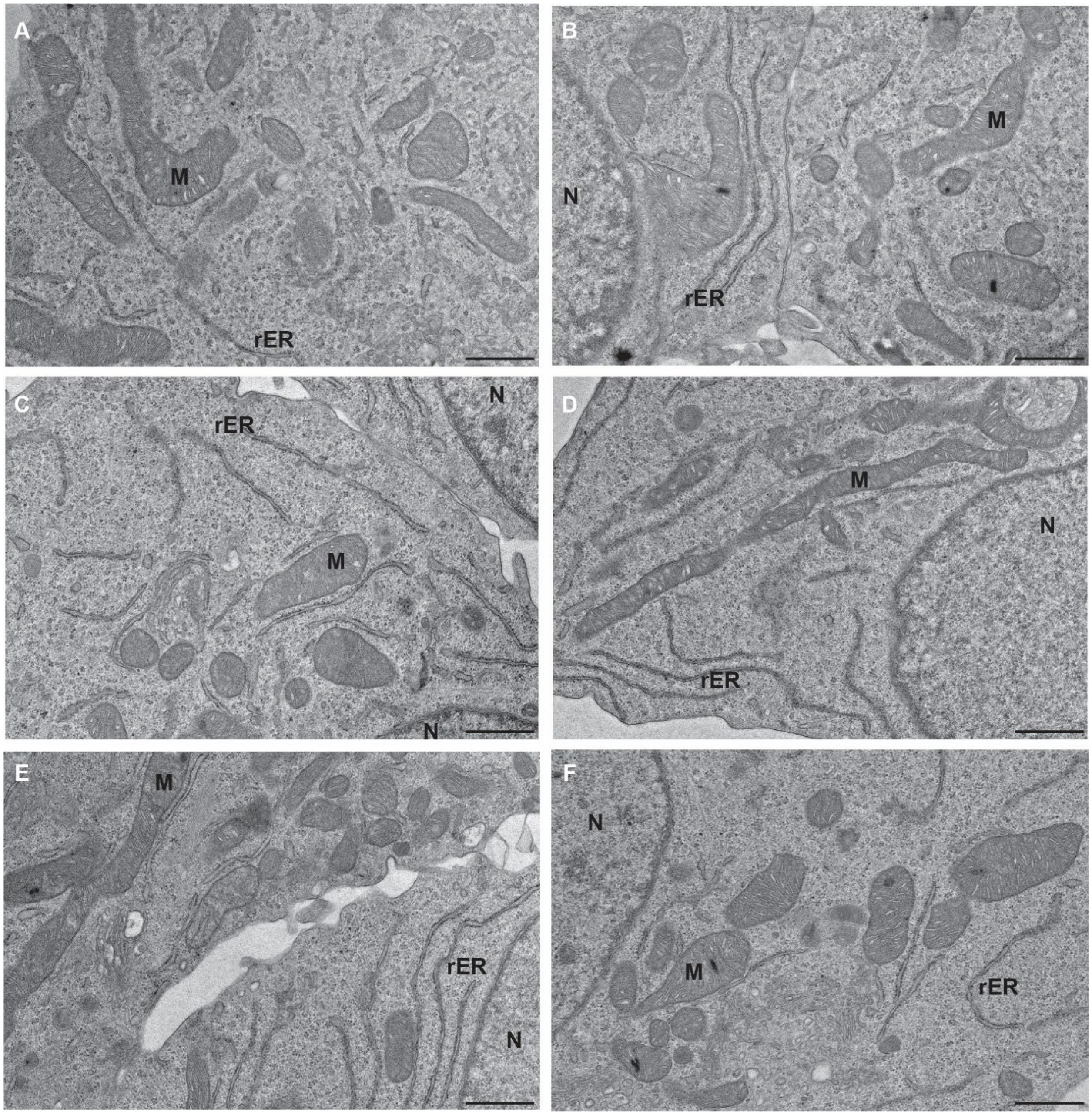

**Figure S17. Electron microscopy establishes normal subcellular organelles in PWS and control INS-1 lines. (A)** Control line 5-9, **(B)** Control line 2, **(C)** Control line 16, **(D)** PWS line 3, **(E)** PWS line 19-1, **(F)** PWS line 19-4. The scale bar in the bottom right corner of each image is 800 nm, while abbreviations for features are: N, nucleus; M, mitochondria; rER, rough endoplasmic reticulum (for M and rER, one label per image).

**Figure S18. Confocal microscopy of the processed C-peptide (CP)-mCherry in PWS and control INS-1 lines.**

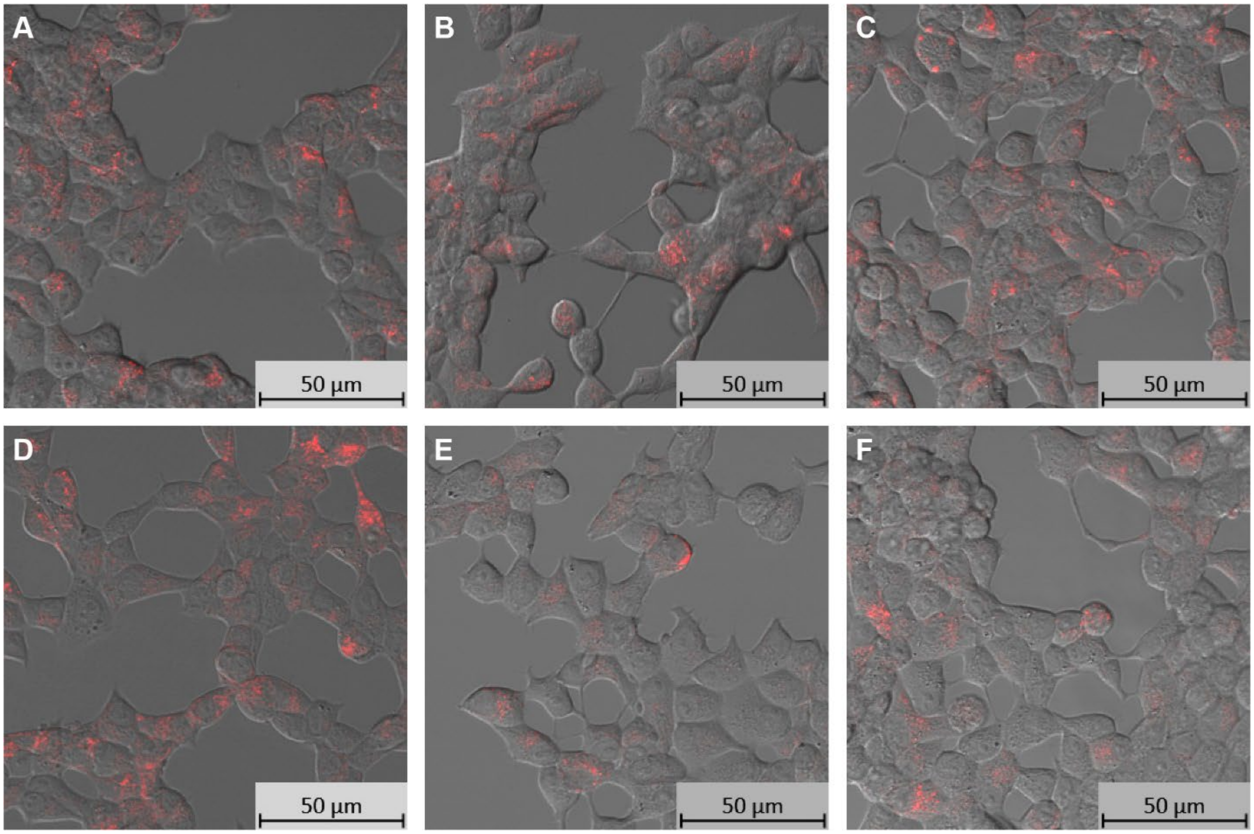

**Figure S18. Confocal microscopy of the processed C-peptide (CP)-mCherry in PWS and control INS-1 lines.** Representative live-cell confocal images obtained at 20x magnification for **(A)** Control line 5-9, **(B)** Control line 2, **(C)** Control line 16, **(D)** PWS line 3, **(E)** PWS line 19-1, **(F)** PWS line 19-4. There is no discernable difference of mCherry localization or total fluorescence between control and PWS INS-1 lines, consistent with western blot results for detection of CP-mCherry (see **Fig. 5B,E**).

**Figure S19. PWS INS-1 β-cell lines are more sensitive to thapsigargin-induced ER stress with earlier and more robust activation of *Xbp1* "splicing".**

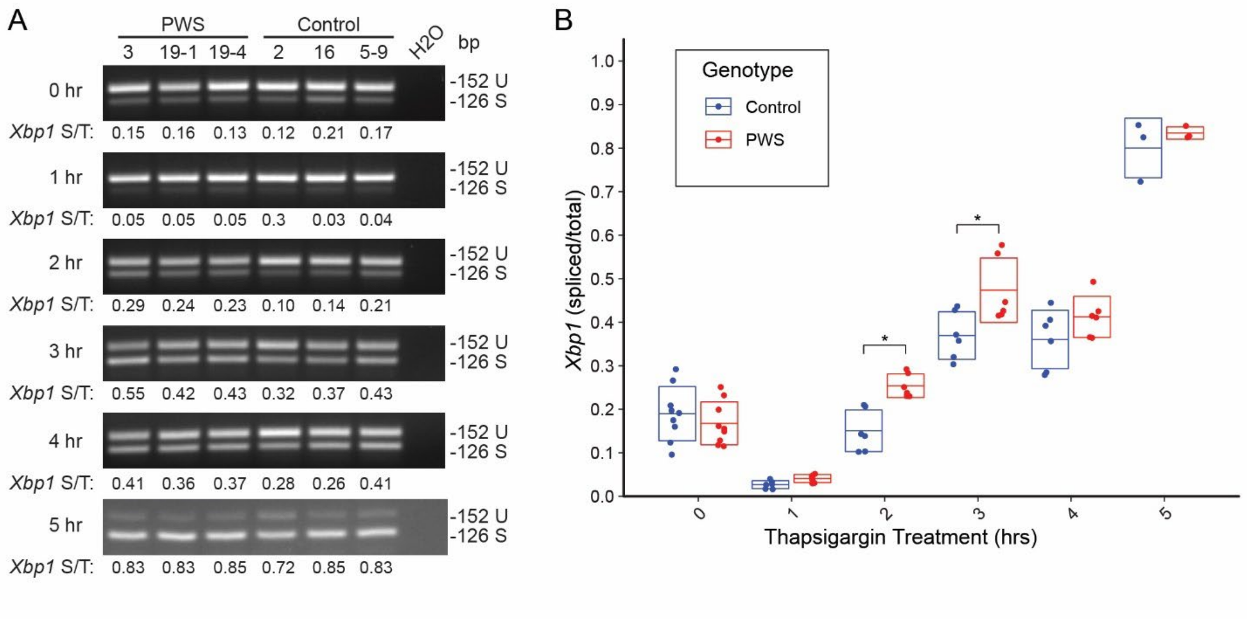

**Figure S19. PWS INS-1 β-cell lines are more sensitive to thapsigargin-induced ER stress with earlier and more robust activation of *Xbp1* "splicing". (A)** Reverse transcription-PCR gel electrophoresis analysis of *Xbp1* mRNA processing for exclusion of 26-nt of exon 4 from time 0 to 5 hours of thapsigargin treatment**.** Abbreviations: S, spliced; T, total; U, unspliced. **(B)** Time-course of *Xbp1* mRNA activation as the ratio of spliced/total mRNA detected by RT-PCR in **(A)**. Initially, at 1 hr of thapsigargin treatment *Xbp1* mRNA levels fall due to mRNA turnover for both control and PWS cell lines. *, $P < 0.05$ as calculated by ANOVA.

**Figure S20. PWS INS-1 β-cell lines are more sensitive to tunicamycin-induced ER stress with earlier and more robust activation of ATF6-N.**

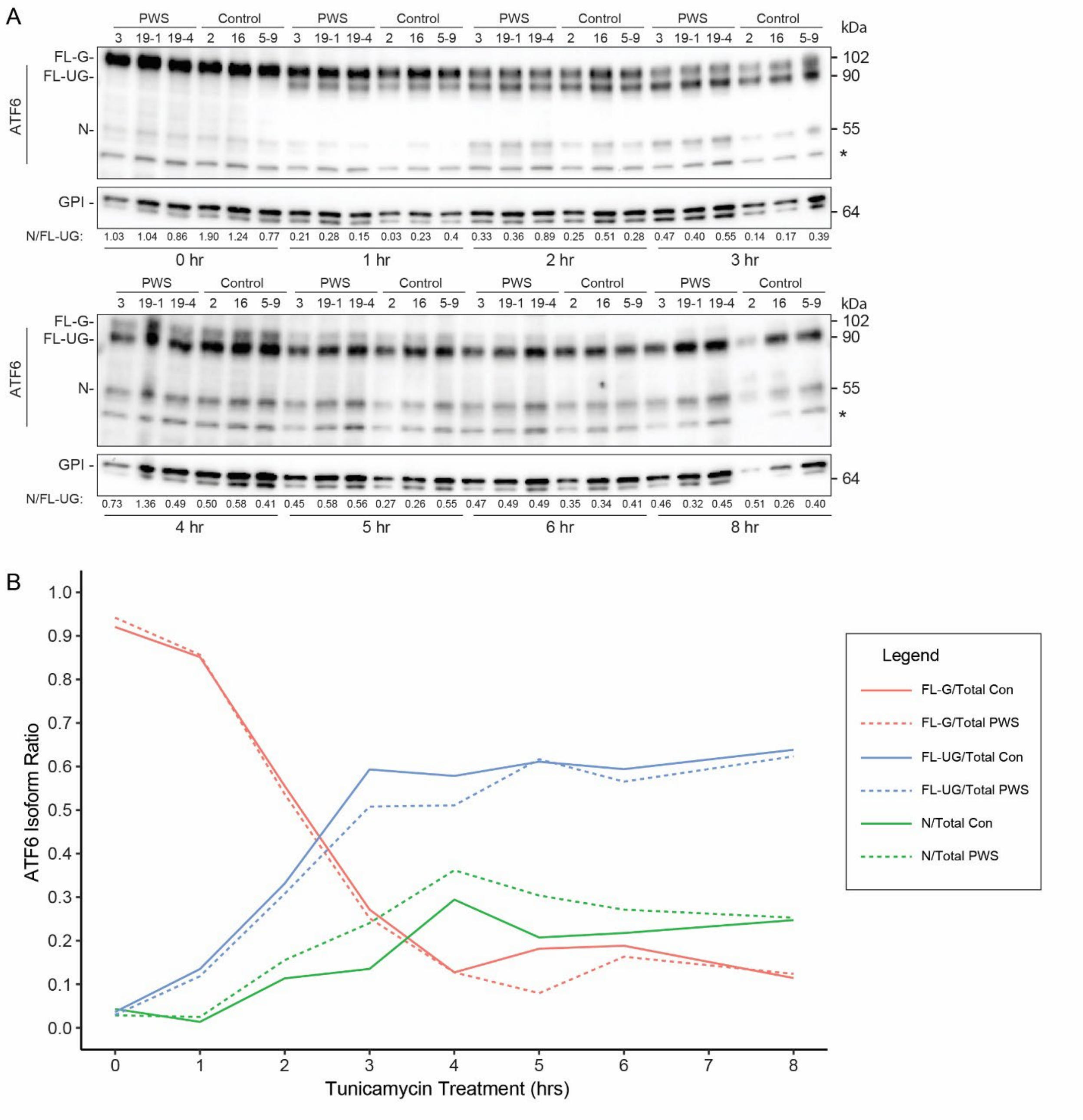

**Figure S20. PWS INS-1 β-cell lines are more sensitive to tunicamycin-induced ER stress with earlier and more robust activation of ATF6-N. (A)** Time-course western blots of ATF6 in PWS and control cell lines treated with tunicamycin for 0, 1, 2, 3, 4, 5, 6 or 8 hours. The ratio of processed nuclear (N) isoform over full-length unglycosylated (FL-UG) is written below each lane. FL-G, full-length glycosylated; *, non-specific band detected by the anti-ATF6 antibody. **(B)** Relative ratio of each ATF6 isoform as a fraction of the total ATF6 during the time-course of tunicamycin treatment. Line graphs represent mean of three measurements at each time point, except n=6 using technical replicates for 4 hr and n=6 using two biological replicates for 5 hr timepoints, for full-length glycosylated (FL-G) over total (Red); FL-UG over total (Blue) and N over total (Green) with control (Con) as solid lines and PWS cell lines as dotted lines. Data indicates that there is more robust processing of FL-UG to N at early tunicamycin timepoints in PWS cell lines.

### SUPPLEMENTARY DATA, REFERENCES

[1] Eichler, E.E., Lu, F., Shen, Y., Antonacci, R., Jurecic, V., Doggett, N.A., et al., 1996. Duplication of a gene-rich cluster between 16p11.1 and Xq28: a novel pericentromeric-directed mechanism for paralogous genome evolution. Hum Mol Genet 5(7):899-912.

[2] Verneau, O., Catzeflis, F., Furano, A.V., 1998. Determining and dating recent rodent speciation events by using L1 (LINE-1) retrotransposons. Proc Natl Acad Sci U S A 95(19):11284-11289.

[3] Robins, J.H., McLenachan, P.A., Phillips, M.J., Craig, L., Ross, H.A., Matisoo-Smith, E., 2008. Dating of divergences within the Rattus genus phylogeny using whole mitochondrial genomes. Mol Phylogenet Evol 49(2):460-466.

[4] Gregor, M.F., Misch, E.S., Yang, L., Hummasti, S., Inouye, K.E., Lee, A.H., et al., 2013. The role of adipocyte XBP1 in metabolic regulation during lactation. Cell Rep 3(5):1430-1439.

[5] Iglesias, J., Barg, S., Vallois, D., Lahiri, S., Roger, C., Yessoufou, A., et al., 2012. PPARbeta/delta affects pancreatic beta cell mass and insulin secretion in mice. J Clin Invest 122(11):4105-4117.

**Table S1. Differentially expressed genes (DEGs) in PWS *vs.* control INS-1 lines from RNA-seq with HTSeq feature counts.**

| gene ID | gene name | Control 5-9 | Control 2 | Control 16 | PWS 3 | PWS 19-1 | PWS 19-4 | baseMean | log2FC | lfcSE | stat | pvalue | padj |
|---|---|---|---|---|---|---|---|---|---|---|---|---|---|
| ENSRNOG00000054391 | Snurf | 3992.45 | 3159.50 | 3911.16 | 2.04 | 0.00 | 3.64 | 1844.80 | -10.810 | 0.603 | -17.931 | 6.74E-72 | 6.61E-68 |
| ENSRNOG00000059764 | Snrpn | 3956.37 | 3149.40 | 3874.19 | 2.04 | 0.00 | 3.64 | 1830.94 | -10.799 | 0.603 | -17.921 | 8.04E-72 | 6.61E-68 |
| Rnor_G_Custom_004_psSnurfSnrpn | psSnurfSnrpn | 502.26 | 362.58 | 470.41 | 27.56 | 43.05 | 45.46 | 241.89 | -3.523 | 0.233 | -15.112 | 1.35E-51 | 7.37E-48 |
| ENSRNOG00000058825 | SNORD116 | 846.91 | 659.99 | 1025.04 | 0.00 | 0.00 | 0.00 | 421.99 | -12.113 | 1.192 | -10.158 | 3.04E-24 | 1.25E-20 |
| ENSRNOG00000054305 | SNORD116 | 225.02 | 167.06 | 367.70 | 0.00 | 0.00 | 0.00 | 126.63 | -10.376 | 1.217 | -8.523 | 1.55E-17 | 5.11E-14 |
| ENSRNOG00000059940 | SNORD116 | 189.89 | 165.23 | 362.56 | 0.00 | 0.00 | 0.00 | 119.61 | -10.293 | 1.221 | -8.428 | 3.50E-17 | 9.59E-14 |
| ENSRNOG00000022595 | Snurf-pseudogene LOC100362965 | 219.32 | 141.36 | 203.36 | 0.00 | 0.00 | 0.00 | 94.01 | -9.947 | 1.207 | -8.244 | 1.66E-16 | 3.91E-13 |
| ENSRNOG00000051811 | SNORD116 | 356.04 | 172.57 | 538.20 | 0.00 | 0.00 | 0.00 | 177.80 | -10.866 | 1.362 | -7.977 | 1.50E-15 | 3.09E-12 |
| ENSRNOG00000051750 | SNORD116 | 100.64 | 85.37 | 110.93 | 0.00 | 0.00 | 0.00 | 49.49 | -9.021 | 1.215 | -7.423 | 1.14E-13 | 2.09E-10 |
| Rnor_G_Custom_003_Ipw | Ipw_3 | 88.30 | 92.71 | 108.87 | 0.00 | 0.00 | 0.00 | 48.31 | -8.986 | 1.215 | -7.396 | 1.40E-13 | 2.31E-10 |
| Rnor_G_Custom_001_Ipw | Ipw_1 | 81.65 | 89.96 | 96.55 | 0.00 | 0.00 | 0.00 | 44.69 | -8.874 | 1.217 | -7.294 | 3.01E-13 | 4.12E-10 |
| Rnor_G_Custom_002_Ipw | Ipw_2 | 82.60 | 89.96 | 96.55 | 0.00 | 0.00 | 0.00 | 44.85 | -8.879 | 1.216 | -7.300 | 2.88E-13 | 4.12E-10 |
| ENSRNOG00000010172 | Mkrn3 | 77.86 | 89.04 | 81.14 | 0.00 | 0.00 | 0.00 | 41.34 | -8.762 | 1.219 | -7.188 | 6.57E-13 | 8.31E-10 |
| ENSRNOG00000052187 | SNORD116 | 82.60 | 66.09 | 84.22 | 0.00 | 0.00 | 0.00 | 38.82 | -8.670 | 1.224 | -7.085 | 1.39E-12 | 1.63E-09 |
| ENSRNOG00000058701 | Snord107 | 75.01 | 60.58 | 65.73 | 0.00 | 0.00 | 0.00 | 33.55 | -8.461 | 1.229 | -6.885 | 5.79E-12 | 6.34E-09 |
| ENSRNOG00000008465 | Tmem176b | 222.17 | 204.70 | 216.72 | 410.40 | 469.93 | 397.31 | 320.20 | 0.988 | 0.144 | 6.867 | 6.56E-12 | 6.73E-09 |
| ENSRNOG00000055753 | SNORD116 | 62.66 | 51.40 | 91.41 | 0.00 | 0.00 | 0.00 | 34.25 | -8.488 | 1.241 | -6.839 | 7.98E-12 | 7.71E-09 |
| ENSRNOG00000054389 | SNORD116 | 64.56 | 51.40 | 67.79 | 0.00 | 0.00 | 0.00 | 30.63 | -8.328 | 1.235 | -6.743 | 1.55E-11 | 1.34E-08 |
| ENSRNOG00000059608 | SNORD116 | 67.41 | 45.90 | 76.00 | 0.00 | 0.00 | 0.00 | 31.55 | -8.371 | 1.241 | -6.743 | 1.55E-11 | 1.34E-08 |
| ENSRNOG00000055348 | Snord64 AABR07005710.1 | 71.21 | 46.81 | 66.76 | 0.00 | 0.00 | 0.00 | 30.80 | -8.336 | 1.240 | -6.725 | 1.76E-11 | 1.44E-08 |
| ENSRNOG00000018685 | AABR07028237.1 | 71.21 | 61.50 | 67.79 | 1.02 | 1.20 | 0.91 | 33.94 | -6.020 | 0.898 | -6.701 | 2.08E-11 | 1.62E-08 |
| ENSRNOG00000061924 | Snord64 AABR07005711.2 | 69.31 | 45.90 | 64.71 | 0.00 | 0.00 | 0.00 | 29.99 | -8.298 | 1.241 | -6.688 | 2.25E-11 | 1.68E-08 |
| ENSRNOG00000057137 | AABR07005752.1 | 69.31 | 43.14 | 50.33 | 0.00 | 0.00 | 0.00 | 27.13 | -8.154 | 1.248 | -6.532 | 6.49E-11 | 4.64E-08 |
| ENSRNOG00000052707 | Cacna1a | 2183.74 | 2051.56 | 2365.39 | 3163.76 | 4333.44 | 3858.54 | 2992.74 | 0.783 | 0.127 | 6.171 | 6.80E-10 | 4.65E-07 |
| ENSRNOG00000061528 | U2 | 722.53 | 727.92 | 839.13 | 480.84 | 436.45 | 478.23 | 614.18 | -0.712 | 0.121 | -5.905 | 3.52E-09 | 2.31E-06 |
| ENSRNOG00000002579 | Parm1 | 271.54 | 222.14 | 236.23 | 393.05 | 603.86 | 540.05 | 377.81 | 1.073 | 0.183 | 5.867 | 4.43E-09 | 2.80E-06 |
| ENSRNOG00000053726 | SNORD116 | 25.64 | 33.96 | 43.14 | 0.00 | 0.00 | 0.00 | 17.12 | -7.489 | 1.282 | -5.842 | 5.17E-09 | 3.15E-06 |
| ENSRNOG00000058531 | SNORD115 | 31.33 | 27.54 | 38.00 | 0.00 | 0.00 | 0.00 | 16.15 | -7.404 | 1.280 | -5.784 | 7.31E-09 | 4.29E-06 |
| ENSRNOG00000017579 | Mylip | 760.51 | 1082.23 | 731.29 | 473.70 | 425.69 | 445.50 | 653.15 | -0.937 | 0.165 | -5.677 | 1.37E-08 | 7.75E-06 |
| ENSRNOG00000057393 | SNORD116 | 26.58 | 25.70 | 38.00 | 0.00 | 0.00 | 0.00 | 15.05 | -7.302 | 1.291 | -5.656 | 1.55E-08 | 8.47E-06 |
| ENSRNOG00000001859 | Sdf2l1 | 1998.60 | 2422.41 | 2694.06 | 1579.33 | 1086.95 | 1051.01 | 1805.39 | -0.936 | 0.169 | -5.535 | 3.12E-08 | 1.65E-05 |
| ENSRNOG00000008996 | Dpysl5 | 430.10 | 402.05 | 418.03 | 590.08 | 734.20 | 675.52 | 541.66 | 0.677 | 0.128 | 5.284 | 1.27E-07 | 6.49E-05 |
| ENSRNOG00000032745 | Slc17a3 | 29.43 | 14.69 | 16.43 | 51.04 | 102.84 | 165.47 | 63.32 | 2.400 | 0.455 | 5.278 | 1.30E-07 | 6.49E-05 |
| ENSRNOG00000011796 | C1r | 52.22 | 41.31 | 73.95 | 104.13 | 254.70 | 216.38 | 123.78 | 1.782 | 0.341 | 5.219 | 1.80E-07 | 8.71E-05 |
| ENSRNOG00000023708 | Tmem176a | 277.24 | 180.83 | 225.96 | 487.99 | 466.35 | 380.94 | 336.55 | 0.964 | 0.186 | 5.186 | 2.14E-07 | 1.01E-04 |
| ENSRNOG00000051822 | SNORD116 | 18.99 | 18.36 | 29.79 | 0.00 | 0.00 | 0.00 | 11.19 | -6.874 | 1.330 | -5.170 | 2.35E-07 | 1.07E-04 |
| ENSRNOG00000061669 | SNORD116 | 16.14 | 19.28 | 28.76 | 0.00 | 0.00 | 0.00 | 10.70 | -6.809 | 1.338 | -5.089 | 3.60E-07 | 1.60E-04 |
| ENSRNOG00000060158 | SNORD116 | 18.04 | 16.52 | 26.70 | 0.00 | 0.00 | 0.00 | 10.21 | -6.742 | 1.341 | -5.029 | 4.94E-07 | 2.14E-04 |
| ENSRNOG00000003614 | Mgat5 | 697.85 | 538.82 | 649.12 | 920.85 | 1092.93 | 974.63 | 812.37 | 0.664 | 0.133 | 4.999 | 5.78E-07 | 2.43E-04 |
| ENSRNOG00000056644 | SNORD116 | 18.04 | 19.28 | 20.54 | 0.00 | 0.00 | 0.00 | 9.64 | -6.662 | 1.338 | -4.979 | 6.40E-07 | 2.56E-04 |
| ENSRNOG00000058803 | SNORD115 | 18.99 | 19.28 | 19.51 | 0.00 | 0.00 | 0.00 | 9.63 | -6.660 | 1.338 | -4.979 | 6.38E-07 | 2.56E-04 |
| ENSRNOG00000009341 | Hivep3 | 28.48 | 29.37 | 38.00 | 72.48 | 145.88 | 87.28 | 66.92 | 1.670 | 0.346 | 4.833 | 1.35E-06 | 5.26E-04 |
| ENSRNOG00000011921 | Dusp4 | 1556.15 | 2348.97 | 1947.36 | 1255.70 | 1204.13 | 960.09 | 1545.40 | -0.776 | 0.161 | -4.808 | 1.52E-06 | 5.60E-04 |
| ENSRNOG00000012482 | Ndrg4 | 505.11 | 512.20 | 763.13 | 324.65 | 319.27 | 338.21 | 460.43 | -0.857 | 0.178 | -4.807 | 1.53E-06 | 5.60E-04 |
| ENSRNOG00000058848 | SNORD116 | 13.29 | 15.60 | 25.68 | 0.00 | 0.00 | 0.00 | 9.10 | -6.575 | 1.367 | -4.808 | 1.52E-06 | 5.60E-04 |
| ENSRNOG00000009031 | Gucy2c | 18371.90 | 21412.46 | 21462.08 | 15760.61 | 13849.30 | 12185.66 | 17173.67 | -0.551 | 0.116 | -4.767 | 1.87E-06 | 6.69E-04 |
| ENSRNOG00000016687 | Ssc5d | 105.39 | 134.94 | 126.33 | 234.81 | 278.61 | 200.02 | 180.02 | 0.957 | 0.201 | 4.762 | 1.91E-06 | 6.69E-04 |
| Rnor_G_Custom_012_Snrpn_U1_5 | Snrpn_U1_5 | 18.04 | 21.11 | 12.33 | 0.00 | 0.00 | 0.00 | 8.58 | -6.496 | 1.369 | -4.746 | 2.07E-06 | 7.10E-04 |
| ENSRNOG00000000457 | Tap1 | 374.08 | 616.85 | 443.70 | 280.75 | 249.91 | 219.11 | 364.07 | -0.938 | 0.201 | -4.672 | 2.99E-06 | 1.00E-03 |
| ENSRNOG00000005018 | Scn2a | 794.69 | 639.79 | 689.18 | 956.58 | 1129.99 | 1088.28 | 883.09 | 0.580 | 0.125 | 4.630 | 3.65E-06 | 1.20E-03 |
| ENSRNOG00000006995 | Ano6 | 826.97 | 1375.97 | 1231.48 | 738.11 | 554.83 | 425.49 | 858.81 | -1.000 | 0.217 | -4.607 | 4.09E-06 | 1.32E-03 |
| ENSRNOG00000005621 | Gxylt2 | 142.42 | 148.70 | 98.60 | 231.74 | 234.37 | 311.85 | 194.61 | 0.997 | 0.217 | 4.598 | 4.27E-06 | 1.35E-03 |
| ENSRNOG00000051787 | SNORD115 | 12.34 | 16.52 | 17.46 | 0.00 | 0.00 | 0.00 | 7.72 | -6.341 | 1.380 | -4.594 | 4.35E-06 | 1.35E-03 |
| ENSRNOG00000011971 | C1s | 30.38 | 15.60 | 22.60 | 74.53 | 50.22 | 151.83 | 57.53 | 2.019 | 0.448 | 4.503 | 6.69E-06 | 2.04E-03 |
| ENSRNOG00000002265 | Casr | 159.51 | 100.05 | 108.87 | 167.43 | 402.97 | 397.31 | 222.69 | 1.393 | 0.311 | 4.480 | 7.47E-06 | 2.23E-03 |
| ENSRNOG00000020151 | Cdh1 | 3711.41 | 2918.09 | 4026.19 | 4600.16 | 7094.45 | 6036.01 | 4731.05 | 0.735 | 0.164 | 4.476 | 7.59E-06 | 2.23E-03 |
| ENSRNOG00000016388 | Sphkap | 5297.94 | 4962.31 | 4667.10 | 6152.95 | 7155.43 | 6681.52 | 5819.54 | 0.421 | 0.094 | 4.461 | 8.15E-06 | 2.35E-03 |
| ENSRNOG00000001337 | Setd1b | 127.23 | 131.26 | 150.98 | 202.14 | 286.98 | 252.75 | 191.89 | 0.857 | 0.194 | 4.410 | 1.04E-05 | 2.94E-03 |
| ENSRNOG00000012040 | Slc25a48 | 287.68 | 319.44 | 353.32 | 221.53 | 174.58 | 168.20 | 254.13 | -0.767 | 0.174 | -4.403 | 1.07E-05 | 2.97E-03 |
| ENSRNOG00000004659 | Creld2 | 5611.26 | 6254.75 | 6806.53 | 4943.18 | 3342.15 | 3655.79 | 5102.28 | -0.645 | 0.147 | -4.394 | 1.11E-05 | 3.05E-03 |
| ENSRNOG00000054515 | Fgd6 | 339.90 | 287.31 | 269.10 | 403.25 | 548.85 | 546.41 | 399.14 | 0.741 | 0.169 | 4.389 | 1.14E-05 | 3.06E-03 |
| ENSRNOG00000019482 | Gnao1 | 410.16 | 450.70 | 474.52 | 545.16 | 1098.90 | 920.08 | 649.92 | 0.941 | 0.215 | 4.373 | 1.23E-05 | 3.25E-03 |
| ENSRNOG00000018294 | Hspa5 | 59385.40 | 70664.70 | 72342.09 | 54201.47 | 40806.52 | 40088.23 | 56248.07 | -0.583 | 0.134 | -4.359 | 1.30E-05 | 3.40E-03 |
| ENSRNOG00000061429 | Rph3al | 1045.35 | 848.16 | 966.49 | 1314.92 | 1677.65 | 1295.57 | 1191.36 | 0.583 | 0.137 | 4.274 | 1.92E-05 | 4.94E-03 |
| ENSRNOG00000029598 | Robo2 | 645.63 | 1230.94 | 877.14 | 469.61 | 567.99 | 322.76 | 685.68 | -1.020 | 0.239 | -4.265 | 2.00E-05 | 5.05E-03 |
| ENSRNOG00000001803 | Dnajb11 | 6138.21 | 6061.07 | 7082.82 | 5238.22 | 4217.45 | 4255.85 | 5498.93 | -0.492 | 0.116 | -4.251 | 2.13E-05 | 5.31E-03 |
| ENSRNOG00000059927 | AABR07044414.1 | 72.16 | 148.70 | 90.38 | 37.77 | 44.24 | 32.73 | 71.00 | -1.447 | 0.341 | -4.247 | 2.17E-05 | 5.32E-03 |
| ENSRNOG00000016112 | Cd274 | 112.98 | 97.30 | 136.60 | 170.49 | 345.57 | 239.11 | 183.68 | 1.121 | 0.264 | 4.240 | 2.24E-05 | 5.40E-03 |
| ENSRNOG00000031249 | Slco1a2 | 166.15 | 149.62 | 208.50 | 107.19 | 75.33 | 81.83 | 131.44 | -0.984 | 0.232 | -4.237 | 2.27E-05 | 5.40E-03 |
| ENSRNOG00000013729 | KIAA1549 RGD1306271 | 924.77 | 1011.55 | 958.28 | 1232.22 | 1463.61 | 1277.39 | 1144.64 | 0.456 | 0.108 | 4.223 | 2.41E-05 | 5.66E-03 |
| ENSRNOG00000001449 | Pom121 | 852.61 | 930.78 | 1101.04 | 1225.08 | 1525.79 | 1546.51 | 1196.97 | 0.576 | 0.137 | 4.190 | 2.79E-05 | 6.46E-03 |
| ENSRNOG00000012747 | Spock1 | 75.01 | 150.54 | 110.93 | 55.13 | 32.29 | 39.09 | 77.16 | -1.408 | 0.337 | -4.177 | 2.96E-05 | 6.75E-03 |
| ENSRNOG00000002028 | Tmem50b | 939.96 | 1318.14 | 1055.85 | 738.11 | 728.22 | 755.52 | 922.63 | -0.577 | 0.138 | -4.166 | 3.10E-05 | 6.98E-03 |
| ENSRNOG00000049433 | rno-mir-344a-1 | 10.44 | 16.52 | 9.24 | 0.00 | 0.00 | 0.00 | 6.04 | -5.988 | 1.442 | -4.152 | 3.30E-05 | 7.32E-03 |
| ENSRNOG00000000456 | Psmb8 | 118.68 | 267.12 | 147.90 | 67.38 | 88.49 | 57.28 | 124.47 | -1.331 | 0.321 | -4.142 | 3.45E-05 | 7.36E-03 |
| ENSRNOG00000016690 | Idi1 | 4601.05 | 4927.43 | 3729.37 | 3357.73 | 2802.86 | 3117.56 | 3756.00 | -0.515 | 0.124 | -4.144 | 3.42E-05 | 7.36E-03 |
| ENSRNOG00000017072 | Slc16a14 | 140.52 | 128.51 | 167.42 | 237.87 | 380.25 | 230.02 | 214.10 | 0.956 | 0.231 | 4.147 | 3.37E-05 | 7.36E-03 |
| ENSRNOG00000006426 | Syt1 | 337.06 | 661.82 | 311.21 | 219.49 | 223.61 | 141.83 | 315.84 | -1.166 | 0.283 | -4.115 | 3.87E-05 | 7.69E-03 |
| ENSRNOG00000010488 | Zmiz1 | 496.56 | 709.56 | 739.51 | 930.04 | 1393.06 | 1004.64 | 878.89 | 0.774 | 0.188 | 4.114 | 3.89E-05 | 7.69E-03 |
| ENSRNOG00000016807 | Oat | 2010.94 | 2143.36 | 2338.69 | 1768.19 | 1395.45 | 1328.30 | 1830.82 | -0.531 | 0.129 | -4.123 | 3.74E-05 | 7.69E-03 |
| ENSRNOG00000018561 | Pprc1 | 1317.84 | 1366.79 | 1319.81 | 1638.54 | 1756.57 | 1730.16 | 1521.62 | 0.356 | 0.086 | 4.114 | 3.89E-05 | 7.69E-03 |
| ENSRNOG00000020952 | Cgn | 748.17 | 657.24 | 468.35 | 833.05 | 1145.54 | 1322.85 | 862.53 | 0.817 | 0.199 | 4.112 | 3.93E-05 | 7.69E-03 |
| ENSRNOG00000049057 | Znrf2 | 3829.14 | 3861.72 | 3890.62 | 4665.50 | 5394.08 | 4812.26 | 4408.89 | 0.360 | 0.087 | 4.125 | 3.71E-05 | 7.69E-03 |
| ENSRNOG00000059154 | SNORD116 | 11.39 | 12.85 | 10.27 | 0.00 | 0.00 | 0.00 | 5.75 | -5.918 | 1.439 | -4.112 | 3.92E-05 | 7.69E-03 |
| ENSRNOG00000006378 | Mga | 3931.68 | 3625.81 | 3941.97 | 4507.26 | 5481.37 | 5525.96 | 4502.34 | 0.432 | 0.105 | 4.107 | 4.00E-05 | 7.73E-03 |
| ENSRNOG00000008666 | Etl4 | 495.61 | 507.61 | 598.79 | 702.38 | 901.60 | 771.89 | 662.98 | 0.568 | 0.139 | 4.094 | 4.24E-05 | 7.87E-03 |

| | | | | | | | | | | | | |
|---|---|---|---|---|---|---|---|---|---|---|---|---|
| ENSRNOG00000010944 | Hyou1 | 14472.50 | 16149.07 | 18002.84 | 13107.30 | 10820.44 | 9485.42 | 13672.93 | -0.541 | 0.132 | -4.095 | 4.23E-05 | 7.87E-03 |
| ENSRNOG00000017593 | Mtr | 957.05 | 1033.58 | 980.87 | 1228.14 | 1364.36 | 1321.94 | 1147.66 | 0.397 | 0.097 | 4.093 | 4.26E-05 | 7.87E-03 |
| ENSRNOG00000047326 | AABR07057443.1 | 1410.89 | 1818.41 | 1514.96 | 1203.64 | 1053.47 | 861.90 | 1310.54 | -0.606 | 0.148 | -4.096 | 4.20E-05 | 7.87E-03 |
| ENSRNOG00000000368 | Grik2 | 123.43 | 361.66 | 158.17 | 57.17 | 104.03 | 54.55 | 143.17 | -1.583 | 0.388 | -4.084 | 4.42E-05 | 7.98E-03 |
| ENSRNOG00000046449 | Npy | 579.17 | 631.53 | 699.45 | 423.67 | 475.91 | 433.68 | 540.57 | -0.520 | 0.127 | -4.086 | 4.39E-05 | 7.98E-03 |
| ENSRNOG00000009768 | Npy-like LOC100912228 | 579.17 | 631.53 | 699.45 | 423.67 | 477.11 | 433.68 | 540.77 | -0.518 | 0.127 | -4.070 | 4.70E-05 | 8.40E-03 |
| ENSRNOG00000054286 | Rrm2 | 2728.73 | 2471.06 | 2766.98 | 2152.05 | 1904.85 | 2048.37 | 2345.34 | -0.383 | 0.095 | -4.054 | 5.04E-05 | 8.80E-03 |
| ENSRNOG00000058591 | SNORD115 | 13.29 | 6.43 | 15.41 | 0.00 | 0.00 | 0.00 | 5.85 | -5.939 | 1.465 | -4.055 | 5.02E-05 | 8.80E-03 |
| ENSRNOG00000028225 | Tnni3k | 207.93 | 78.94 | 166.39 | 329.75 | 487.87 | 270.02 | 256.82 | 1.262 | 0.312 | 4.045 | 5.23E-05 | 8.85E-03 |
| ENSRNOG00000052740 | SNORD116 | 15.19 | 11.93 | 7.19 | 0.00 | 0.00 | 0.00 | 5.72 | -5.910 | 1.461 | -4.046 | 5.22E-05 | 8.85E-03 |
| ENSRNOG00000053691 | Lama5 | 544.04 | 354.32 | 651.18 | 742.19 | 972.15 | 1094.65 | 726.42 | 0.859 | 0.212 | 4.050 | 5.12E-05 | 8.85E-03 |
| ENSRNOG00000006228 | Pdia4 | 12993.26 | 13606.42 | 16444.75 | 11547.37 | 8317.71 | 9465.41 | 12062.49 | -0.553 | 0.137 | -4.035 | 5.46E-05 | 9.07E-03 |
| ENSRNOG00000060933 | SNORD115 | 7.60 | 11.02 | 15.41 | 0.00 | 0.00 | 0.00 | 5.67 | -5.894 | 1.461 | -4.035 | 5.47E-05 | 9.07E-03 |
| ENSRNOG00000012802 | Tenm3 | 814.63 | 1055.61 | 818.59 | 1251.62 | 1241.20 | 1345.58 | 1087.87 | 0.513 | 0.128 | 4.013 | 6.00E-05 | 9.71E-03 |
| ENSRNOG00000016352 | Cbfa2t2 | 829.82 | 797.68 | 877.14 | 1011.71 | 1234.02 | 1211.93 | 993.72 | 0.465 | 0.116 | 4.013 | 6.00E-05 | 9.71E-03 |
| ENSRNOG00000056562 | Olfml1 | 186.09 | 69.76 | 70.87 | 277.68 | 264.26 | 265.48 | 189.03 | 1.304 | 0.325 | 4.012 | 6.03E-05 | 9.71E-03 |
| ENSRNOG00000028274 | Myrf | 668.41 | 774.73 | 951.09 | 1075.00 | 1388.28 | 1181.93 | 1006.57 | 0.606 | 0.153 | 3.961 | 7.47E-05 | 1.19E-02 |
| ENSRNOG00000018681 | Nes | 60.76 | 155.13 | 63.68 | 39.81 | 23.92 | 16.37 | 59.94 | -1.808 | 0.457 | -3.958 | 7.56E-05 | 1.19E-02 |
| ENSRNOG00000017967 | Asb13 | 257.30 | 247.84 | 247.53 | 333.83 | 418.52 | 374.58 | 313.27 | 0.581 | 0.147 | 3.951 | 7.77E-05 | 1.22E-02 |
| ENSRNOG00000049361 | Gas7 | 29.43 | 12.85 | 13.35 | 55.13 | 44.24 | 94.55 | 41.59 | 1.806 | 0.458 | 3.945 | 7.98E-05 | 1.24E-02 |
| ENSRNOG00000015321 | Moxd1 | 358.89 | 234.07 | 270.12 | 424.69 | 552.44 | 457.31 | 382.92 | 0.732 | 0.186 | 3.943 | 8.06E-05 | 1.24E-02 |
| ENSRNOG00000061106 | U2 | 436.75 | 467.22 | 508.41 | 345.06 | 236.76 | 317.30 | 385.25 | -0.648 | 0.165 | -3.929 | 8.53E-05 | 1.30E-02 |
| ENSRNOG00000003213 | Helz | 857.36 | 876.62 | 970.60 | 1118.90 | 1333.27 | 1211.93 | 1061.45 | 0.438 | 0.112 | 3.920 | 8.84E-05 | 1.32E-02 |
| ENSRNOG00000004290 | Grb10 | 44.62 | 100.05 | 57.52 | 19.40 | 25.11 | 22.73 | 44.91 | -1.593 | 0.406 | -3.921 | 8.80E-05 | 1.32E-02 |
| ENSRNOG00000013057 | Prc1 | 8208.97 | 8097.94 | 7402.24 | 6668.50 | 6063.70 | 5805.99 | 7041.22 | -0.355 | 0.091 | -3.914 | 9.08E-05 | 1.34E-02 |
| ENSRNOG00000010545 | Mrap2 | 194.64 | 122.08 | 160.23 | 235.83 | 283.40 | 333.67 | 221.64 | 0.840 | 0.216 | 3.897 | 9.75E-05 | 1.42E-02 |
| ENSRNOG00000057887 | U2 | 433.90 | 459.88 | 503.27 | 337.92 | 261.87 | 331.85 | 388.12 | -0.581 | 0.149 | -3.898 | 9.70E-05 | 1.42E-02 |
| ENSRNOG00000057989 | Zp2 | 79.75 | 107.40 | 142.77 | 50.02 | 52.61 | 21.82 | 75.73 | -1.417 | 0.364 | -3.892 | 9.95E-05 | 1.43E-02 |
| ENSRNOG00000022893 | Rimbp2 | 653.22 | 663.66 | 841.19 | 902.47 | 1118.04 | 1213.75 | 898.72 | 0.585 | 0.151 | 3.881 | 1.04E-04 | 1.49E-02 |
| ENSRNOG00000013598 | Melk | 1968.21 | 2532.56 | 2037.75 | 1655.90 | 1567.64 | 1540.14 | 1883.70 | -0.457 | 0.118 | -3.866 | 1.10E-04 | 1.56E-02 |
| ENSRNOG00000032297 | Msmo1 | 3226.24 | 3918.63 | 2790.60 | 2436.88 | 2171.50 | 2403.86 | 2824.62 | -0.503 | 0.131 | -3.848 | 1.19E-04 | 1.67E-02 |
| ENSRNOG00000022505 | Slc17a4 | 595.31 | 444.28 | 599.82 | 724.84 | 828.66 | 891.90 | 680.80 | 0.578 | 0.150 | 3.842 | 1.22E-04 | 1.70E-02 |
| ENSRNOG00000030714 | Bsn | 1363.41 | 1602.70 | 2045.96 | 2392.98 | 3132.89 | 2271.12 | 2134.84 | 0.637 | 0.166 | 3.833 | 1.27E-04 | 1.75E-02 |
| ENSRNOG00000026186 | Syde2 | 572.52 | 581.05 | 637.82 | 756.48 | 795.18 | 911.90 | 709.16 | 0.461 | 0.121 | 3.817 | 1.35E-04 | 1.85E-02 |
| ENSRNOG00000007033 | Sorcs2 | 4028.53 | 3930.56 | 4090.90 | 5247.41 | 4860.77 | 4764.98 | 4487.19 | 0.304 | 0.080 | 3.812 | 1.38E-04 | 1.87E-02 |
| ENSRNOG00000030515 | Nfasc | 2724.93 | 2722.57 | 2985.75 | 3475.13 | 4012.97 | 3440.32 | 3226.95 | 0.374 | 0.098 | 3.808 | 1.40E-04 | 1.89E-02 |
| ENSRNOG00000042245 | Dcaf7 | 3663.94 | 3490.87 | 3606.11 | 4153.01 | 4634.77 | 4778.62 | 4054.55 | 0.334 | 0.088 | 3.803 | 1.43E-04 | 1.91E-02 |
| ENSRNOG00000008639 | Pabpc1 | 6801.88 | 7415.01 | 8762.11 | 9170.72 | 11938.47 | 11149.21 | 9206.23 | 0.489 | 0.129 | 3.791 | 1.50E-04 | 1.99E-02 |
| ENSRNOG00000012067 | Fam111a | 7785.51 | 7014.79 | 7256.39 | 6197.87 | 5734.87 | 5845.99 | 6639.24 | -0.311 | 0.082 | -3.788 | 1.52E-04 | 2.00E-02 |
| ENSRNOG00000001104 | Foxk1 | 390.22 | 521.38 | 502.25 | 618.66 | 699.52 | 824.62 | 592.78 | 0.601 | 0.159 | 3.781 | 1.56E-04 | 2.04E-02 |
| ENSRNOG00000027839 | Ptk2b | 605.75 | 433.26 | 708.69 | 833.05 | 877.69 | 1167.38 | 770.97 | 0.721 | 0.191 | 3.779 | 1.57E-04 | 2.04E-02 |
| ENSRNOG00000015160 | Gem | 255.40 | 324.95 | 344.08 | 206.22 | 202.08 | 144.56 | 246.21 | -0.745 | 0.197 | -3.772 | 1.62E-04 | 2.07E-02 |
| ENSRNOG00000021021 | Ffar2 | 203.18 | 369.01 | 201.31 | 105.15 | 153.06 | 127.28 | 193.17 | -1.008 | 0.267 | -3.771 | 1.62E-04 | 2.07E-02 |
| ENSRNOG00000021433 | Arhgef39 | 930.46 | 1050.11 | 844.27 | 755.46 | 626.58 | 598.24 | 800.85 | -0.513 | 0.136 | -3.766 | 1.66E-04 | 2.10E-02 |
| ENSRNOG00000033262 | Reep6 | 258.25 | 240.50 | 234.18 | 356.29 | 357.53 | 337.30 | 297.34 | 0.519 | 0.138 | 3.760 | 1.70E-04 | 2.12E-02 |
| ENSRNOG00000058589 | AABR07046778.1 | 735.83 | 788.50 | 843.24 | 944.33 | 1177.82 | 1138.29 | 938.00 | 0.462 | 0.123 | 3.759 | 1.71E-04 | 2.12E-02 |
| ENSRNOG00000018784 | Jph3 | 141.47 | 197.35 | 187.96 | 103.11 | 66.96 | 107.28 | 134.02 | -0.918 | 0.244 | -3.755 | 1.73E-04 | 2.14E-02 |
| ENSRNOG00000007584 | Ehd4 | 496.56 | 591.14 | 481.71 | 370.59 | 387.43 | 298.21 | 437.61 | -0.575 | 0.153 | -3.745 | 1.80E-04 | 2.18E-02 |
| ENSRNOG00000008992 | Grik3 | 29.43 | 239.58 | 18.49 | 3.06 | 4.78 | 5.46 | 50.13 | -4.433 | 1.183 | -3.747 | 1.79E-04 | 2.18E-02 |
| ENSRNOG00000014751 | Ret | 275.34 | 199.19 | 298.88 | 140.88 | 136.32 | 172.74 | 203.89 | -0.777 | 0.207 | -3.746 | 1.79E-04 | 2.18E-02 |
| ENSRNOG00000028619 | Hoxc8 | 106.34 | 214.79 | 139.68 | 56.15 | 83.70 | 72.73 | 112.23 | -1.120 | 0.299 | -3.741 | 1.84E-04 | 2.19E-02 |
| Rnor_G_Custom_013_Snrpn_U1_6 | Snrpn_U1_6 | 7.60 | 13.77 | 7.19 | 0.00 | 0.00 | 0.00 | 4.76 | -5.646 | 1.509 | -3.741 | 1.84E-04 | 2.19E-02 |
| ENSRNOG00000013147 | Cacna1d | 1620.72 | 1505.40 | 1979.20 | 2204.12 | 2529.03 | 2290.21 | 2021.45 | 0.460 | 0.123 | 3.737 | 1.86E-04 | 2.19E-02 |
| ENSRNOG00000051533 | SNORD116 | 8.55 | 4.59 | 17.46 | 0.00 | 0.00 | 0.00 | 5.10 | -5.738 | 1.535 | -3.738 | 1.86E-04 | 2.19E-02 |
| ENSRNOG00000008445 | Dact1 | 298.13 | 214.79 | 239.31 | 350.17 | 420.91 | 390.04 | 318.89 | 0.625 | 0.168 | 3.734 | 1.89E-04 | 2.20E-02 |
| ENSRNOG00000023720 | Ntm | 57.92 | 67.93 | 53.41 | 28.59 | 25.11 | 20.00 | 42.16 | -1.289 | 0.346 | -3.729 | 1.92E-04 | 2.23E-02 |
| ENSRNOG00000009956 | Wnk1 | 4654.22 | 4724.57 | 4973.17 | 5555.72 | 7337.19 | 6299.67 | 5590.76 | 0.419 | 0.113 | 3.723 | 1.97E-04 | 2.27E-02 |
| ENSRNOG00000049093 | Hsp90b1 AABR07056495.1 | 2367.93 | 2552.75 | 2795.74 | 2173.49 | 1411.00 | 1602.87 | 2150.63 | -0.572 | 0.154 | -3.718 | 2.01E-04 | 2.29E-02 |
| ENSRNOG00000008956 | Cdkn2c | 1521.97 | 2003.83 | 1404.03 | 1201.60 | 1162.28 | 892.81 | 1364.42 | -0.599 | 0.161 | -3.713 | 2.05E-04 | 2.32E-02 |
| ENSRNOG00000029399 | Bcam | 446.24 | 388.28 | 388.24 | 524.74 | 564.40 | 632.79 | 490.78 | 0.495 | 0.133 | 3.710 | 2.07E-04 | 2.33E-02 |
| ENSRNOG00000003189 | Cited1 | 821.28 | 831.64 | 829.89 | 928.00 | 1409.80 | 1393.76 | 1035.73 | 0.588 | 0.159 | 3.707 | 2.10E-04 | 2.35E-02 |
| ENSRNOG00000010352 | Dnajc3 | 3465.50 | 3352.27 | 4271.67 | 3020.83 | 2533.82 | 2443.86 | 3181.32 | -0.471 | 0.127 | -3.703 | 2.13E-04 | 2.36E-02 |
| ENSRNOG00000003148 | Timp2 | 122.48 | 177.16 | 157.14 | 89.84 | 82.51 | 83.64 | 118.80 | -0.835 | 0.226 | -3.698 | 2.18E-04 | 2.40E-02 |
| ENSRNOG00000016781 | Ppib | 4803.28 | 4730.07 | 5405.58 | 4166.28 | 3685.33 | 3878.54 | 4444.85 | -0.349 | 0.094 | -3.694 | 2.21E-04 | 2.42E-02 |
| ENSRNOG00000022953 | Ccdc163 | 457.64 | 504.86 | 483.76 | 336.90 | 327.64 | 375.49 | 414.38 | -0.474 | 0.129 | -3.680 | 2.33E-04 | 2.52E-02 |
| ENSRNOG00000036459 | Pdia4 AABR07060522.1 | 846.91 | 914.25 | 1084.61 | 739.13 | 520.16 | 655.51 | 793.43 | -0.570 | 0.155 | -3.677 | 2.36E-04 | 2.52E-02 |
| ENSRNOG00000050834 | Fam102a | 1254.23 | 1028.08 | 1297.22 | 1494.59 | 1675.26 | 1674.70 | 1404.01 | 0.437 | 0.119 | 3.677 | 2.36E-04 | 2.52E-02 |
| ENSRNOG00000053379 | SNORD116 | 10.44 | 6.43 | 10.27 | 0.00 | 0.00 | 0.00 | 4.52 | -5.568 | 1.512 | -3.682 | 2.32E-04 | 2.52E-02 |
| ENSRNOG00000023549 | Samd5 | 61.71 | 122.08 | 78.06 | 39.81 | 38.26 | 32.73 | 62.11 | -1.245 | 0.340 | -3.666 | 2.46E-04 | 2.61E-02 |
| ENSRNOG00000018755 | Acss2 | 1603.63 | 1987.31 | 1555.01 | 1306.75 | 1247.18 | 1261.02 | 1493.48 | -0.432 | 0.118 | -3.659 | 2.53E-04 | 2.67E-02 |
| ENSRNOG00000020281 | Kif22 | 4704.54 | 4328.94 | 4273.72 | 3799.78 | 3381.61 | 3408.49 | 3982.85 | -0.329 | 0.090 | -3.650 | 2.62E-04 | 2.74E-02 |
| ENSRNOG00000055471 | Ywhah | 5041.59 | 5057.77 | 5133.40 | 4204.05 | 4360.94 | 4114.92 | 4652.11 | -0.265 | 0.073 | -3.641 | 2.72E-04 | 2.82E-02 |
| ENSRNOG00000058561 | Srrm2 | 5672.03 | 6133.58 | 7462.84 | 8103.88 | 10604.00 | 8456.23 | 7738.76 | 0.495 | 0.136 | 3.640 | 2.72E-04 | 2.82E-02 |
| ENSRNOG00000031053 | Mt-nd4l | 1582.74 | 1720.19 | 1919.63 | 2080.59 | 2509.90 | 2373.85 | 2031.15 | 0.415 | 0.114 | 3.637 | 2.76E-04 | 2.84E-02 |
| ENSRNOG00000021269 | Chgb | 23243.54 | 9354.58 | 12274.76 | 41825.14 | 41728.46 | 41745.65 | 28362.02 | 1.481 | 0.408 | 3.633 | 2.80E-04 | 2.86E-02 |
| ENSRNOG00000001726 | Tmem44 | 353.20 | 332.29 | 382.08 | 254.20 | 255.89 | 180.93 | 293.10 | -0.631 | 0.174 | -3.616 | 2.99E-04 | 2.99E-02 |
| ENSRNOG00000029191 | Gbp7 | 815.58 | 727.00 | 479.65 | 1078.07 | 929.11 | 1224.66 | 875.68 | 0.676 | 0.187 | 3.617 | 2.98E-04 | 2.99E-02 |
| ENSRNOG00000050282 | Srcap | 932.36 | 1164.85 | 1308.51 | 1497.66 | 2126.06 | 1541.05 | 1428.42 | 0.600 | 0.166 | 3.618 | 2.97E-04 | 2.99E-02 |
| ENSRNOG00000047746 | AABR07000398.1 | 12069.44 | 16097.67 | 20812.96 | 22092.21 | 26025.73 | 26518.80 | 20602.80 | 0.608 | 0.168 | 3.613 | 3.03E-04 | 3.01E-02 |
| ENSRNOG00000057044 | Garnl3 | 114.88 | 62.42 | 95.52 | 166.41 | 184.15 | 156.38 | 129.96 | 0.894 | 0.248 | 3.609 | 3.08E-04 | 3.05E-02 |
| ENSRNOG00000006729 | Slc24a4 | 22.79 | 38.55 | 44.16 | 9.19 | 8.37 | 11.82 | 22.48 | -1.833 | 0.509 | -3.603 | 3.14E-04 | 3.06E-02 |
| ENSRNOG00000007459 | Pcnx1 | 1641.60 | 1554.05 | 1681.35 | 1898.87 | 2417.83 | 2129.29 | 1887.16 | 0.402 | 0.112 | 3.604 | 3.14E-04 | 3.06E-02 |
| ENSRNOG00000009433 | Mcub | 243.06 | 283.64 | 324.56 | 188.87 | 193.71 | 170.92 | 234.13 | -0.622 | 0.173 | -3.603 | 3.15E-04 | 3.06E-02 |
| ENSRNOG00000005619 | Misp3 | 695.95 | 872.95 | 796.00 | 600.29 | 593.10 | 510.05 | 678.05 | -0.475 | 0.132 | -3.592 | 3.28E-04 | 3.16E-02 |
| ENSRNOG00000005726 | Pclo | 4565.92 | 5021.97 | 5908.85 | 6239.72 | 8817.54 | 7053.37 | 6267.90 | 0.513 | 0.143 | 3.591 | 3.29E-04 | 3.16E-02 |
| ENSRNOG00000008030 | Shc2 | 3463.60 | 3225.59 | 2930.29 | 3899.83 | 4033.30 | 4210.39 | 3627.17 | 0.336 | 0.094 | 3.587 | 3.35E-04 | 3.18E-02 |
| ENSRNOG00000033321 | Hmgb2l1 | 1400.44 | 1452.16 | 1214.02 | 1096.44 | 1003.24 | 999.18 | 1194.25 | -0.392 | 0.109 | -3.586 | 3.36E-04 | 3.18E-02 |

| | | | | | | | | | | | | | |
|---|---|---|---|---|---|---|---|---|---|---|---|---|---|
| ENSRNOG00000050535 | RragB LOC108348096 | 4635.23 | 4943.95 | 4400.06 | 5328.06 | 6220.35 | 6510.60 | 5339.71 | 0.369 | 0.103 | 3.586 | 3.35E-04 | 3.18E-02 |
| ENSRNOG00000018288 | Ncoa6 | 1429.88 | 1324.57 | 1715.24 | 1857.01 | 2350.87 | 2002.00 | 1779.93 | 0.474 | 0.132 | 3.581 | 3.42E-04 | 3.22E-02 |
| ENSRNOG00000015921 | Esco2 | 1838.14 | 1700.92 | 1677.24 | 1384.34 | 1422.96 | 1384.67 | 1568.04 | -0.316 | 0.089 | -3.568 | 3.60E-04 | 3.36E-02 |
| ENSRNOG00000024905 | Drc1 | 69.31 | 68.84 | 77.03 | 103.11 | 167.41 | 130.92 | 102.77 | 0.897 | 0.251 | 3.566 | 3.63E-04 | 3.37E-02 |
| ENSRNOG00000003160 | RragB | 4658.01 | 4964.14 | 4416.49 | 5344.40 | 6227.52 | 6526.96 | 5356.25 | 0.366 | 0.103 | 3.564 | 3.66E-04 | 3.38E-02 |
| ENSRNOG00000051204 | Dop1b | 491.82 | 559.02 | 545.39 | 721.77 | 1089.34 | 679.15 | 681.08 | 0.640 | 0.180 | 3.562 | 3.68E-04 | 3.38E-02 |
| ENSRNOG00000021763 | Ffar4 | 30.38 | 22.03 | 15.41 | 37.77 | 69.35 | 85.46 | 43.40 | 1.504 | 0.422 | 3.560 | 3.71E-04 | 3.39E-02 |
| ENSRNOG00000004778 | Cnga1 | 218.37 | 142.28 | 235.20 | 309.33 | 301.33 | 389.13 | 265.94 | 0.750 | 0.211 | 3.554 | 3.79E-04 | 3.40E-02 |
| ENSRNOG00000013481 | Cdh11 | 32.28 | 14.69 | 24.65 | 60.23 | 70.55 | 51.82 | 42.37 | 1.349 | 0.379 | 3.555 | 3.78E-04 | 3.40E-02 |
| ENSRNOG00000031475 | Col16a1 | 225.02 | 324.95 | 474.52 | 188.87 | 186.54 | 73.64 | 245.59 | -1.193 | 0.335 | -3.557 | 3.75E-04 | 3.40E-02 |
| ENSRNOG00000060753 | Esyt1 | 1623.56 | 2401.29 | 1773.79 | 1447.63 | 1341.64 | 1177.38 | 1627.55 | -0.548 | 0.154 | -3.552 | 3.82E-04 | 3.41E-02 |
| ENSRNOG00000054165 | Dpysl5 LOC103692570 | 151.91 | 120.25 | 141.74 | 188.87 | 253.50 | 224.57 | 180.14 | 0.687 | 0.194 | 3.549 | 3.87E-04 | 3.44E-02 |
| ENSRNOG00000027410 | Ccdc125 | 6.65 | 5.51 | 8.22 | 6.13 | 141.10 | 301.85 | 78.24 | 4.464 | 1.259 | 3.546 | 3.91E-04 | 3.45E-02 |
| ENSRNOG00000006770 | Brd4 | 541.19 | 567.28 | 707.67 | 747.30 | 1152.71 | 884.63 | 766.79 | 0.616 | 0.174 | 3.539 | 4.01E-04 | 3.52E-02 |
| ENSRNOG00000025551 | Rgs22 | 847.86 | 644.38 | 772.37 | 915.74 | 1164.67 | 1137.38 | 913.73 | 0.507 | 0.143 | 3.537 | 4.05E-04 | 3.54E-02 |
| ENSRNOG00000018194 | Srrm1 | 686.45 | 639.79 | 751.83 | 856.53 | 988.89 | 913.72 | 806.20 | 0.409 | 0.116 | 3.535 | 4.08E-04 | 3.55E-02 |
| ENSRNOG00000015473 | Phactr2 | 157.61 | 87.20 | 139.68 | 187.85 | 248.72 | 259.11 | 180.03 | 0.857 | 0.244 | 3.515 | 4.40E-04 | 3.80E-02 |
| ENSRNOG00000016099 | Id4 | 1090.92 | 2651.89 | 1639.24 | 757.51 | 709.09 | 732.79 | 1263.57 | -1.291 | 0.368 | -3.510 | 4.48E-04 | 3.86E-02 |
| ENSRNOG00000016295 | Lyzl1 | 47.47 | 252.43 | 78.06 | 37.77 | 46.63 | 20.91 | 80.55 | -1.851 | 0.528 | -3.505 | 4.57E-04 | 3.89E-02 |
| ENSRNOG00000020310 | Grik5 | 485.17 | 525.05 | 515.60 | 603.35 | 822.68 | 717.34 | 611.53 | 0.489 | 0.140 | 3.505 | 4.57E-04 | 3.89E-02 |
| ENSRNOG00000016885 | Klf6 | 656.07 | 775.65 | 671.72 | 564.56 | 503.42 | 442.77 | 602.36 | -0.479 | 0.137 | -3.502 | 4.62E-04 | 3.91E-02 |
| ENSRNOG00000004281 | Cobl | 450.04 | 496.60 | 429.32 | 594.16 | 662.45 | 596.42 | 538.17 | 0.428 | 0.122 | 3.500 | 4.65E-04 | 3.92E-02 |
| ENSRNOG00000001424 | Cux1 | 2705.94 | 2782.23 | 3065.87 | 3307.71 | 3688.92 | 4109.47 | 3276.69 | 0.377 | 0.108 | 3.494 | 4.76E-04 | 3.99E-02 |
| ENSRNOG00000001352 | Hectd4 | 1784.97 | 1844.11 | 2238.03 | 2315.39 | 2896.13 | 2757.53 | 2306.03 | 0.442 | 0.127 | 3.492 | 4.79E-04 | 3.99E-02 |
| ENSRNOG00000061230 | L1cam | 215.53 | 301.08 | 264.99 | 185.80 | 162.62 | 135.47 | 210.91 | -0.694 | 0.199 | -3.490 | 4.82E-04 | 4.00E-02 |
| ENSRNOG00000008450 | Rrm2 LOC100359539 | 3818.70 | 3344.92 | 3798.18 | 3034.11 | 2813.62 | 2942.09 | 3291.94 | -0.318 | 0.091 | -3.486 | 4.90E-04 | 4.00E-02 |
| ENSRNOG00000010975 | Adnp | 3293.65 | 3138.39 | 4009.76 | 4036.63 | 5464.63 | 5048.65 | 4165.28 | 0.479 | 0.137 | 3.489 | 4.85E-04 | 4.00E-02 |
| Rnor_G_CustomTg_mINS2mcherry | mINS2mcherry | 34168.89 | 38160.02 | 47714.52 | 25332.54 | 16273.11 | 13026.65 | 29112.62 | -1.136 | 0.326 | -3.487 | 4.88E-04 | 4.00E-02 |
| ENSRNOG00000048980 | Gng2 | 466.18 | 379.10 | 656.31 | 356.29 | 275.02 | 174.56 | 384.58 | -0.899 | 0.258 | -3.480 | 5.01E-04 | 4.06E-02 |
| ENSRNOG00000056228 | Atp10a | 244.96 | 221.22 | 184.88 | 162.32 | 101.64 | 75.46 | 165.08 | -0.942 | 0.271 | -3.481 | 5.00E-04 | 4.06E-02 |
| ENSRNOG00000019100 | Kif2c | 2891.08 | 3399.08 | 2803.96 | 2566.54 | 2062.69 | 2188.38 | 2651.95 | -0.416 | 0.120 | -3.473 | 5.15E-04 | 4.15E-02 |
| ENSRNOG00000001288 | Gpr146 | 96.84 | 84.45 | 116.06 | 132.72 | 174.58 | 283.66 | 148.05 | 0.995 | 0.287 | 3.469 | 5.22E-04 | 4.19E-02 |
| ENSRNOG00000001004 | Ncor2 | 806.09 | 1222.68 | 1165.75 | 1327.17 | 1946.70 | 1634.70 | 1350.51 | 0.619 | 0.179 | 3.467 | 5.27E-04 | 4.20E-02 |
| ENSRNOG00000023412 | Tmem241 | 114.88 | 88.12 | 89.36 | 178.66 | 161.43 | 146.38 | 129.80 | 0.733 | 0.212 | 3.456 | 5.48E-04 | 4.35E-02 |
| ENSRNOG00000005007 | Scn3a | 4752.96 | 4384.93 | 4571.58 | 5166.76 | 5892.71 | 6312.40 | 5180.22 | 0.342 | 0.099 | 3.452 | 5.56E-04 | 4.38E-02 |
| ENSRNOG00000016684 | Wnk2 | 430.10 | 593.90 | 524.84 | 677.88 | 949.43 | 704.61 | 646.79 | 0.589 | 0.171 | 3.452 | 5.56E-04 | 4.38E-02 |
| ENSRNOG00000022325 | Smc2 | 12474.85 | 15025.53 | 11834.14 | 10485.63 | 9581.63 | 10331.86 | 11622.27 | -0.372 | 0.108 | -3.447 | 5.66E-04 | 4.43E-02 |
| ENSRNOG00000057005 | SNORD116 | 8.55 | 6.43 | 8.22 | 0.00 | 0.00 | 0.00 | 3.86 | -5.342 | 1.554 | -3.438 | 5.86E-04 | 4.57E-02 |
| ENSRNOG00000029707 | Mt-nd4 | 6986.07 | 7298.43 | 9706.00 | 9488.22 | 13625.69 | 11677.44 | 9796.98 | 0.536 | 0.156 | 3.435 | 5.93E-04 | 4.58E-02 |
| ENSRNOG00000042944 | Cenpw | 1352.97 | 1194.22 | 1210.94 | 1050.50 | 906.39 | 944.63 | 1109.94 | -0.373 | 0.109 | -3.435 | 5.92E-04 | 4.58E-02 |
| ENSRNOG00000009411 | Chn2 | 269.64 | 324.95 | 369.75 | 237.87 | 206.87 | 160.01 | 261.52 | -0.675 | 0.197 | -3.431 | 6.01E-04 | 4.60E-02 |
| ENSRNOG00000020836 | Rorc | 418.71 | 212.04 | 183.85 | 482.88 | 454.39 | 591.87 | 390.62 | 0.909 | 0.265 | 3.432 | 6.00E-04 | 4.60E-02 |
| ENSRNOG00000020372 | Hdac4 | 231.67 | 268.03 | 262.94 | 357.31 | 365.90 | 341.85 | 304.62 | 0.481 | 0.141 | 3.418 | 6.30E-04 | 4.80E-02 |
| ENSRNOG00000021589 | Nexmif | 1590.33 | 1659.61 | 1731.67 | 1291.43 | 1422.96 | 1248.30 | 1490.72 | -0.331 | 0.097 | -3.417 | 6.34E-04 | 4.80E-02 |
| ENSRNOG00000047007 | AABR07054909.1 | 151.91 | 168.90 | 132.49 | 217.45 | 212.85 | 330.03 | 202.27 | 0.748 | 0.219 | 3.412 | 6.44E-04 | 4.86E-02 |
| ENSRNOG00000001684 | Ripply3 | 1971.06 | 2067.17 | 2273.98 | 1788.61 | 1261.53 | 1535.60 | 1816.32 | -0.460 | 0.135 | -3.410 | 6.51E-04 | 4.88E-02 |
| ENSRNOG00000024428 | Kif20a | 5355.86 | 4979.75 | 4895.11 | 4459.28 | 3659.03 | 3890.36 | 4539.90 | -0.343 | 0.101 | -3.408 | 6.54E-04 | 4.88E-02 |
| ENSRNOG00000002414 | Tfcp2l1 | 523.15 | 436.01 | 628.58 | 698.29 | 953.02 | 726.43 | 660.91 | 0.582 | 0.171 | 3.401 | 6.71E-04 | 4.93E-02 |
| ENSRNOG00000003686 | Tbl1x | 2402.11 | 2141.52 | 2541.02 | 2837.07 | 3254.86 | 2942.09 | 2686.45 | 0.351 | 0.103 | 3.400 | 6.74E-04 | 4.93E-02 |
| ENSRNOG00000011203 | Farp1 | 182.29 | 203.78 | 221.85 | 271.56 | 294.16 | 318.21 | 248.64 | 0.542 | 0.159 | 3.398 | 6.79E-04 | 4.93E-02 |
| ENSRNOG00000011885 | Rhpn2 | 1113.71 | 1197.89 | 1345.49 | 942.29 | 912.37 | 972.82 | 1080.76 | -0.370 | 0.109 | -3.404 | 6.64E-04 | 4.93E-02 |
| ENSRNOG00000013663 | Tmem86a | 546.88 | 309.34 | 464.24 | 577.83 | 872.91 | 725.52 | 582.79 | 0.721 | 0.212 | 3.397 | 6.80E-04 | 4.93E-02 |
| ENSRNOG00000018019 | Hspa12a | 377.88 | 340.55 | 236.23 | 487.99 | 413.73 | 820.98 | 446.23 | 0.853 | 0.251 | 3.397 | 6.81E-04 | 4.93E-02 |
| ENSRNOG00000032112 | AY172581.14 | 742.47 | 842.66 | 911.03 | 1016.81 | 1092.93 | 1336.49 | 990.40 | 0.466 | 0.137 | 3.400 | 6.74E-04 | 4.93E-02 |
| ENSRNOG00000061121 | Wdfy3 | 1756.49 | 2047.89 | 2310.95 | 2542.03 | 3418.68 | 2621.15 | 2449.53 | 0.489 | 0.144 | 3.393 | 6.90E-04 | 4.97E-02 |
| ENSRNOG00000001323 | Zfp157 | 1428.93 | 1434.72 | 1335.22 | 1666.10 | 1769.73 | 2304.76 | 1656.57 | 0.452 | 0.133 | 3.389 | 7.01E-04 | 5.03E-02 |
| ENSRNOG00000020532 | Kcnq1 | 195.59 | 385.53 | 269.10 | 169.47 | 166.21 | 147.29 | 222.20 | -0.818 | 0.241 | -3.386 | 7.10E-04 | 5.05E-02 |
| ENSRNOG00000026196 | Lrrd1 | 6.65 | 7.34 | 8.22 | 9.19 | 46.63 | 49.10 | 21.19 | 2.241 | 0.662 | 3.385 | 7.13E-04 | 5.05E-02 |
| ENSRNOG00000057955 | U2 | 640.88 | 885.80 | 627.55 | 548.22 | 453.19 | 467.32 | 603.83 | -0.553 | 0.163 | -3.385 | 7.11E-04 | 5.05E-02 |
| ENSRNOG00000000455 | Tap2 | 169.95 | 356.16 | 154.06 | 106.17 | 129.14 | 93.64 | 168.19 | -1.052 | 0.311 | -3.381 | 7.21E-04 | 5.07E-02 |
| ENSRNOG00000017893 | Baiap3 | 5191.61 | 5308.37 | 6096.81 | 6721.59 | 7750.92 | 6680.61 | 6291.65 | 0.350 | 0.103 | 3.382 | 7.19E-04 | 5.07E-02 |
| ENSRNOG00000007234 | Cyp51 | 4916.26 | 5396.49 | 4652.72 | 3965.16 | 3595.65 | 4254.03 | 4463.39 | -0.341 | 0.101 | -3.377 | 7.32E-04 | 5.12E-02 |
| ENSRNOG00000005180 | Vstm2a | 76.91 | 119.33 | 85.25 | 56.15 | 41.85 | 42.73 | 70.37 | -1.000 | 0.297 | -3.373 | 7.42E-04 | 5.17E-02 |
| ENSRNOG00000010274 | Smc4 | 4977.98 | 5007.29 | 4679.42 | 4156.07 | 4112.22 | 4027.64 | 4493.44 | -0.254 | 0.075 | -3.372 | 7.46E-04 | 5.17E-02 |
| ENSRNOG00000025001 | Pcolce | 22.79 | 2.75 | 12.33 | 96.99 | 49.03 | 24.55 | 34.74 | 2.173 | 0.648 | 3.353 | 8.00E-04 | 5.50E-02 |
| ENSRNOG00000037613 | Kdm6b | 196.54 | 208.37 | 282.45 | 329.75 | 446.02 | 317.30 | 296.74 | 0.668 | 0.199 | 3.353 | 7.99E-04 | 5.50E-02 |
| ENSRNOG00000004304 | Syap1 | 1328.28 | 1364.04 | 1391.71 | 1132.17 | 1126.41 | 1034.64 | 1229.54 | -0.311 | 0.093 | -3.350 | 8.10E-04 | 5.54E-02 |
| ENSRNOG00000009760 | Palm | 621.89 | 634.29 | 643.99 | 774.86 | 838.23 | 797.35 | 718.43 | 0.343 | 0.102 | 3.347 | 8.17E-04 | 5.57E-02 |
| ENSRNOG00000029682 | Clic1 | 2508.45 | 2590.39 | 2685.84 | 2203.10 | 2170.31 | 1991.09 | 2358.20 | -0.291 | 0.087 | -3.343 | 8.28E-04 | 5.60E-02 |
| ENSRNOG00000037483 | Ep400 | 3359.16 | 3678.13 | 3955.33 | 4123.40 | 4995.89 | 5367.77 | 4246.61 | 0.398 | 0.119 | 3.344 | 8.25E-04 | 5.60E-02 |
| ENSRNOG00000056637 | Abhd16a | 3156.93 | 4634.61 | 3089.49 | 2826.86 | 2469.25 | 2006.55 | 3030.61 | -0.576 | 0.172 | -3.342 | 8.33E-04 | 5.61E-02 |
| ENSRNOG00000005509 | Pak7 | 67.41 | 97.30 | 54.44 | 28.59 | 35.87 | 35.46 | 53.18 | -1.137 | 0.340 | -3.339 | 8.42E-04 | 5.63E-02 |
| ENSRNOG00000059408 | Prr12 | 121.53 | 150.54 | 177.69 | 205.20 | 324.05 | 230.93 | 201.66 | 0.755 | 0.226 | 3.339 | 8.42E-04 | 5.63E-02 |
| ENSRNOG00000014080 | Kif23 | 2884.44 | 2878.62 | 2704.33 | 2451.17 | 2234.88 | 1982.91 | 2522.72 | -0.345 | 0.103 | -3.336 | 8.49E-04 | 5.65E-02 |
| ENSRNOG00000007587 | Tcp11l2 | 303.82 | 246.00 | 261.91 | 375.69 | 368.29 | 386.40 | 323.69 | 0.478 | 0.144 | 3.333 | 8.58E-04 | 5.69E-02 |
| ENSRNOG00000010217 | Prrc2b | 1850.48 | 2138.77 | 2223.65 | 2551.22 | 3186.70 | 2532.96 | 2413.96 | 0.412 | 0.124 | 3.332 | 8.63E-04 | 5.69E-02 |
| ENSRNOG00000022066 | Fam117b | 450.99 | 463.55 | 452.95 | 531.89 | 672.02 | 645.51 | 536.15 | 0.435 | 0.131 | 3.325 | 8.86E-04 | 5.80E-02 |
| ENSRNOG00000026963 | Hsp90b1 | 75710.28 | 67011.36 | 93290.63 | 64969.90 | 52119.61 | 56287.90 | 68231.61 | -0.445 | 0.134 | -3.325 | 8.84E-04 | 5.80E-02 |
| ENSRNOG00000022863 | Lingo4 | 443.39 | 216.63 | 185.90 | 507.39 | 469.93 | 603.69 | 404.49 | 0.902 | 0.272 | 3.319 | 9.05E-04 | 5.88E-02 |
| ENSRNOG00000047401 | Gfy | 250.66 | 132.18 | 311.21 | 385.90 | 414.93 | 438.22 | 322.18 | 0.838 | 0.253 | 3.318 | 9.06E-04 | 5.88E-02 |
| ENSRNOG00000025589 | Jph4 | 121.53 | 62.42 | 92.44 | 172.53 | 159.04 | 162.74 | 128.45 | 0.840 | 0.254 | 3.313 | 9.23E-04 | 5.97E-02 |
| ENSRNOG00000000977 | Pnpla6 | 1323.54 | 1334.66 | 1442.04 | 1592.60 | 1726.68 | 1764.71 | 1530.70 | 0.311 | 0.094 | 3.301 | 9.64E-04 | 6.03E-02 |
| ENSRNOG00000006137 | Arid1a | 700.70 | 768.30 | 858.65 | 919.83 | 1245.98 | 1047.37 | 923.47 | 0.465 | 0.141 | 3.304 | 9.52E-04 | 6.03E-02 |
| ENSRNOG00000009722 | Chd3 | 6866.44 | 6717.38 | 6467.59 | 7568.93 | 8232.81 | 8145.29 | 7333.07 | 0.256 | 0.077 | 3.306 | 9.46E-04 | 6.03E-02 |
| ENSRNOG00000011800 | F3 | 1016.86 | 1130.88 | 1100.01 | 908.60 | 753.33 | 551.87 | 910.26 | -0.554 | 0.167 | -3.309 | 9.37E-04 | 6.03E-02 |
| ENSRNOG00000013515 | Ptpru | 54.12 | 166.14 | 62.65 | 31.65 | 40.66 | 32.73 | 64.66 | -1.435 | 0.435 | -3.302 | 9.61E-04 | 6.03E-02 |
| ENSRNOG00000014268 | Abca2 | 2434.40 | 3024.57 | 2702.28 | 3246.45 | 4274.84 | 3461.23 | 3190.63 | 0.428 | 0.130 | 3.302 | 9.61E-04 | 6.03E-02 |

| | | | | | | | | | | | | | |
|---|---|---|---|---|---|---|---|---|---|---|---|---|---|
| ENSRNOG00000019207 | Shank1 | 207.93 | 213.88 | 298.88 | 356.29 | 399.38 | 332.76 | 301.52 | 0.595 | 0.180 | 3.302 | 9.60E-04 | 6.03E-02 |
| ENSRNOG00000028243 | Derl3 LOC103694875 | 129.13 | 149.62 | 143.79 | 98.01 | 51.42 | 82.73 | 109.12 | -0.856 | 0.260 | -3.300 | 9.68E-04 | 6.03E-02 |
| ENSRNOG00000037514 | Qser1 | 1351.07 | 1466.85 | 1518.04 | 1640.58 | 2072.25 | 1925.63 | 1662.40 | 0.379 | 0.115 | 3.304 | 9.52E-04 | 6.03E-02 |
| ENSRNOG00000055498 | Derl3 | 129.13 | 149.62 | 143.79 | 98.01 | 51.42 | 82.73 | 109.12 | -0.856 | 0.260 | -3.300 | 9.68E-04 | 6.03E-02 |
| ENSRNOG00000024923 | Nnat | 7963.06 | 5684.72 | 9111.32 | 8288.66 | 14469.90 | 13436.69 | 9825.72 | 0.669 | 0.203 | 3.297 | 9.76E-04 | 6.05E-02 |
| ENSRNOG00000008431 | Gabbr2 | 950.40 | 1014.31 | 849.40 | 1187.30 | 1375.12 | 1139.20 | 1085.96 | 0.394 | 0.120 | 3.292 | 9.95E-04 | 6.13E-02 |
| ENSRNOG00000025423 | Map3k13 | 444.34 | 418.57 | 419.05 | 553.33 | 528.53 | 600.06 | 493.98 | 0.393 | 0.119 | 3.291 | 9.98E-04 | 6.13E-02 |
| ENSRNOG00000059984 | Dll1 | 65.51 | 196.44 | 92.44 | 57.17 | 51.42 | 27.28 | 81.71 | -1.390 | 0.422 | -3.291 | 9.99E-04 | 6.13E-02 |
| ENSRNOG00000003769 | Tmem163 | 700.70 | 654.48 | 951.09 | 609.48 | 438.84 | 491.86 | 641.07 | -0.581 | 0.177 | -3.286 | 1.02E-03 | 6.20E-02 |
| ENSRNOG00000006526 | Sema3c | 1569.44 | 1710.10 | 1580.69 | 1162.80 | 1430.13 | 1143.74 | 1432.82 | -0.381 | 0.116 | -3.278 | 1.05E-03 | 6.31E-02 |
| ENSRNOG00000009360 | Sh3bp1 | 385.48 | 317.60 | 381.05 | 456.34 | 557.22 | 490.04 | 431.29 | 0.471 | 0.144 | 3.279 | 1.04E-03 | 6.31E-02 |
| ENSRNOG00000009598 | Ncaph2 | 4556.42 | 4371.16 | 4611.64 | 3938.62 | 3767.84 | 3580.33 | 4137.67 | -0.263 | 0.080 | -3.275 | 1.06E-03 | 6.31E-02 |
| ENSRNOG00000011474 | Ppp1r3b | 584.86 | 578.29 | 467.33 | 689.11 | 706.69 | 966.45 | 665.46 | 0.536 | 0.164 | 3.276 | 1.05E-03 | 6.31E-02 |
| ENSRNOG00000013167 | Hmgb2 | 7334.52 | 7516.90 | 6481.97 | 6140.70 | 5561.48 | 5517.78 | 6425.56 | -0.309 | 0.094 | -3.277 | 1.05E-03 | 6.31E-02 |
| ENSRNOG00000027906 | Ankrd11 | 1803.01 | 1733.04 | 2136.35 | 2318.46 | 2723.94 | 2351.12 | 2177.65 | 0.382 | 0.117 | 3.278 | 1.04E-03 | 6.31E-02 |
| ENSRNOG00000001479 | Gtf2i | 6074.60 | 6045.46 | 6629.87 | 7193.24 | 7827.45 | 10063.65 | 7305.71 | 0.420 | 0.128 | 3.273 | 1.06E-03 | 6.31E-02 |
| ENSRNOG00000027317 | Ylpm1 | 589.61 | 632.45 | 721.02 | 776.90 | 1031.94 | 866.44 | 769.73 | 0.461 | 0.141 | 3.273 | 1.06E-03 | 6.31E-02 |
| ENSRNOG00000011861 | Aadat | 153.81 | 80.78 | 131.47 | 196.01 | 198.50 | 243.66 | 167.37 | 0.805 | 0.246 | 3.270 | 1.07E-03 | 6.33E-02 |
| ENSRNOG00000053599 | Pogz | 1010.22 | 987.69 | 1084.61 | 1207.72 | 1531.77 | 1292.85 | 1185.81 | 0.387 | 0.118 | 3.271 | 1.07E-03 | 6.33E-02 |
| ENSRNOG00000027540 | Fam102b | 242.11 | 386.45 | 345.10 | 168.45 | 239.15 | 194.56 | 262.64 | -0.696 | 0.213 | -3.266 | 1.09E-03 | 6.40E-02 |
| ENSRNOG00000059900 | Bst2 | 174.70 | 141.36 | 177.69 | 228.68 | 253.50 | 245.48 | 203.57 | 0.560 | 0.172 | 3.264 | 1.10E-03 | 6.43E-02 |
| ENSRNOG00000013305 | Atp2c1 | 4827.97 | 4311.50 | 4182.31 | 5123.88 | 5611.70 | 5766.89 | 4970.71 | 0.309 | 0.095 | 3.263 | 1.10E-03 | 6.43E-02 |
| ENSRNOG00000013780 | Nf1 | 2718.28 | 2507.77 | 3225.06 | 3357.73 | 4205.49 | 3664.88 | 3279.87 | 0.410 | 0.126 | 3.261 | 1.11E-03 | 6.45E-02 |
| ENSRNOG00000012417 | Iapp | 210391.05 | 194172.57 | 246567.45 | ####### | 150687.35 | 97751.70 | 180132.79 | -0.600 | 0.185 | -3.250 | 1.15E-03 | 6.62E-02 |
| ENSRNOG00000017172 | Mvb12b | 140.52 | 146.87 | 170.50 | 213.37 | 267.85 | 212.75 | 191.97 | 0.598 | 0.184 | 3.249 | 1.16E-03 | 6.62E-02 |
| ENSRNOG00000048411 | Uhrf1 | 3193.01 | 3433.04 | 3088.46 | 2805.43 | 2619.91 | 2494.77 | 2939.10 | -0.295 | 0.091 | -3.251 | 1.15E-03 | 6.62E-02 |
| ENSRNOG00000056678 | Nckap5l | 88.30 | 81.70 | 108.87 | 135.78 | 180.56 | 145.47 | 123.45 | 0.727 | 0.224 | 3.250 | 1.15E-03 | 6.62E-02 |
| ENSRNOG00000058739 | Snn | 425.35 | 471.81 | 488.90 | 353.23 | 310.90 | 359.12 | 401.55 | -0.435 | 0.134 | -3.248 | 1.16E-03 | 6.62E-02 |
| ENSRNOG00000002104 | Scaf4 | 387.38 | 435.10 | 427.27 | 516.57 | 632.56 | 538.23 | 489.52 | 0.431 | 0.133 | 3.244 | 1.18E-03 | 6.70E-02 |
| ENSRNOG00000010170 | Tubb4b | 13490.77 | 14634.50 | 12848.90 | 11947.56 | 10676.95 | 10935.55 | 12422.37 | -0.288 | 0.089 | -3.239 | 1.20E-03 | 6.80E-02 |
| ENSRNOG00000005390 | Nup210 | 3188.26 | 3390.82 | 3289.77 | 3840.62 | 4777.06 | 3923.09 | 3734.94 | 0.345 | 0.107 | 3.238 | 1.21E-03 | 6.81E-02 |
| ENSRNOG00000014204 | Pacsin3 | 236.41 | 355.24 | 247.53 | 211.33 | 138.71 | 156.38 | 224.27 | -0.728 | 0.226 | -3.227 | 1.25E-03 | 7.03E-02 |
| ENSRNOG00000050206 | Shank2 | 1272.27 | 1183.21 | 1587.88 | 1845.78 | 2093.78 | 1613.78 | 1599.45 | 0.458 | 0.142 | 3.226 | 1.25E-03 | 7.03E-02 |
| ENSRNOG00000012502 | Stk17b | 762.41 | 656.32 | 831.94 | 606.41 | 556.03 | 497.32 | 651.74 | -0.440 | 0.136 | -3.221 | 1.28E-03 | 7.12E-02 |
| ENSRNOG00000029740 | Slc35g1 | 2295.78 | 2685.85 | 2103.48 | 2596.14 | 3418.68 | 4637.70 | 2956.27 | 0.588 | 0.183 | 3.221 | 1.28E-03 | 7.12E-02 |
| ENSRNOG00000030880 | Hs6st2 | 313.32 | 473.65 | 406.73 | 266.45 | 294.16 | 230.93 | 330.87 | -0.595 | 0.185 | -3.217 | 1.29E-03 | 7.18E-02 |
| ENSRNOG00000030644 | Mt-nd1 | 6358.48 | 7009.28 | 7117.74 | 7920.12 | 8158.67 | 8767.17 | 7555.24 | 0.279 | 0.087 | 3.216 | 1.30E-03 | 7.20E-02 |
| ENSRNOG00000004099 | R3hdm1 | 1076.68 | 1079.48 | 1242.78 | 1303.69 | 1670.48 | 1497.41 | 1311.75 | 0.395 | 0.123 | 3.214 | 1.31E-03 | 7.23E-02 |
| ENSRNOG00000028834 | Polr2a | 1301.70 | 1466.85 | 1840.55 | 1789.63 | 2996.58 | 2173.84 | 1928.19 | 0.594 | 0.185 | 3.210 | 1.33E-03 | 7.28E-02 |
| ENSRNOG00000036852 | Pcbp2 | 3055.34 | 3127.37 | 3569.14 | 3676.25 | 4690.97 | 4248.57 | 3727.94 | 0.371 | 0.116 | 3.210 | 1.33E-03 | 7.28E-02 |
| ENSRNOG00000039469 | Pcdhb19 | 118.68 | 110.15 | 108.87 | 152.11 | 170.99 | 203.66 | 144.08 | 0.643 | 0.201 | 3.209 | 1.33E-03 | 7.28E-02 |
| ENSRNOG00000053959 | U3 | 5120.40 | 4904.48 | 5452.82 | 6276.48 | 5902.27 | 6467.87 | 5687.39 | 0.269 | 0.084 | 3.205 | 1.35E-03 | 7.34E-02 |
| ENSRNOG00000047045 | Sdhaf1 LOC108348111 | 1390.95 | 1830.34 | 1319.81 | 1171.99 | 1095.32 | 1082.83 | 1315.21 | -0.439 | 0.137 | -3.204 | 1.35E-03 | 7.34E-02 |
| ENSRNOG00000060367 | Rfx7 | 397.82 | 428.67 | 451.92 | 474.72 | 674.41 | 682.79 | 518.39 | 0.519 | 0.162 | 3.201 | 1.37E-03 | 7.40E-02 |
| ENSRNOG00000014205 | Klf2 | 51.27 | 127.59 | 74.98 | 33.69 | 44.24 | 31.82 | 60.60 | -1.217 | 0.381 | -3.191 | 1.42E-03 | 7.55E-02 |
| ENSRNOG00000030920 | Rtn4r | 42.73 | 36.72 | 45.19 | 63.30 | 92.07 | 83.64 | 60.61 | 0.939 | 0.294 | 3.192 | 1.41E-03 | 7.55E-02 |
| ENSRNOG00000031033 | Mt-nd2 | 4762.45 | 5019.22 | 5996.15 | 6159.07 | 7578.73 | 6785.17 | 6050.13 | 0.379 | 0.119 | 3.192 | 1.41E-03 | 7.55E-02 |
| ENSRNOG00000058438 | AABR07030184.3 | 5127.04 | 4910.90 | 5456.93 | 6279.54 | 5902.27 | 6471.50 | 5691.37 | 0.268 | 0.084 | 3.193 | 1.41E-03 | 7.55E-02 |
| ENSRNOG00000015986 | Rassf8 | 2595.80 | 2320.52 | 2906.67 | 3303.62 | 3327.80 | 3196.66 | 2941.84 | 0.329 | 0.103 | 3.187 | 1.44E-03 | 7.64E-02 |
| ENSRNOG00000051952 | Tes | 1357.72 | 1352.11 | 1438.95 | 1186.28 | 1115.64 | 1056.46 | 1251.19 | -0.305 | 0.096 | -3.184 | 1.45E-03 | 7.69E-02 |
| ENSRNOG00000029134 | Adgrl1 | 2281.53 | 2582.13 | 2696.11 | 3194.39 | 3587.28 | 2902.99 | 2874.07 | 0.357 | 0.112 | 3.182 | 1.47E-03 | 7.74E-02 |
| ENSRNOG00000014201 | Manf | 5043.49 | 5337.74 | 6134.81 | 4880.91 | 3883.83 | 3665.79 | 4824.43 | -0.410 | 0.129 | -3.179 | 1.48E-03 | 7.77E-02 |
| ENSRNOG00000015420 | Stxbp1 | 2281.53 | 1956.10 | 2392.09 | 2695.17 | 3134.09 | 2679.34 | 2523.05 | 0.360 | 0.113 | 3.177 | 1.49E-03 | 7.81E-02 |
| ENSRNOG00000052552 | SNORD116 | 3.80 | 7.34 | 9.24 | 0.00 | 0.00 | 0.00 | 3.40 | -5.155 | 1.626 | -3.171 | 1.52E-03 | 7.94E-02 |
| ENSRNOG00000001851 | Far2 | 101.59 | 78.02 | 87.30 | 64.32 | 28.70 | 28.18 | 64.69 | -1.137 | 0.359 | -3.166 | 1.55E-03 | 8.03E-02 |
| ENSRNOG00000005841 | Erp44 | 3751.29 | 5100.92 | 4165.88 | 3598.66 | 3169.96 | 3057.55 | 3807.38 | -0.406 | 0.128 | -3.165 | 1.55E-03 | 8.03E-02 |
| ENSRNOG00000033556 | Spen | 437.70 | 484.66 | 548.47 | 577.83 | 900.41 | 679.15 | 604.70 | 0.552 | 0.174 | 3.166 | 1.54E-03 | 8.03E-02 |
| ENSRNOG00000001094 | Zfp316 | 275.34 | 254.27 | 313.26 | 378.75 | 376.66 | 391.85 | 331.69 | 0.447 | 0.141 | 3.160 | 1.58E-03 | 8.16E-02 |
| ENSRNOG00000012295 | Pla2g6 | 1285.56 | 1141.90 | 1320.84 | 1536.45 | 1518.62 | 1582.87 | 1397.71 | 0.308 | 0.098 | 3.155 | 1.61E-03 | 8.25E-02 |
| ENSRNOG00000052445 | Kdm7a | 1481.15 | 1507.23 | 1429.71 | 1647.73 | 2148.78 | 1893.81 | 1684.74 | 0.364 | 0.115 | 3.155 | 1.60E-03 | 8.25E-02 |
| ENSRNOG00000053047 | Top2a | 31487.64 | 28322.61 | 28879.73 | 25978.77 | 24394.71 | 23877.65 | 27156.85 | -0.256 | 0.081 | -3.151 | 1.63E-03 | 8.34E-02 |
| ENSRNOG00000054757 | Adcy6 | 3149.33 | 2866.68 | 3251.77 | 3599.68 | 3801.32 | 3802.17 | 3411.83 | 0.274 | 0.087 | 3.150 | 1.63E-03 | 8.34E-02 |
| ENSRNOG00000054212 | Pde1a | 112.04 | 150.54 | 152.01 | 87.80 | 69.35 | 90.01 | 110.29 | -0.741 | 0.235 | -3.148 | 1.64E-03 | 8.37E-02 |
| ENSRNOG00000001706 | Kalrn | 2788.54 | 2357.23 | 2648.87 | 3361.81 | 3712.84 | 2945.73 | 2969.17 | 0.362 | 0.115 | 3.146 | 1.65E-03 | 8.37E-02 |
| ENSRNOG00000012616 | Ppt1 | 1193.46 | 1321.81 | 1107.20 | 955.56 | 963.78 | 962.82 | 1084.11 | -0.330 | 0.105 | -3.146 | 1.66E-03 | 8.37E-02 |
| ENSRNOG00000017940 | Rere | 680.76 | 724.24 | 868.92 | 884.10 | 1176.63 | 1050.10 | 897.46 | 0.452 | 0.144 | 3.138 | 1.70E-03 | 8.57E-02 |
| ENSRNOG00000004889 | Vangl2 | 112.04 | 121.17 | 106.82 | 160.28 | 216.43 | 159.11 | 145.97 | 0.651 | 0.207 | 3.137 | 1.71E-03 | 8.58E-02 |
| ENSRNOG00000002496 | Stxbp5l | 376.93 | 430.51 | 471.43 | 492.07 | 639.73 | 656.42 | 511.18 | 0.484 | 0.155 | 3.123 | 1.79E-03 | 8.90E-02 |
| ENSRNOG00000012613 | F13b | 182.29 | 104.64 | 194.12 | 188.87 | 326.44 | 375.49 | 228.64 | 0.891 | 0.285 | 3.124 | 1.78E-03 | 8.90E-02 |
| ENSRNOG00000014573 | Ckmt1 | 1557.10 | 1520.09 | 1636.16 | 1384.34 | 1061.84 | 1206.47 | 1394.33 | -0.367 | 0.117 | -3.123 | 1.79E-03 | 8.90E-02 |
| ENSRNOG00000033195 | A1cf | 3473.10 | 2940.12 | 4186.42 | 4616.50 | 5239.82 | 4369.49 | 4137.57 | 0.424 | 0.136 | 3.123 | 1.79E-03 | 8.90E-02 |
| ENSRNOG00000008945 | Nans | 4347.54 | 5347.84 | 4410.33 | 3946.79 | 3474.88 | 3713.98 | 4206.89 | -0.341 | 0.109 | -3.121 | 1.80E-03 | 8.92E-02 |
| ENSRNOG00000018666 | Gpsm1 | 836.47 | 834.39 | 757.99 | 1072.96 | 1067.81 | 937.36 | 917.83 | 0.340 | 0.109 | 3.120 | 1.81E-03 | 8.92E-02 |
| ENSRNOG00000018730 | Nectin2 | 485.17 | 471.81 | 573.12 | 670.73 | 687.56 | 640.06 | 588.08 | 0.386 | 0.124 | 3.117 | 1.83E-03 | 8.99E-02 |
| ENSRNOG00000053258 | Faim2 | 60.76 | 17.44 | 44.16 | 104.13 | 49.03 | 210.02 | 80.92 | 1.575 | 0.506 | 3.114 | 1.85E-03 | 9.06E-02 |
| ENSRNOG00000042213 | Zscan25 | 138.62 | 118.41 | 145.85 | 180.70 | 199.69 | 221.84 | 167.52 | 0.583 | 0.187 | 3.111 | 1.87E-03 | 9.13E-02 |
| ENSRNOG00000004563 | Sec24a | 1711.86 | 1715.60 | 2044.94 | 2158.18 | 2500.34 | 2283.85 | 2069.13 | 0.343 | 0.110 | 3.108 | 1.88E-03 | 9.13E-02 |
| ENSRNOG00000013791 | Enpp3 | 224.07 | 276.30 | 208.50 | 300.14 | 333.62 | 420.95 | 293.93 | 0.574 | 0.185 | 3.109 | 1.88E-03 | 9.13E-02 |
| ENSRNOG00000018200 | Gad2 | 402.57 | 1342.93 | 430.35 | 251.14 | 179.36 | 127.28 | 455.61 | -1.964 | 0.632 | -3.109 | 1.87E-03 | 9.13E-02 |
| ENSRNOG00000002418 | Tgfb2 | 39.88 | 61.50 | 110.93 | 33.69 | 23.92 | 28.18 | 49.68 | -1.301 | 0.420 | -3.101 | 1.93E-03 | 9.23E-02 |
| ENSRNOG00000009995 | Rapgef6 | 2084.05 | 2479.32 | 2046.99 | 1714.09 | 1873.76 | 1573.78 | 1962.00 | -0.358 | 0.115 | -3.103 | 1.92E-03 | 9.23E-02 |
| ENSRNOG00000027234 | Zfp367 | 5786.91 | 7448.05 | 5657.21 | 4769.63 | 4870.33 | 5019.55 | 5591.95 | -0.366 | 0.118 | -3.102 | 1.92E-03 | 9.23E-02 |
| ENSRNOG00000046031 | Tap2 LOC103689996 | 148.11 | 328.62 | 145.85 | 100.05 | 126.75 | 86.37 | 155.96 | -0.996 | 0.321 | -3.102 | 1.92E-03 | 9.23E-02 |
| ENSRNOG00000014475 | Slc31a1 | 774.75 | 944.55 | 793.94 | 681.96 | 634.95 | 615.51 | 740.94 | -0.380 | 0.123 | -3.100 | 1.94E-03 | 9.25E-02 |
| ENSRNOG00000028415 | Cdc20 | 2849.31 | 4103.13 | 2748.49 | 2602.27 | 2220.53 | 2064.74 | 2764.74 | -0.494 | 0.160 | -3.096 | 1.96E-03 | 9.34E-02 |
| ENSRNOG00000019491 | Stard10 | 2690.75 | 3188.88 | 2842.99 | 2539.99 | 2197.81 | 2201.11 | 2610.25 | -0.330 | 0.107 | -3.094 | 1.98E-03 | 9.36E-02 |
| ENSRNOG00000037541 | Sdhaf1 | 442.44 | 625.11 | 431.38 | 379.77 | 327.64 | 323.67 | 421.67 | -0.541 | 0.175 | -3.094 | 1.98E-03 | 9.36E-02 |

| | | | | | | | | | | | | | |
|---|---|---|---|---|---|---|---|---|---|---|---|---|---|
| ENSRNOG00000050197 | Pdia6 | 11809.29 | 10480.88 | 15429.98 | 10797.01 | 8101.28 | 7799.81 | 10736.37 | -0.499 | 0.161 | -3.090 | 2.00E-03 | 9.45E-02 |
| ENSRNOG00000015553 | Gatad2b | 704.49 | 759.12 | 804.21 | 896.35 | 1133.58 | 940.09 | 872.97 | 0.388 | 0.126 | 3.083 | 2.05E-03 | 9.65E-02 |
| ENSRNOG00000019215 | Cd151-like LOC100911730 | 6843.65 | 7452.64 | 6885.61 | 6042.69 | 6119.90 | 5663.25 | 6501.29 | -0.249 | 0.081 | -3.082 | 2.06E-03 | 9.66E-02 |
| ENSRNOG00000021314 | Fdft1 | 884.89 | 1110.69 | 739.51 | 694.21 | 591.90 | 666.42 | 781.27 | -0.486 | 0.158 | -3.080 | 2.07E-03 | 9.69E-02 |
| ENSRNOG00000013451 | Drosha | 933.31 | 972.08 | 1070.23 | 1146.47 | 1525.79 | 1256.48 | 1150.73 | 0.400 | 0.130 | 3.078 | 2.08E-03 | 9.69E-02 |
| ENSRNOG00000015308 | Pbk | 1913.15 | 1744.06 | 1839.52 | 1591.58 | 1494.70 | 1372.85 | 1659.31 | -0.302 | 0.098 | -3.079 | 2.08E-03 | 9.69E-02 |
| ENSRNOG00000019667 | Ppfibp2 | 94.95 | 49.57 | 61.63 | 123.53 | 113.60 | 135.47 | 96.46 | 0.856 | 0.278 | 3.077 | 2.09E-03 | 9.70E-02 |
| ENSRNOG00000050374 | Pigg | 961.80 | 973.92 | 1089.74 | 1152.59 | 1336.86 | 1339.21 | 1142.35 | 0.340 | 0.111 | 3.076 | 2.10E-03 | 9.71E-02 |
| ENSRNOG00000022598 | Trerf1 | 125.33 | 130.35 | 190.01 | 216.43 | 252.31 | 222.75 | 189.53 | 0.635 | 0.207 | 3.074 | 2.11E-03 | 9.74E-02 |
| ENSRNOG00000049895 | Pigg LOC100910143 | 961.80 | 973.92 | 1090.77 | 1152.59 | 1338.06 | 1341.03 | 1143.03 | 0.341 | 0.111 | 3.074 | 2.11E-03 | 9.74E-02 |
| ENSRNOG00000013397 | Foxo1 | 351.30 | 380.94 | 427.27 | 462.47 | 694.74 | 518.23 | 472.49 | 0.529 | 0.172 | 3.072 | 2.13E-03 | 9.76E-02 |
| ENSRNOG00000014130 | Cks2 | 4132.97 | 5354.26 | 3777.64 | 3606.83 | 3051.58 | 3248.48 | 3861.96 | -0.421 | 0.137 | -3.071 | 2.13E-03 | 9.76E-02 |
| ENSRNOG00000001708 | Dvl3 | 480.42 | 493.84 | 611.12 | 682.98 | 937.48 | 650.97 | 642.80 | 0.518 | 0.169 | 3.070 | 2.14E-03 | 9.76E-02 |
| ENSRNOG00000011879 | Nfat5 | 1412.79 | 1382.40 | 1868.28 | 1834.55 | 2537.40 | 2099.28 | 1855.78 | 0.473 | 0.154 | 3.067 | 2.16E-03 | 9.83E-02 |
| ENSRNOG00000000543 | Frk | 80.70 | 99.14 | 70.87 | 118.42 | 142.30 | 149.10 | 110.09 | 0.707 | 0.231 | 3.063 | 2.19E-03 | 9.93E-02 |

**Table S2. Differentially expressed genes (DEGs) in PWS *vs.* control INS-1 lines from RNA-seq with RSEM feature counts.**

| gene_ID | gene_name | Control 5-9 | Control 2 | Control 16 | PWS 3 | PWS 19-1 | PWS 19-4 | baseMean | log2FC | lfcSE | stat | pvalue | padj |
|---|---|---|---|---|---|---|---|---|---|---|---|---|---|
| Rnor_G_Custom_004_psSnurfSnrpn | psSnurfSnrpn | 405.43 | 293.12 | 393.20 | 22.42 | 31.37 | 29.13 | 195.78 | -3.72 | 0.25 | -14.71 | 5.24E-49 | 7.46E-45 |
| ENSRNOG00000054391 | Snurf | 1599.61 | 1188.54 | 1550.32 | 0.00 | 0.00 | 0.00 | 723.08 | -12.89 | 1.19 | -10.85 | 2.07E-27 | 1.47E-23 |
| ENSRNOG00000059764 | Snrpn | 1537.94 | 1340.17 | 1554.41 | 0.00 | 0.00 | 1.82 | 739.06 | -11.07 | 1.03 | -10.77 | 4.81E-27 | 2.28E-23 |
| ENSRNOG00000051811 | SNORD116 | 398.05 | 299.46 | 692.56 | 0.00 | 0.00 | 0.00 | 231.68 | -11.33 | 1.22 | -9.31 | 1.26E-20 | 2.55E-17 |
| ENSRNOG00000054305 | SNORD116 | 398.05 | 299.46 | 692.56 | 0.00 | 0.00 | 0.00 | 231.68 | -11.33 | 1.22 | -9.31 | 1.26E-20 | 2.55E-17 |
| ENSRNOG00000058825 | SNORD116 | 398.05 | 299.46 | 692.56 | 0.00 | 0.00 | 0.00 | 231.68 | -11.33 | 1.22 | -9.31 | 1.26E-20 | 2.55E-17 |
| ENSRNOG00000059940 | SNORD116 | 398.05 | 299.46 | 692.56 | 0.00 | 0.00 | 0.00 | 231.68 | -11.33 | 1.22 | -9.31 | 1.26E-20 | 2.55E-17 |
| ENSRNOG00000022595 | LOC100362965 | 120.71 | 70.70 | 93.98 | 0.00 | 0.00 | 0.00 | 47.56 | -8.99 | 1.23 | -7.31 | 2.74E-13 | 4.87E-10 |
| ENSRNOG00000008465 | Tmem176b | 193.42 | 172.05 | 170.94 | 351.91 | 403.36 | 357.10 | 274.80 | 1.05 | 0.15 | 6.91 | 4.90E-12 | 7.74E-09 |
| ENSRNOG00000010172 | Mkrn3 | 58.85 | 58.77 | 58.37 | 0.00 | 0.00 | 0.00 | 29.33 | -8.26 | 1.24 | -6.68 | 2.39E-11 | 3.40E-08 |
| ENSRNOG00000052707 | Cacna1a | 1739.12 | 1632.18 | 1859.79 | 2464.99 | 3407.41 | 3074.14 | 2362.94 | 0.77 | 0.13 | 5.91 | 3.51E-09 | 4.54E-06 |
| ENSRNOG00000002579 | Parm1 | 255.13 | 208.92 | 228.37 | 360.74 | 569.18 | 500.01 | 353.72 | 1.05 | 0.19 | 5.41 | 6.18E-08 | 7.32E-05 |
| ENSRNOG00000001859 | Sdf2l1 | 1874.10 | 2238.89 | 2503.49 | 1474.79 | 1004.43 | 966.84 | 1677.09 | -0.94 | 0.18 | -5.37 | 8.06E-08 | 8.26E-05 |
| ENSRNOG00000017579 | Mylip | 667.84 | 952.59 | 627.75 | 411.70 | 360.00 | 395.81 | 569.28 | -0.95 | 0.18 | -5.35 | 8.71E-08 | 8.26E-05 |
| ENSRNOG00000057137 | AABR07005752.1 | 39.86 | 17.43 | 24.58 | 0.00 | 0.00 | 0.00 | 13.64 | -7.16 | 1.33 | -5.36 | 8.23E-08 | 8.26E-05 |
| ENSRNOG00000057393 | SNORD116 | 24.69 | 19.35 | 24.32 | 0.00 | 0.00 | 0.00 | 11.39 | -6.98 | 1.33 | -5.26 | 1.47E-07 | 1.30E-04 |
| ENSRNOG00000016687 | Ssc5d | 87.26 | 105.38 | 92.17 | 200.75 | 244.09 | 174.26 | 150.65 | 1.12 | 0.21 | 5.22 | 1.80E-07 | 1.51E-04 |
| ENSRNOG00000008996 | Dpysl5 | 385.06 | 362.86 | 369.70 | 525.82 | 636.84 | 596.73 | 479.50 | 0.65 | 0.13 | 5.10 | 3.46E-07 | 2.59E-04 |
| ENSRNOG00000016099 | Id4 | 726.13 | 1783.09 | 1093.64 | 513.64 | 463.50 | 485.51 | 844.25 | -1.30 | 0.26 | -5.10 | 3.43E-07 | 2.59E-04 |
| ENSRNOG00000011796 | C1r | 50.50 | 35.60 | 65.23 | 98.39 | 220.02 | 196.04 | 110.96 | 1.77 | 0.35 | 5.06 | 4.15E-07 | 2.95E-04 |
| ENSRNOG00000023708 | Tmem176a | 251.71 | 164.64 | 201.71 | 438.26 | 429.28 | 339.28 | 304.15 | 0.96 | 0.20 | 4.92 | 8.47E-07 | 5.73E-04 |
| ENSRNOG00000003614 | Mgat5 | 599.59 | 454.85 | 557.09 | 799.97 | 940.95 | 837.97 | 698.40 | 0.68 | 0.14 | 4.86 | 1.15E-06 | 7.44E-04 |
| ENSRNOG00000032745 | Slc17a3 | 14.42 | 7.95 | 11.31 | 29.43 | 65.33 | 110.07 | 39.75 | 2.61 | 0.54 | 4.81 | 1.54E-06 | 9.50E-04 |
| ENSRNOG00000009341 | Hivep3 | 22.59 | 23.56 | 31.41 | 61.83 | 129.56 | 77.25 | 57.70 | 1.79 | 0.38 | 4.71 | 2.49E-06 | 1.48E-03 |
| ENSRNOG00000009031 | Gucy2c | 15776.96 | 18563.33 | 18560.46 | 13679.66 | 11854.09 | 10560.73 | 14832.54 | -0.55 | 0.12 | -4.66 | 3.17E-06 | 1.80E-03 |
| ENSRNOG00000012482 | Ndrg4 | 442.00 | 448.56 | 685.78 | 285.82 | 268.68 | 302.84 | 405.61 | -0.88 | 0.19 | -4.52 | 6.31E-06 | 3.45E-03 |
| ENSRNOG00000011921 | Dusp4 | 1363.96 | 2052.99 | 1736.85 | 1113.81 | 1098.33 | 855.70 | 1370.27 | -0.75 | 0.17 | -4.50 | 6.81E-06 | 3.59E-03 |
| ENSRNOG00000001337 | Setd1b | 79.66 | 84.29 | 101.39 | 149.80 | 213.91 | 173.39 | 133.74 | 1.02 | 0.23 | 4.47 | 7.68E-06 | 3.64E-03 |
| Rnor_G_Custom_001_Ipw | Ipw_1 | 12.46 | 11.28 | 22.50 | 0.00 | 0.00 | 0.00 | 7.71 | -6.35 | 1.42 | -4.48 | 7.59E-06 | 3.64E-03 |
| Rnor_G_Custom_002_Ipw | Ipw_2 | 12.46 | 11.28 | 22.50 | 0.00 | 0.00 | 0.00 | 7.71 | -6.35 | 1.42 | -4.48 | 7.59E-06 | 3.64E-03 |
| ENSRNOG00000029598 | Robo2 | 579.31 | 1046.86 | 795.57 | 388.48 | 497.74 | 311.78 | 603.29 | -1.02 | 0.23 | -4.45 | 8.62E-06 | 3.95E-03 |
| ENSRNOG00000011971 | C1s | 24.67 | 11.92 | 18.43 | 63.18 | 45.90 | 136.79 | 50.15 | 2.17 | 0.49 | 4.44 | 8.92E-06 | 3.96E-03 |
| ENSRNOG00000016112 | Cd274 | 90.15 | 75.25 | 99.33 | 138.60 | 271.67 | 202.31 | 146.22 | 1.21 | 0.27 | 4.41 | 1.05E-05 | 4.39E-03 |
| ENSRNOG00000047898 | Hist2h2aa2 | 5678.74 | 8311.76 | 5858.15 | 4468.97 | 4015.76 | 4185.88 | 5419.88 | -0.65 | 0.15 | -4.41 | 1.03E-05 | 4.39E-03 |
| ENSRNOG00000005621 | Gxylt2 | 97.79 | 102.91 | 69.63 | 170.20 | 165.30 | 204.83 | 135.11 | 1.00 | 0.23 | 4.39 | 1.16E-05 | 4.71E-03 |
| ENSRNOG00000020151 | Cdh1 | 3373.70 | 2641.09 | 3708.23 | 4211.68 | 6420.05 | 5558.95 | 4318.95 | 0.74 | 0.17 | 4.37 | 1.26E-05 | 4.97E-03 |
| ENSRNOG00000005018 | Scn2a | 641.89 | 494.95 | 552.36 | 779.70 | 892.61 | 849.72 | 701.87 | 0.58 | 0.13 | 4.35 | 1.36E-05 | 5.15E-03 |
| ENSRNOG00000014030 | Synm | 156.15 | 162.62 | 179.81 | 89.52 | 99.71 | 80.66 | 128.08 | -0.89 | 0.20 | -4.35 | 1.38E-05 | 5.15E-03 |
| ENSRNOG00000002028 | Tmem50b | 866.31 | 1208.16 | 960.55 | 678.71 | 648.51 | 657.24 | 836.58 | -0.61 | 0.14 | -4.32 | 1.56E-05 | 5.43E-03 |
| ENSRNOG00000016388 | Sphkap | 4845.81 | 4527.10 | 4222.05 | 5590.38 | 6548.59 | 6036.50 | 5295.07 | 0.42 | 0.10 | 4.32 | 1.53E-05 | 5.43E-03 |
| ENSRNOG00000018561 | Pprc1 | 1066.00 | 1129.72 | 1086.57 | 1375.69 | 1456.24 | 1438.09 | 1258.72 | 0.38 | 0.09 | 4.32 | 1.56E-05 | 5.43E-03 |
| ENSRNOG00000013729 | RGD1306271 | 841.11 | 905.85 | 869.47 | 1136.20 | 1338.06 | 1147.32 | 1039.67 | 0.47 | 0.11 | 4.31 | 1.66E-05 | 5.56E-03 |
| ENSRNOG00000047854 | LOC108348072 | 17.36 | 33.08 | 2.04 | 0.00 | 0.00 | 0.00 | 8.75 | -6.70 | 1.56 | -4.30 | 1.68E-05 | 5.56E-03 |
| ENSRNOG00000001803 | Dnajb11 | 5618.10 | 5508.12 | 6493.35 | 4761.24 | 3847.94 | 3905.73 | 5022.41 | -0.49 | 0.12 | -4.24 | 2.25E-05 | 6.53E-03 |
| ENSRNOG00000002265 | Casr | 143.54 | 87.74 | 94.83 | 148.13 | 366.39 | 358.50 | 199.85 | 1.42 | 0.33 | 4.24 | 2.20E-05 | 6.53E-03 |
| ENSRNOG00000003213 | Helz | 678.07 | 691.68 | 736.33 | 894.70 | 1030.92 | 961.86 | 832.26 | 0.45 | 0.11 | 4.25 | 2.18E-05 | 6.53E-03 |
| ENSRNOG00000017593 | Mtr | 690.73 | 689.49 | 691.92 | 890.06 | 943.37 | 916.55 | 803.69 | 0.41 | 0.10 | 4.25 | 2.09E-05 | 6.53E-03 |
| ENSRNOG00000018294 | Hspa5 | 55896.21 | 66418.97 | 68306.52 | 51195.12 | 38597.43 | 38032.35 | 53074.43 | -0.58 | 0.14 | -4.26 | 2.03E-05 | 6.53E-03 |
| ENSRNOG00000019482 | Gnao1 | 296.07 | 291.78 | 354.32 | 399.48 | 778.85 | 653.48 | 462.33 | 0.96 | 0.23 | 4.24 | 2.23E-05 | 6.53E-03 |
| ENSRNOG00000006426 | Syt1 | 309.20 | 587.48 | 292.23 | 200.13 | 190.75 | 134.10 | 285.65 | -1.18 | 0.28 | -4.20 | 2.61E-05 | 7.43E-03 |
| ENSRNOG00000004659 | Creld2 | 4272.55 | 4823.79 | 5170.04 | 3803.49 | 2525.33 | 2830.34 | 3904.26 | -0.64 | 0.15 | -4.19 | 2.77E-05 | 7.72E-03 |
| ENSRNOG00000015921 | Esco2 | 1676.23 | 1535.26 | 1552.77 | 1240.81 | 1195.65 | 1248.33 | 1408.17 | -0.37 | 0.09 | -4.17 | 3.05E-05 | 8.34E-03 |
| ENSRNOG00000006995 | Ano6 | 721.32 | 1147.66 | 1053.26 | 656.71 | 496.26 | 351.21 | 737.74 | -0.96 | 0.23 | -4.15 | 3.38E-05 | 9.07E-03 |
| ENSRNOG00000061429 | Rph3al | 839.84 | 672.63 | 790.55 | 1078.20 | 1346.30 | 1038.02 | 960.92 | 0.59 | 0.14 | 4.14 | 3.46E-05 | 9.11E-03 |
| ENSRNOG00000019207 | Shank1 | 137.41 | 146.33 | 207.18 | 243.35 | 419.06 | 323.53 | 246.14 | 1.00 | 0.24 | 4.13 | 3.69E-05 | 9.55E-03 |
| ENSRNOG00000050877 | Cetn3 | 12.35 | 8.28 | 16.38 | 0.00 | 0.00 | 0.00 | 6.17 | -6.01 | 1.46 | -4.12 | 3.82E-05 | 9.69E-03 |
| ENSRNOG00000016690 | Idi1 | 4289.06 | 4586.68 | 3497.17 | 3131.58 | 2641.78 | 2935.37 | 3513.61 | -0.51 | 0.12 | -4.10 | 4.12E-05 | 1.01E-02 |
| ENSRNOG00000049057 | Znrf2 | 3446.39 | 3494.31 | 3509.59 | 4230.99 | 4874.04 | 4293.50 | 3974.80 | 0.36 | 0.09 | 4.10 | 4.13E-05 | 1.01E-02 |
| ENSRNOG00000010944 | Hyou1 | 13281.99 | 14703.75 | 16342.47 | 11978.74 | 9819.55 | 8708.85 | 12472.56 | -0.54 | 0.13 | -4.07 | 4.70E-05 | 1.13E-02 |
| ENSRNOG00000012802 | Tenm3 | 723.55 | 919.67 | 723.02 | 1098.50 | 1096.27 | 1200.18 | 960.20 | 0.52 | 0.13 | 4.06 | 5.01E-05 | 1.19E-02 |
| ENSRNOG00000054515 | Fgd6 | 306.33 | 252.87 | 239.64 | 354.62 | 474.95 | 487.28 | 352.62 | 0.72 | 0.18 | 4.05 | 5.14E-05 | 1.20E-02 |
| ENSRNOG00000000457 | Tap1 | 320.27 | 534.74 | 375.96 | 262.82 | 210.82 | 178.90 | 313.92 | -0.92 | 0.23 | -4.04 | 5.25E-05 | 1.20E-02 |
| ENSRNOG00000003189 | Cited1 | 696.51 | 705.31 | 703.93 | 831.67 | 1211.72 | 1232.72 | 896.98 | 0.64 | 0.16 | 4.04 | 5.40E-05 | 1.21E-02 |
| ENSRNOG00000057989 | Zp2 | 74.01 | 89.91 | 128.00 | 42.80 | 41.06 | 20.05 | 65.97 | -1.50 | 0.37 | -4.04 | 5.44E-05 | 1.21E-02 |
| ENSRNOG00000003148 | Timp2 | 118.65 | 160.67 | 134.14 | 81.53 | 65.18 | 61.94 | 103.69 | -0.99 | 0.24 | -4.03 | 5.53E-05 | 1.21E-02 |
| ENSRNOG00000006378 | Mga | 3440.46 | 3128.38 | 3482.94 | 3927.58 | 4849.01 | 4938.41 | 3961.13 | 0.45 | 0.11 | 3.99 | 6.59E-05 | 1.39E-02 |
| ENSRNOG00000016807 | Oat | 1843.71 | 1959.71 | 2141.26 | 1626.47 | 1307.80 | 1233.23 | 1685.36 | -0.51 | 0.13 | -3.99 | 6.64E-05 | 1.39E-02 |
| ENSRNOG00000017967 | Asb13 | 214.48 | 212.00 | 214.02 | 291.46 | 373.08 | 326.23 | 271.88 | 0.63 | 0.16 | 3.99 | 6.53E-05 | 1.39E-02 |
| ENSRNOG00000003160 | RragB | 2842.70 | 2953.97 | 2645.21 | 3312.91 | 4396.92 | 4033.98 | 3364.28 | 0.48 | 0.12 | 3.98 | 6.94E-05 | 1.43E-02 |
| ENSRNOG00000008639 | Pabpc1 | 5375.32 | 5827.58 | 6998.99 | 7472.82 | 9718.32 | 9066.28 | 7409.89 | 0.53 | 0.13 | 3.98 | 7.02E-05 | 1.43E-02 |
| ENSRNOG00000008666 | Etl4 | 415.36 | 425.95 | 488.50 | 577.78 | 749.37 | 640.69 | 549.61 | 0.56 | 0.14 | 3.95 | 7.72E-05 | 1.55E-02 |
| ENSRNOG00000053691 | Lama5 | 458.94 | 312.30 | 547.90 | 616.50 | 837.70 | 929.26 | 617.10 | 0.85 | 0.22 | 3.94 | 8.24E-05 | 1.63E-02 |
| ENSRNOG00000000368 | Grik2 | 98.78 | 271.10 | 121.83 | 41.77 | 78.41 | 42.49 | 109.06 | -1.60 | 0.41 | -3.91 | 9.14E-05 | 1.72E-02 |
| ENSRNOG00000010488 | Zmiz1 | 393.55 | 573.42 | 605.25 | 752.03 | 1144.61 | 826.89 | 715.96 | 0.79 | 0.20 | 3.92 | 8.94E-05 | 1.72E-02 |
| ENSRNOG00000031249 | Slco1a2 | 140.40 | 123.77 | 180.24 | 89.68 | 59.20 | 70.23 | 110.58 | -1.01 | 0.26 | -3.91 | 9.21E-05 | 1.72E-02 |
| ENSRNOG00000042245 | Dcaf7 | 3420.82 | 3274.53 | 3339.59 | 3884.53 | 4372.42 | 4534.29 | 3804.36 | 0.35 | 0.09 | 3.91 | 9.05E-05 | 1.72E-02 |
| ENSRNOG00000001449 | Pom121 | 733.10 | 796.19 | 967.77 | 1047.56 | 1337.82 | 1335.00 | 1036.24 | 0.58 | 0.15 | 3.89 | 9.85E-05 | 1.82E-02 |

| | | | | | | | | | | | | | |
|---|---|---|---|---|---|---|---|---|---|---|---|---|---|
| ENSRNOG00000006228 | Pdia4 | 11210.46 | 11775.38 | 14331.53 | 10058.33 | 7226.86 | 8238.28 | 10473.47 | -0.55 | 0.14 | -3.88 | 1.05E-04 | 1.91E-02 |
| ENSRNOG00000016352 | Cbfa2t2 | 682.15 | 644.72 | 694.31 | 817.29 | 977.15 | 961.93 | 796.26 | 0.45 | 0.12 | 3.87 | 1.09E-04 | 1.96E-02 |
| ENSRNOG00000020952 | Cgn | 687.60 | 598.33 | 407.59 | 753.06 | 1046.53 | 1231.82 | 787.49 | 0.84 | 0.22 | 3.86 | 1.14E-04 | 2.03E-02 |
| ENSRNOG00000032297 | Msmo1 | 3023.66 | 3657.50 | 2608.18 | 2268.53 | 2008.96 | 2236.07 | 2633.82 | -0.51 | 0.13 | -3.85 | 1.18E-04 | 2.07E-02 |
| ENSRNOG00000016781 | Ppib | 4495.66 | 4434.85 | 5048.85 | 3867.96 | 3444.01 | 3623.05 | 4152.40 | -0.35 | 0.09 | -3.82 | 1.31E-04 | 2.23E-02 |
| ENSRNOG00000026186 | Syde2 | 494.20 | 515.16 | 560.17 | 672.57 | 703.16 | 810.90 | 626.03 | 0.48 | 0.13 | 3.83 | 1.31E-04 | 2.23E-02 |
| ENSRNOG00000028274 | Myrf | 551.01 | 634.94 | 810.06 | 905.91 | 1207.31 | 1008.33 | 852.93 | 0.65 | 0.17 | 3.83 | 1.29E-04 | 2.23E-02 |
| ENSRNOG00000007033 | Sorcs2 | 3665.39 | 3598.93 | 3735.93 | 4784.26 | 4433.51 | 4324.20 | 4090.37 | 0.30 | 0.08 | 3.81 | 1.36E-04 | 2.26E-02 |
| ENSRNOG00000013598 | Melk | 1638.39 | 2092.48 | 1708.14 | 1386.95 | 1302.18 | 1283.96 | 1568.68 | -0.45 | 0.12 | -3.81 | 1.38E-04 | 2.26E-02 |
| ENSRNOG00000018685 | AABR07028237.1 | 14.26 | 8.29 | 8.19 | 0.00 | 0.00 | 0.00 | 5.12 | -5.75 | 1.51 | -3.81 | 1.37E-04 | 2.26E-02 |
| ENSRNOG00000009411 | Chn2 | 213.93 | 250.56 | 277.70 | 166.49 | 153.89 | 122.66 | 197.54 | -0.75 | 0.20 | -3.80 | 1.46E-04 | 2.37E-02 |
| ENSRNOG00000030515 | Nfasc | 2424.20 | 2410.24 | 2619.80 | 3068.71 | 3522.34 | 3018.36 | 2843.94 | 0.37 | 0.10 | 3.79 | 1.53E-04 | 2.45E-02 |
| ENSRNOG00000009956 | Wnk1 | 4095.56 | 4209.84 | 4460.13 | 4984.77 | 6532.19 | 5639.08 | 4986.93 | 0.43 | 0.11 | 3.77 | 1.63E-04 | 2.49E-02 |
| ENSRNOG00000013057 | Prc1 | 7340.81 | 7238.97 | 6573.48 | 5993.10 | 5460.13 | 5262.81 | 6311.55 | -0.34 | 0.09 | -3.77 | 1.60E-04 | 2.49E-02 |
| ENSRNOG00000028225 | Tnni3k | 144.20 | 58.68 | 126.98 | 233.36 | 337.04 | 197.90 | 183.03 | 1.22 | 0.32 | 3.77 | 1.62E-04 | 2.49E-02 |
| ENSRNOG00000030714 | Bsn | 1211.72 | 1441.75 | 1815.38 | 2138.91 | 2839.30 | 2034.99 | 1913.68 | 0.65 | 0.17 | 3.77 | 1.61E-04 | 2.49E-02 |
| ENSRNOG00000004290 | Grb10 | 43.72 | 91.33 | 54.42 | 14.18 | 25.58 | 22.33 | 41.93 | -1.62 | 0.43 | -3.76 | 1.72E-04 | 2.57E-02 |
| ENSRNOG00000012067 | Fam111a | 7202.87 | 6466.06 | 6759.90 | 5761.74 | 5339.75 | 5449.46 | 6163.30 | -0.30 | 0.08 | -3.76 | 1.73E-04 | 2.57E-02 |
| ENSRNOG00000046449 | Npy | 512.14 | 580.85 | 614.20 | 384.33 | 442.34 | 373.04 | 484.48 | -0.51 | 0.14 | -3.76 | 1.73E-04 | 2.57E-02 |
| ENSRNOG00000010545 | Mrap2 | 139.79 | 83.05 | 114.23 | 176.02 | 218.37 | 229.41 | 160.15 | 0.89 | 0.24 | 3.75 | 1.76E-04 | 2.57E-02 |
| ENSRNOG00000015321 | Moxd1 | 327.29 | 212.71 | 242.70 | 390.30 | 490.46 | 407.65 | 345.19 | 0.72 | 0.19 | 3.75 | 1.79E-04 | 2.57E-02 |
| ENSRNOG00000018784 | Jph3 | 132.80 | 181.49 | 173.07 | 89.68 | 62.83 | 100.34 | 123.37 | -0.94 | 0.25 | -3.75 | 1.79E-04 | 2.57E-02 |
| ENSRNOG00000027839 | Ptk2b | 486.59 | 351.04 | 585.77 | 691.93 | 731.00 | 969.70 | 636.01 | 0.75 | 0.20 | 3.72 | 2.00E-04 | 2.84E-02 |
| ENSRNOG00000055471 | Ywhah | 4699.37 | 4704.19 | 4787.66 | 3912.52 | 4076.77 | 3855.96 | 4339.41 | -0.26 | 0.07 | -3.71 | 2.07E-04 | 2.89E-02 |
| ENSRNOG00000056562 | Olfml1 | 165.07 | 61.43 | 65.54 | 244.57 | 230.73 | 226.16 | 165.58 | 1.26 | 0.34 | 3.71 | 2.07E-04 | 2.89E-02 |
| ENSRNOG00000013147 | Cacna1d | 1314.68 | 1172.05 | 1582.22 | 1769.12 | 2076.74 | 1861.10 | 1629.32 | 0.49 | 0.13 | 3.71 | 2.11E-04 | 2.92E-02 |
| ENSRNOG00000008992 | Grik3 | 22.77 | 208.99 | 14.34 | 3.06 | 3.62 | 3.65 | 42.74 | -4.58 | 1.24 | -3.70 | 2.17E-04 | 2.94E-02 |
| ENSRNOG00000058561 | Srrm2 | 4908.33 | 5347.45 | 6523.58 | 7080.12 | 9433.87 | 7595.58 | 6814.82 | 0.52 | 0.14 | 3.70 | 2.15E-04 | 2.94E-02 |
| ENSRNOG00000017072 | Slc16a14 | 131.01 | 123.53 | 136.59 | 204.93 | 311.19 | 189.83 | 182.85 | 0.85 | 0.23 | 3.69 | 2.26E-04 | 3.03E-02 |
| ENSRNOG00000020025 | LOC108348052 | 0.00 | 0.00 | 0.00 | 3.06 | 41.03 | 1.82 | 7.65 | 6.42 | 1.75 | 3.67 | 2.38E-04 | 3.14E-02 |
| ENSRNOG00000020281 | Kif22 | 4222.26 | 3862.66 | 3827.89 | 3385.38 | 3038.61 | 3088.62 | 3570.90 | -0.32 | 0.09 | -3.67 | 2.39E-04 | 3.14E-02 |
| ENSRNOG00000019100 | Kif2c | 2483.91 | 2889.38 | 2386.20 | 2135.93 | 1711.63 | 1862.28 | 2244.89 | -0.44 | 0.12 | -3.67 | 2.43E-04 | 3.14E-02 |
| ENSRNOG00000029399 | Bcam | 402.29 | 358.63 | 350.23 | 484.06 | 506.05 | 569.80 | 445.18 | 0.49 | 0.13 | 3.67 | 2.43E-04 | 3.14E-02 |
| ENSRNOG00000021269 | Chgb | 21332.11 | 8601.89 | 11257.43 | 38555.61 | 38557.09 | 38679.00 | 26163.85 | 1.49 | 0.41 | 3.66 | 2.52E-04 | 3.23E-02 |
| ENSRNOG00000007459 | Pcnx1 | 1407.26 | 1342.83 | 1433.75 | 1643.67 | 2077.74 | 1825.41 | 1621.78 | 0.41 | 0.11 | 3.65 | 2.64E-04 | 3.35E-02 |
| ENSRNOG00000001104 | Foxk1 | 348.06 | 470.93 | 446.51 | 551.29 | 634.47 | 734.58 | 530.97 | 0.60 | 0.17 | 3.64 | 2.68E-04 | 3.37E-02 |
| ENSRNOG00000012747 | Spock1 | 58.86 | 112.05 | 89.09 | 46.88 | 24.14 | 29.14 | 60.02 | -1.37 | 0.38 | -3.64 | 2.76E-04 | 3.44E-02 |
| ENSRNOG00000001424 | Cux1 | 2202.32 | 2131.51 | 2349.26 | 2634.21 | 2834.87 | 3236.18 | 2564.72 | 0.38 | 0.11 | 3.63 | 2.84E-04 | 3.51E-02 |
| ENSRNOG00000018681 | Nes | 52.49 | 137.69 | 56.38 | 35.19 | 23.28 | 14.63 | 53.28 | -1.76 | 0.49 | -3.62 | 2.93E-04 | 3.60E-02 |
| ENSRNOG00000007584 | Ehd4 | 454.37 | 528.93 | 432.16 | 341.38 | 346.74 | 275.46 | 396.51 | -0.56 | 0.15 | -3.61 | 3.05E-04 | 3.71E-02 |
| ENSRNOG00000054286 | Rrm2 | 1394.13 | 1262.30 | 1461.17 | 1110.91 | 980.60 | 1070.06 | 1213.19 | -0.38 | 0.11 | -3.61 | 3.10E-04 | 3.73E-02 |
| ENSRNOG00000001352 | Hectd4 | 1460.25 | 1521.13 | 1837.27 | 1938.15 | 2393.49 | 2294.94 | 1907.54 | 0.46 | 0.13 | 3.60 | 3.19E-04 | 3.75E-02 |
| ENSRNOG00000014751 | Ret | 226.66 | 165.83 | 243.74 | 114.13 | 114.81 | 139.62 | 167.46 | -0.78 | 0.22 | -3.60 | 3.18E-04 | 3.75E-02 |
| ENSRNOG00000018755 | Acss2 | 1424.92 | 1766.02 | 1384.53 | 1176.00 | 1095.63 | 1109.80 | 1326.15 | -0.44 | 0.12 | -3.60 | 3.18E-04 | 3.75E-02 |
| ENSRNOG00000011752 | Sh3d19 | 248.48 | 187.49 | 222.50 | 315.91 | 372.27 | 313.97 | 276.77 | 0.61 | 0.17 | 3.60 | 3.24E-04 | 3.78E-02 |
| ENSRNOG00000033262 | Reep6 | 181.24 | 176.16 | 172.04 | 264.96 | 263.25 | 248.83 | 217.75 | 0.55 | 0.15 | 3.59 | 3.29E-04 | 3.80E-02 |
| ENSRNOG00000029191 | Gbp7 | 568.47 | 519.46 | 346.12 | 757.19 | 673.70 | 872.95 | 622.98 | 0.68 | 0.19 | 3.59 | 3.31E-04 | 3.80E-02 |
| ENSRNOG00000050282 | Srcap | 732.05 | 960.75 | 1072.25 | 1238.10 | 1750.30 | 1293.43 | 1174.48 | 0.63 | 0.18 | 3.58 | 3.47E-04 | 3.95E-02 |
| ENSRNOG00000002414 | Tfcp2l1 | 425.31 | 374.56 | 540.77 | 607.01 | 830.17 | 648.36 | 571.03 | 0.64 | 0.18 | 3.57 | 3.51E-04 | 3.96E-02 |
| ENSRNOG00000000977 | Pnpla6 | 992.53 | 995.68 | 1057.52 | 1209.00 | 1325.75 | 1311.72 | 1148.70 | 0.34 | 0.09 | 3.57 | 3.58E-04 | 4.01E-02 |
| ENSRNOG00000005726 | Pclo | 3715.58 | 3922.60 | 4981.29 | 5118.48 | 7550.04 | 6006.53 | 5215.75 | 0.57 | 0.16 | 3.56 | 3.77E-04 | 4.18E-02 |
| ENSRNOG00000027410 | Ccdc125 | 5.69 | 5.50 | 5.12 | 5.10 | 117.14 | 235.19 | 62.29 | 4.45 | 1.25 | 3.55 | 3.79E-04 | 4.18E-02 |
| ENSRNOG00000000852 | Prrc2a | 1613.36 | 2125.56 | 2044.90 | 2537.96 | 3117.99 | 2475.51 | 2319.21 | 0.49 | 0.14 | 3.54 | 3.96E-04 | 4.24E-02 |
| ENSRNOG00000001479 | Gtf2i | 5211.30 | 5288.56 | 5773.17 | 6159.84 | 7517.90 | 8873.15 | 6470.65 | 0.47 | 0.13 | 3.54 | 3.97E-04 | 4.24E-02 |
| ENSRNOG00000022953 | Ccdc163 | 324.63 | 359.03 | 352.26 | 234.40 | 224.50 | 263.23 | 293.01 | -0.52 | 0.15 | -3.54 | 3.94E-04 | 4.24E-02 |
| ENSRNOG00000050834 | Fam102a | 1102.17 | 893.61 | 1142.88 | 1308.45 | 1497.07 | 1458.70 | 1233.81 | 0.44 | 0.12 | 3.55 | 3.91E-04 | 4.24E-02 |
| ENSRNOG00000010975 | Adnp | 2967.49 | 2836.57 | 3630.43 | 3686.84 | 5022.62 | 4629.21 | 3795.53 | 0.50 | 0.14 | 3.54 | 4.03E-04 | 4.27E-02 |
| ENSRNOG00000021433 | Arhgef39 | 681.94 | 730.34 | 636.98 | 564.55 | 450.73 | 419.70 | 580.71 | -0.51 | 0.15 | -3.53 | 4.09E-04 | 4.31E-02 |
| ENSRNOG00000027540 | Fam102b | 204.07 | 314.27 | 280.39 | 117.14 | 189.24 | 140.03 | 207.52 | -0.84 | 0.24 | -3.53 | 4.16E-04 | 4.35E-02 |
| Rnor_G_CustomTg_mINS2mcherry | mINS2mcherry | 22001.25 | 24899.31 | 30855.73 | 16370.01 | 10433.83 | 8477.59 | 18839.62 | -1.14 | 0.33 | -3.51 | 4.55E-04 | 4.72E-02 |
| ENSRNOG00000006770 | Brd4 | 426.26 | 436.78 | 569.43 | 603.65 | 965.65 | 716.78 | 619.76 | 0.67 | 0.19 | 3.49 | 4.80E-04 | 4.91E-02 |
| ENSRNOG00000026196 | Lrrd1 | 5.69 | 2.75 | 7.17 | 9.17 | 39.86 | 43.77 | 18.07 | 2.58 | 0.74 | 3.49 | 4.79E-04 | 4.91E-02 |
| ENSRNOG00000054757 | Adcy6 | 2670.28 | 2474.64 | 2792.51 | 3125.05 | 3325.00 | 3311.98 | 2949.91 | 0.30 | 0.09 | 3.49 | 4.92E-04 | 4.96E-02 |
| ENSRNOG00000060849 | Rcor2 | 16.14 | 22.96 | 35.84 | 61.15 | 77.25 | 53.74 | 44.51 | 1.36 | 0.39 | 3.49 | 4.91E-04 | 4.96E-02 |
| ENSRNOG00000061121 | Wdfy3 | 1422.93 | 1685.38 | 1831.82 | 2119.46 | 2770.54 | 2098.40 | 1988.09 | 0.50 | 0.14 | 3.48 | 5.04E-04 | 5.05E-02 |
| ENSRNOG00000021021 | Ffar2 | 133.95 | 257.66 | 140.27 | 71.35 | 102.47 | 83.65 | 131.56 | -1.05 | 0.30 | -3.47 | 5.12E-04 | 5.09E-02 |
| ENSRNOG00000060753 | Esyt1 | 1382.12 | 2010.43 | 1485.93 | 1225.92 | 1123.54 | 1003.28 | 1371.87 | -0.54 | 0.16 | -3.47 | 5.23E-04 | 5.16E-02 |
| ENSRNOG00000024428 | Kif20a | 4681.70 | 4331.48 | 4320.53 | 3881.61 | 3168.56 | 3407.91 | 3965.30 | -0.35 | 0.10 | -3.46 | 5.34E-04 | 5.24E-02 |
| ENSRNOG00000009722 | Chd3 | 6136.58 | 6055.12 | 5823.07 | 6793.80 | 7481.44 | 7417.05 | 6617.85 | 0.27 | 0.08 | 3.45 | 5.55E-04 | 5.29E-02 |
| ENSRNOG00000020310 | Grik5 | 389.85 | 438.12 | 407.58 | 492.20 | 675.41 | 602.06 | 500.87 | 0.52 | 0.15 | 3.45 | 5.58E-04 | 5.29E-02 |
| ENSRNOG00000022325 | Smc2 | 11229.00 | 13490.64 | 10710.02 | 9445.37 | 8590.01 | 9346.68 | 10468.62 | -0.37 | 0.11 | -3.45 | 5.52E-04 | 5.29E-02 |
| ENSRNOG00000022893 | Rimbp2 | 504.63 | 486.67 | 638.91 | 667.35 | 855.48 | 846.48 | 666.59 | 0.54 | 0.16 | 3.46 | 5.46E-04 | 5.29E-02 |
| ENSRNOG00000029682 | Clic1 | 2341.92 | 2395.90 | 2510.71 | 2032.25 | 2022.70 | 1866.23 | 2194.95 | -0.29 | 0.08 | -3.45 | 5.57E-04 | 5.29E-02 |
| ENSRNOG00000049361 | Gas7 | 24.50 | 10.39 | 9.67 | 47.97 | 31.93 | 68.86 | 32.22 | 1.76 | 0.51 | 3.44 | 5.72E-04 | 5.38E-02 |
| ENSRNOG00000004304 | Syap1 | 1227.82 | 1256.77 | 1279.03 | 1034.37 | 1037.38 | 948.95 | 1130.72 | -0.32 | 0.09 | -3.43 | 5.96E-04 | 5.47E-02 |
| ENSRNOG00000008030 | Shc2 | 2862.63 | 2701.02 | 2440.39 | 3183.49 | 3359.00 | 3456.51 | 3000.51 | 0.32 | 0.09 | 3.43 | 5.96E-04 | 5.47E-02 |
| ENSRNOG00000031475 | Col16a1 | 183.04 | 250.14 | 364.58 | 150.82 | 143.81 | 63.87 | 192.71 | -1.16 | 0.34 | -3.43 | 5.94E-04 | 5.47E-02 |
| ENSRNOG00000052445 | Kdm7a | 1081.57 | 1088.59 | 1072.19 | 1254.48 | 1556.92 | 1395.03 | 1241.46 | 0.37 | 0.11 | 3.44 | 5.86E-04 | 5.47E-02 |

| | | | | | | | | | | | | | |
|---|---|---|---|---|---|---|---|---|---|---|---|---|---|
| ENSRNOG00000023549 | Samd5 | 55.98 | 101.81 | 66.56 | 30.57 | 36.23 | 31.00 | 53.69 | -1.20 | 0.35 | -3.42 | 6.17E-04 | 5.63E-02 |
| ENSRNOG00000010217 | Prrc2b | 1527.94 | 1783.85 | 1867.76 | 2185.41 | 2754.76 | 2138.96 | 2043.11 | 0.45 | 0.13 | 3.42 | 6.27E-04 | 5.68E-02 |
| ENSRNOG00000001004 | Ncor2 | 632.77 | 964.32 | 925.66 | 1044.13 | 1602.90 | 1335.88 | 1084.28 | 0.66 | 0.19 | 3.41 | 6.48E-04 | 5.76E-02 |
| ENSRNOG00000013780 | Nf1 | 2349.54 | 2168.32 | 2778.06 | 2929.81 | 3680.09 | 3250.28 | 2859.35 | 0.43 | 0.13 | 3.41 | 6.46E-04 | 5.76E-02 |
| ENSRNOG00000061230 | L1cam | 191.32 | 265.50 | 231.82 | 154.39 | 151.16 | 119.03 | 185.54 | -0.70 | 0.21 | -3.41 | 6.44E-04 | 5.76E-02 |
| ENSRNOG00000007234 | Cyp51 | 4589.30 | 4984.21 | 4330.81 | 3682.85 | 3340.49 | 3955.75 | 4147.24 | -0.34 | 0.10 | -3.40 | 6.62E-04 | 5.85E-02 |
| ENSRNOG00000014080 | Kif23 | 2415.32 | 2363.76 | 2214.01 | 2007.36 | 1830.22 | 1649.44 | 2080.02 | -0.35 | 0.10 | -3.40 | 6.82E-04 | 5.95E-02 |
| ENSRNOG00000028619 | Hoxc8 | 83.50 | 171.57 | 102.40 | 43.82 | 61.59 | 58.33 | 86.87 | -1.13 | 0.33 | -3.40 | 6.81E-04 | 5.95E-02 |
| ENSRNOG00000056637 | Abhd16a | 2177.41 | 3246.99 | 2198.80 | 1932.60 | 1706.25 | 1412.91 | 2112.49 | -0.59 | 0.18 | -3.39 | 6.90E-04 | 5.96E-02 |
| ENSRNOG00000059984 | Dll1 | 57.86 | 173.24 | 77.83 | 43.82 | 41.08 | 23.72 | 69.59 | -1.51 | 0.45 | -3.39 | 6.91E-04 | 5.96E-02 |
| ENSRNOG00000001323 | Zfp157 | 1205.69 | 1215.63 | 1129.55 | 1410.37 | 1516.16 | 1966.39 | 1407.30 | 0.46 | 0.14 | 3.36 | 7.66E-04 | 6.07E-02 |
| ENSRNOG00000009598 | Ncaph2 | 3722.54 | 3494.03 | 3754.48 | 3160.33 | 3024.00 | 2886.96 | 3340.39 | -0.27 | 0.08 | -3.38 | 7.28E-04 | 6.07E-02 |
| ENSRNOG00000009760 | Palm | 512.32 | 532.83 | 527.39 | 652.20 | 709.01 | 661.96 | 599.29 | 0.36 | 0.11 | 3.37 | 7.49E-04 | 6.07E-02 |
| ENSRNOG00000010274 | Smc4 | 4200.83 | 4180.82 | 3903.82 | 3464.73 | 3461.89 | 3383.80 | 3765.98 | -0.25 | 0.07 | -3.38 | 7.20E-04 | 6.07E-02 |
| ENSRNOG00000014285 | Ssh2 | 494.09 | 480.96 | 539.70 | 627.72 | 831.51 | 657.98 | 605.33 | 0.48 | 0.14 | 3.37 | 7.56E-04 | 6.07E-02 |
| ENSRNOG00000018019 | Hspa12a | 350.89 | 315.14 | 221.21 | 454.48 | 395.21 | 776.62 | 418.92 | 0.88 | 0.26 | 3.36 | 7.72E-04 | 6.07E-02 |
| ENSRNOG00000018194 | Srrm1 | 491.15 | 468.53 | 555.12 | 619.10 | 744.26 | 684.97 | 593.85 | 0.44 | 0.13 | 3.36 | 7.73E-04 | 6.07E-02 |
| ENSRNOG00000021589 | Nexmif | 1418.67 | 1472.05 | 1534.11 | 1135.19 | 1272.72 | 1107.07 | 1323.30 | -0.33 | 0.10 | -3.37 | 7.50E-04 | 6.07E-02 |
| ENSRNOG00000025589 | Jph4 | 82.56 | 50.48 | 62.47 | 133.50 | 114.70 | 113.90 | 92.94 | 0.89 | 0.26 | 3.38 | 7.15E-04 | 6.07E-02 |
| ENSRNOG00000027906 | Ankrd11 | 1571.28 | 1540.59 | 1883.34 | 2062.48 | 2402.96 | 2096.58 | 1926.21 | 0.39 | 0.12 | 3.37 | 7.60E-04 | 6.07E-02 |
| ENSRNOG00000029707 | Mt-nd4 | 5942.39 | 6199.52 | 8338.25 | 8112.43 | 11911.89 | 10043.20 | 8424.61 | 0.55 | 0.16 | 3.37 | 7.64E-04 | 6.07E-02 |
| ENSRNOG00000051822 | SNORD116 | 7.90 | 4.07 | 11.63 | 0.00 | 0.00 | 0.00 | 3.93 | -5.45 | 1.62 | -3.36 | 7.67E-04 | 6.07E-02 |
| ENSRNOG00000052552 | SNORD116 | 7.90 | 4.07 | 11.63 | 0.00 | 0.00 | 0.00 | 3.93 | -5.45 | 1.62 | -3.36 | 7.67E-04 | 6.07E-02 |
| ENSRNOG00000059408 | Prr12 | 95.78 | 133.75 | 146.45 | 178.33 | 277.98 | 208.08 | 173.39 | 0.82 | 0.24 | 3.38 | 7.30E-04 | 6.07E-02 |
| ENSRNOG00000060158 | SNORD116 | 7.90 | 4.07 | 11.63 | 0.00 | 0.00 | 0.00 | 3.93 | -5.45 | 1.62 | -3.36 | 7.67E-04 | 6.07E-02 |
| ENSRNOG00000061669 | SNORD116 | 7.90 | 4.07 | 11.63 | 0.00 | 0.00 | 0.00 | 3.93 | -5.45 | 1.62 | -3.36 | 7.67E-04 | 6.07E-02 |
| ENSRNOG00000025423 | Map3k13 | 356.89 | 315.65 | 358.55 | 469.37 | 431.54 | 510.14 | 407.02 | 0.46 | 0.14 | 3.36 | 7.84E-04 | 6.09E-02 |
| ENSRNOG00000053599 | Pogz | 847.84 | 830.78 | 936.73 | 1046.39 | 1295.04 | 1112.98 | 1011.63 | 0.40 | 0.12 | 3.36 | 7.84E-04 | 6.09E-02 |
| ENSRNOG00000003686 | Tbl1x | 2186.81 | 1907.43 | 2281.01 | 2568.84 | 2974.68 | 2642.85 | 2426.94 | 0.36 | 0.11 | 3.36 | 7.92E-04 | 6.09E-02 |
| ENSRNOG00000051204 | Dop1b | 403.03 | 472.67 | 453.68 | 598.16 | 922.20 | 566.76 | 569.42 | 0.65 | 0.19 | 3.36 | 7.90E-04 | 6.09E-02 |
| ENSRNOG00000005007 | Scn3a | 3951.73 | 3635.39 | 3805.54 | 4264.83 | 4892.10 | 5263.77 | 4302.23 | 0.34 | 0.10 | 3.35 | 8.21E-04 | 6.23E-02 |
| ENSRNOG00000016885 | Klf6 | 379.32 | 432.08 | 376.33 | 313.49 | 274.58 | 246.78 | 337.10 | -0.51 | 0.15 | -3.34 | 8.23E-04 | 6.23E-02 |
| ENSRNOG00000018288 | Ncoa6 | 1230.93 | 1120.37 | 1481.88 | 1586.61 | 2003.88 | 1695.65 | 1519.89 | 0.46 | 0.14 | 3.34 | 8.24E-04 | 6.23E-02 |
| ENSRNOG00000005390 | Nup210 | 2818.42 | 2968.95 | 2911.53 | 3423.91 | 4272.51 | 3478.98 | 3312.38 | 0.36 | 0.11 | 3.33 | 8.71E-04 | 6.55E-02 |
| ENSRNOG00000013481 | Cdh11 | 28.45 | 12.83 | 22.53 | 51.97 | 62.84 | 47.45 | 37.68 | 1.35 | 0.40 | 3.33 | 8.79E-04 | 6.58E-02 |
| ENSRNOG00000013305 | Atp2c1 | 3406.90 | 3137.39 | 3008.54 | 3660.60 | 4020.91 | 4106.99 | 3556.89 | 0.30 | 0.09 | 3.31 | 9.22E-04 | 6.86E-02 |
| ENSRNOG00000006137 | Arid1a | 596.48 | 671.42 | 742.48 | 798.91 | 1087.82 | 932.78 | 804.98 | 0.49 | 0.15 | 3.31 | 9.27E-04 | 6.86E-02 |
| ENSRNOG00000030644 | Mt-nd1 | 5922.08 | 6570.07 | 6595.92 | 7393.51 | 7640.73 | 8169.64 | 7048.66 | 0.28 | 0.09 | 3.31 | 9.47E-04 | 6.94E-02 |
| ENSRNOG00000058739 | Snn | 390.85 | 438.29 | 442.40 | 317.95 | 282.67 | 326.47 | 366.44 | -0.45 | 0.14 | -3.31 | 9.47E-04 | 6.94E-02 |
| ENSRNOG00000019189 | Acat2 | 519.35 | 533.61 | 396.28 | 356.41 | 312.15 | 341.33 | 409.86 | -0.52 | 0.16 | -3.30 | 9.64E-04 | 7.01E-02 |
| ENSRNOG00000037514 | Qser1 | 1247.92 | 1343.66 | 1396.89 | 1510.18 | 1936.39 | 1781.69 | 1536.12 | 0.39 | 0.12 | 3.30 | 9.67E-04 | 7.01E-02 |
| ENSRNOG00000055111 | AABR07000658.2 | 1486.62 | 1421.78 | 1438.71 | 1228.06 | 1023.39 | 1177.49 | 1296.01 | -0.34 | 0.10 | -3.30 | 9.72E-04 | 7.02E-02 |
| ENSRNOG00000017893 | Baiap3 | 4216.50 | 4374.84 | 5020.14 | 5459.94 | 6383.44 | 5520.78 | 5162.61 | 0.35 | 0.11 | 3.30 | 9.82E-04 | 7.03E-02 |
| ENSRNOG00000048980 | Gng2 | 410.74 | 315.34 | 546.86 | 298.58 | 247.67 | 160.53 | 329.95 | -0.85 | 0.26 | -3.30 | 9.84E-04 | 7.03E-02 |
| ENSRNOG00000001684 | Ripply3 | 1645.25 | 1759.12 | 1929.18 | 1514.45 | 1073.98 | 1298.44 | 1536.74 | -0.46 | 0.14 | -3.29 | 1.01E-03 | 7.05E-02 |
| ENSRNOG00000006526 | Sema3c | 1414.08 | 1537.52 | 1441.93 | 1038.39 | 1288.22 | 1043.85 | 1294.00 | -0.38 | 0.12 | -3.29 | 1.01E-03 | 7.05E-02 |
| ENSRNOG00000022505 | Slc17a4 | 353.93 | 244.39 | 332.27 | 406.94 | 483.62 | 498.04 | 386.53 | 0.58 | 0.18 | 3.29 | 9.94E-04 | 7.05E-02 |
| ENSRNOG00000029134 | Adgrl1 | 1888.02 | 2136.96 | 2185.69 | 2626.80 | 2983.03 | 2414.87 | 2372.56 | 0.37 | 0.11 | 3.29 | 1.01E-03 | 7.05E-02 |
| ENSRNOG00000037483 | Ep400 | 2859.47 | 3167.00 | 3398.17 | 3546.34 | 4273.84 | 4633.38 | 3646.37 | 0.40 | 0.12 | 3.29 | 1.00E-03 | 7.05E-02 |
| ENSRNOG00000060367 | Rfx7 | 344.23 | 359.98 | 399.40 | 419.83 | 606.76 | 610.60 | 456.80 | 0.57 | 0.17 | 3.29 | 1.02E-03 | 7.05E-02 |
| ENSRNOG00000013515 | Ptpru | 48.75 | 150.31 | 53.11 | 27.37 | 32.50 | 26.79 | 56.47 | -1.55 | 0.47 | -3.28 | 1.02E-03 | 7.06E-02 |
| ENSRNOG00000009433 | Mcub | 204.17 | 247.28 | 270.32 | 161.03 | 171.28 | 138.32 | 198.74 | -0.62 | 0.19 | -3.28 | 1.04E-03 | 7.17E-02 |
| ENSRNOG00000012613 | F13b | 146.73 | 85.76 | 160.39 | 166.65 | 283.74 | 321.09 | 194.06 | 0.97 | 0.30 | 3.28 | 1.05E-03 | 7.19E-02 |
| ENSRNOG00000008956 | Cdkn2c | 1130.69 | 1529.44 | 1079.28 | 918.25 | 931.60 | 682.92 | 1045.36 | -0.56 | 0.17 | -3.27 | 1.06E-03 | 7.23E-02 |
| ENSRNOG00000008450 | LOC100359539 | 3759.02 | 3277.88 | 3758.28 | 3024.61 | 2777.97 | 2942.20 | 3256.66 | -0.30 | 0.09 | -3.27 | 1.07E-03 | 7.24E-02 |
| ENSRNOG00000031033 | Mt-nd2 | 3901.34 | 4080.96 | 4955.76 | 5122.55 | 6293.77 | 5660.32 | 5002.45 | 0.40 | 0.12 | 3.27 | 1.07E-03 | 7.24E-02 |
| ENSRNOG00000015160 | Gem | 211.67 | 284.63 | 294.91 | 181.40 | 179.85 | 132.99 | 214.24 | -0.68 | 0.21 | -3.27 | 1.09E-03 | 7.30E-02 |
| ENSRNOG00000028834 | Polr2a | 1051.71 | 1212.00 | 1500.31 | 1486.75 | 2567.07 | 1829.78 | 1607.94 | 0.64 | 0.20 | 3.26 | 1.10E-03 | 7.33E-02 |
| ENSRNOG00000031053 | Mt-nd4l | 1022.59 | 1114.67 | 1266.02 | 1348.52 | 1644.66 | 1498.94 | 1315.90 | 0.40 | 0.12 | 3.25 | 1.14E-03 | 7.58E-02 |
| ENSRNOG00000024923 | Nnat | 7027.53 | 5032.21 | 8023.65 | 7360.55 | 12701.56 | 11959.71 | 8684.20 | 0.67 | 0.21 | 3.25 | 1.16E-03 | 7.67E-02 |
| ENSRNOG00000042944 | Cenpw | 1209.29 | 1067.89 | 1075.20 | 947.79 | 812.32 | 848.01 | 993.42 | -0.36 | 0.11 | -3.24 | 1.18E-03 | 7.79E-02 |
| ENSRNOG00000013663 | Tmem86a | 510.60 | 279.03 | 414.73 | 526.88 | 799.21 | 672.43 | 533.81 | 0.73 | 0.23 | 3.24 | 1.21E-03 | 7.92E-02 |
| ENSRNOG00000020532 | Kcnq1 | 175.72 | 342.95 | 230.45 | 144.84 | 154.75 | 119.25 | 194.66 | -0.84 | 0.26 | -3.23 | 1.22E-03 | 7.94E-02 |
| ENSRNOG00000015420 | Stxbp1 | 2018.83 | 1765.29 | 2115.28 | 2491.95 | 2871.30 | 2315.02 | 2262.94 | 0.38 | 0.12 | 3.23 | 1.24E-03 | 7.99E-02 |
| ENSRNOG00000037613 | Kdm6b | 145.12 | 141.14 | 208.91 | 237.44 | 357.66 | 243.58 | 222.31 | 0.76 | 0.23 | 3.23 | 1.23E-03 | 7.99E-02 |
| ENSRNOG00000014268 | Abca2 | 2059.68 | 2625.19 | 2306.30 | 2823.69 | 3746.95 | 2953.52 | 2752.55 | 0.45 | 0.14 | 3.22 | 1.27E-03 | 8.16E-02 |
| ENSRNOG00000001288 | Gpr146 | 81.62 | 74.35 | 102.40 | 110.06 | 159.36 | 238.72 | 127.75 | 0.98 | 0.30 | 3.22 | 1.28E-03 | 8.23E-02 |
| ENSRNOG00000001706 | Kalrn | 2189.18 | 1847.55 | 2092.05 | 2656.39 | 2936.63 | 2350.68 | 2345.41 | 0.37 | 0.12 | 3.22 | 1.29E-03 | 8.26E-02 |
| ENSRNOG00000012616 | Ppt1 | 1025.50 | 1143.25 | 951.26 | 815.33 | 817.95 | 818.28 | 928.60 | -0.35 | 0.11 | -3.21 | 1.34E-03 | 8.53E-02 |
| ENSRNOG00000001726 | Tmem44 | 276.15 | 284.46 | 317.45 | 206.88 | 219.76 | 144.00 | 241.45 | -0.63 | 0.20 | -3.20 | 1.36E-03 | 8.58E-02 |
| ENSRNOG00000021314 | Fdft1 | 816.77 | 1024.06 | 686.13 | 624.68 | 535.18 | 605.59 | 715.40 | -0.52 | 0.16 | -3.20 | 1.36E-03 | 8.58E-02 |
| ENSRNOG00000004281 | Cobl | 416.64 | 440.20 | 377.89 | 520.57 | 571.75 | 535.20 | 477.04 | 0.40 | 0.12 | 3.19 | 1.40E-03 | 8.79E-02 |
| ENSRNOG00000002248 | Fryl | 2092.78 | 2397.75 | 2176.25 | 2717.70 | 3005.01 | 2592.34 | 2496.97 | 0.32 | 0.10 | 3.18 | 1.47E-03 | 8.90E-02 |
| ENSRNOG00000014201 | Manf | 4318.82 | 4614.48 | 5242.07 | 4169.84 | 3328.75 | 3129.45 | 4133.90 | -0.42 | 0.13 | -3.18 | 1.45E-03 | 8.90E-02 |
| ENSRNOG00000016684 | Wnk2 | 383.52 | 512.78 | 452.46 | 583.20 | 805.16 | 590.32 | 554.57 | 0.55 | 0.17 | 3.18 | 1.46E-03 | 8.90E-02 |
| ENSRNOG00000023720 | Ntm | 39.84 | 48.59 | 38.91 | 20.38 | 18.12 | 12.77 | 29.77 | -1.32 | 0.42 | -3.18 | 1.48E-03 | 8.90E-02 |
| ENSRNOG00000025551 | Rgs22 | 678.11 | 515.86 | 607.29 | 706.19 | 923.27 | 898.78 | 721.58 | 0.49 | 0.15 | 3.19 | 1.43E-03 | 8.90E-02 |
| ENSRNOG00000026942 | RGD1311595 | 3833.08 | 3411.31 | 4129.20 | 4402.16 | 5245.54 | 4787.81 | 4301.52 | 0.34 | 0.11 | 3.18 | 1.48E-03 | 8.90E-02 |

| | | | | | | | | | | | | | |
|---|---|---|---|---|---|---|---|---|---|---|---|---|---|
| ENSRNOG00000026963 | Hsp90b1 | 68324.65 | 60205.86 | 84641.48 | 58720.39 | 47392.51 | 51606.35 | 61815.21 | -0.43 | 0.14 | -3.18 | 1.46E-03 | 8.90E-02 |
| ENSRNOG00000033556 | Spen | 362.15 | 387.52 | 483.86 | 502.94 | 798.22 | 589.94 | 520.77 | 0.62 | 0.19 | 3.19 | 1.44E-03 | 8.90E-02 |
| ENSRNOG00000059592 | LOC103689960 | 8.54 | 16.52 | 14.34 | 39.75 | 32.59 | 36.44 | 24.70 | 1.47 | 0.46 | 3.18 | 1.46E-03 | 8.90E-02 |
| ENSRNOG00000001982 | Cblb | 134.89 | 135.52 | 157.88 | 204.28 | 218.19 | 206.34 | 176.19 | 0.55 | 0.17 | 3.17 | 1.50E-03 | 8.92E-02 |
| ENSRNOG00000012417 | Iapp | 79691.25 | 77197.19 | 93672.54 | 71038.20 | 58062.59 | 39749.92 | 69901.95 | -0.57 | 0.18 | -3.18 | 1.49E-03 | 8.92E-02 |
| ENSRNOG00000014573 | Ckmt1 | 1426.91 | 1391.77 | 1498.39 | 1263.04 | 974.22 | 1086.73 | 1273.51 | -0.38 | 0.12 | -3.17 | 1.51E-03 | 8.92E-02 |
| ENSRNOG00000051553 | LOC103690980 | 5.70 | 5.53 | 6.14 | 15.29 | 32.53 | 21.80 | 14.50 | 2.00 | 0.63 | 3.17 | 1.51E-03 | 8.92E-02 |
| ENSRNOG00000053047 | Top2a | 28377.50 | 25569.44 | 26059.23 | 23430.48 | 22138.90 | 21791.87 | 24561.24 | -0.25 | 0.08 | -3.17 | 1.51E-03 | 8.92E-02 |
| ENSRNOG00000047045 | LOC108348111 | 1292.18 | 1723.14 | 1245.05 | 1094.66 | 1022.11 | 1009.01 | 1231.02 | -0.45 | 0.14 | -3.17 | 1.52E-03 | 8.96E-02 |
| ENSRNOG00000008445 | Dact1 | 209.68 | 147.67 | 166.92 | 230.31 | 300.73 | 278.07 | 222.23 | 0.62 | 0.20 | 3.17 | 1.53E-03 | 8.97E-02 |
| ENSRNOG00000004234 | Mgat2 | 836.83 | 937.35 | 849.96 | 709.28 | 682.40 | 698.38 | 785.70 | -0.33 | 0.10 | -3.16 | 1.56E-03 | 9.05E-02 |
| ENSRNOG00000018094 | Sv2c | 44.59 | 64.18 | 76.80 | 28.53 | 25.37 | 30.09 | 44.93 | -1.14 | 0.36 | -3.16 | 1.56E-03 | 9.05E-02 |
| ENSRNOG00000000543 | Frk | 69.24 | 88.89 | 62.47 | 108.02 | 129.30 | 144.16 | 100.34 | 0.79 | 0.25 | 3.15 | 1.61E-03 | 9.15E-02 |
| ENSRNOG00000005841 | Erp44 | 3405.86 | 4573.98 | 3759.10 | 3248.98 | 2868.94 | 2752.46 | 3434.89 | -0.40 | 0.13 | -3.15 | 1.61E-03 | 9.15E-02 |
| ENSRNOG00000007587 | Tcp11l2 | 281.81 | 222.92 | 228.36 | 335.28 | 334.52 | 360.07 | 293.83 | 0.49 | 0.16 | 3.16 | 1.60E-03 | 9.15E-02 |
| ENSRNOG00000011800 | F3 | 893.07 | 992.20 | 976.91 | 804.08 | 666.42 | 486.56 | 803.21 | -0.55 | 0.17 | -3.16 | 1.59E-03 | 9.15E-02 |
| ENSRNOG00000011885 | Rhpn2 | 901.19 | 958.96 | 1099.85 | 766.33 | 740.56 | 793.48 | 876.73 | -0.36 | 0.11 | -3.15 | 1.61E-03 | 9.15E-02 |
| ENSRNOG00000030880 | Hs6st2 | 248.63 | 356.96 | 301.06 | 198.72 | 220.96 | 185.00 | 251.89 | -0.59 | 0.19 | -3.16 | 1.59E-03 | 9.15E-02 |
| ENSRNOG00000004921 | Nusap1 | 4678.00 | 4184.36 | 4384.24 | 3852.48 | 3646.53 | 3370.36 | 4019.33 | -0.29 | 0.09 | -3.15 | 1.65E-03 | 9.33E-02 |
| ENSRNOG00000027317 | Ylpm1 | 491.03 | 522.11 | 613.51 | 649.29 | 897.85 | 736.75 | 651.76 | 0.49 | 0.16 | 3.14 | 1.69E-03 | 9.48E-02 |
| ENSRNOG00000057040 | Spns2 | 1763.30 | 1872.50 | 1581.18 | 2114.51 | 2463.66 | 2075.35 | 1978.42 | 0.35 | 0.11 | 3.14 | 1.70E-03 | 9.54E-02 |
| ENSRNOG00000012820 | Add3 | 2174.67 | 2766.19 | 2269.55 | 1892.91 | 1962.28 | 1580.13 | 2107.62 | -0.41 | 0.13 | -3.13 | 1.73E-03 | 9.63E-02 |
| ENSRNOG00000010170 | Tubb4b | 8214.16 | 9004.14 | 7842.86 | 7363.15 | 6478.35 | 6648.89 | 7591.93 | -0.29 | 0.09 | -3.13 | 1.76E-03 | 9.68E-02 |
| ENSRNOG00000018391 | Cdk2ap2 | 668.73 | 706.49 | 649.16 | 561.58 | 526.91 | 449.28 | 593.69 | -0.40 | 0.13 | -3.13 | 1.76E-03 | 9.68E-02 |
| ENSRNOG00000018730 | Nectin2 | 423.37 | 400.53 | 493.77 | 589.04 | 578.18 | 569.10 | 509.00 | 0.40 | 0.13 | 3.13 | 1.76E-03 | 9.68E-02 |
| ENSRNOG00000020836 | Rorc | 349.11 | 179.71 | 154.63 | 395.40 | 364.82 | 491.55 | 322.54 | 0.87 | 0.28 | 3.13 | 1.76E-03 | 9.68E-02 |
| ENSRNOG00000006729 | Slc24a4 | 20.30 | 34.29 | 33.31 | 6.24 | 7.98 | 12.12 | 19.04 | -1.73 | 0.55 | -3.13 | 1.77E-03 | 9.68E-02 |
| ENSRNOG00000002496 | Stxbp5l | 261.82 | 305.28 | 353.31 | 371.95 | 482.04 | 482.49 | 376.15 | 0.54 | 0.17 | 3.12 | 1.78E-03 | 9.71E-02 |
| ENSRNOG00000003769 | Tmem163 | 509.65 | 491.07 | 683.02 | 448.41 | 310.27 | 365.37 | 467.96 | -0.58 | 0.19 | -3.12 | 1.83E-03 | 9.94E-02 |
| ENSRNOG00000024905 | Drc1 | 54.08 | 59.62 | 58.37 | 76.43 | 130.44 | 111.23 | 81.69 | 0.88 | 0.28 | 3.11 | 1.84E-03 | 9.96E-02 |

**Table S3. Significant differentially expressed small RNAs (DE sRNAs) in PWS *vs.* control INS-1 lines from small RNA-seq.**

| gene_ID | gene_name | Control 5-9 | Control 2 | Control 16 | PWS 3 | PWS 19-1 | PWS 19-4 | baseMean | log2FC | lfcSE | pvalue | padj |
|---|---|---|---|---|---|---|---|---|---|---|---|---|
| ENSRNOG00000046685 | Mir344a-2 | 1293.19 | 879.80 | 987.64 | 1.19 | 0.00 | 1.32 | 527.19 | -10.08 | 0.85 | 4.89E-31 | 4.35E-27 |
| ENSRNOG00000049433 | Mir344a-1 | 1259.71 | 877.19 | 1040.11 | 0.59 | 0.00 | 1.32 | 529.82 | -10.61 | 1.01 | 4.33E-25 | 1.93E-21 |
| ENSRNOG00000059940 | Snord116 | 765.28 | 436.63 | 671.29 | 0.00 | 0.00 | 0.00 | 312.20 | -13.22 | 3.28 | 2.61E-22 | 7.74E-19 |
| ENSRNOG00000051811 | Snord116 | 708.15 | 415.72 | 668.20 | 0.00 | 0.00 | 0.00 | 298.68 | 0.00 | 0.00 | 4.36E-22 | 9.70E-19 |
| ENSRNOG00000061924 | Snord64 AABR07005711.2 | 560.42 | 486.31 | 578.70 | 0.00 | 0.00 | 0.00 | 270.90 | -13.00 | 3.26 | 7.41E-22 | 1.27E-18 |
| ENSRNOG00000055348 | Snord64 AABR07005710.1 | 542.69 | 466.70 | 608.79 | 0.00 | 0.00 | 0.00 | 269.70 | 0.00 | 0.00 | 8.53E-22 | 1.27E-18 |
| ENSRNOG00000058825 | Snord116 | 687.47 | 420.95 | 660.48 | 0.59 | 0.00 | 0.00 | 294.92 | -10.61 | 1.34 | 1.52E-19 | 1.94E-16 |
| ENSRNOG00000054305 | Snord116 | 635.27 | 383.03 | 644.28 | 0.00 | 0.00 | 1.32 | 277.32 | 0.00 | 0.00 | 3.36E-19 | 3.74E-16 |
| ENSRNOG00000053918 | Snord116 | 340.78 | 251.00 | 307.09 | 0.00 | 0.00 | 0.00 | 149.81 | -12.04 | 3.16 | 8.12E-19 | 8.03E-16 |
| ENSRNOG00000061669 | Snord116 | 334.87 | 198.71 | 349.53 | 0.00 | 0.00 | 0.00 | 147.19 | -12.00 | 3.16 | 1.58E-18 | 1.41E-15 |
| ENSRNOG00000052552 | Snord116 | 344.72 | 192.17 | 335.64 | 0.00 | 0.00 | 0.00 | 145.42 | -11.98 | 3.16 | 1.91E-18 | 1.54E-15 |
| ENSRNOG00000052740 | Snord116 | 309.26 | 183.02 | 266.97 | 0.00 | 0.00 | 0.00 | 126.54 | -11.74 | 3.12 | 7.23E-18 | 5.37E-15 |
| ENSRNOG00000055753 | Snord116 | 265.93 | 186.94 | 240.74 | 0.00 | 0.00 | 0.00 | 115.60 | -11.60 | 3.10 | 1.52E-17 | 1.04E-14 |
| ENSRNOG00000051750 | Snord116 | 417.60 | 290.22 | 342.59 | 0.00 | 1.05 | 0.00 | 175.24 | -9.82 | 1.33 | 3.76E-17 | 2.25E-14 |
| ENSRNOG00000052187 | Snord116 | 229.49 | 197.40 | 201.39 | 0.00 | 0.00 | 0.00 | 104.71 | -11.44 | 3.09 | 3.80E-17 | 2.25E-14 |
| ENSRNOG00000053521 | Snord116 | 261.99 | 168.64 | 183.64 | 0.00 | 0.00 | 0.00 | 102.38 | -11.40 | 3.09 | 6.71E-17 | 3.73E-14 |
| ENSRNOG00000051822 | Snord116 | 354.57 | 241.85 | 413.57 | 0.59 | 0.00 | 0.00 | 168.43 | -9.78 | 1.33 | 7.41E-17 | 3.88E-14 |
| ENSRNOG00000059154 | Snord116 | 246.23 | 149.03 | 185.95 | 0.00 | 0.00 | 0.00 | 96.87 | -11.31 | 3.08 | 1.25E-16 | 6.20E-14 |
| ENSRNOG00000060158 | Snord116 | 366.39 | 217.01 | 382.71 | 0.00 | 0.00 | 1.32 | 161.24 | -9.68 | 1.33 | 1.34E-16 | 6.28E-14 |
| ENSRNOG00000061909 | Snord116 | 340.78 | 234.00 | 330.24 | 0.00 | 0.00 | 1.32 | 151.06 | -9.59 | 1.33 | 1.95E-16 | 8.67E-14 |
| ENSRNOG00000057393 | Snord116 | 205.85 | 129.42 | 201.39 | 0.00 | 0.00 | 0.00 | 89.44 | -11.18 | 3.07 | 2.89E-16 | 1.23E-13 |
| ENSRNOG00000058701 | Snord107 | 203.88 | 172.56 | 274.69 | 0.59 | 0.00 | 1.32 | 108.84 | -8.26 | 1.01 | 8.39E-16 | 3.40E-13 |
| ENSRNOG00000056157 | Snord116 | 161.53 | 91.51 | 155.86 | 0.00 | 0.00 | 0.00 | 68.15 | -10.73 | 3.03 | 5.70E-15 | 2.21E-12 |
| ENSRNOG00000056713 | Snord116 | 150.69 | 103.28 | 111.11 | 0.00 | 0.00 | 0.00 | 60.85 | -10.53 | 3.00 | 1.44E-14 | 5.34E-12 |
| ENSRNOG00000057005 | Snord116 | 143.80 | 91.51 | 128.08 | 0.00 | 0.00 | 0.00 | 60.57 | -10.53 | 3.00 | 1.56E-14 | 5.58E-12 |
| ENSRNOG00000053726 | Snord116 | 151.68 | 94.12 | 109.57 | 0.00 | 0.00 | 0.00 | 59.23 | -10.49 | 2.99 | 2.08E-14 | 7.12E-12 |
| ENSRNOG00000049100 | Mir344b-1 | 235.39 | 160.80 | 174.38 | 0.00 | 0.00 | 1.32 | 95.31 | -8.89 | 1.33 | 2.30E-14 | 7.58E-12 |
| ENSRNOG00000054389 | Snord116 | 229.49 | 151.64 | 182.87 | 0.59 | 0.00 | 0.00 | 94.10 | -8.90 | 1.33 | 2.57E-14 | 8.18E-12 |
| ENSRNOG00000059567 | Snord116 | 125.08 | 87.59 | 124.23 | 0.00 | 0.00 | 0.00 | 56.15 | -10.40 | 2.98 | 3.16E-14 | 9.71E-12 |
| ENSRNOG00000053379 | Snord116 | 144.78 | 90.20 | 102.62 | 0.00 | 0.00 | 0.00 | 56.27 | -10.40 | 2.98 | 3.52E-14 | 1.01E-11 |
| ENSRNOG00000056644 | Snord116 | 140.84 | 87.59 | 108.02 | 0.00 | 0.00 | 0.00 | 56.08 | -10.40 | 2.98 | 3.53E-14 | 1.01E-11 |
| ENSRNOG00000059451 | Snord116 | 95.54 | 75.82 | 97.22 | 0.00 | 0.00 | 0.00 | 44.76 | -10.02 | 2.94 | 2.83E-13 | 7.87E-11 |
| ENSRNOG00000060200 | Snord116 | 90.61 | 75.82 | 100.31 | 0.00 | 0.00 | 0.00 | 44.46 | -10.00 | 2.94 | 3.04E-13 | 8.21E-11 |
| ENSRNOG00000053833 | Snord116 | 94.55 | 65.36 | 109.57 | 0.00 | 0.00 | 0.00 | 44.91 | -10.02 | 2.94 | 3.35E-13 | 8.65E-11 |
| ENSRNOG00000058848 | Snord116 | 165.47 | 100.66 | 176.70 | 0.00 | 1.05 | 0.00 | 73.98 | -8.51 | 1.33 | 3.40E-13 | 8.65E-11 |
| ENSRNOG00000049846 | Mir344g | 152.66 | 118.96 | 145.06 | 0.59 | 0.00 | 0.00 | 69.55 | -8.44 | 1.32 | 4.44E-13 | 1.10E-10 |

| | | | | | | | | | | | | |
|---|---|---|---|---|---|---|---|---|---|---|---|---|
| ENSRNOG00000059608 | Snord116 | 122.13 | 57.52 | 80.25 | 0.00 | 0.00 | 0.00 | 43.32 | -9.95 | 2.94 | 6.60E-13 | 1.59E-10 |
| ENSRNOG00000052049 | Snord116 | 84.70 | 47.06 | 50.93 | 0.00 | 0.00 | 0.00 | 30.45 | -9.34 | 2.87 | 1.74E-11 | 4.08E-09 |
| ENSRNOG00000058910 | Snord116 | 55.16 | 30.07 | 44.75 | 0.00 | 0.00 | 0.00 | 21.66 | -8.75 | 2.81 | 3.80E-10 | 8.69E-08 |
| ENSRNOG00000058803 | Snord115 | 47.28 | 20.92 | 54.78 | 0.00 | 0.00 | 0.00 | 20.50 | -8.65 | 2.81 | 8.95E-10 | 1.99E-07 |
| ENSRNOG00000059948 | Snord115 | 46.29 | 18.30 | 53.24 | 0.00 | 0.00 | 0.00 | 19.64 | -8.57 | 2.81 | 1.48E-09 | 3.21E-07 |
| ENSRNOG00000058531 | Snord115 | 37.43 | 27.45 | 42.44 | 0.00 | 0.00 | 0.00 | 17.89 | -8.41 | 2.78 | 1.93E-09 | 4.09E-07 |
| ENSRNOG00000056852 | Snord115 | 40.38 | 24.84 | 39.35 | 0.00 | 0.00 | 0.00 | 17.43 | -8.36 | 2.78 | 2.57E-09 | 5.31E-07 |
| ENSRNOG00000052538 | Snord115 | 23.64 | 15.69 | 30.86 | 0.00 | 0.00 | 0.00 | 11.70 | 0.00 | 0.00 | 8.98E-08 | 1.82E-05 |
| ENSRNOG00000059788 | Snord115 | 19.70 | 14.38 | 24.69 | 0.00 | 0.00 | 0.00 | 9.79 | 0.00 | 0.00 | 3.69E-07 | 7.30E-05 |
| ENSRNOG00000050488 | miR-344 AABR07003935.14 | 16.74 | 18.30 | 14.66 | 0.00 | 0.00 | 0.00 | 8.28 | 0.00 | 0.00 | 1.37E-06 | 2.65E-04 |
| ENSRNOG00000061343 | Snord115 | 21.67 | 7.84 | 20.83 | 0.00 | 0.00 | 0.00 | 8.39 | 0.00 | 0.00 | 1.69E-06 | 3.21E-04 |
| ENSRNOG00000047759 | Mir344b-3 | 26.59 | 16.99 | 7.72 | 0.00 | 0.00 | 0.00 | 8.55 | 0.00 | 0.00 | 1.85E-06 | 3.44E-04 |
| ENSRNOG00000049216 | Mir344b-2 | 12.80 | 19.61 | 14.66 | 0.00 | 0.00 | 0.00 | 7.85 | 0.00 | 0.00 | 2.25E-06 | 4.09E-04 |
| ENSRNOG00000061156 | Snord116 | 18.71 | 6.54 | 23.15 | 0.00 | 0.00 | 0.00 | 8.07 | 0.00 | 0.00 | 2.58E-06 | 4.60E-04 |
| ENSRNOG00000046092 | miR-344 AABR07003935.1 | 18.71 | 18.30 | 8.49 | 0.00 | 0.00 | 0.00 | 7.58 | 0.00 | 0.00 | 3.63E-06 | 6.34E-04 |
| ENSRNOG00000051533 | Snord116 | 14.77 | 3.92 | 23.15 | 0.00 | 0.00 | 0.00 | 6.97 | 0.00 | 0.00 | 1.02E-05 | 1.75E-03 |
| ENSRNOG00000035621 | Mir135b | 94.55 | 346.43 | 81.02 | 22.57 | 40.97 | 52.73 | 106.38 | -1.92 | 0.54 | 1.19E-05 | 2.01E-03 |
| ENSRNOG00000046914 | miR-344 AABR07003935.4 | 17.73 | 11.77 | 8.49 | 0.00 | 0.00 | 0.00 | 6.33 | 0.00 | 0.00 | 1.29E-05 | 2.13E-03 |
| ENSRNOG00000059473 | Snord115 | 22.65 | 11.77 | 19.29 | 0.00 | 0.00 | 1.32 | 9.17 | -5.03 | 1.44 | 1.63E-05 | 2.65E-03 |
| ENSRNOG00000035465 | Mir3065 | 653.98 | 632.73 | 668.20 | 377.13 | 310.95 | 266.30 | 484.88 | 0.00 | 0.00 | 4.66E-05 | 7.41E-03 |
| ENSRNOG00000047713 | miR-344 AABR07003935.6 | 11.82 | 9.15 | 9.26 | 0.00 | 0.00 | 0.00 | 5.04 | 0.00 | 0.00 | 5.69E-05 | 8.89E-03 |
| ENSRNOG00000035476 | Mir212 | 1953.09 | 1261.53 | 2612.62 | 1121.90 | 814.14 | 942.61 | 1450.98 | 0.00 | 0.00 | 6.31E-04 | 9.70E-02 |

**Table S4. Highly expressed miRNAs in INS-1 lines from small RNA-seq.**

| gene_ID | gene_name | Control 5-9 | Control 2 | Control 16 | PWS 3 | PWS 19-1 | PWS 19-4 | Control_AVG |
|---|---|---|---|---|---|---|---|---|
| ENSRNOG00000035613 | Mir7a1 | 1258260.92 | 620376.88 | 1121962.86 | 922445.32 | 1001917.94 | 740374.29 | 1000200.22 |
| ENSRNOG00000035477 | Mir7a2 | 1264776.14 | 623497.37 | 1105213.09 | 929051.41 | 1007153.68 | 749767.44 | 997828.87 |
| ENSRNOG00000035504 | Mir16 | 671204.08 | 859472.60 | 653032.62 | 650940.59 | 789242.34 | 828869.00 | 727903.10 |
| ENSRNOG00000035455 | Mir375 | 806138.47 | 657529.89 | 697051.31 | 683473.93 | 833397.35 | 558021.87 | 720239.89 |
| ENSRNOG00000035464 | Mirlet7f2 | 660821.10 | 635055.06 | 547815.72 | 502281.91 | 597436.16 | 469426.97 | 614563.96 |
| ENSRNOG00000046475 | Mir148a | 532328.05 | 602307.60 | 705603.66 | 501868.55 | 466810.59 | 751290.12 | 613413.10 |
| ENSRNOG00000035642 | Mirlet7i | 527687.13 | 580125.60 | 476862.22 | 450704.17 | 594884.48 | 456635.14 | 528224.99 |
| ENSRNOG00000035451 | Mir183 | 355488.46 | 390677.92 | 322713.90 | 322305.04 | 352005.69 | 523439.25 | 356293.43 |
| ENSRNOG00000035611 | Mir30d | 331949.97 | 326251.04 | 311342.14 | 304930.72 | 329561.56 | 271312.50 | 323181.05 |
| ENSRNOG00000035576 | Mir27b | 293892.84 | 300509.31 | 268982.39 | 261355.95 | 279482.73 | 266739.19 | 287794.85 |
| ENSRNOG00000035531 | Mir25 | 295459.84 | 268959.32 | 248187.15 | 266792.03 | 257743.49 | 325939.13 | 270868.77 |
| ENSRNOG00000035485 | Mir93 | 287324.44 | 246560.31 | 232232.12 | 270585.95 | 250824.84 | 328984.49 | 255372.29 |
| ENSRNOG00000035632 | Mirlet7a1 | 151437.63 | 171839.88 | 137001.19 | 136415.17 | 172216.28 | 134147.42 | 153426.23 |
| ENSRNOG00000035500 | Mirlet7c1 | 150681.22 | 161007.72 | 123913.41 | 115417.39 | 138760.72 | 87567.92 | 145200.78 |
| ENSRNOG00000035461 | Mirlet7c2 | 150459.61 | 157666.30 | 122673.46 | 114771.22 | 138398.29 | 87152.64 | 143599.79 |
| ENSRNOG00000035532 | Mir101-2 | 160343.24 | 124559.33 | 141000.36 | 138142.86 | 118370.33 | 149291.16 | 141967.64 |
| ENSRNOG00000035496 | Mir103a2 | 145277.97 | 126300.63 | 134246.60 | 123091.34 | 137590.45 | 107780.67 | 135275.07 |
| ENSRNOG00000035503 | Mir103a1 | 145311.46 | 125543.72 | 131738.15 | 122598.39 | 138328.96 | 108513.66 | 134197.77 |
| ENSRNOG00000035636 | Mirlet7b | 126037.65 | 138962.99 | 107689.10 | 114365.58 | 135406.44 | 91828.79 | 124229.91 |
| ENSRNOG00000047216 | Mir26a-2 | 139891.47 | 115264.54 | 112694.43 | 103268.91 | 110891.76 | 79975.62 | 122616.81 |
| ENSRNOG00000035582 | Mir26a | 140129.82 | 113638.28 | 110443.69 | 103256.44 | 110360.20 | 79300.63 | 121403.93 |
| ENSRNOG00000035530 | Mir192 | 158732.90 | 66206.09 | 122627.16 | 159281.40 | 164949.91 | 134707.71 | 115855.39 |
| ENSRNOG00000035572 | Mir125a | 101528.03 | 121135.56 | 102998.57 | 96403.88 | 97683.70 | 54751.87 | 108554.05 |
| ENSRNOG00000035525 | Mir148b | 109013.39 | 95056.55 | 100882.09 | 93152.80 | 96969.36 | 94671.12 | 101650.68 |
| ENSRNOG00000035577 | Mir141 | 87613.15 | 51925.32 | 91999.50 | 70813.97 | 64630.48 | 68055.21 | 77179.32 |
| ENSRNOG00000035654 | Mir99b | 71887.00 | 76151.91 | 74387.86 | 72154.43 | 76547.46 | 67701.90 | 74142.26 |
| ENSRNOG00000035542 | Mir129-2 | 72876.84 | 66951.25 | 59202.88 | 54426.14 | 59692.04 | 56260.05 | 66343.66 |
| ENSRNOG00000035598 | Mir129-1 | 70351.52 | 64464.79 | 56871.13 | 51813.52 | 57329.44 | 54456.56 | 63895.81 |
| ENSRNOG00000035646 | Mir30a | 61362.20 | 45497.41 | 50987.72 | 46807.43 | 46306.45 | 44778.65 | 52615.78 |
| ENSRNOG00000035555 | Mir384 | 57926.81 | 42967.81 | 51436.02 | 49313.74 | 44314.68 | 25430.73 | 50776.88 |
| ENSRNOG00000035612 | Mir342 | 51109.23 | 49343.44 | 46443.03 | 45527.55 | 51959.24 | 35193.02 | 48965.23 |
| ENSRNOG00000035480 | Mir21 | 51704.12 | 42370.38 | 52196.04 | 47635.94 | 43752.66 | 56008.24 | 48756.85 |
| ENSRNOG00000035484 | Mir425 | 49951.95 | 39383.24 | 47303.36 | 45609.51 | 45201.31 | 30358.67 | 45546.18 |

| | | | | | | | | |
|---|---|---|---|---|---|---|---|---|
| ENSRNOG00000047385 | Mir340-2 | 56806.96 | 29668.81 | 44308.03 | 47134.68 | 45419.82 | 37584.48 | 43594.60 |
| ENSRNOG00000045609 | Mir340-1 | 56280.03 | 29332.84 | 43409.89 | 46681.52 | 45114.12 | 36999.14 | 43007.59 |
| ENSRNOG00000035467 | Mir23b | 45390.81 | 42548.17 | 34048.91 | 33744.92 | 33936.70 | 15290.08 | 40662.63 |
| ENSRNOG00000035594 | Mirlet7d | 42416.37 | 38719.14 | 34435.48 | 35821.83 | 38406.62 | 25297.58 | 38523.66 |
| ENSRNOG00000050242 | Mirlet7g | 41139.92 | 35160.71 | 34574.37 | 32681.82 | 35422.12 | 22009.64 | 36958.33 |
| ENSRNOG00000035640 | Mir30e | 39222.29 | 36640.56 | 31228.73 | 31773.73 | 31713.82 | 26104.40 | 35697.19 |
| ENSRNOG00000035516 | Mir132 | 28314.35 | 21925.77 | 41186.93 | 20748.33 | 17319.75 | 20481.69 | 30475.68 |
| ENSRNOG00000035609 | Mir140 | 30725.42 | 25422.75 | 27968.75 | 29418.86 | 28657.89 | 27045.69 | 28038.97 |
| ENSRNOG00000035540 | Mir106b | 31067.19 | 26880.37 | 25736.53 | 27915.08 | 26785.88 | 32282.13 | 27894.70 |
| ENSRNOG00000035608 | Mir26b | 31413.88 | 27684.35 | 24282.84 | 27776.10 | 29528.76 | 17847.65 | 27793.69 |
| ENSRNOG00000035624 | Mir96 | 22781.09 | 17005.15 | 20403.27 | 19312.85 | 14593.67 | 10107.69 | 20063.17 |
| ENSRNOG00000040448 | Mir146b | 20091.29 | 17233.93 | 18288.33 | 17765.71 | 20767.52 | 14050.84 | 18537.85 |
| ENSRNOG00000040357 | Mir672 | 15998.97 | 21381.94 | 16998.22 | 15103.20 | 17631.75 | 21926.59 | 18126.38 |
| ENSRNOG00000055316 | Mir1839 | 21067.34 | 14312.15 | 18652.52 | 16283.30 | 14677.72 | 14111.48 | 18010.67 |
| ENSRNOG00000035656 | Mir98 | 18771.50 | 18651.02 | 15598.55 | 17461.62 | 20058.42 | 14632.23 | 17673.69 |
| ENSRNOG00000035614 | Mir196a | 16969.11 | 19024.90 | 15657.19 | 11189.91 | 14878.36 | 20800.73 | 17217.07 |
| ENSRNOG00000035472 | Mir204 | 15144.06 | 17162.03 | 13918.02 | 12800.00 | 14078.92 | 7725.46 | 15408.03 |
| ENSRNOG00000046413 | rno-mir-598-1 | 16845.99 | 15092.59 | 12309.24 | 11376.99 | 11230.99 | 9095.21 | 14749.28 |
| ENSRNOG00000046841 | Mir598 | 16500.29 | 14982.78 | 12178.07 | 11219.60 | 11056.61 | 8819.68 | 14553.71 |
| ENSRNOG00000041219 | Mir872 | 10100.31 | 21470.83 | 11059.26 | 9637.42 | 13202.80 | 15519.47 | 14210.13 |
| ENSRNOG00000035479 | Mir339 | 14361.05 | 10281.79 | 13202.75 | 15653.75 | 19382.95 | 28534.09 | 12615.20 |
| ENSRNOG00000036245 | Mir532 | 13655.85 | 8207.13 | 12917.26 | 9523.39 | 7240.11 | 6434.80 | 11593.41 |
| ENSRNOG00000035588 | Mir194-2 | 15733.04 | 6827.94 | 11621.75 | 13585.75 | 13024.21 | 8636.43 | 11394.25 |
| ENSRNOG00000035544 | Mir361 | 12300.61 | 9885.68 | 11692.74 | 12595.70 | 12489.50 | 8031.31 | 11293.01 |
| ENSRNOG00000035501 | Mir24-2 | 10950.29 | 13187.88 | 8938.92 | 8681.22 | 11315.04 | 5517.24 | 11025.70 |
| ENSRNOG00000035599 | Mir101a | 11287.13 | 10356.30 | 10344.76 | 9906.46 | 8113.08 | 9509.17 | 10662.73 |
| ENSRNOG00000035639 | Mir107 | 10256.91 | 10804.70 | 8979.81 | 8213.22 | 9606.91 | 4829.07 | 10013.81 |
| ENSRNOG00000035482 | Mir194-1 | 12898.45 | 5489.29 | 9626.41 | 10749.22 | 9263.39 | 5319.49 | 9338.05 |
| ENSRNOG00000035617 | Mir186 | 10138.72 | 7408.38 | 9071.63 | 10035.93 | 9170.95 | 5182.38 | 8872.91 |
| ENSRNOG00000046704 | Mir3099 | 8462.39 | 9272.57 | 8655.74 | 9171.20 | 9313.81 | 7026.74 | 8796.90 |
| ENSRNOG00000035550 | Mir125b1 | 7973.87 | 7926.06 | 5972.14 | 6275.87 | 8429.29 | 3493.59 | 7290.69 |
| ENSRNOG00000035539 | Mir152 | 8049.71 | 4321.88 | 8641.08 | 8412.18 | 5142.24 | 6765.71 | 7004.23 |
| ENSRNOG00000035637 | Mir29b2 | 7367.17 | 4761.13 | 6631.08 | 5384.41 | 5600.26 | 3320.89 | 6253.13 |
| ENSRNOG00000049256 | Mir29b-3 | 7268.68 | 4949.38 | 6536.17 | 5253.16 | 5521.48 | 3334.08 | 6251.41 |

| | | | | | | | | |
|---|---|---|---|---|---|---|---|---|
| ENSRNOG00000035463 | Mir29b1 | 7119.95 | 4736.29 | 6570.90 | 5176.54 | 5494.16 | 3260.25 | 6142.38 |
| ENSRNOG00000053467 | Mir30b | 9174.49 | 4055.20 | 4918.91 | 4960.95 | 4595.98 | 2118.57 | 6049.53 |
| ENSRNOG00000035585 | Mir185 | 6382.25 | 5453.99 | 5699.00 | 6468.90 | 7249.56 | 3630.70 | 5845.08 |
| ENSRNOG00000035534 | Mir128-1 | 5930.18 | 5891.93 | 5368.75 | 5044.69 | 6016.27 | 3245.75 | 5730.29 |
| ENSRNOG00000049112 | Mir744 | 4694.11 | 7601.86 | 4717.53 | 5694.43 | 7494.33 | 5501.42 | 5671.16 |
| ENSRNOG00000040346 | Mir671 | 5791.30 | 5952.06 | 5227.55 | 6290.13 | 6775.78 | 4250.32 | 5656.97 |
| ENSRNOG00000061112 | Mir676 | 5456.43 | 5393.85 | 5708.25 | 5390.94 | 5155.90 | 4602.32 | 5519.51 |
| ENSRNOG00000050872 | Mir3559 | 6246.33 | 4261.75 | 4879.56 | 5253.16 | 7311.54 | 6536.32 | 5129.21 |
| ENSRNOG00000035644 | Mir23a | 6044.43 | 4334.95 | 4987.59 | 5430.14 | 4229.35 | 2169.98 | 5122.32 |
| ENSRNOG00000036413 | Mir328 | 3656.99 | 7088.09 | 3846.40 | 4324.87 | 5173.76 | 3753.31 | 4863.83 |
| ENSRNOG00000035562 | Mir30c2 | 6225.65 | 4524.51 | 3694.39 | 3268.89 | 4177.88 | 1879.95 | 4814.85 |
| ENSRNOG00000057450 | Mir196b-2 | 4434.09 | 5396.47 | 4550.86 | 4617.08 | 8343.14 | 10551.97 | 4793.81 |
| ENSRNOG00000049663 | Mir196b-2 | 4352.34 | 5481.44 | 4507.65 | 4569.56 | 8388.32 | 10359.49 | 4780.48 |
| ENSRNOG00000049258 | Mir196b | 4341.51 | 5365.09 | 4604.10 | 4576.10 | 8284.32 | 10354.22 | 4770.23 |
| ENSRNOG00000035607 | Mir128-2 | 5060.50 | 4657.85 | 4410.43 | 4173.42 | 5069.76 | 2578.67 | 4709.59 |
| ENSRNOG00000035567 | Mir30c1 | 5924.27 | 4218.61 | 3321.71 | 3006.98 | 3857.47 | 1560.91 | 4488.19 |
| ENSRNOG00000035522 | Mir184 | 4516.82 | 4585.95 | 4115.68 | 5095.18 | 3867.98 | 5421.00 | 4406.15 |
| ENSRNOG00000040434 | Mir582 | 6103.52 | 2494.30 | 4600.24 | 4952.04 | 4155.81 | 4702.51 | 4399.35 |
| ENSRNOG00000035627 | Mir325 | 5343.17 | 3414.63 | 4212.90 | 3842.02 | 4176.82 | 2902.98 | 4323.57 |
| ENSRNOG00000035478 | Mir34c | 4894.04 | 3723.15 | 4298.55 | 3610.39 | 4627.49 | 4078.94 | 4305.25 |
| ENSRNOG00000036417 | Mir181d | 5003.37 | 3482.60 | 4352.56 | 3700.07 | 3667.33 | 3974.79 | 4279.51 |
| ENSRNOG00000036423 | Mir92b | 3897.31 | 4159.78 | 4590.21 | 4573.13 | 4572.87 | 7278.54 | 4215.77 |
| ENSRNOG00000035483 | Mir15b | 5100.88 | 3844.72 | 3438.22 | 3902.01 | 3966.72 | 2011.78 | 4127.94 |
| ENSRNOG00000041321 | Mir879 | 4727.59 | 3929.70 | 3215.23 | 3748.18 | 3588.54 | 3497.55 | 3957.51 |
| ENSRNOG00000035537 | Mir423 | 3160.59 | 4040.82 | 4062.44 | 3782.03 | 5153.80 | 6495.45 | 3754.62 |
| ENSRNOG00000035601 | Mir139 | 4366.13 | 2963.61 | 2998.42 | 4462.66 | 3631.61 | 4122.44 | 3442.72 |
| ENSRNOG00000035569 | Mir181c | 3877.61 | 2476.00 | 3228.35 | 2626.88 | 2212.37 | 1500.27 | 3193.99 |
| ENSRNOG00000036265 | Mir488 | 3074.91 | 2451.16 | 2825.58 | 2829.99 | 2332.13 | 2402.01 | 2783.88 |
| ENSRNOG00000036266 | Mir455 | 2677.00 | 3504.83 | 2102.59 | 2382.78 | 2413.02 | 1067.85 | 2761.47 |
| ENSRNOG00000035512 | Mir32 | 2677.00 | 2643.33 | 2912.77 | 2495.62 | 1647.20 | 1837.76 | 2744.37 |
| ENSRNOG00000035519 | Mir324 | 2618.89 | 2317.81 | 2300.89 | 2095.92 | 2690.35 | 2076.38 | 2412.53 |
| ENSRNOG00000040458 | Mir652 | 2009.23 | 2104.73 | 2272.35 | 2214.11 | 3396.30 | 3303.75 | 2128.77 |
| ENSRNOG00000035586 | Mir301a | 2358.87 | 1813.20 | 2080.99 | 2217.08 | 2071.60 | 1446.22 | 2084.36 |
| ENSRNOG00000035597 | Mir27a | 2343.11 | 1700.78 | 1834.85 | 2232.52 | 2052.70 | 1501.59 | 1959.58 |

| | | | | | | | | |
|---|---|---|---|---|---|---|---|---|
| ENSRNOG00000035476 | Mir212 | 1953.09 | 1261.53 | 2612.62 | 1121.90 | 814.14 | 942.61 | 1942.41 |
| ENSRNOG00000036246 | Mir483 | 1610.34 | 2300.82 | 1793.96 | 1754.42 | 1894.07 | 1186.50 | 1901.70 |
| ENSRNOG00000035638 | Mir195 | 1889.07 | 2316.51 | 1403.53 | 1902.90 | 2032.74 | 1202.32 | 1869.70 |
| ENSRNOG00000040376 | Mir802 | 1945.21 | 1362.19 | 1791.64 | 2110.77 | 2311.12 | 1597.82 | 1699.68 |
| ENSRNOG00000035629 | Mir320a | 1591.62 | 1733.46 | 1746.89 | 1858.95 | 2449.79 | 2258.31 | 1690.66 |
| ENSRNOG00000035618 | Mir345 | 1904.83 | 1584.43 | 1449.83 | 1602.97 | 1240.65 | 1491.04 | 1646.36 |
| ENSRNOG00000035481 | Mir218-2 | 1854.60 | 1823.66 | 1214.49 | 1764.51 | 2491.81 | 1394.80 | 1630.92 |
| ENSRNOG00000036562 | Mir374b | 1965.89 | 1015.76 | 1568.65 | 1611.28 | 1363.56 | 1111.36 | 1516.77 |
| ENSRNOG00000035543 | Mir331 | 1786.64 | 1394.87 | 1340.26 | 1501.41 | 1890.92 | 1232.65 | 1507.26 |
| ENSRNOG00000035521 | Mir187 | 1270.54 | 2002.76 | 1039.34 | 1134.37 | 1216.49 | 1111.36 | 1437.55 |
| ENSRNOG00000035570 | Mir130a | 1937.33 | 1092.89 | 1259.24 | 1535.86 | 1380.37 | 835.83 | 1429.82 |
| ENSRNOG00000035589 | Mir383 | 1103.11 | 1588.35 | 1070.20 | 845.73 | 902.39 | 311.13 | 1253.89 |
| ENSRNOG00000040385 | Mir760 | 968.17 | 1204.01 | 1036.25 | 1112.99 | 1534.79 | 1384.25 | 1069.48 |
| ENSRNOG00000049433 | Mir344a-1 | 1259.71 | 877.19 | 1040.11 | 0.59 | 0.00 | 1.32 | 1059.00 |
| ENSRNOG00000046685 | Mir344a-2 | 1293.19 | 879.80 | 987.64 | 1.19 | 0.00 | 1.32 | 1053.55 |
| ENSRNOG00000050903 | mir-378 AABR07030200.1 | 1212.43 | 818.36 | 1106.47 | 874.24 | 755.32 | 850.33 | 1045.75 |
| ENSRNOG00000041292 | miR17 AABR07041787.1 | 1247.89 | 465.39 | 1379.61 | 1650.48 | 1535.84 | 1564.87 | 1030.96 |
| ENSRNOG00000041535 | Mir1224 | 914.99 | 1058.90 | 982.24 | 1074.98 | 1373.02 | 1502.90 | 985.38 |
| ENSRNOG00000040447 | Mir708 | 510.19 | 1249.76 | 1193.66 | 494.14 | 423.36 | 881.97 | 984.54 |
| ENSRNOG00000049078 | Mir7b | 1288.27 | 640.57 | 1000.76 | 1100.52 | 1652.45 | 939.97 | 976.53 |
| ENSRNOG00000036338 | Mir702 | 1124.77 | 873.27 | 895.05 | 1076.17 | 1200.73 | 1156.18 | 964.36 |
| ENSRNOG00000035536 | Mir28 | 1014.46 | 878.49 | 985.33 | 1190.79 | 939.16 | 1372.39 | 959.43 |
| ENSRNOG00000046719 | Mir664-2 | 1034.16 | 878.49 | 918.97 | 846.33 | 839.36 | 912.29 | 943.87 |
| ENSRNOG00000041223 | Mir877 | 962.26 | 956.93 | 910.48 | 1057.76 | 1036.85 | 1686.15 | 943.23 |
| ENSRNOG00000047763 | Mir3547 | 874.60 | 732.08 | 718.35 | 820.79 | 896.08 | 562.93 | 775.01 |
| ENSRNOG00000035546 | Mir421 | 869.68 | 628.80 | 789.34 | 884.34 | 815.20 | 707.95 | 762.61 |
| ENSRNOG00000036273 | Mir363 | 1033.18 | 296.75 | 889.65 | 1249.59 | 1100.93 | 431.10 | 739.86 |
| ENSRNOG00000050728 | Mir190a-2 | 736.72 | 669.33 | 605.70 | 494.73 | 503.19 | 448.23 | 670.58 |
| ENSRNOG00000047497 | Mir190 | 672.70 | 683.71 | 612.65 | 527.39 | 535.76 | 494.38 | 656.35 |
| ENSRNOG00000035465 | Mir3065 | 653.98 | 632.73 | 668.20 | 377.13 | 310.95 | 266.30 | 651.64 |
| ENSRNOG00000035515 | Mir153 | 840.13 | 305.90 | 749.22 | 694.28 | 638.71 | 453.51 | 631.75 |
| ENSRNOG00000035547 | Mir33 | 678.61 | 634.03 | 543.97 | 488.20 | 518.95 | 541.84 | 618.87 |
| ENSRNOG00000035491 | Mir137 | 798.77 | 377.81 | 661.26 | 542.24 | 672.33 | 428.46 | 612.61 |
| ENSRNOG00000036268 | Mir362 | 688.46 | 439.25 | 695.21 | 481.07 | 316.20 | 160.84 | 607.64 |

| | | | | | | | | |
|---|---|---|---|---|---|---|---|---|
| ENSRNOG00000041554 | Mir1306 | 471.77 | 666.71 | 534.71 | 479.88 | 516.85 | 735.63 | 557.73 |
| ENSRNOG00000035561 | Mir34b | 551.55 | 594.81 | 468.36 | 374.76 | 564.12 | 437.69 | 538.24 |
| ENSRNOG00000035655 | Mir365b | 487.53 | 572.59 | 470.67 | 296.36 | 400.24 | 291.35 | 510.27 |
| ENSRNOG00000036260 | Mir497 | 510.19 | 543.83 | 430.55 | 553.53 | 528.41 | 344.09 | 494.86 |
| ENSRNOG00000047135 | Mir365-1 | 482.61 | 505.92 | 392.74 | 238.75 | 368.73 | 209.62 | 460.42 |
| ENSRNOG00000035549 | Mir92a2 | 458.97 | 232.70 | 602.62 | 779.81 | 709.09 | 433.73 | 431.43 |
| ENSRNOG00000035524 | Mir99a | 517.08 | 301.98 | 428.23 | 655.09 | 755.32 | 537.88 | 415.77 |
| ENSRNOG00000052819 | Mir346 | 464.88 | 398.72 | 334.10 | 295.17 | 262.63 | 228.07 | 399.23 |
| ENSRNOG00000035653 | Mir219a1 | 544.66 | 317.67 | 332.56 | 353.97 | 420.20 | 300.58 | 398.30 |
| ENSRNOG00000036558 | Mir20b | 500.34 | 162.10 | 490.73 | 772.09 | 669.17 | 325.63 | 384.39 |
| ENSRNOG00000046478 | Mir6324 | 436.32 | 367.35 | 335.64 | 367.04 | 386.59 | 283.44 | 379.77 |
| ENSRNOG00000040429 | Mir674 | 358.51 | 309.83 | 428.23 | 249.44 | 308.85 | 179.29 | 365.52 |
| ENSRNOG00000036262 | Mir500 | 356.54 | 284.99 | 378.85 | 291.02 | 220.61 | 143.70 | 340.13 |
| ENSRNOG00000046203 | Mir322-2 | 375.25 | 298.06 | 327.93 | 304.68 | 369.78 | 254.44 | 333.75 |
| ENSRNOG00000050456 | Mir351 | 356.54 | 292.83 | 348.76 | 339.72 | 307.80 | 284.76 | 332.71 |
| ENSRNOG00000046692 | Mir322 | 385.10 | 274.53 | 337.19 | 332.59 | 352.97 | 249.17 | 332.27 |
| ENSRNOG00000046106 | Mir351-2 | 349.64 | 292.83 | 341.82 | 326.65 | 316.20 | 267.62 | 328.10 |
| ENSRNOG00000043820 | Mir664-1 | 470.79 | 235.31 | 266.97 | 235.78 | 193.29 | 166.11 | 324.36 |
| ENSRNOG00000040395 | Mir484 | 326.01 | 354.27 | 253.08 | 245.29 | 370.83 | 164.79 | 311.12 |
| ENSRNOG00000047361 | Mir6329 | 307.29 | 313.75 | 295.52 | 201.34 | 226.91 | 369.13 | 305.52 |
| ENSRNOG00000052523 | rno-mir-484 | 303.35 | 333.36 | 259.26 | 251.22 | 354.02 | 171.38 | 298.66 |
| ENSRNOG00000060270 | Mir224 | 225.55 | 320.28 | 273.92 | 223.31 | 360.32 | 203.02 | 273.25 |
| ENSRNOG00000035641 | Mir138-1 | 256.08 | 291.52 | 239.19 | 207.87 | 146.02 | 122.61 | 262.27 |
| ENSRNOG00000048587 | Mir350-2 | 294.49 | 213.09 | 222.22 | 174.02 | 220.61 | 108.10 | 243.27 |
| ENSRNOG00000048754 | Mir378b | 250.17 | 244.46 | 230.71 | 156.20 | 195.39 | 180.61 | 241.78 |
| ENSRNOG00000047684 | Mir350 | 306.31 | 181.71 | 220.68 | 177.58 | 248.97 | 113.38 | 236.23 |
| ENSRNOG00000035508 | Mir200b | 309.26 | 171.25 | 222.22 | 176.99 | 185.94 | 197.75 | 234.25 |
| ENSRNOG00000041245 | Mir190b | 207.82 | 266.69 | 179.01 | 168.67 | 202.75 | 246.53 | 217.84 |
| ENSRNOG00000035620 | Mir22 | 229.49 | 164.72 | 217.59 | 212.62 | 197.50 | 100.19 | 203.93 |
| ENSRNOG00000035643 | Mir138-2 | 223.58 | 224.85 | 147.37 | 161.54 | 153.37 | 79.10 | 198.60 |
| ENSRNOG00000035471 | Mir124-1 | 264.94 | 135.96 | 187.50 | 148.48 | 111.35 | 55.37 | 196.13 |
| ENSRNOG00000035499 | Mir124-2 | 258.05 | 139.88 | 182.87 | 165.11 | 128.16 | 46.14 | 193.60 |
| ENSRNOG00000049100 | Mir344b-1 | 235.39 | 160.80 | 174.38 | 0.00 | 0.00 | 1.32 | 190.19 |
| ENSRNOG00000035593 | Mir124-3 | 254.11 | 107.20 | 195.21 | 150.26 | 147.07 | 50.10 | 185.51 |

| | | | | | | | | |
|---|---|---|---|---|---|---|---|---|
| ENSRNOG00000048126 | Mir503-2 | 228.50 | 137.26 | 188.27 | 179.36 | 171.23 | 122.61 | 184.68 |
| ENSRNOG00000048299 | Mir503 | 220.62 | 130.73 | 196.76 | 167.48 | 152.32 | 133.15 | 182.70 |
| ENSRNOG00000035506 | Mir326 | 183.19 | 201.32 | 155.09 | 131.25 | 133.41 | 69.87 | 179.87 |
| ENSRNOG00000035592 | Mir106a | 250.17 | 82.36 | 205.24 | 302.30 | 212.20 | 129.20 | 179.26 |
| ENSRNOG00000035621 | Mir135b | 94.55 | 346.43 | 81.02 | 22.57 | 40.97 | 52.73 | 174.00 |
| ENSRNOG00000035565 | Mir330 | 199.94 | 151.64 | 152.78 | 217.97 | 221.66 | 181.93 | 168.12 |
| ENSRNOG00000036443 | Mir499a | 221.61 | 166.03 | 86.42 | 149.07 | 135.52 | 56.69 | 158.02 |
| ENSRNOG00000035568 | Mir3074 | 124.10 | 193.48 | 148.15 | 149.07 | 132.36 | 125.24 | 155.24 |
| ENSRNOG00000036514 | Mir615 | 131.98 | 169.95 | 157.41 | 118.78 | 161.78 | 102.83 | 153.11 |
| ENSRNOG00000049846 | Mir344g | 152.66 | 118.96 | 145.06 | 0.59 | 0.00 | 0.00 | 138.89 |
| ENSRNOG00000050256 | Mir1843b | 129.02 | 120.27 | 104.17 | 109.87 | 87.19 | 75.15 | 117.82 |
| ENSRNOG00000049508 | Mir6318 | 131.98 | 126.81 | 93.36 | 112.25 | 102.95 | 97.56 | 117.38 |
| ENSRNOG00000046738 | Mir542-3 | 130.99 | 78.44 | 113.42 | 71.27 | 90.34 | 79.10 | 107.62 |
| ENSRNOG00000049278 | Mir542 | 132.96 | 75.82 | 106.48 | 78.40 | 86.14 | 90.97 | 105.09 |
| ENSRNOG00000046539 | Mir542-2 | 125.08 | 84.97 | 97.99 | 73.05 | 88.24 | 92.28 | 102.68 |

#### Table S5. Highly expressed snoRNAs in INS-1 lines from small RNA-seq.

| gene_ID | gene_name | Control 5-9 | Control 2 | Control 16 | PWS 3 | PWS 19-1 | PWS 19-4 | Control_AVG |
|---|---|---|---|---|---|---|---|---|
| ENSRNOG00000044894 | Scarna15 | 21059.46 | 14308.22 | 18644.03 | 16274.39 | 14670.36 | 14104.89 | 18003.91 |
| ENSRNOG00000054739 | Snord110 | 14760.93 | 16790.76 | 17104.70 | 13993.77 | 12294.11 | 21177.77 | 16218.79 |
| ENSRNOG00000056237 | Snord57 | 16744.55 | 14637.66 | 15327.72 | 14892.36 | 12641.83 | 15707.99 | 15569.98 |
| ENSRNOG00000052611 | Snord43 AC127784.1 | 13443.11 | 13929.11 | 14172.64 | 13738.38 | 12049.34 | 19657.73 | 13848.29 |
| ENSRNOG00000052013 | Snord81 | 12638.43 | 11700.19 | 13933.45 | 13031.03 | 11901.22 | 13648.75 | 12757.36 |
| ENSRNOG00000061138 | Snord30 | 11203.41 | 10707.96 | 10996.76 | 11003.42 | 9490.30 | 11205.87 | 10969.38 |
| ENSRNOG00000057682 | snoZ40 Gm24357 | 10008.71 | 11840.07 | 10834.73 | 10127.40 | 9656.28 | 13619.74 | 10894.50 |
| ENSRNOG00000053795 | Snord45 Gm24494 | 10036.29 | 9677.82 | 9651.87 | 10367.34 | 9475.59 | 13995.47 | 9788.66 |
| ENSRNOG00000056420 | Snord45c | 9623.61 | 10229.50 | 9452.80 | 10615.00 | 9969.33 | 14542.58 | 9768.64 |
| ENSRNOG00000057925 | Snord91a | 8668.24 | 8758.80 | 8545.41 | 8240.54 | 6228.47 | 9420.84 | 8657.48 |
| ENSRNOG00000006961 | Snrpb (Snord119) | 8908.56 | 7754.81 | 9212.84 | 8728.73 | 7299.99 | 13632.93 | 8625.40 |
| ENSRNOG00000060919 | Snord72 | 7887.20 | 8850.31 | 8340.16 | 8099.18 | 7199.14 | 15842.46 | 8359.23 |
| ENSRNOG00000053988 | Snord52 | 5365.82 | 7999.27 | 6506.08 | 6242.02 | 5968.99 | 8948.87 | 6623.72 |
| ENSRNOG00000051436 | Snord24 | 4337.57 | 7799.26 | 6825.52 | 4887.90 | 5648.59 | 9887.53 | 6320.78 |
| ENSRNOG00000058679 | Snord21 | 5805.09 | 5344.18 | 6033.10 | 5880.33 | 4570.77 | 5674.12 | 5727.45 |
| ENSRNOG00000060818 | Snord37 | 4560.16 | 5859.25 | 5070.15 | 4772.68 | 4411.09 | 8880.32 | 5163.18 |
| ENSRNOG00000060290 | Snord87 | 5285.06 | 5162.46 | 4616.45 | 4416.33 | 3721.95 | 3823.18 | 5021.32 |
| ENSRNOG00000056422 | Snord104 | 4030.27 | 5172.92 | 4786.20 | 4260.13 | 4223.05 | 5794.09 | 4663.13 |
| ENSRNOG00000057993 | Snord38 AC119459.2 | 3152.71 | 4865.71 | 3513.07 | 3458.35 | 3282.84 | 4601.00 | 3843.83 |
| ENSRNOG00000053621 | Snord2 | 3346.74 | 3623.79 | 3626.49 | 3548.03 | 3336.42 | 4330.74 | 3532.34 |
| ENSRNOG00000057012 | Snord95 | 2663.21 | 4485.29 | 3425.88 | 2648.26 | 2868.94 | 6326.70 | 3524.79 |
| ENSRNOG00000052660 | Snord95 Gm25296 | 2646.47 | 4438.23 | 3322.48 | 2711.21 | 2985.55 | 6454.58 | 3469.06 |
| ENSRNOG00000061471 | Snord48 Gm25744 | 3077.86 | 4002.90 | 3000.73 | 3622.87 | 3558.07 | 6486.22 | 3360.50 |
| ENSRNOG00000054315 | Snord27 | 2631.69 | 3414.63 | 3104.90 | 3409.06 | 3114.76 | 3033.49 | 3050.41 |
| ENSRNOG00000055952 | Snord75 AC113837.4 | 2692.76 | 3168.86 | 2940.55 | 2720.12 | 2425.63 | 3245.75 | 2934.05 |
| ENSRNOG00000060216 | Snord20 Gm24148 | 3025.66 | 2810.66 | 2572.50 | 2990.35 | 2712.41 | 3378.90 | 2802.94 |
| ENSRNOG00000059148 | Snord25 | 2814.89 | 2492.99 | 2775.42 | 2944.62 | 2405.67 | 3530.51 | 2694.43 |
| ENSRNOG00000054872 | snoR38 AC123144.4 | 2436.68 | 2247.22 | 2452.90 | 2661.32 | 2111.52 | 3096.77 | 2378.93 |
| ENSRNOG00000053586 | Snord19 | 2289.93 | 2265.52 | 2421.26 | 2382.78 | 2021.18 | 2607.67 | 2325.57 |
| ENSRNOG00000058269 | Snord83b | 2240.68 | 2265.52 | 2467.56 | 2412.47 | 2819.57 | 4568.04 | 2324.59 |
| ENSRNOG00000056942 | Snord31 | 2334.25 | 1916.48 | 2230.68 | 2506.90 | 2157.75 | 2329.50 | 2160.47 |
| ENSRNOG00000047709 | Snord93 AABR07059190.1 | 2212.12 | 1822.35 | 2064.01 | 2124.43 | 1866.75 | 1985.42 | 2032.83 |
| ENSRNOG00000056913 | Snord91 Gm22771 | 1706.86 | 2150.48 | 1856.46 | 2033.56 | 1825.79 | 2096.16 | 1904.60 |
| ENSRNOG00000056002 | Snord50 AABR07071004.1 | 1864.44 | 1856.34 | 1844.88 | 1619.60 | 1672.41 | 1352.61 | 1855.22 |
| ENSRNOG00000058779 | Snord50 AABR07071007.1 | 1899.90 | 1811.90 | 1802.44 | 1602.97 | 1604.13 | 1367.12 | 1838.08 |
| ENSRNOG00000060079 | Snord26 | 1733.45 | 1601.42 | 2096.42 | 2294.88 | 1970.76 | 3070.41 | 1810.43 |
| ENSRNOG00000054737 | Snord50 AABR07071006.1 | 1835.88 | 1813.20 | 1753.06 | 1565.55 | 1656.65 | 1248.47 | 1800.72 |
| ENSRNOG00000051978 | Snord1b | 1838.84 | 1478.54 | 1907.38 | 2013.36 | 1585.22 | 1075.76 | 1741.59 |
| ENSRNOG00000056119 | Snord66 | 1383.81 | 1809.28 | 1920.50 | 1681.37 | 1678.71 | 2759.28 | 1704.53 |
| ENSRNOG00000054347 | Snord74 Gm26224 | 1785.65 | 1636.72 | 1516.18 | 1872.01 | 1858.35 | 1524.00 | 1646.18 |
| ENSRNOG00000061827 | Snord50 AC120938.1 | 1642.84 | 1502.07 | 1587.17 | 1433.70 | 1463.36 | 1110.04 | 1577.36 |
| ENSRNOG00000059872 | Snora58 Gm50450 | 1562.08 | 1338.66 | 1560.16 | 1585.75 | 1391.92 | 1367.12 | 1486.97 |
| ENSRNOG00000035409 | Snora58 | 1535.48 | 1321.66 | 1532.39 | 1557.24 | 1374.07 | 1352.61 | 1463.18 |
| ENSRNOG00000052293 | Snord15 | 1324.71 | 1211.85 | 1775.44 | 1877.36 | 1515.88 | 1568.82 | 1437.33 |
| ENSRNOG00000054324 | Snord39 | 1056.81 | 1813.20 | 1322.51 | 1269.78 | 1299.48 | 1357.89 | 1397.51 |
| ENSRNOG00000056475 | Snord41 Gm25506 | 1219.33 | 1455.01 | 1266.96 | 1344.02 | 1158.71 | 2808.06 | 1313.76 |
| ENSRNOG00000057809 | Snord36 | 1146.44 | 1485.07 | 1305.54 | 962.14 | 1264.81 | 1081.04 | 1312.35 |
| ENSRNOG00000060311 | Snord34 | 1048.93 | 1626.26 | 1218.35 | 1133.19 | 1262.71 | 1692.75 | 1297.85 |
| ENSRNOG00000054079 | Snord36 | 1087.35 | 1486.38 | 1294.74 | 938.38 | 1237.50 | 1003.25 | 1289.49 |
| ENSRNOG00000051927 | Snord98 | 1346.38 | 1162.18 | 1314.80 | 1187.23 | 1065.22 | 1342.07 | 1274.45 |
| ENSRNOG00000051481 | Scarna18 AABR07032532.1 | 1108.03 | 1302.05 | 1346.43 | 1256.72 | 1265.86 | 2322.91 | 1252.17 |
| ENSRNOG00000057240 | Snord49b | 1371.00 | 1192.24 | 1175.91 | 1253.16 | 1053.66 | 636.76 | 1246.38 |
| ENSRNOG00000053764 | Snord33 | 1122.80 | 1256.30 | 1334.09 | 1276.32 | 1149.26 | 1103.45 | 1237.73 |
| ENSRNOG00000055658 | Snord69 | 1225.23 | 1153.02 | 1273.13 | 1278.10 | 901.34 | 1032.26 | 1217.13 |
| ENSRNOG00000055464 | Snord82 | 970.14 | 1054.98 | 1081.00 | 1219.30 | 1018.99 | 1026.98 | 1035.37 |
| ENSRNOG00000057905 | Snord60 | 1173.03 | 924.25 | 920.51 | 1139.72 | 858.27 | 660.49 | 1005.93 |
| ENSRNOG00000059556 | Snora36b | 1042.04 | 878.49 | 922.06 | 851.67 | 840.41 | 912.29 | 947.53 |
| ENSRNOG00000055378 | Snord38a | 668.76 | 1221.00 | 797.06 | 923.53 | 1016.89 | 971.61 | 895.61 |
| ENSRNOG00000055722 | Snord42 | 832.25 | 841.89 | 985.33 | 971.05 | 1010.59 | 746.18 | 886.49 |

| | | | | | | | | |
|---|---|---|---|---|---|---|---|---|
| ENSRNOG00000051367 | Snord70 Gm26293 | 832.25 | 996.15 | 826.38 | 858.20 | 783.68 | 1537.18 | 884.93 |
| ENSRNOG00000060388 | Snord45 AABR07013860.4 | 916.96 | 800.06 | 844.90 | 1015.59 | 817.30 | 791.00 | 853.97 |
| ENSRNOG00000060532 | Snord105 Gm24067 | 683.53 | 1041.91 | 818.66 | 854.64 | 770.02 | 841.10 | 848.03 |
| ENSRNOG00000058132 | Snord29 | 760.35 | 941.24 | 827.92 | 889.68 | 784.73 | 779.14 | 843.17 |
| ENSRNOG00000061361 | Snord46 Gm26330 | 780.05 | 452.32 | 1293.96 | 1321.46 | 570.43 | 773.86 | 842.11 |
| ENSRNOG00000054607 | Snord56 Gm23650 | 608.68 | 900.72 | 894.28 | 755.46 | 849.86 | 1095.54 | 801.22 |
| ENSRNOG00000055561 | Snord58 | 772.17 | 725.54 | 799.37 | 928.29 | 794.18 | 750.13 | 765.70 |
| ENSRNOG00000059435 | Snord18 | 998.70 | 661.49 | 616.50 | 685.97 | 637.66 | 1144.32 | 758.90 |
| ENSRNOG00000056657 | Snord62 | 613.60 | 762.15 | 851.07 | 808.91 | 764.77 | 843.74 | 742.27 |
| ENSRNOG00000053438 | Snora7a | 967.19 | 559.52 | 668.20 | 660.43 | 516.85 | 510.20 | 731.64 |
| ENSRNOG00000056323 | Snora7 AC118412.1 | 965.22 | 571.28 | 639.65 | 693.10 | 518.95 | 503.60 | 725.38 |
| ENSRNOG00000052983 | Snord18 | 522.01 | 977.85 | 637.34 | 621.23 | 827.80 | 2179.21 | 712.40 |
| ENSRNOG00000053819 | Snord88 AABR07003250.2 | 467.83 | 917.71 | 646.60 | 654.49 | 743.76 | 1319.66 | 677.38 |
| ENSRNOG00000052573 | Snord17 | 707.17 | 468.01 | 840.27 | 950.85 | 609.30 | 416.59 | 671.81 |
| ENSRNOG00000057120 | Snora7 AC129753.3 | 885.44 | 492.85 | 611.10 | 579.66 | 476.93 | 481.19 | 663.13 |
| ENSRNOG00000055569 | Snord15 | 635.27 | 461.47 | 817.89 | 910.47 | 628.20 | 611.71 | 638.21 |
| ENSRNOG00000059940 | Snord116 | 765.28 | 436.63 | 671.29 | 0.00 | 0.00 | 0.00 | 624.40 |
| ENSRNOG00000060868 | Scarna3b | 666.79 | 626.19 | 571.75 | 422.87 | 448.57 | 363.86 | 621.58 |
| ENSRNOG00000045015 | Scarna3 | 662.85 | 617.04 | 565.58 | 418.11 | 438.06 | 362.54 | 615.15 |
| ENSRNOG00000059171 | Snord58 | 596.86 | 643.18 | 578.70 | 688.94 | 597.74 | 688.17 | 606.25 |
| ENSRNOG00000051811 | Snord116 | 708.15 | 415.72 | 668.20 | 0.00 | 0.00 | 0.00 | 597.36 |
| ENSRNOG00000059926 | Snord71 | 604.74 | 539.91 | 638.11 | 566.00 | 467.48 | 445.60 | 594.25 |
| ENSRNOG00000058825 | Snord116 | 687.47 | 420.95 | 660.48 | 0.59 | 0.00 | 0.00 | 589.63 |
| ENSRNOG00000057316 | Snord88 Gm26247 | 362.45 | 644.49 | 747.67 | 530.96 | 638.71 | 652.58 | 584.87 |
| ENSRNOG00000052577 | Snora54 Gm23297 | 625.42 | 485.00 | 594.90 | 462.66 | 434.91 | 412.64 | 568.44 |
| ENSRNOG00000054305 | Snord116 | 635.27 | 383.03 | 644.28 | 0.00 | 0.00 | 1.32 | 554.20 |
| ENSRNOG00000055390 | Snord10 Gm25835 | 565.34 | 423.56 | 672.83 | 612.92 | 440.16 | 396.82 | 553.91 |
| ENSRNOG00000052156 | Snord51 Gm26457 | 607.69 | 529.45 | 514.65 | 574.31 | 450.67 | 705.31 | 550.60 |
| ENSRNOG00000058899 | Snord49a | 516.10 | 522.91 | 601.07 | 596.88 | 565.17 | 309.81 | 546.69 |
| ENSRNOG00000061924 | Snord64 AABR07005711.2 | 560.42 | 486.31 | 578.70 | 0.00 | 0.00 | 0.00 | 541.81 |
| ENSRNOG00000055348 | Snord64 AABR07005710.1 | 542.69 | 466.70 | 608.79 | 0.00 | 0.00 | 0.00 | 539.39 |
| ENSRNOG00000058488 | Snord92 | 517.08 | 477.16 | 619.59 | 652.71 | 501.09 | 991.39 | 537.94 |
| ENSRNOG00000040595 | Snord79 | 531.85 | 505.92 | 564.04 | 777.43 | 761.62 | 905.70 | 533.94 |
| ENSRNOG00000059478 | snoU54 Gm24016 | 475.71 | 602.66 | 420.52 | 386.64 | 378.18 | 353.31 | 499.63 |
| ENSRNOG00000060627 | Snord62 | 439.27 | 465.39 | 565.58 | 525.61 | 468.53 | 544.47 | 490.08 |
| ENSRNOG00000056397 | Snord22 | 477.68 | 533.37 | 415.12 | 421.68 | 384.49 | 332.22 | 475.39 |
| ENSRNOG00000060589 | Snord53_Snord92 Gm22858 | 404.80 | 541.22 | 416.66 | 408.02 | 366.63 | 297.94 | 454.23 |
| ENSRNOG00000059509 | snosnR60_Z15 AC113837.7 | 430.41 | 564.75 | 367.28 | 512.55 | 521.05 | 457.46 | 454.14 |
| ENSRNOG00000060075 | Snord77 AC113837.8 | 430.41 | 564.75 | 367.28 | 512.55 | 521.05 | 457.46 | 454.14 |
| ENSRNOG00000053606 | Scarna6 | 497.38 | 414.41 | 448.30 | 453.16 | 414.95 | 448.23 | 453.36 |
| ENSRNOG00000052356 | snoZ196 | 475.71 | 427.48 | 420.52 | 465.63 | 352.97 | 661.81 | 441.24 |
| ENSRNOG00000059804 | Snord102 Gm25091 | 430.41 | 376.50 | 475.30 | 554.12 | 405.50 | 611.71 | 427.40 |
| ENSRNOG00000057398 | Snord19B Gm24916 | 294.49 | 453.63 | 453.70 | 379.51 | 482.18 | 697.40 | 400.60 |
| ENSRNOG00000058148 | Snord58 | 455.03 | 363.42 | 363.42 | 452.56 | 388.69 | 267.62 | 393.96 |
| ENSRNOG00000060842 | Snord12 AC130053.4 | 462.91 | 338.59 | 359.56 | 411.58 | 348.77 | 197.75 | 387.02 |
| ENSRNOG00000055560 | Snord33 | 302.37 | 456.24 | 402.00 | 410.99 | 461.17 | 413.96 | 386.87 |
| ENSRNOG00000054081 | Snord12 | 334.87 | 430.10 | 384.25 | 361.69 | 305.70 | 362.54 | 383.07 |
| ENSRNOG00000055151 | Snord34 | 235.39 | 475.85 | 358.79 | 267.85 | 347.72 | 357.27 | 356.68 |
| ENSRNOG00000051750 | Snord116 | 417.60 | 290.22 | 342.59 | 0.00 | 1.05 | 0.00 | 350.14 |
| ENSRNOG00000054970 | Snord53 | 355.55 | 339.89 | 339.50 | 348.63 | 308.85 | 440.32 | 344.98 |
| ENSRNOG00000061767 | Snord83 | 238.35 | 454.93 | 330.24 | 329.62 | 362.43 | 487.78 | 341.18 |
| ENSRNOG00000058017 | Snord90 | 349.64 | 277.14 | 385.80 | 447.22 | 262.63 | 406.05 | 337.53 |
| ENSRNOG00000051822 | Snord116 | 354.57 | 241.85 | 413.57 | 0.59 | 0.00 | 0.00 | 336.66 |
| ENSRNOG00000055429 | Snord4a | 369.34 | 311.13 | 304.01 | 345.06 | 267.88 | 152.93 | 328.16 |
| ENSRNOG00000054381 | Snora36 AC136161.1 | 470.79 | 235.31 | 266.97 | 235.78 | 193.29 | 166.11 | 324.36 |
| ENSRNOG00000060158 | Snord116 | 366.39 | 217.01 | 382.71 | 0.00 | 0.00 | 1.32 | 322.04 |
| ENSRNOG00000057952 | Snord101 Gm23130 | 353.58 | 318.98 | 282.40 | 355.16 | 304.65 | 292.67 | 318.32 |
| ENSRNOG00000054310 | snoMBII-202 AC119635.1 | 295.47 | 349.04 | 291.66 | 303.49 | 308.85 | 341.45 | 312.06 |
| ENSRNOG00000058660 | Snora79 Gm25848 | 298.43 | 269.30 | 344.13 | 353.38 | 310.95 | 535.24 | 303.95 |
| ENSRNOG00000061909 | Snord116 | 340.78 | 234.00 | 330.24 | 0.00 | 0.00 | 1.32 | 301.68 |

| | | | | | | | | |
|---|---|---|---|---|---|---|---|---|
| ENSRNOG00000057278 | Snord47 AC113837.6 | 346.69 | 313.75 | 242.28 | 354.57 | 310.95 | 218.84 | 300.91 |
| ENSRNOG00000053918 | Snord116 | 340.78 | 251.00 | 307.09 | 0.00 | 0.00 | 0.00 | 299.62 |
| ENSRNOG00000055242 | Snora7 AABR07028813.1 | 404.80 | 198.71 | 293.98 | 262.51 | 221.66 | 233.35 | 299.16 |
| ENSRNOG00000054326 | Snord88 AABR07003250.3 | 280.70 | 312.44 | 293.21 | 283.89 | 272.08 | 485.15 | 295.45 |
| ENSRNOG00000061669 | Snord116 | 334.87 | 198.71 | 349.53 | 0.00 | 0.00 | 0.00 | 294.37 |
| ENSRNOG00000055465 | Snora62 | 242.29 | 281.07 | 353.39 | 346.84 | 289.94 | 424.50 | 292.25 |
| ENSRNOG00000052552 | Snord116 | 344.72 | 192.17 | 335.64 | 0.00 | 0.00 | 0.00 | 290.84 |
| ENSRNOG00000058093 | Snord59a | 228.50 | 355.58 | 248.45 | 288.64 | 266.83 | 274.21 | 277.51 |
| ENSRNOG00000057313 | Snord22 | 300.40 | 253.61 | 262.34 | 219.15 | 217.46 | 213.57 | 272.12 |
| ENSRNOG00000061678 | Snord100 | 264.94 | 243.15 | 285.49 | 284.48 | 303.60 | 208.30 | 264.53 |
| ENSRNOG00000057249 | Snord59 Gm50449 | 267.90 | 295.45 | 222.22 | 305.27 | 237.42 | 352.00 | 261.85 |
| ENSRNOG00000058211 | Snord22 | 278.73 | 248.38 | 250.00 | 212.62 | 201.70 | 205.66 | 259.04 |
| ENSRNOG00000052740 | Snord116 | 309.26 | 183.02 | 266.97 | 0.00 | 0.00 | 0.00 | 253.08 |
| ENSRNOG00000055652 | Snord44 Gm23212 | 332.90 | 172.56 | 231.48 | 242.91 | 181.74 | 175.34 | 245.65 |
| ENSRNOG00000055753 | Snord116 | 265.93 | 186.94 | 240.74 | 0.00 | 0.00 | 0.00 | 231.20 |
| ENSRNOG00000052487 | Snord73 | 198.95 | 290.22 | 167.44 | 247.07 | 265.78 | 300.58 | 218.87 |
| ENSRNOG00000053089 | Snora3 AABR07005004.1 | 196.00 | 248.38 | 212.19 | 243.50 | 219.56 | 254.44 | 218.86 |
| ENSRNOG00000058701 | Snord107 | 203.88 | 172.56 | 274.69 | 0.59 | 0.00 | 1.32 | 217.04 |
| ENSRNOG00000057770 | Snord111 Gm22193 | 240.32 | 185.63 | 219.90 | 270.82 | 224.81 | 230.71 | 215.29 |
| ENSRNOG00000047430 | Snord93 AABR07059193.1 | 207.82 | 224.85 | 208.33 | 234.00 | 262.63 | 266.30 | 213.67 |
| ENSRNOG00000058816 | Snord73 | 206.83 | 205.24 | 223.76 | 241.13 | 191.19 | 188.52 | 211.95 |
| ENSRNOG00000060559 | Scarna10 | 232.44 | 142.49 | 258.48 | 310.62 | 196.45 | 263.67 | 211.14 |
| ENSRNOG00000060924 | Snora3 AABR07005004.4 | 189.10 | 239.23 | 202.93 | 236.97 | 221.66 | 251.80 | 210.42 |
| ENSRNOG00000052187 | Snord116 | 229.49 | 197.40 | 201.39 | 0.00 | 0.00 | 0.00 | 209.42 |
| ENSRNOG00000060031 | Snord35a | 126.07 | 277.14 | 222.22 | 212.62 | 277.33 | 346.72 | 208.48 |
| ENSRNOG00000060695 | Snord88a | 174.33 | 243.15 | 199.07 | 217.97 | 195.39 | 424.50 | 205.52 |
| ENSRNOG00000053521 | Snord116 | 261.99 | 168.64 | 183.64 | 0.00 | 0.00 | 0.00 | 204.76 |
| ENSRNOG00000055949 | snoU105B Gm23008 | 223.58 | 179.10 | 194.44 | 201.93 | 210.10 | 171.38 | 199.04 |
| ENSRNOG00000053973 | Snord121A Gm24837 | 137.89 | 265.38 | 187.50 | 204.90 | 196.45 | 280.81 | 196.92 |
| ENSRNOG00000059154 | Snord116 | 246.23 | 149.03 | 185.95 | 0.00 | 0.00 | 0.00 | 193.74 |
| ENSRNOG00000054389 | Snord116 | 229.49 | 151.64 | 182.87 | 0.59 | 0.00 | 0.00 | 188.00 |
| ENSRNOG00000053674 | snoU18 Gm22571 | 242.29 | 130.73 | 182.10 | 224.50 | 153.37 | 155.56 | 185.04 |
| ENSRNOG00000055361 | Snord18 | 242.29 | 130.73 | 182.10 | 224.50 | 153.37 | 155.56 | 185.04 |
| ENSRNOG00000058606 | Snora1 Gm22620 | 154.63 | 181.71 | 211.42 | 200.74 | 190.14 | 216.21 | 182.59 |
| ENSRNOG00000057393 | Snord116 | 205.85 | 129.42 | 201.39 | 0.00 | 0.00 | 0.00 | 178.88 |
| ENSRNOG00000061656 | Snord14 | 186.15 | 147.72 | 202.16 | 231.63 | 133.41 | 106.79 | 178.68 |
| ENSRNOG00000060649 | Snord24 | 142.81 | 197.40 | 192.90 | 153.82 | 191.19 | 110.74 | 177.70 |
| ENSRNOG00000055640 | Snord65 | 174.33 | 163.41 | 187.50 | 219.75 | 210.10 | 221.48 | 175.08 |
| ENSRNOG00000061024 | Snord121A Gm22888 | 117.20 | 213.09 | 189.81 | 162.73 | 147.07 | 212.25 | 173.37 |
| ENSRNOG00000053443 | SNORD36 | 148.72 | 205.24 | 147.37 | 136.01 | 151.27 | 152.93 | 167.11 |
| ENSRNOG00000051850 | Scarna2 | 166.45 | 88.90 | 244.60 | 248.85 | 131.31 | 148.97 | 166.65 |
| ENSRNOG00000059404 | Snora64 | 207.82 | 155.57 | 133.49 | 174.61 | 182.79 | 98.88 | 165.62 |
| ENSRNOG00000057883 | Snord36 | 149.71 | 177.79 | 162.81 | 158.57 | 155.48 | 148.97 | 163.43 |
| ENSRNOG00000053898 | Snora24 AABR07013208.1 | 207.82 | 137.26 | 133.49 | 200.74 | 160.73 | 114.70 | 159.52 |
| ENSRNOG00000052161 | Snord5 AC105648.4 | 184.18 | 124.19 | 158.95 | 199.55 | 96.65 | 143.70 | 155.77 |
| ENSRNOG00000058848 | Snord116 | 165.47 | 100.66 | 176.70 | 0.00 | 1.05 | 0.00 | 147.61 |
| ENSRNOG00000056200 | Snord12 Gm25878 | 124.10 | 147.72 | 165.89 | 162.73 | 128.16 | 105.47 | 145.91 |
| ENSRNOG00000054774 | Snord14 | 143.80 | 104.58 | 185.18 | 175.80 | 112.40 | 69.87 | 144.52 |
| ENSRNOG00000060578 | Snord70 | 157.59 | 147.72 | 124.23 | 186.49 | 133.41 | 105.47 | 143.18 |
| ENSRNOG00000054559 | Snord61 | 208.80 | 86.28 | 133.49 | 184.11 | 122.91 | 112.06 | 142.86 |
| ENSRNOG00000057500 | Snora79 Gm25776 | 172.36 | 99.35 | 138.12 | 127.10 | 118.71 | 238.62 | 136.61 |
| ENSRNOG00000056157 | Snord116 | 161.53 | 91.51 | 155.86 | 0.00 | 0.00 | 0.00 | 136.30 |
| ENSRNOG00000051605 | Snord86 Gm23187 | 147.74 | 116.35 | 126.54 | 129.47 | 151.27 | 187.20 | 130.21 |
| ENSRNOG00000052114 | Snord29 | 128.04 | 156.87 | 104.17 | 116.41 | 99.80 | 59.33 | 129.69 |
| ENSRNOG00000060156 | Snord28 | 135.92 | 111.12 | 136.57 | 149.07 | 136.57 | 208.30 | 127.87 |
| ENSRNOG00000061095 | ScaRNA20 Gm25492 | 124.10 | 122.88 | 119.60 | 109.87 | 112.40 | 154.25 | 122.19 |
| ENSRNOG00000053634 | Scarna3a | 130.99 | 128.11 | 107.25 | 114.03 | 88.24 | 80.42 | 122.12 |
| ENSRNOG00000056713 | Snord116 | 150.69 | 103.28 | 111.11 | 0.00 | 0.00 | 0.00 | 121.69 |
| ENSRNOG00000057005 | Snord116 | 143.80 | 91.51 | 128.08 | 0.00 | 0.00 | 0.00 | 121.13 |
| ENSRNOG00000054329 | Snord16 | 130.99 | 98.05 | 131.17 | 168.67 | 96.65 | 145.02 | 120.07 |

| | | | | | | | | |
|---|---|---|---|---|---|---|---|---|
| ENSRNOG00000053726 | Snord116 | 151.68 | 94.12 | 109.57 | 0.00 | 0.00 | 0.00 | 118.46 |
| ENSRNOG00000053927 | ScaRNA7 Gm22009 | 122.13 | 111.12 | 121.14 | 147.88 | 147.07 | 90.97 | 118.13 |
| ENSRNOG00000060502 | Snord96 | 113.27 | 99.35 | 140.43 | 148.48 | 142.87 | 208.30 | 117.68 |
| ENSRNOG00000060242 | Snord103 AABR07050041.2 | 112.28 | 150.34 | 83.33 | 105.12 | 93.50 | 84.37 | 115.32 |
| ENSRNOG00000055168 | Snora79 AABR07051827.1 | 116.22 | 101.97 | 121.14 | 124.13 | 89.29 | 214.89 | 113.11 |
| ENSRNOG00000053379 | Snord116 | 144.78 | 90.20 | 102.62 | 0.00 | 0.00 | 0.00 | 112.54 |
| ENSRNOG00000059567 | Snord116 | 125.08 | 87.59 | 124.23 | 0.00 | 0.00 | 0.00 | 112.30 |
| ENSRNOG00000056644 | Snord116 | 140.84 | 87.59 | 108.02 | 0.00 | 0.00 | 0.00 | 112.15 |
| ENSRNOG00000055929 | Snord103 AABR07050041.1 | 102.43 | 146.42 | 81.79 | 108.09 | 88.24 | 85.69 | 110.21 |
| ENSRNOG00000053003 | Snord11 | 89.63 | 118.96 | 115.74 | 155.61 | 123.96 | 156.88 | 108.11 |
| ENSRNOG00000055027 | Snord83 | 91.60 | 103.28 | 118.05 | 121.16 | 116.61 | 135.79 | 104.31 |

**Table S6. sgRNA oligonucleotides.** *Bbs* I site oligonucleotide cloning adapters for sgRNAs. Abbreviations: b/w, between; F, forward.

| sgRNA Oligo and Location | Oligo # | OligoSequence |
|---|---|---|
| sgRNA1 5’ of *Frat 3* sense | 4602 | 5’-CACCgGTCAGTGTGTTATTGCTCTG-3’ |
| sgRNA1 5’ of *Frat3* complementary | 4603 | 5’-AAACCAGAGCAATAACACACTGACc-3’ |
| sgRNA2 *Frat3* 3’-UTR sense | 4604 | 5’-CACCgAAAGACCATCTCATCTTTTG-3’ |
| sgRNA2 *Frat3* 3’-UTR complementary | 4605 | 5’-AAACCAAAAGATGAGATGGTCTTTc-3’ |
| sgRNA3 b/w *Snord115* /*Ube3a* #1 sense | 4618 | 5’-CACCgTACTTATGACAATTAACCTC-3’ |
| sgRNA3 b/w *Snord115* /*Ube3a* #1 complementary | 4619 | 5’-AAACGAGGTTAATTGTCATAAGTAc-3’ |
| sgRNA4 b/w *Snord115* /*Ube3a* #2 sense | 4620 | 5’-CACCgGCCAGTGACATGGTTTAGCA-3’ |
| sgRNA4 b/w *Snord115* /*Ube3a* #2 complementary | 4621 | 5’-AAACTGCTAAACCATGTCACTGGCc-3’ |
| sgRNA64-4 5' of *Frat3* #1 sense (3' of gRNA1) | 4990 | 5’-CACCgTTCAACATATTGGCCACCAG-3’ |
| sgRNA64-4 5' of *Frat3* #1 complementary | 4991 | 5’-AAACCTGGTGGCCAATATGTTGAAc-3’ |
| sgRNA70-3 5' of *Frat3* #2 (3' of gRNA1) | 4992 | 5’-CACCgGCTTGTATGAATCCCTCTGG-3’ |
| sgRNA70-3 5' of *Frat3* #2 complementary | 4993 | 5’-AAACCCAGAGGGATTCATACAAGCc-3’ |
| sgRNA79-1 inside b/w *Snord115* /*Ube3a* #1 sense | 4994 | 5’-CACCgGCCAGGTACAGGTAGTTCAC-3’ |
| sgRNA79-1 inside b/w *Snord115* /*Ube3a* #1 complementary | 4995 | 5’-AAACGTGAACTACCTGTACCTGGCc-3’ |
| sgRNA58-7 inside b/w *Snord115* /*Ube3a* #2 | 4996 | 5’-CACCgTGTTCAAACCAGGGGCACTG-3’ |
| sgRNA58-7 inside b/w *Snord115* /*Ube3a* #2 complementary | 4997 | 5’-AAACCAGTGCCCCTGGTTTGAACAc-3’ |
| pX330 U6 promoter F sequencing primer | 4454 | 5’-GAGGGCCTATTTCCCATGATTCC-3’ |

**Table S7. Genomic PCR, Copy number ddPCR, and DNA methylation PCR primers.** Abbreviations: F, forward; R, reverse; un, unmethylated; me, methylated; IC, imprinting cente

| Genomic PCR Primer | Primer # | Primer Sequence | Use |
|---|---|---|---|
| sgRNA1 deletion-PCR F | 4642 | 5′-TGGGAACTACAGTTCACACCTC-3′ | Genome editing |
| sgRNA3 deletion-PCR R | 4644 | 5′-TTGTCACCCTGGTAAGATACTG-3′ | Genome editing |
| sgRNA1 rescreen deletion-PCR F | 4805 | 5′-AAGTTACAGCAAATCTTTATTCTG-3′ | Genome editing |
| sgRNA3 rescreen deletion-PCR R | 4806 | 5′-AACACGAAAATATGTAACCACAG-3′ | Genome editing |
| sgRNA1 scarred allele PCR R | 4988 | 5′-TATAATTCAGCTTTGTGAATAGTATC-3′ | Genome editing |
| sgRNA1 scarred allele PCR R | 5353 | 5′-ATTAGAGTCCTAACTTCAATCAATC-3′ | Genome editing |
| sgRNA3 scarred allele PCR F | 4989 | 5′-TACTACAGATGTAAGGGCCAAG-3′ | Genome editing |
| sgRNA2 deletion-PCR F | 4643 | 5′-GTTGGTTCATGACTCACTTTGG-3′ | Genome editing |
| sgRNA4 deletion-PCR R | 4645 | 5′-AAAACCAGTGTGCTTCTGTCAC-3′ | Genome editing |
| sgRNA70-3, sgRNA64-4 deletion-PCR F | 4998 | 5′-GAGCTTGAAGTCAGATTACACC-3′ | Genome editing |
| sgRNA79-1 deletion-PCR R | 4999 | 5′-TGAAGTTGGCACCTTTGTTTTGC-3′ | Genome editing |
| alt-sgRNA70-3 deletion-PCR F | 5193 | 5′-TTCACCCTAGCAGATAAAGCTG-3′ | Genome editing |
| sgRNA58-7 deletion-PCR R | 5001 | 5′CTTGTCAGTGAACTTCCATCAC--3′ | Genome editing |
| sgRNA70-3 scarred allele PCR R | 5002 | 5′-ATACAGAATGTCCTGGCTTATAC-3′ | Genome editing |
| sgRNA79-1 scarred allele PCR F | 5003 | 5′-GTACAATGTGATGATTATGTCCTG-3′ | Genome editing |
| sgRNA79-1 scarred allele PCR R | 5107 | 5′-GCTCATCTAATTCCCTGGCTTG-3′ | Genome editing |
| alt-sgRNA70-3 scarred allele PCR R | 6023 | 5′-GCCAGGTGCCCATCTGTGG-3′ | Genome editing |
| *Ube3a* exon 13 genomic PCR F | 4467 | 5′-GTTACCTACATCTCATACTTGCT-3′ | Control (genome editing) |
| *Ube3a* exon 13 genomic PCR R | 4468 | 5′-TGTGGTTCACTATCTTACAGCC-3′ | Control (genome editing) |
| *Snord107* genomic PCR F | 6028 | 5′-CAGAGACCTTGACTAGATTTATG-3′ | Test ddPCR |
| *Snord107* genomic PCR R | 6029 | 5′-TGAGGCACATTGACTAGATTTC-3′ | Test ddPCR |
| *Snord107* genomic Taqman probe | 6027 | /56-FAM/ACCTTGTCT/ZEN/GAACACAATGATTTCA/3IABkFQ/ | Test ddPCR probe |
| *Snurf-Snrpn* promoter F | 4678 | 5'-CTTTTGACAGGACATTGCAGTC-3' | Test ddPCR |
| *Snurf-Snrpn* promoter R | 4679 | 5'-CTCCATTGCGTTGCAAAATCTC-3' | Test ddPCR |
| *Mirh1* exon 1 F | 3493 | 5'-GATGAAATCCTGATCATCTGTG-3' | Test ddPCR |
| *Mirh1* exon 1 R | 3494 | 5'-CTCTCGTCGAGTCTCATCTG-3' | Test ddPCR |
| *Ube3a* exon 3 genomic PCR F | 6025 | 5′-TAGCAGCTAGACTGGTGTGAC-3′ | Control ddPCR |
| *Ube3a* exon 3 genomic PCR R | 6026 | 5′-CTTTTACAAGCTGTGGCCATTC-3′ | Control ddPCR |
| *Ube3a* exon 3 genomic Taqman probe | 5444 | /5HEX/CTGTTTTTA/ZEN/ATCAGTGACTCAAAGCT/3IABkFQ/ | Control ddPCR probe |
| F outer (un/me) *Snurf-Snrpn* /IC | 5037 | 5′-GTAGGAAtTtTGAAGATtAGATAGtTG-3′ | DNA methylation |
| R outer (un/me) *Snurf-Snrpn* /IC | 5038 | 5′-AAACCAAATACCCAACTAACCTTCC-3′ | DNA methylation |
| F un *Snurf-Snrpn* /IC | 5041 | 5′-GAtGtATGtGTAGGGAGtAGtAtG-3′ | DNA methylation |
| R un *Snurf-Snrpn* /IC | 5042 | 5′-CTACTCTCTACACTAAACTTACCA-3′ | DNA methylation |
| F me *Snurf-Snrpn* /IC | 5043 | 5′-CGtATGCGTAGGGAGtAGtACG-3′ | DNA methylation |

| | | | |
|---|---|---|---|
| R me *Snurf* -*Snrpn* /IC | 5044 | 5′-TACTCTCTACGCTAAACTTACCG-3′ | DNA methylation |
| *ψSnurf* PCR F | 5302 | 5′-TCAGACATCTTGTGAAGACCAATG-3′ | ψgene genomic analysis |
| *ψSnurf* PCR R | 5303 | 5′-CTTCCTCGCTCCATTGCGTTG-3′ | ψgene genomic analysis |
| sgRNA1 off-target F (chr 2) | RC0005 | 5'-TGAGAATGGCCCCCATAGGA-3' | Off target PCR |
| sgRNA1 off-target R (chr 2) | RC0006 | 5'-TTGCTGCCTTCAACCCAAGAT-3' | Off target PCR |
| sgRNA1 off-target F (chr 6) | RC0007 | 5'-TCTGCCTCCAACTTCCACAC-3' | Off target PCR |
| sgRNA1 off-target R (chr 6) | RC0008 | 5'-TAACTGCAGCTTGTCAGTGGA-3' | Off target PCR |
| sgRNA1 off-target F (chr 1) | RC0009 | 5'-AACTGCACGGTAGGGAATGG-3' | Off target PCR |
| sgRNA1 off-target R (chr 1) | RC0010 | 5'-ATGCAAGCATGGGCCTAGAG-3' | Off target PCR |
| sgRNA3 off-target F (chr 19) | RC0013 | 5'-TCAATTTGAAGCCCAAAGGTCAC-3' | Off target PCR |
| sgRNA3 off-target R (chr 19) | RC0014 | 5'-TGCTCTGGACAACCAAATGTACT-3' | Off target PCR |
| sgRNA3 off-target F (chr 6) | RC0015 | 5'-TTGAGGGGATACTCTGGGGT-3' | Off target PCR |
| sgRNA3 off-target R (chr 6) | RC0016 | 5'-TGTAGAGCTGCTGGGAACAC-3' | Off target PCR |
| sgRNA3 off-target F (chr 9) | RC0017 | 5'-TTTCTGGTTTCGCTAATGACAC-3' | Off target PCR |
| sgRNA3 off-target R (chr 9) | RC0018 | 5'-GTGCTCAGGGGGAGCTAATC-3' | Off target PCR |
| sgRNA70-3 off-target F (chr 1) | RC0021 | 5'-GCAGTGCCATGCTTCAGAAC-3' | Off target PCR |
| sgRNA70-3 off-target R (chr 1) | RC0022 | 5'-TTGTACAGGTGACCCTCTGC-3' | Off target PCR |
| sgRNA70-3 off-target F (chr 8) | RC0023 | 5'-AAGGGCTGCTCGTAAATCCC-3' | Off target PCR |
| sgRNA70-3 off-target R (chr 8) | RC0024 | 5'-GGCCAGAGGGAGGGAAGATA-3' | Off target PCR |
| sgRNA79-1 off-target F (chr 9) | RC0029 | 5'-ACCAATAGAGCCCAGTTCGC-3' | Off target PCR |
| sgRNA79-1 off-target R (chr 9) | RC0030 | 5'-AGTGGTAAGGCAACAGCCAAG-3' | Off target PCR |
| sgRNA79-1 off-target F (chr 3) | RC0031 | 5'-CCACACTACTTCTCAGGGTTG-3' | Off target PCR |
| sgRNA79-1 off-target R (chr 3) | RC0032 | 5'-GGTGAACTGTGGAATCCTGTAC-3' | Off target PCR |
| sgRNA79-1 off-target F (chr 2) | RC0033 | 5'-GCTGTGCTCCATTTTCCTAGC-3' | Off target PCR |
| sgRNA79-1 off-target R (chr 2) | RC0034 | 5'-CACGTGCTACCATAGCAACAG-3' | Off target PCR |

**Table S8. RT-PCR and RT-ddPCR primers.** Abbreviations: F, forward; R, reverse; RT-PCR, reverse transcription-PCR; RT-ddPCR, reverse transcription-droplet digital PCR.

| RT-PCR or RT-ddPCR Primer | Primer # | Primer Sequence |
|---|---|---|
| *Gapdh* exon 4 RT-PCR, RT-ddPCR F | 4833 | 5′-TGCACCACCAACTGCTTAGC-3′ |
| *Gapdh* exon 4/5 RT-PCR, RT-ddPCR R | 4834 | 5′-GGCATGGACTGTGGTCATGA-3′ |
| *Ins2* exon 1 RT-PCR | 4284 | 5′-CTAAGTGACCAGCTACAGTCG-3′ |
| *Ins2* exon 2 RT-PCR | 4291 | 5′-TCCACAGGGCCATGTTGGAAC-3′ |
| 5.8S RNA RT-PCR F | 4205 | 5′-CGACTCTTAGCGGTGGATCA-3′ |
| 5.8S RNA RT-PCR R | 4206 | 5′-GACGCTCAGACAGGCGTAG-3′ |
| *Mkrn3* RT-PCR F | 3617 | 5′-CTCCTCTGGCTTTGTCATTCCTA-3′ |
| *Mkrn3* RT-PCR R | 3498 | 5′-AGAATTCTCCAAATGGGCAGTG-3′ |
| *Magel2* RT-PCR F | 3497 | 5′-CCAACACTAAGCTGGAGTGCAC-3′ |
| *Magel2* RT-PCR R | 3616 | 5′-TGACGATGATGTAAGTGTGGGTTT-3′ |
| *Ndn* RT-PCR F | 3495 | 5′-GAAGAAGCACTCCACCTTCG-3′ |
| *Ndn* RT-PCR R | 3496 | 5′-TCCATGATCTGCATCTTGGTG-3′ |
| U1A RT-PCR F | 3488 | 5′-GAATGGTCAGCAAGGCAATGC-3′ |
| *Snurf* exon 1 RT-PCR F | 3490 | 5′-GACGGTTGGTTCTGAGGAGA-3′ |
| *Snurf* exon 2 RT-PCR R | 3489 | 5′-ACCTGGACCTCAATCTCTGG-3′ |
| *Snurf* exon 3 RT-PCR R | 4242 | 5′-GTCTCAGCTCTGCCTGGAAATC-3′ |
| *Snurf* exon 4 RT-PCR R | 5294 | 5′-GATTGCTGTTCCACAATAGCAG-3′ |
| *Snurf* exon 5 RT-PCR R | 5295 | 5′-ATGCTTGTCAAAAGCCTTAAAGG-3′ |
| *Snrpn* exon 8 RT-PCR F | 2825 | 5′-ACCTGTAGGCAGAGCAACC-3′ |
| *Snrpn* exon 10 RT-PCR R | 2826 | 5′-TCACAAGAAGCATTGTAGGG-3′ |
| *Snrpn* exon 8/9 RT-PCR F | 5191 | 5′-ACCTCCAGGCATTATGGCTC-3′ |
| *Snrpn* exon 10 RT-PCR R | 5192 | 5′-TCAACTGTATCTTAGGGTCTTG-3′ |
| *Snord107* RT-PCR F | 3499 | 5′-ATGATGACATGGGACCTTGTC-3′ |
| *Snord107* RT-PCR R | 3279 | 5′-GTGAGGCACATTGACTAGAT-3′ |
| *Snord107* RT-PCR R | 5971 | 5′-ATTTCAGAGTCACTCTAAGCTC-3′ |
| *Snord64* RT-PCR F | 3500 | 5′-AATGATGAGCTGTGTTTACTG-3′ |
| *Snord64* RT-PCR R | 3501 | 5′-CTTCAGAGTAATTATTTTGAGC-3′ |
| *Snord116* RT-PCR, RT-ddPCR F | 3502 | 5′-GGATCGATGATGATTTCCAATAAA-3′ |
| *Snord116* RT-PCR, RT-ddPCR R | 3503 | 5′-CTCAGTCACGATGATAGTGGC-3′ |
| *Ipw* RT-PCR F | 3491 | 5′-CATAGATGGTGCCACTTCTTCA-3′ |
| *Ipw* RT-PCR R | 3492 | 5′-TCTGAAGGTGGTGTCTGCTG-3′ |
| *Snord115* RT-PCR F | 3504 | 5′-GTCAATGATGACAACATTAAG-3′ |
| *Snord115* RT-PCR R | 3285 | 5′-GGCCTCAGCGTAATCCTATT-3′ |
| *Ube3a* exon 12 RT-PCR F | 4430 | 5′-CTGCAGTTTACAACAGGCACAG-3′ |
| *Ube3a* exon 13 RT-PCR R | 4192 | 5′-AGCAAGTATGAGATGTAGGTAAC-3′ |
| *Ube3a-ats* RT-PCR R (*Ube3a* int 12 F) | 4435 | 5′-AATAGGAGCATTTAACACTGAGC-3′ |
| *Xbp1* exon 3 RT-PCR F | 5677 | 5′-GTCTCAGAGGCAGAGTCCAAG-3′ |
| *Xbp1* exon 4 RT-PCR R | 5678 | 5′-GGCAACAGCGTCAGAATCCATG-3′ |
| *Gpi* exon 13/14 RT-PCR F | 5883 | 5′-CACCAAGGCACCAAGATGATAC-3′ |
| *Gpi* exon 14/15 RT-PCR R | 5884 | 5′-AGGAGGATCTTGTGATGCAGAC-3′ |
| *Hspa5* (Grp78) exon 8 RT-ddPCR F | 5726 | 5′-CTCCACAGCTTCTGATAATCAG-3′ |
| *Hspa5* (Grp78) exon 9 RT-ddPCR R | 5727 | 5′-GATTCCAGTCAGATCAAATGTAC-3′ |
| *Hsp90b1* (Grp94) exon 17 RT-ddPCR F | 5732 | 5′-GATGAAGAAGAGACAGACGCAG-3′ |
| *Hsp90b1* (Grp94) exon 18 RT-ddPCR R | 5733 | 5′-TTACAATTCATCCTTCTCTGTAGG-3′ |
| *Pdia4* (ERP72) exon 9 RT-ddPCR F | 5728 | 5′-GAGCCAGAGGAGTTTGATTCAG-3′ |
| *Pdia4* (ERP72) exon 10 RT-ddPCR R | 5729 | 5′-TGTTGTTCTTGGGAACTGGCTG-3′ |
| *Pdia6* (ERp5) exon 11/12 RT-ddPCR F | 5738 | 5′-GTTTCTCAGGGAACTGTCTTTCG-3′ |
| *Pdia6* (ERp5) exon 13 RT-ddPCR R | 5739 | 5′-ATCACTGAGGTCAATGTCGTCC-3′ |
| *Ppib* (Cyclophilin B) exon 4 RT-ddPCR F | 6256 | 5′-ACAGTCAAGACCTCCTGGCT-3′ |
| *Ppib* (Cyclophilin B) exon 5 RT-ddPCR R | 6257 | 5′-TCCACCTTCCGTACCACATC-3′ |
| *Creld2* exon 9 RT-ddPCR F | 5734 | 5′-GATGGCTTTGAGGAGACAGAAG-3′ |
| *Creld2* exon 10 RT-ddPCR R | 5735 | 5′-CTTCTGAATCCAGATGTCTATCAC-3′ |
| *Sdf2l1* exon 2 RT-ddPCR F | 5675 | 5′-AACCTGCACACGCACCACTTC-3′ |
| *Sdf2l1* exon 3 RT-ddPCR R | 5676 | 5′-AACATCGTACTGTCCACAGGTC-3′ |
| *Dnajb11* (ERdj3) exon 9 RT-ddPCR F | 5736 | 5′-TTGATAATCACCTTTGATGTGGAC-3′ |
| *Dnajb11* (ERdj3) exonx 10 RT-ddPCR R | 5737 | 5′-CCGTTGTATACCTTCTGCACTG-3′ |
| *Dnajc3* (P58[IPK]) exon 11 RT-ddPCR F | 6260 | 5′-GAAAAAGTTCATTGACATAGCAGC-3′ |
| *Dnajc3* (P58[IPK]) exon 12 RT-ddPCR R | 6261 | 5′-ACGGTCCGCCTGAGCTGAAG-3′ |
| *Hyou1* (ORP150, GRP170) exon 23 RT-ddPCR F | 5730 | 5′-GGAAAAGGTCATTCCACCTACAG-3′ |
| *Hyou1* (ORP150, GRP170) exon 26 RT-ddPCR R | 5731 | 5′-CTGTCTGCTCTGCCTGTTCAG-3′ |
| Rat *Ins1* (1) exon 2 RT-ddPCR F | 5499 | 5'-TGTGGTCCTCACCTGGTGGAG-3' |
| Rat *Ins1* (1) exon 2 RT-ddPCR F | 5500 | 5'-GGCGGGGAGTGGTGGACTCA-3' |
| Rat *Ins1* (2) exon 1 RT-ddPCR F | 5466 | 5'-AGTGACCAGCTACAATCATAG-3' |
| Rat *Ins1* (2) exon 2 RT-ddPCR R | 5467 | 5'-GACAAAAGCCTGGGCAGGCTT-3' |
| Rat *Ins2* (1) exon 1 RT-ddPCR F | 4284 | 5'-CTAAGTGACCAGCTACAGTCG-3' |
| Rat *Ins2* (1) exon 2 RT-ddPCR R | 6194 | 5'-ACAAAAGCCTGGGCAGGGCG-3' |
| Rat *Ins2* (2) exon 2 RT-ddPCR F | 5497 | 5'-TGTGGTTCTCACTTGGTGGAA-3' |
| Rat *Ins2* (2) exon 3 RT-ddPCR R | 5498 | 5'-TGGACAGGGTAGTGGTGGGC-3' |
| Mouse *Ins2* exon 1 RT-ddPCR F | 5479 | 5'-CCTAAGTGATCCGCTACAATC-3' |

| | | |
|---|---|---|
| Mouse *Ins2* exon 2 RT-ddPCR R | 6195 | 5'-ACAAAAGCCTGGGTGGGGTG-3' |
| *mCherry* (1) RT-PCR, 3' RT-ddPCR F | 4294 | 5'-GCCAAGCTGAAGGTGACCAAG-3' |
| *mCherry* (1) RT-PCR, 3' RT-ddPCR R | 4295 | 5'-GAAGGACAGCTTCAAGTAGTCG-3' |
| *mCherry* (2) RT-ddPCR F | 6232 | 5'-GAGCAGCAGCAATGGTGAGC-3' |
| *mCherry* (2) RT-ddPCR R | 4391 | 5'-GCCCTCGATCTCGAACTCGT-3' |
| Human *INS* (1) exon 2 RT-ddPCR F | 5489 | 5'-CATGGCCCTGTGGATGCGC-3' |
| Human *INS* (1) exon 2 RT-ddPCR R | 5478 | 5'-GTGTTGGTTCACAAAGGCTGC-3' |
| Human *INS* (2) exon 2 RT-ddPCR F | 5490 | 5'-GCAGCCTTTGTGAACCAACAC-3' |
| Human *INS* (2) exon 3 RT-ddPCR R | 5492 | 5'-TGTTCCACAATGCCACGCTTC-3' |
| *NeoR* RT-ddPCR F | 6202 | 5'-CCTTCTATCGCCTTCTTGACG-3' |
| *hGH* polyA RT-ddPCR R | 5291 | 5'-CACTGGAGTGGCAACTTCCAG-3' |
| *Npy* exon 2 RT-ddPCR F | 5679 | 5′-TACTCCGCTCTGCGACACTAC-3′ |
| *Npy* exon 4 RT-ddPCR R | 5680 | 5′-CACATGGAAGGGTCTTCAAGC-3′ |
| *Iapp* exon 2 RT-ddPCR F | 4614 | 5′-CAGGCTGCCAGCTGTTCTCC-3′ |
| *Iapp* exon 3 RT-ddPCR R | 4615 | 5′-ACATGTGGCTGTGTTGCACTTC-3′ |
| *Pcsk1* exon 12 RT-ddPCR F | 4745 | 5′-TAAGAATTGGGACTTCATGTCTG-3′ |
| *Pcsk1* exon 13 RT-ddPCR R | 4746 | 5′-ACACGAGGCTGCTTCATGTG-3′ |
| *Chgb* exon 3 RT-ddPCR F | 4585 | 5′-AACCATCACCCCTGAGTGCC-3′ |
| *Chgb* exon 4 RT-ddPCRR R | 5476 | 5′-GTCTCTTAGCAACCGTACTTC-3′ |
| *Cacna1a* (Cav2.1) exon 47 RT-ddPCR F | 5757 | 5′-CACAGACAGGGCAGTAGTTC-3′ |
| *Cacna1a* (Cav2.1) exon 48 RT-ddPCR R | 5758 | 5′-GGCGAGTAGGACACAAGCG-3′ |
| *Tmem176a* exon 6 RT-ddPCR F | 5786 | 5′-CTGGAAAAGATTCTTCACAAAGG-3′ |
| *Tmem176a* exon7 RT-ddPCR R | 5787 | 5′-CTGACGCTTCTTTCAGGCTTC-3′ |
| *Tmem176b* exon 7 RT-ddPCR F | 5788 | 5′-CACGGTTATCTGTATCCTGAAG-3′ |
| *Tmem176b* exon 8 RT-ddPCR R | 5789 | 5′-TGGGTTTGGACAGCTCATGGG-3′ |
| *Derl3* exon 5/6 RT-ddPCR F | 6283 | 5′-ATCACAGACCTGCTAGGGATC-3′ |
| *Derl3* exon 7 RT-ddPCR F | 6284 | 5′-TGAGGGTCATCTAGTAGTAGCT-3′ |
| *Mylip* exon 6 RT-ddPCR F | 6269 | 5′-TGCGAGGAGGAGATCAACTC-3′ |
| *Mylip* exon 7 RT-ddPCR F | 6270 | 5′-GCAGGTAGACGTGCTGGACA-3′ |
| *Atp10a* exon 21 RT-ddPCR F | 4807 | 5′-GCCTACTATGACTCCGATGTG-3′ |
| *Atp10a* exon21/22 RT-ddPCR F | 4808 | 5′-TTGAGCCAGGTCCAGGTTTTG-3′ |
| *Jph3* exon 4 RT-ddPCR F | 6289 | 5′-GAAGTCCCTGCCTGTTGCAC-3′ |
| *Jph3* exon 5 RT-ddPCR F | 6290 | 5′-CGACCAAGATGGGAGCTGAG-3′ |
| *Ndrg4* 3'-UTR RT-ddPCR F | 6285 | 5′-GAGTGGGAGTCAGACCAGTG-3′ |
| *Ndrg4* 3'-UTR RT-ddPCR F | 6286 | 5′-CAGAGAGCACCAGGAAAATGG-3′ |
| *Tap1* exon 8/9 RT-ddPCR F | 6277 | 5′-TCAGGGCTATGACACAGAGGT-3′ |
| *Tap1* exon 9 RT-ddPCR R | 6278 | 5′-AGCTGGTTGCCAGCATCCAG-3′ |
| *Tap2* exon 10 RT-ddPCR F | 6287 | 5′-CTGTGCAGACGACTTCATAGG-3′ |
| *Tap2* exon 11 RT-ddPCR R | 6288 | 5′-TAGCCTCATCCAGGATGAGGA-3′ |
| *Syt1* exon 9/10 RT-ddPCR F | 6262 | 5′-TGTGGGTGGCTTATCTGATCC-3′ |
| *Syt1* exon 10/11 RT-ddPCR R | 6263 | 5′-ACCACCACTTGCACTTTCTGG-3′ |
| *Robo2* exon 28 RT-ddPCR F | 5458 | 5′-GAATGCCAATGACCTTCTCGAC-3′ |
| *Robo2* exon 29 RT-ddPCR F | 5459 | 5′-TTCCATGAGTTTGGACCTGATG-3′ |
| *Id4* exon 1 RT-ddPCR F | 5454 | 5′-CTTACGGCGCTCAACACTGAC-3′ |
| *Id4* exon 2 RT-ddPCR F | 5455 | 5′-CCTGGCTCGCTGGTGCTCAG-3′ |
| *Stard10* exon 6 RT-ddPCR F | 6281 | 5′-AGAGCTGTGTCATCACCTACC-3′ |
| *Stard10* exon 7/8 RT-ddPCR F | 6282 | 5′-TACATCTTCTTCATGGCCTTGG-3′ |
| *Parm1* exon 2/3 RT-ddPCR R | 6267 | 5′-CCTTAAGTTCAGGCAGCATCG-3′ |
| *Parm1* exon 4 RT-ddPCR F | 6268 | 5′-CAGGACCCGTAGTCATGGTC-3′ |
| *Rph3al* (Noc2) exon 8 RT-ddPCR F | 5784 | 5′-CCATCTTTCTGGCAGCCAGAG-3′ |
| *Rph3al* (Noc2) exon 9 RT-ddPCR R | 5785 | 5′-GTTGCCCAGGTATGCCTTTTGC-3′ |
| *Bsn* exon 9 RT-ddPCR F | 5776 | 5′-ATTTTCTAAGATCCTCCCTGGTG-3′ |
| *Bsn* exon 10 RT-ddPCR R | 5777 | 5′-TCTGGACACAATCACCAGAATG-3′ |
| *Pclo* exon 15 RT-ddPCR F | 5778 | 5′-CTACTCTGACCCGTTTGTTAAG-3′ |
| *Pclo* exon 18 RT-ddPCR F | 5779 | 5′-CCTCCAAGGTCTTCTTCATGAG-3′ |
| *Jph4* exon 4 RT-ddPCR F | 6258 | 5′-CTGGGTGCCTGACAGAAGAG-3′ |
| *Jph4* exon 5 RT-ddPCR R | 6259 | 5′-TGAGGAGCTGGGAGAACAGG-3′ |
| *Cacna1d* (Cav1.3) exon 38 RT-ddPCR F | 5782 | 5'-ACGGCTCTCAAGATCAAGACTG-3' |
| *Cacna1d* (Cav1.3) exon 40 RT-ddPCR R | 5783 | 5'-TCAGGAAAGTGGCATAGAACTTC-3' |
| *Srrm1* ex 11 RT-ddPCR F | 6047 | 5'-CAGAACCAGCAGTCTTCATCTG-3' |
| *Srrm1* ex 12 RT-ddPCR R | 6048 | 5'-CGCTTTCGAGGTGATGGAGAG-3' |
| *Srrm2* ex 12 RT-ddPCR F | 6006 | 5'-TGCTAAGCCTGGCCCTCAGG-3' |
| *Srrm2* ex 14/15 RT-ddPCR R | 6007 | 5'-TTATGGAGACCTGGAGGAGCG-3' |
| *Sim1* exon 10 RT-PCR F | 5428 | 5′-AACCTATGAGAACAGCATGCC-3′ |
| *Sim1* exon 11 RT-PCR R | 5430 | 5′-AGCTGACCACACTATCTTCATC-3′ |
| *Mon2* exon 2 RT-PCR F | 5579 | 5′-ACAATAGCTGCAAGAAACACCG-3′ |
| *Mon2* exon 3 RT-PCR R | 5580 | 5′-GCTAAACACAGCTGAGTGACC-3′ |
| *ψSnurf* RT-PCR F | 5300 | 5′-CTTGGTTCTGAGGAGGGATTTG-3′ |

**Table S9. Antibodies used in this study.**

| Antibody | Type | Supplier | Catalog No. or Reference | Dilution |
|---|---|---|---|---|
| Anti-α-tubulin | rabbit monoclonal | Cell Signaling Technology, Danvers, MA | # 2125 (11H10) | 1:1000 in 5% milk TBST |
| Anti-GPI | rabbit polyclonal | Sigma-Aldrich, Saint Louis, MO | HPA024305 | 1:500 in 5% milk TBST |
| Anti-GAPDH | mouse monoclonal | Sigma-Aldrich, Saint Louis, MO | G8795 (GAPDH-71.1) | 1:5000 in 5% milk TBST |
| Anti-SmB/B'/Anti-SmN | rabbit polyclonal | Sigma-Aldrich, Saint Louis, MO | HPA003482 | 1:1000 in 5% milk TBST |
| Anti-KDEL | mouse monoclonal | Enzo Life Sciences, Farmingdale, NY | # ADI-SPA-827 (10C3) | 1:1000 in 5% milk TBST |
| Anti-insulin | mouse monoclonal | Cell Signaling Technology, Danvers, MA | # 8138 (L6B10) | 1:1000 in 5% BSA TBST |
| Anti-mCherry | rabbit polyclonal | Millipore Sigma, Burlington, MA | AB356482 | 1:1000 in 5% milk TBST |
| Anti-phosphoEIF2α (Ser51) | rabbit polyclonal | Cell Signaling Technology, Danvers, MA | # 9721 | 1:100 in 5% milk TBST |
| Anti-EIF2α | rabbit polyclonal | Cell Signaling Technology, Danvers, MA | # 9722 | 1:100 in 5% milk TBST |
| Anti-ATF6 | rabbit polyclonal | Ronald C. Wek | Teske et al. 2011 | 1:500 in 5% milk TBST |
| Anti-mouse IgG (HRP) | Horse | Cell Signaling Technology, Danvers, MA | # 7076S | 1:3000 in 5% milk TBST |
| Anti-rabbit IgG (HRP) | Goat | Santa Cruz Biotechnology, Dallas, TX | # 2301 | 1:3000-1:5000 5% milk TBST |

| |
|---|
| Dilution of antibodies in TBST (20mM Tris pH 7.4, 150 mM NaCl and 0.1% Tween 20) included addition of sodium azide (0.05%) to prevent bacterial contamination |
| After transfer membranes were stained with Ponceau S for 10 minutes, then washed with TBST 3x for 3 min; then blocked with 5% milk TBST for 1 hr at room temp |
| After blocking antibodies were added at noted dilution and incubated overnight on a shaker at 4°C. |
| For 5% BSA TBST diluted antibodies a 10 minute blocking step in 5% BSA TBST was included following the first block in 5% milk |
| After primary antibody incubation membranes were washed 3x for 15 min with TBST followed by addition of secondary HRP-conjugated antibody for 1 hr at room temp |
| Membranes were washed an additional 3x for 15 min with TBST prior to ECL detection |
